# Impacts of global food supply on biodiversity via land use and climate change

**DOI:** 10.1101/2023.05.03.539201

**Authors:** Elizabeth H Boakes, Carole Dalin, Adrienne Etard, Tim Newbold

## Abstract

Land-use change is currently the greatest driver of biodiversity change, with climate change predicted to match or surpass its impacts by mid-century. The global food system is a key driver of both these anthropogenic pressures, thus the development of sustainable food systems will be critical to halting and reversing biodiversity loss. Previous studies of the biodiversity footprint of food tend to focus on land use alone. We use the multi-regional input-output model EXIOBASE to estimate the impacts of biodiversity embedded within the global food system. We build on prior analyses, calculating the impacts of both agricultural land-use and greenhouse gas (GHG) emission footprints for the same two metrics of biodiversity: local species richness and rarity-weighted species richness. Our biodiversity models capture regional variation in the sensitivity of biodiversity both to land-use differences and to climate change. We find that the footprint of land area does not capture the biodiversity impact embedded within trade that is provided by our metric of land-driven species richness change, and that our metric of rarity-weighted richness places a greater emphasis on the biodiversity costs in Central and South America. We find that methane emissions are responsible for 70% of the overall GHG-driven biodiversity footprint and that, in several regions, emissions from a single year’s food production cause biodiversity loss equivalent to 2% or more of that region’s total historic land use. The measures we present are simple to calculate and could be incorporated into decision making and environmental impact assessments by governments and businesses.

## 1. Introduction

Anthropogenic pressures continue to drive biodiversity loss despite increasing conservation efforts^1^. Land use is currently the greatest driver of biodiversity change^1,2^, and has thus been the focus of much previous conservation research. However, the impacts of climate change on biodiversity are expected to increase considerably^3^, and, by 2070, climate may match or surpass land use as the greatest driver of biodiversity change^4,5^. Agricultural land used for food production covers an estimated 38-55% of Earth’s habitable land^6,7^, while the global food system is responsible for 21-37% of anthropogenic greenhouse gas (GHG) emissions^8–10^. The development of sustainable food systems will be critical in halting/reversing land-driven^11^. and climate-driven biodiversity loss^12^.

Increasingly, food grown in one country is traded internationally to satisfy demand elsewhere^13–15^. The associated biodiversity impact of this international consumption is driven by upstream economic activities that are often geographically distant from the locations of biodiversity loss^16^. Reducing the environmental impacts of food therefore requires both supply-side and demand-side changes^16^.

Understanding the separate contributions of agricultural land use and greenhouse gas emissions to biodiversity loss is an important step toward developing a sustainable international trade in food. A strategy which considers land use alone might well differ from one that incorporates GHG-driven biodiversity loss. Importantly, the land and GHG footprint per tonne of crops arising from N_2_O tend to be negatively correlated – intensive farming uses less land than extensive farming, but has higher GHG emissions (N_2_O) arising from higher fertilizer use^17^. Moreover, land use and climate differ in their spatial impacts on biodiversity loss. Land use affects biodiversity that is local to the land conversion. In contrast, climate change affects biodiversity globally, no matter the location of the emissions source.

Environmentally-extended multi-regional input-output models (EEMRIOs) are used to link downstream environmental impacts to upstream drivers, allowing the ‘footprints’ of commodities to be followed back through complex supply chains^18^. The footprint of a hamburger, for example, would contain impacts not only from cattle but from cattle feed, fertilizer, machinery, water, packaging, fuel etc.^19^. Summing the impacts associated with each upstream product would, in theory, give the hamburger’s total footprint. However, in reality this simple addition process is impossible given the complications of tracking supply chains, double counting of recycled products, and infinite loops, for example electricity production requiring water which requires electricity to pump it. EEMRIOs resolve these problems by using input-output tables to infer production recipes, allocating environmental costs to sectors to avoid double counting and approximating infinite sums with the Leontief inverse matrix^19^. Regional production data and their associated environmental costs are combined with information on international trade, allowing the calculation of a variety of consumption footprints, e.g. land use^20^, GHG emissions^21^ and water^22^.

EEMRIO models have been used to follow the effects of international trade along supply chains, to estimate biodiversity footprints. Lenzen et al.^23^ performed the first global biodiversity footprint analysis, using the IUCN Red List to count species’ threats within trade regions. This method assumes that species are equally threatened across their ranges, and excludes non-threatened species. Other methods use a similar philosophy to calculate footprints based on biodiversity threat hotspots^24^, bird ranges and the number of individual birds lost from an area^16^. An alternative method, derived from land-use data, uses the countryside species-area relationship to estimate the potential number of extinctions caused by land conversion and international trade^15,25,26^. Since many of the extinctions have yet to be realized, it is unclear how these extinctions would be allocated to different actors across time^16^. Wilting et al.^27^ take a step further, deriving biodiversity footprints from GHG emissions as well as land use, using the biodiversity metric Mean Species Abundance (although climate impacts are based on model predictions of changes in species richness, so land-use and climate impacts are not entirely comparable). The GLOBIO biodiversity model^28^, on which Wilting et al.’s^27^ analysis was based, assumes that effects of land use and climate change are even across all terrestrial regions; in reality, biodiversity tends to be more sensitive to land use and climate change in tropical regions^29,30^.

We build on these prior analyses, introducing three novel aspects. (i) We calculate the biodiversity impacts of agricultural land use and GHG-emission footprints using models that directly output metrics of terrestrial biodiversity change in the same units, allowing the drivers’ impacts to be compared and splitting emissions into carbon dioxide (CO_2_), methane (CH_4_) and nitrous dioxide (N_2_O). (ii) We consider change in local rarity-weighted species richness relative to an unimpacted baseline in addition to local species richness. Species richness, although easy to measure, captures only one of the many dimensions of biodiversity, and does not always decline with global biodiversity loss^31^. Rarity-weighted richness gives greater weight to species with small geographic range size (range size correlates with species extinction risks^32^) and so declines if rare species are replaced by more common ones. (iii) We use biodiversity models that allow us for the first time to capture regional variation in the sensitivity of biodiversity both to land-use differences and to climate change^29^. We base our biodiversity metrics on local measures of biodiversity in the relevant agricultural areas as opposed to a value averaged across an entire trade region, meaning that we account for the wide variation in species richness that occurs within regions.

We examine the international production and consumption footprints of food-related commodities in terms of: a) land area; b) species richness (land-driven and GHG-driven); and c) rarity-weighted species richness (land-driven and GHG-driven). We calculate footprints for 33 food-related products that span food’s journey from field to farm gate to household to waste in 49 regions (44 countries and 5 “rest of world” regions). We ask whether the different footprints would give the same conclusions with respect to sustainable production, consumption and trade. We identify the regions and food groups with the highest biodiversity footprints, examining the contributions of land use and GHG emissions to these footprints. We explore biodiversity footprints per km^2^ and per capita for each region and look at the proportion of regions’ footprints which are imported. Our analysis provides a detailed examination into the pathways by which regions’ consumption of particular food-related products affects biodiversity worldwide, giving insight into the trade-offs between land use and GHG emissions, and into priorities for demand-side changes.

## 2. Methods

### 2.1 EEMRIO analysis

Input-output analysis is a top-down approach that uses sectoral transactions data (either in financial terms or units of product) to account for the complex interdependencies of industries. An input-output table can be environmentally extended by adding information on exchanges with the environment, e.g. GHG emissions or land use, by each industry sector. EEMRIO analysis traces the production and supply chain of traded goods and services and their associated materials back to the source of primary extraction, thus capturing the direct and indirect environmental pressures associated with a country’s final consumption^19^. Full details of the calculations underlying EEMRIO analysis can be found in Kitzes^19^ and Miller and Blair^18^.

We used the standard environmentally-extended Leontief model to calculate the effects of consumption on biodiversity loss

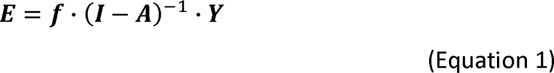

Where, for *i* regions and *m* economic sectors:

***E*** is a (1 × *i*) matrix, representing the environmental impacts associated with the final demand of each region (CO_2_, CH_4_ and N_2_O emissions, in kg, and agricultural area, in km^2^).

***f*** is the (1 × *im*) direct intensity vector, representing the environmental pressures (e.g. area of land, mass of CO_2_ emissions) associated with one unit (€1M) of production for each product sector in each region.

***I*** is the (*im* × *im*) identity matrix.

***A*** is the (*im* × *im*) technical coefficient matrix which gives the amount of input (€1M) that every sector must receive from every other sector in order to produce one unit (€1M) of output.

***Y*** is the (*im* × *i*) matrix of final demand given in monetary terms (€1M).

The direct intensity vector ***f*** allows the MRIO to be extended to include environmental costs of production and consumption. Sections 2.2 and 2.3 describe our environmental ‘characterisation factors’ based on species richness, rarity-weighted richness, land use and GHG emissions. The vector relating to land use is a sparse vector populated in the entries for production activities that involve use of cropland or pasture.

We use the EEMRIO database EXIOBASE3.8.1 (product by product version)^33^, because of its superior balance of sectoral and regional disaggregation relative to other MRIOs. EXIOBASE3 provides a harmonized time-series of MRIO tables and environmental extensions over the period 1995-2022, covering 200 products, 44 countries (EU countries and other major economies), plus 5 ‘Rest of World’ (RoW) regions. We chose 2011 as our year for analysis since land use data are available up to 2011 only and spatial crop data were not available after this date (see below).

Our analysis focuses on the production and consumption of the 33 product-sectors associated with food (Supplementary Table 1) in the year 2011, including fertiliser production and food waste processing. ‘Hotel and restaurant services’ was excluded since it encompasses energy and materials, for example, as well as food. Full details of EXIOBASE’s land-use and emissions accounts are given in Stadler et al.^33^. Our land-driven biodiversity footprints are based on agricultural land, and exclude impacts of forestry, infrastructure or other uses. Emissions from crops derive from nitrogen and phosphorous fertilisers, while emissions from livestock derive from the animals, the manure excreted and the cultivation of feed crops. Impacts of fertiliser refer to the production of the fertiliser only and impacts of food waste do not include the production impacts of the wasted food. For simplicity, we regard land use as a one-off cost that results in a change in biodiversity and we assume that once land has been converted, it can be used repeatedly without the biodiversity cost increasing. In contrast, GHG-driven biodiversity loss accrues year on year. GHG emissions that are released during land conversion are not considered, because, without detailed land-history knowledge, we cannot estimate the proportion of emissions that have dissipated since conversion, nor apportion food-production emissions across years. To put this gap in our coverage of GHGs into context, direct emissions from agriculture contribute 5.1-6.1 Pg CO_2_-eq,/yr while the clearing of native land for agriculture contributes around 5.9 (SD 2.9) Pg CO_2_-eq/yr^34^.

### 2.2 Characterisation factors for land-driven biodiversity impacts

Maps for a) the 49 EXIOBASE trade regions, b) the six biomes used in Newbold et al.^29^ (but here also including boreal forest within the temperate-forest grouping), and c) the 15 EXIOBASE agricultural land-use categories were identified, and map masks for each unique 3-way combination were generated. Shapefiles mapping the borders of individual countries^35^ were aggregated and rasterised to match EXIOBASE’s 49 regions. Biomes were obtained from The Nature Conservancy^36^ and were aggregated into tropical forest, temperate/boreal forest, tropical grassland, temperate/montane grassland, Mediterranean, and drylands (tundra, mangroves, flooded grasslands, inland water and rock & ice were excluded; Supplementary Table 2). Cropland data were obtained from the Spatial Production Allocation Model (SPAM)^37^ and from EarthStat^38^ and pasture data from EarthStat^39^. All spatial data were reprojected to an equal-area Behrmann grid. A summary schematic is given in Figure 1a and further details of the preparation of datasets are given in the Supplementary Information.

**Figure 1.**
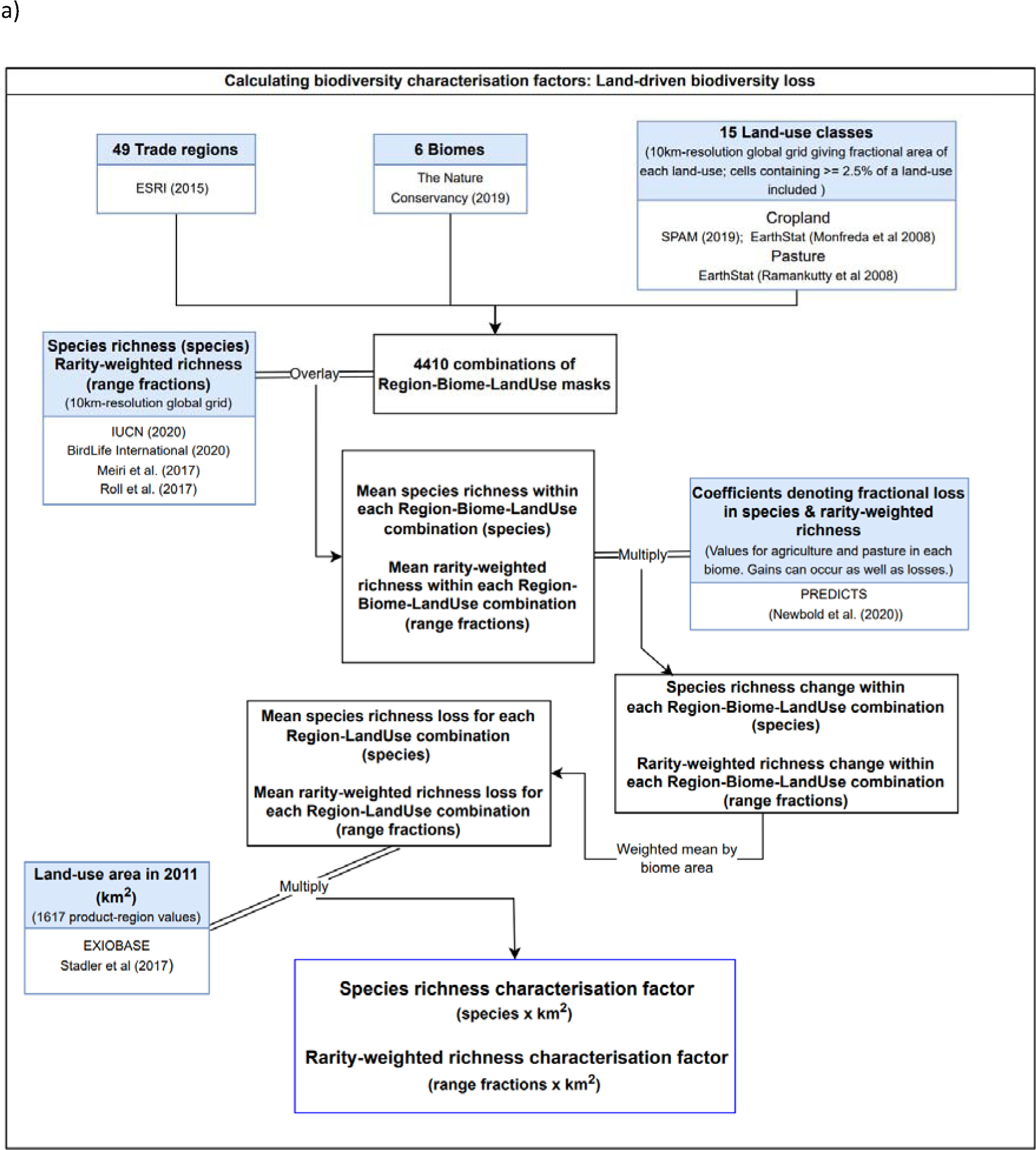
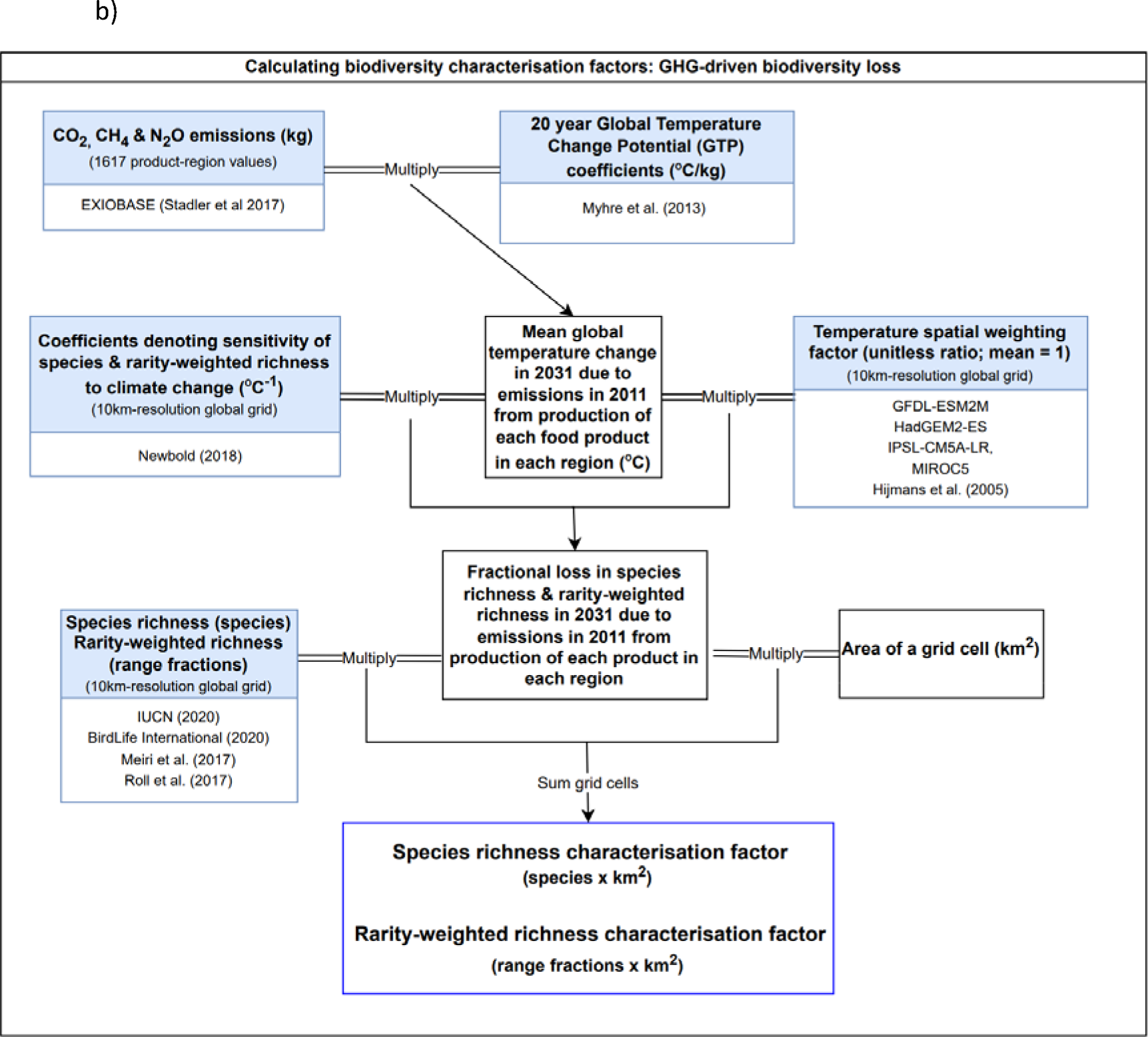
Schematics summarising the characterisation factor calculations. Characterisation factors for a) land-driven biodiversity loss and b) GHG-driven biodiversity loss. The methods allow for biodiversity gain which would be represented as a negative loss.

Mapped spatial estimates of species richness and rarity-weighted richness for terrestrial vertebrates were obtained by stacking species geographical distributions at a 10-km equal-area resolution. Extent-of-occurrence distribution maps were obtained from the IUCN^40^, BirdLife International^41^, Meiri et al.^42^, and Roll et al.^43^. We selected only areas where species are extant or probably extant, and resident during the breeding season. As in Etard et al.^44^, we further excluded areas outside the known elevational limits of species^40^. Species richness was calculated as the number of species occurring within a grid cell. Rarity-weighted species richness weights were calculated as the inverse of a species’ estimated range size, and rarity-weighted richness as the sum of these weights across species occurring within a grid cell. Mean species richness and rarity-weighted richness were calculated for each land-use type within each biome within each trade region.

We estimated the sensitivity of biodiversity to land use using models of the PREDICTS database^45,46^, following the methods given in Newbold et al.^29^ (see Supplementary Information). The PREDICTS database contains 3.25 million samples of biodiversity in six different land uses from 26, 114 locations and 47,044 different species. For the purposes of this study, we modelled biodiversity differences among four land-use classes: primary vegetation, secondary vegetation, agriculture (including both plantations and herbaceous cropland) and pasture, allowing the response of biodiversity (i.e. species richness and rarity-weighted richness) to these land-use categories to vary among the six broad biome groupings listed above. For further details, see the Supplementary Information.

Characterisation factors for species richness were calculated as follows. We used the species richness map to estimate the mean species richness *S_i,j,k_* that would occur in undisturbed land across the area covered by each EXIOBASE land-use type *i*, within each biome *j* and each EXIOBASE trade region *k*. The change in species richness Δ*S_i,j,k_* due to land-use was calculated as:

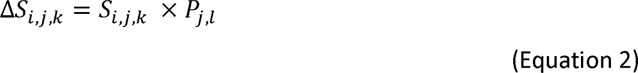

where *P_j,l_* is the proportional change in species richness in PREDICTS land-use type *l* and biome *j*, compared to primary vegetation in that biome (see Supplementary Table 3 for the correspondence between EXIOBASE and PREDICTS land-use types, and Supplementary Tables 4 and 5 for values of *P_j,l_*). The change in species richness Δ*S_i,k_* within each EXIOBASE land-use type *i* and each region *k* was calculated as:

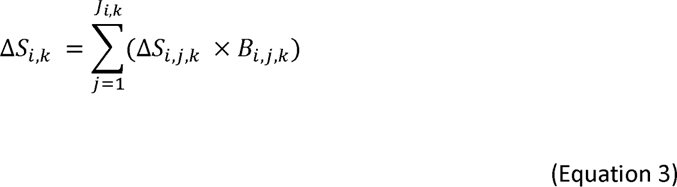

where *J_I,k_* is the number of biomes covered by the EXIOBASE land use type *i* within the region *k* and *B_i,j,k_* is the proportion of the EXIOBASE land-use type *i* within the region *k* that is covered by the biome *j*. The characterisation factor *CF_i,k_* was then calculated as:

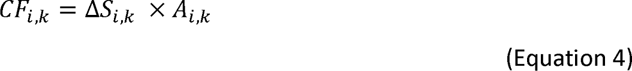

where *A_i,k_* is the area of agricultural land used to produce €1M of product in land-use type *i* in region *k*). Characterisation factors for rarity-weighted richness were calculated using the same method, but using the map of rarity-weighted species richness, and corresponding modelled estimates of land-use sensitivity (for characterisation factor values, see Supplementary Tables 6 and 7).

The characterisation factors for species richness have units of number of species × km^2^ and can be thought of as the count of the species lost, with this loss extending over a certain area (i.e., the area of land use given in EXIOBASE). Rarity-weighted richness characterisation factors are less intuitive, but can be thought of as having a unit of species range fractions × km^2^.

### 2.3 Characterisation factors for GHG-driven biodiversity change

We calculated GHG-driven biodiversity change in a three-step process. First, we calculated the GHG emissions (split into carbon dioxide (CO_2_), methane (CH_4_) and nitrous oxide (N_2_O)) associated with each region’s production of the 33 different food-related products in 2011, using EXIOBASE. Second, we calculated the warming that would result from these emissions. Third, we calculated the biodiversity change that would result from this warming, allowing for non-uniformity of warming across the globe. A summary schematic is given in Figure 1b.

EXIOBASE gives emissions for six GHGs: carbon dioxide, methane, nitrous oxide, and three fluorinated gases. We calculated footprints using the first three gases only since their contribution to warming via food-related emissions is much greater than that of the fluorinated gases. We summed methane emissions over the four categories given by EXIOBASE (‘Combustion – air’, ‘non-combustion’, ‘agriculture’ and ‘waste’). Footprints were calculated using the environmentally-extended Leontief model described in section 2.1.

We calculated the increase in global surface temperature due to GHG emissions using the Global Temperature Change Potential (GTP), a metric designed to be an intuitive measure of climate response^47^. The GTP is defined as the ratio between the global mean surface temperature change at a given future time horizon following an emission (pulse or sustained) of a compound relative to a reference gas^47^. We chose a 20-year time horizon, to capture warming due to the relatively short-lived methane emissions, and to represent a time that is tangible to today’s policy and decision-makers. GTP values in units of degrees of warming (°C) per kilogram of emissions were taken from the AR5 IPCC report^48^, Table 8.A.1 (CO 6.84 x 10^-16^ °C /kg, CH 4.62 x 10^-14^ °C /kg, N O 1.89 x 10^-13^ °C /kg). The warming in °C that would result by 2031 Δ*T_p,k_* due to the emissions *Eg,p,k* associated with each gas *g* due to the production of €1M of each product *p* in each region *k* was calculated as:

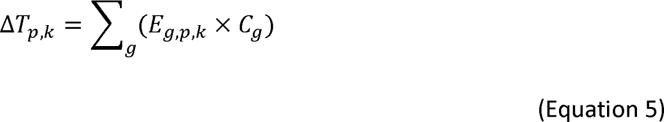

where *C_g_* is the GTP value for each gas *g*.

We estimated the sensitivity of biodiversity to climate change based on future projections of changes in the distributions of terrestrial vertebrates under the Representative Concentration Pathways (RCP) climate-change scenarios^5^. It is necessary to use future projections rather than observed responses of species to climate change, because we do not yet have enough data for a wide range of species from locations that have experienced extensive historical climate change^29^. Full details are given in the Supplementary Information.

To account for the non-uniformity of warming across the globe, we calculated a grid of spatial weighting factors to up or down weight temperature change depending on the difference between the projected local and mean global temperature change. The projected temperature change Δ*T_x_* in each cell *x* between 2011 and 2031 under RCP8.5 was calculated by averaging the projections of four different climate models (GFDL-ESM2M, HadGEM2-ES, IPSL-CM5A-LR, MIROC5^49^). The temperature weighting factor *F_x_* for each cell *x* was calculated as

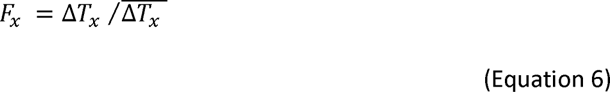

where 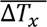 is the mean global temperature change. The fractional loss in species richness Δ*S_x,p,k_* in 2031 due to Δ*T_p,k_*, the temperature increase from emissions from food production of each product *p* in each region *k,* was calculated as:

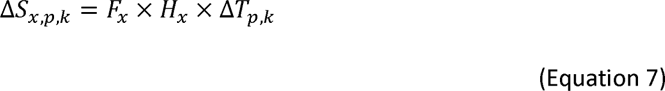

where *H_x_* is the grid estimating sensitivity of species richness to climate change.

This fractional loss in species richness Δ*S_x,p,k_* was then multiplied by our species richness grid and by the area of each grid cell to give a characterisation factor for GHG-driven biodiversity loss, *CF_p,k_*, with a unit of species × km^2^ that is comparable to our metric for land-driven biodiversity loss. This characterisation factor was calculated as:

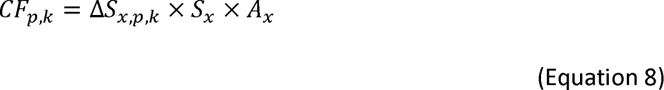

where *S_x_* is the species richness in undisturbed vegetation under a natural climate in grid cell *x* and *A_x_* is the area of grid cell *x*. The same basic method was used to calculate the characterisation factors for change in rarity-weighted richness (see Supplementary Information for details). The GHG-driven characterisation factors for change in species and rarity-weighted richness are directly correlated (for characterisation factor values, see Supplementary Tables 8 and 9.)

Global maps illustrating the proportional richness changes that would occur due to land-driven and GHG-driven processes are given in Supplementary Figure 3. Our metrics of land-driven and GHG-driven biodiversity loss are modelled and presented in the same units, and hence are comparable. However, there are differences in the way that the two drivers impact biodiversity, and in the methods we used to model these impacts. We consider land use as a one-off cost that has already occurred, but GHG emissions as repeated annual costs that occur 20 years after emission. We would expect the biodiversity loss caused by a single year of emissions to be much lower than that caused by the one-off cost of total land conversion. We calculate the ratio of land-driven to GHG-driven biodiversity loss for products and regions. We can crudely approximate a ratio of 100, for example, to mean that within a century, assuming the same emissions repeat annually, the global biodiversity loss caused by emissions from a region’s food production will equal the biodiversity loss due to total land conversion in that region. In reality, this may be an underestimate since we do not consider the emissions released by land conversion.

### 2.4 Per land area and per capita footprints

The regions within EXIOBASE have unequal land areas and human populations. To make a fairer comparison of footprints between regions, we converted production footprints to footprint per km^2^ and consumption footprints to footprint per capita. Per-km^2^ footprints were calculated using EXIOBASE trade regions’ areas (Supplementary Table 10), which were calculated in R from ESRI country shapefiles^35^. Per-capita footprints were calculated using population data for the year 2011 for the EXIOBASE trade regions (Supplementary Table 11), which were obtained from CountryEconomy.com^50^ (Taiwan) and the World Bank^51^ (all other regions).

## 3. Results

### 3.1 Regional production and consumption-based footprints for total food

For all footprints, regions with a high production-based footprint tend to have a high consumption-based footprint. Our land-driven biodiversity footprints show a different picture from a simple land-area footprint. Important information is missed if the area of agricultural land alone is used as a proxy for the impact of food production on biodiversity (Figure 2). Differences between land-associated footprints are driven in part by differences in species richness between regions’ agricultural areas but also by our characterisation factors allowing for biome-specific sensitivity to land use. Furthermore, footprints also vary according to the measure of biodiversity used. The regions with the greatest land-area footprints in 2011 were Rest of World (RoW) Africa, China, and RoW Asia & Pacific (Figure 2a). However, while RoW Africa also has the highest land-driven species richness footprint, Brazil and RoW Central & South America (RoW CS America) have the second and third highest species richness footprints, respectively (Figure 2b). RoW CS America hosts many narrow-ranged species, which is reflected in its high rarity-weighted richness footprint (Figure 2c). Decisions regarding land use and food production should therefore consider not only the land area involved but also the land-driven loss of different facets of biodiversity (see Supplementary Figure 5 and Supplementary Table 12 for the footprints of individual products within regions).

**Figure 2.**
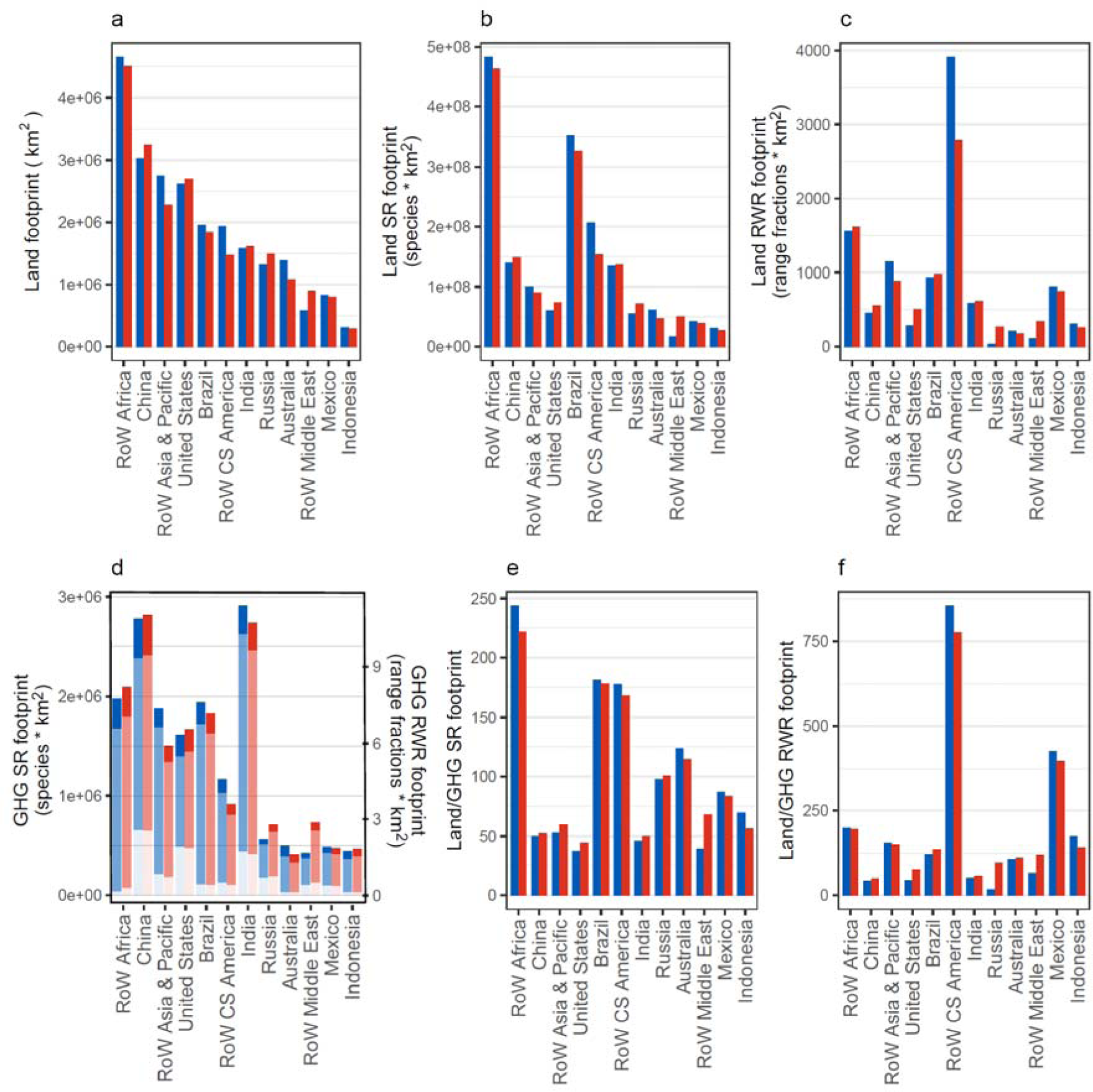
Total production-based (blue bars) and consumption-based (red bars) footprints of food-related products. Footprints relate to a) land area, b) land-driven species richness (SR) loss, c) land-driven rarity-weighted richness (RWR) loss, d) GHG-driven biodiversity loss split by emissions type: carbon dioxide (dark blue/red), methane (mid blue/red), nitrous oxide (light blue/red) (right-hand axis – species richness; left hand axis – rarity-weighted richness), e) the ratio of land-driven species richness loss to total GHG-driven species richness loss and f) the ratio of land-driven rarity-weighted richness loss to total GHG-driven rarity-weighted richness loss. Regions which are in the highest ten for one or more footprints are shown.

India and China are the regions responsible for the highest GHG-driven biodiversity loss, followed by RoW Africa (Figure 2). However, compared to India and China, RoW Africa has a much lower GHG-driven footprint relative to its land-driven biodiversity footprint. We expected the GHG-driven footprint to be much lower than the land-driven footprint since it relates to the global biodiversity loss caused by a single year of emissions whereas our land-driven footprint relates to the historic conversion of all agricultural land used in 2011. The ratio of land-driven to GHG-driven biodiversity loss varied by region from 16 for rarity-weighted richness production footprint in Russia to 855 for production in RoW C&S America, with several regions, including China, India and RoW Asia, having ratios around 50. Finding ratios of 50 or lower is concerning as it shows that direct emissions from a single year of a region’s food production will cause biodiversity loss equivalent to 2% or more of the biodiversity loss caused by that region’s total historic land use. Furthermore, we substantially underestimate biodiversity losses from GHG emissions since our analysis does not include emissions from land clearance.

Some regions are net importers of land-driven biodiversity loss but net exporters of GHG-driven biodiversity loss, e.g. India, or *vice versa*, e.g. Indonesia. RoW Asia & Pacific, RoW CS America, Australia and Mexico are all net exporters of both land-driven and GHG-driven biodiversity meaning their international exports are harming both their domestic biodiversity and, via climate change, global biodiversity. China, the United States, Russia and RoW Middle East are net importers of both footprints (Figure 2). The top ten footprints stem from regions with very large land areas and/or populations and, aside from Russia, do not include regions in continental Europe.

Methane emissions account for 70% of the total GHG-driven biodiversity footprint from food-related products, compared to 42% of the GHG-driven footprint of all EXIOBASE’s products. Carbon dioxide contributed to 18% of food’s total footprint versus 54% of the footprints of all products and nitrous oxide contributed to 12% of food’s footprint and 4% of the footprints of all products. Food-related products account for 23% of the total emissions in EXIOBASE and the total GHG-driven footprint of all food-related products was approximately 1% of the land-driven richness footprints.

### 3.2 Production-based footprints for world regions aggregated by food-related sector

Production-based footprints vary considerably among food-related groups and, within food-related groups, among world regions (Figure 3). As for total food footprints, land-driven biodiversity footprints do not mirror land-area footprints, and the relative size of biodiversity footprints is affected by the richness metric. Using animal-derived products as an example, Asia & Pacific has a land footprint 60% higher than CS America’s (and Africa’s, Figure 3a), but its land-driven species-richness footprint is less than half (Figure 3b), and its rarity-weighted richness footprint less than a third that of CS America (Figure 3c). This result reflects Asia & Pacific’s lower average natural species richness compared to CS America’s and the lower biodiversity sensitivity of some of its biomes to land use.

**Figure 3.**
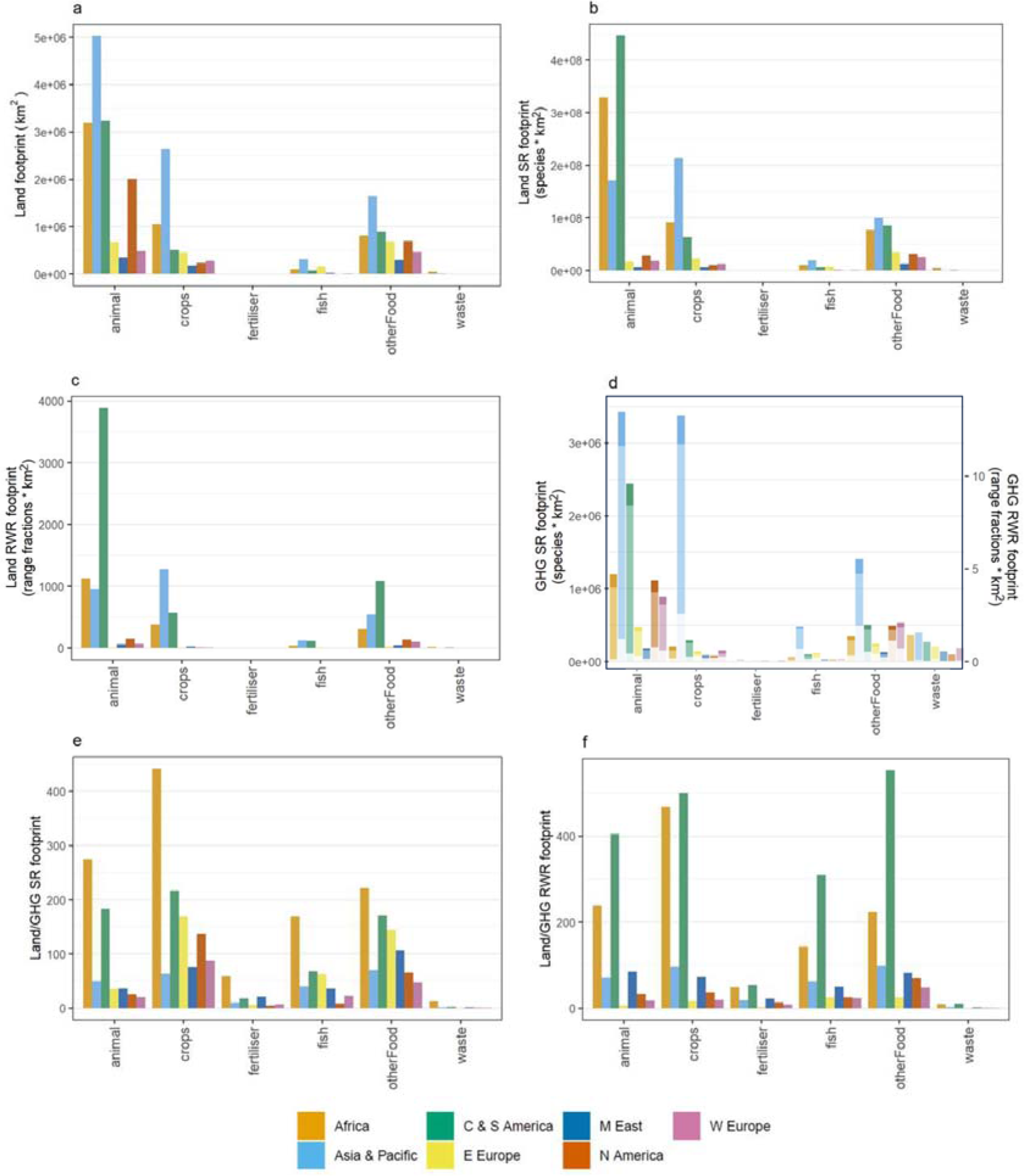
Production-based footprints of aggregated food-related groups within world regions. Footprints relate to a) land area, b) land-driven species richness (SR) loss, c) land-driven rarity-weighted richness (RWR) loss, d) GHG-driven biodiversity loss split by emissions type: carbon dioxide (dark shade), methane (mid shade), nitrous oxide (light shade) (right-hand axis – species richness; left-hand axis – rarity-weighted richness), e) the ratio of land-driven species richness loss to total GHG-driven species richness loss and f) the ratio of land-driven rarity-weighted richness loss to total GHG-driven rarity-weighted richness loss.

Again, GHG-driven and land-driven biodiversity footprints show different patterns, with tropical regions no longer having consistently higher footprints. Production in Western Europe and North America also drives relatively high GHG-driven biodiversity loss, particularly in the ‘Other Food’ sector although Asia & Pacific has the highest GHG-driven footprints across all food groups (Figure 3d). Asia & Pacific’s ratio of land-driven to GHG-driven ratios ranges from around 40-100 (Figures 3e, f) which is extremely concerning given the high land-driven biodiversity footprints this region has. If emissions continue at this rate their impact on global biodiversity will, in under a century, equal that already caused by land conversion in Asia & Pacific.

Animal-derived products tend to have much higher footprints than plant-derived (i.e. crops), although Asia & Pacific is a notable exception for both land-driven and GHG-driven biodiversity footprints, in part due to production of paddy rice, wheat and other cereals in India (for individual product footprints see Supplementary Figure 5 and Supplementary Table 13). As would be expected, land-driven biodiversity loss from fertiliser production and the processing of food waste is much lower than that stemming from the production of food itself since the processes use very little agricultural land.

In many instances, the ‘Other Food’ category has higher production-based GHG-driven biodiversity footprints than the footprints from crops (Figures 3a-d). Other Food includes ‘Food products not elsewhere classified (nec)’ (coded in EXIOBASE as ‘OFOD’) and beverages (‘BEVR’). ‘Food products nec’ contributes the vast majority of the Other Food footprint and covers a broad range of processed foods such as soups, sandwiches and sauces. The EXIOBASE land-use types that contribute most to the Food Products nec land-driven footprints are cereal grains nec, cattle pasture, wheat, oil seeds, raw milk pasture and fruit/vegetables/nuts (Supplementary Figure 6).

### 3.3 Per-area and per-capita footprints

Taiwan stands out as having the highest per-area land-related production-based footprints (Figure 4a-c). The agriculturally dense but biodiversity poor UK is an example of a region with relatively lower per-area land-driven biodiversity footprints than per-area land-use footprints (Figure 4). In contrast, the species-rich Brazil has a lower per-area land-use footprint but a relatively higher per-area land-driven species richness footprint. Some regions (e.g. Brazil, India, Mexico) have both high total production-based footprints (Figure 2) and high per-area production-based footprints (Figure 4). The other regions with high per-area production tend to be smaller regions with wide agricultural coverage, for example the Netherlands and Belgium. Small agricultural regions have the highest per-area production-based GHG-driven footprints, for example Taiwan, the Netherlands and Malta. The relative contributions of the different GHGs differs between regions, for example nitrous oxide contributing a larger proportion of the Netherlands’ and Belgium’s GHG-driven footprint than Malta’s (Figure 4d). For per-area footprints for all products and regions see Supplementary Figure 7 and Supplementary Table 14.

**Figure 4.**
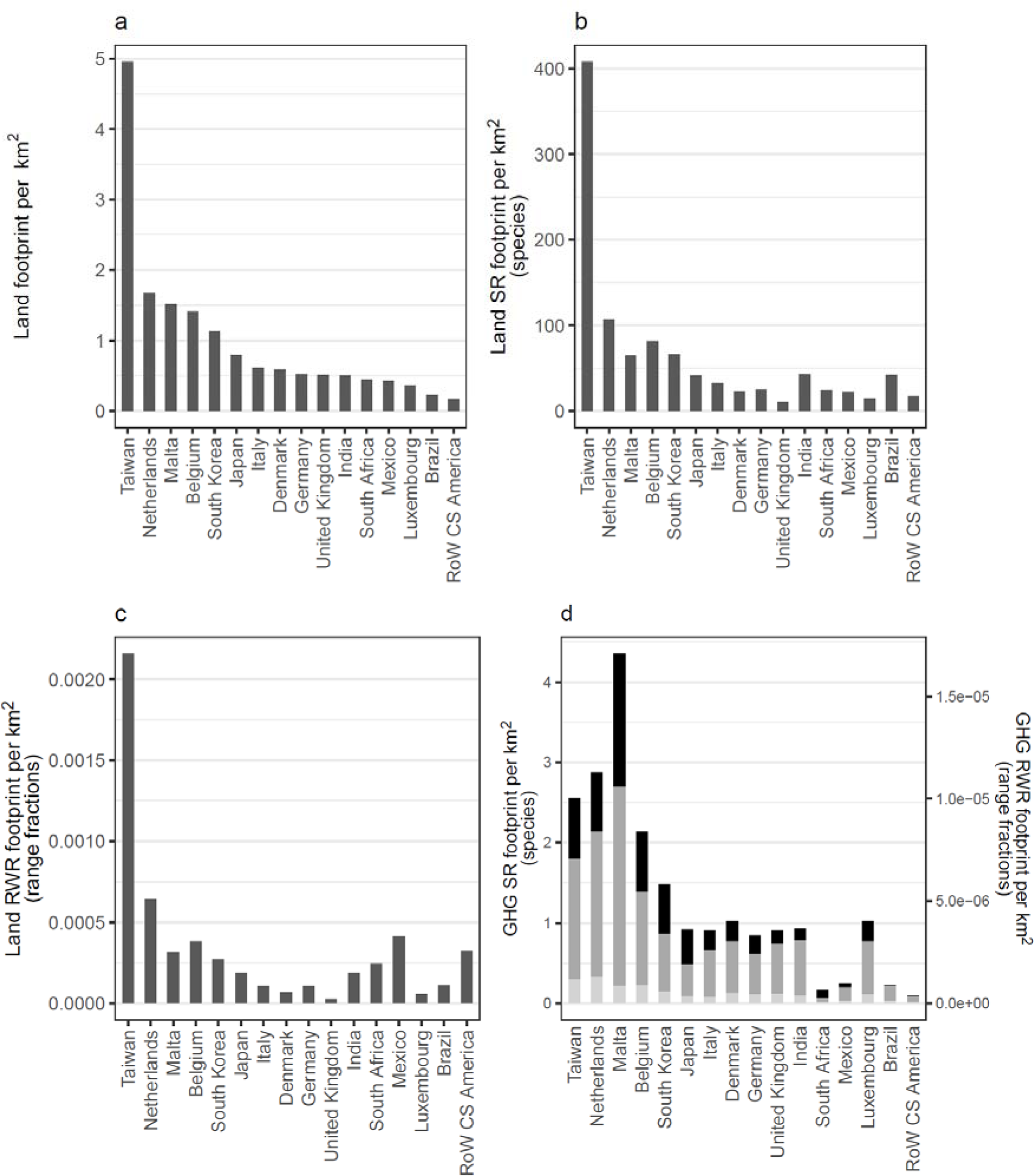
Total per-area production-based footprints of food-related products. Footprints relate to a) land use, b) land-driven species richness (SR) loss, c) land-driven rarity-weighted richness (RWR) loss and d) GHG-driven biodiversity loss split by emissions type: carbon dioxide (black), methane (dark grey), nitrous oxide (light grey) (right-hand axis - species richness; left-hand axis - rarity-weighted richness). Regions which are in the highest ten for one or more footprints are shown.

Australia is notable for having extremely high land-driven consumption-based footprints per capita (Figure 5), driven particularly by animal-based products (notably ‘Products of Cattle’), but also by plant-based products and ‘Other Food’ (Supplementary Figure 8, Supplementary Tables 15, 16). Luxembourg also has consistently high per-capita consumption-based footprints, probably in part due its highly affluent population^51^. In contrast to Australia, Luxembourg’s consumption-based footprint is primarily driven by plant-derived products (fruit/vegetables/nuts, oil seeds and other crops, i.e., coffee, cocoa, spices). Despite having high total consumption due to their large populations (Figure 2), China, India and RoW Asia & Pacific are not within the top ten highest per-capita consumption-based footprints for any category. However, some regions with high total biodiversity consumption also have high per-capita footprints: Australia, Brazil, Mexico, Russia, the United States, RoW Africa and RoW CS America. Again, we see contrasts between different footprint types. For example, Brazil’s land-driven species richness consumption-based footprint per capita is much larger than RoW CS America’s, but the latter’s per-capita rarity-weighted footprint is almost twice that of Brazil (Figures 5b,c). (Brazil has particularly high per-capita species richness footprints relative to RoW CS America for cattle, processed cattle, vegetable oils and dairy, see Supplementary Figure 8). Regions with relatively low species richness can have high per-capita land-driven biodiversity consumption-based footprints as a result of importing food.

**Figure 5.**
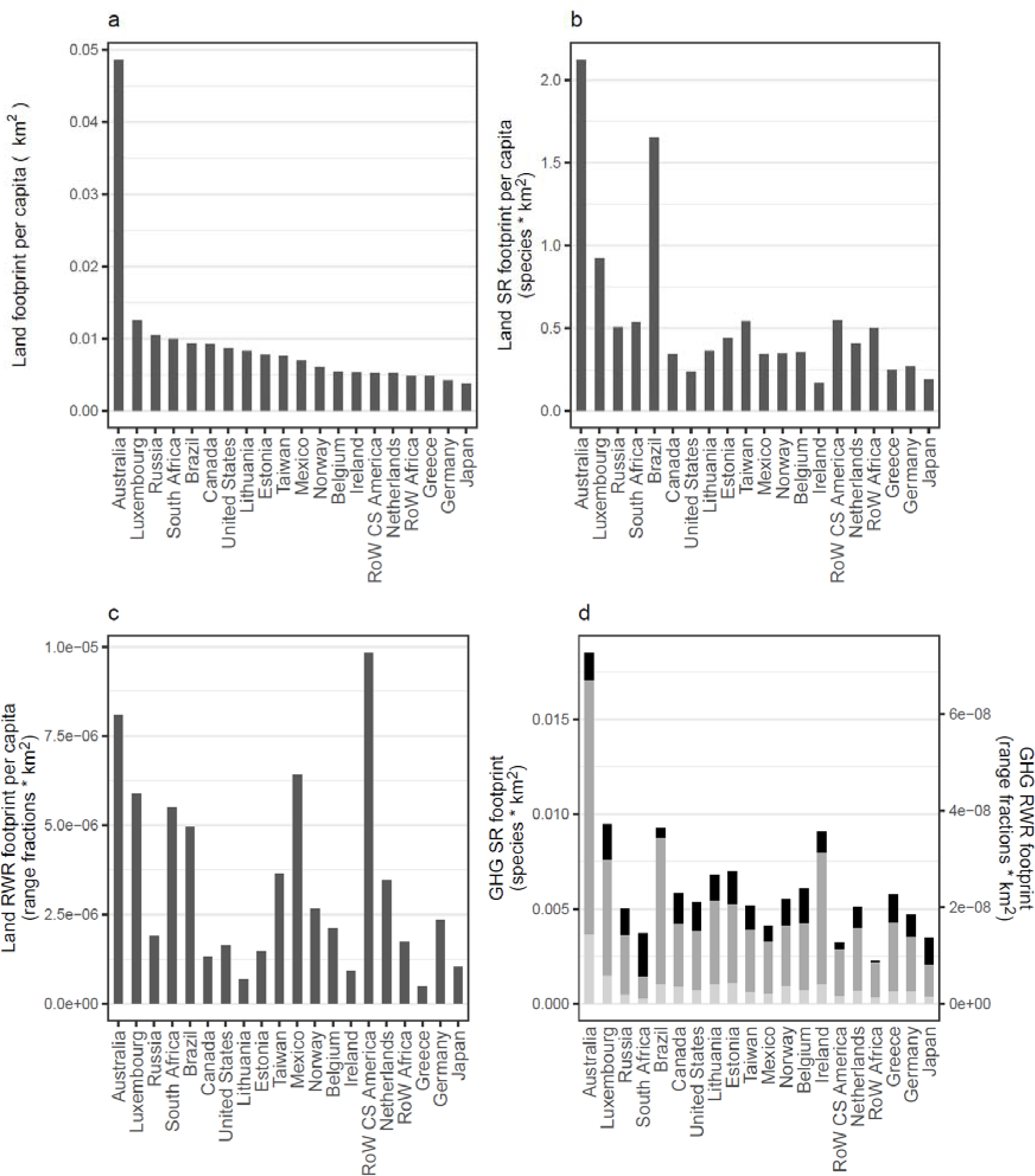
Total consumption-based footprints per-capita of food-related products. Footprints are in terms of a) land use, b) land-driven species richness (SR) loss, c) land-driven rarity-weighted richness (RWR) loss and d) GHG-driven biodiversity loss split by emissions type: carbon dioxide (black), methane (dark grey), nitrous oxide (light grey) (right-hand axis – species richness; left-hand axis – rarity-weighted richness). Regions which are in the highest ten for one or more footprints are shown.

The products underpinning high per-capita consumption-based GHG-driven biodiversity footprints vary among regions (Figure 5d). Belgium’s high footprint, for example, is largely driven by other foods (e.g. pizza, ready-meals, sauces) and dairy products, Estonia’s by dairy products, raw milk, cattle and other foods, Luxembourg’s by fruit/vegetable/nuts, other crops (i.e., coffee, cocoa, spices), cattle and chemical fertiliser, and the Republic of Ireland’s by processed cattle products and food waste disposed of in land fill. Some regions, e.g. Australia, Luxembourg and Lithuania, have high footprints across multiple products and at least one animal-based product is associated with particularly high consumption footprints for all regions in Figure 5, except Taiwan (< ∼0.0004 species x km^2^ per capita for every animal-based product) (see also Supplementary Figure 8 and Supplementary Tables 15, 16 for further breakdowns). Relative consumption of the different GHGs differs across regions, carbon dioxide contributing relatively highly to South Africa’s footprint but relatively little to Brazil’s, for example.

### 3.4 Biodiversity loss embedded in export/import trade

We found a near-reversal of net imports (i.e. imports minus exports, or consumption-based footprint minus production-based footprint) for land-driven versus GHG-driven biodiversity footprints (Figure 6, Supplementary Figure 9). The United States, the United Kingdom, Germany, Russia, Japan, China and RoW Middle East Ill regions that tend to be relatively biodiversity poor and highly industrialised in their land use systems Ill are all net importers of land-driven biodiversity loss but net exporters of GHG-driven biodiversity loss. The different land-driven footprints also show different messages. For example, RoW CS America exports a similar land-area footprint to that exported by RoW Asia & Pacific but approximately four times RoW Asia & Pacific’s exported rarity-weighted richness footprint. RoW Africa is a net exporter of land-driven species richness impacts but a net importer of land-driven rarity-weighted richness impacts. The three greatest net importers of both land-driven biodiversity footprints are RoW Middle East, Russia and the United States.

**Figure 6.**
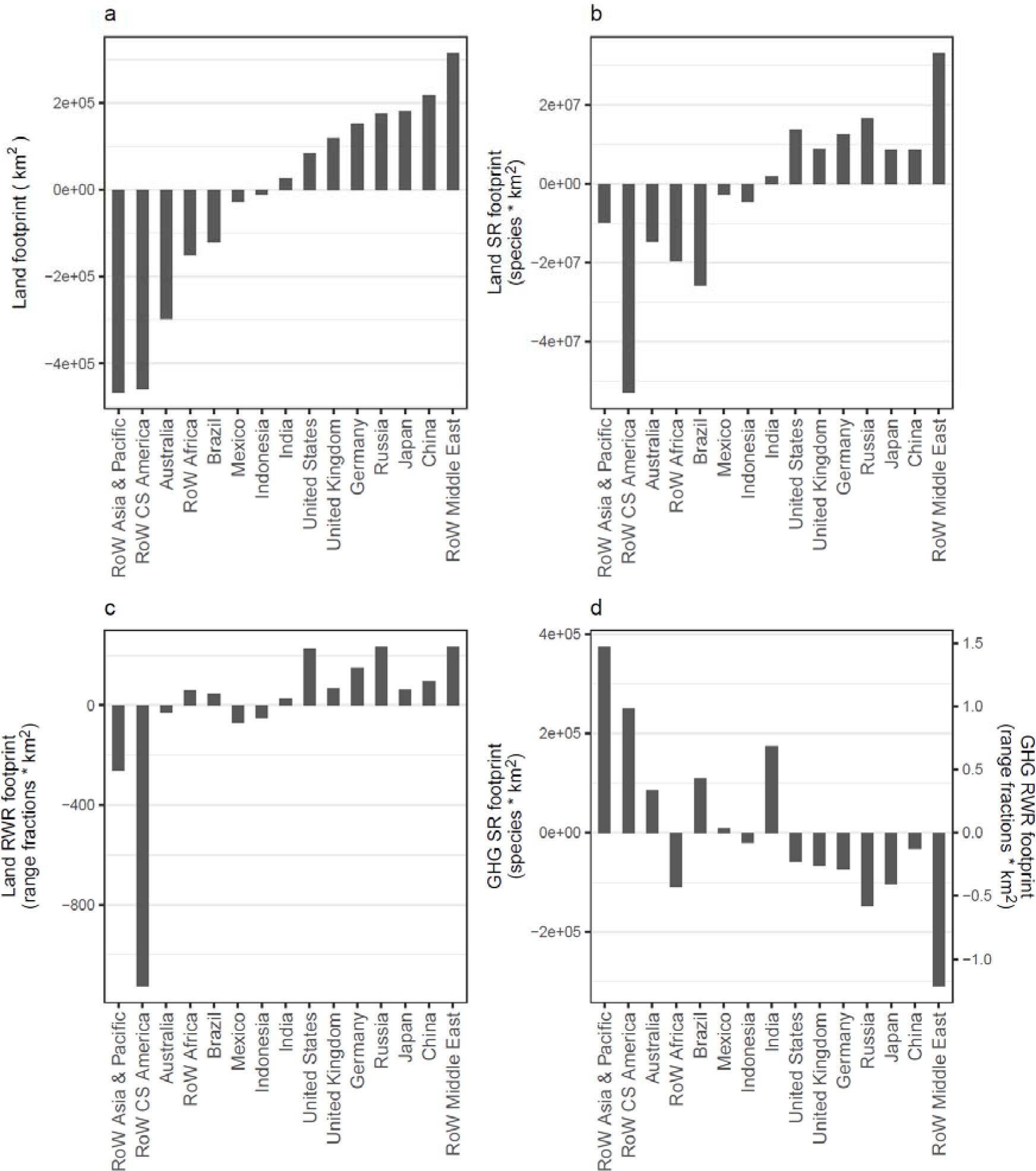
Regions’ net import footprints for food-related products. Footprints are in terms of a) land use, b) land-driven species richness (SR) loss, c) land-driven rarity-weighted richness (RWR) loss and d) GHG-driven richness loss. Regions which are in the highest five (net importers) or lowest five (net exporters) for the net import footprints are shown.

Understanding the percentage of a region’s footprint that is imported is important in devising policies to reduce footprints. Imports make up 5% or less of land-driven species richness footprints for highly biodiverse regions such as Brazil and Mexico (Figure 7b, Supplementary Figure 10); reducing domestic consumption in these regions might actually lower their consumption-based footprints. In contrast, the regions with the highest percentages of imported footprints are European. Luxembourg, the Republic of Ireland and the United Kingdom import 98%, 88% and 83% of their land-driven species richness footprints respectively. These high percentages will in part be due to the regions’ relatively low domestic food production footprints but may also reflect imports grown in highly biodiverse regions, which could potentially be sourced more sustainably. Luxembourg, Ireland and Norway import high percentages of their GHG-driven biodiversity footprints (69%, 65% and 48% respectively) (Figure 7d) and also have particularly high per capita consumption-based GHG-driven footprints (Figure 5d).

**Figure 7.**
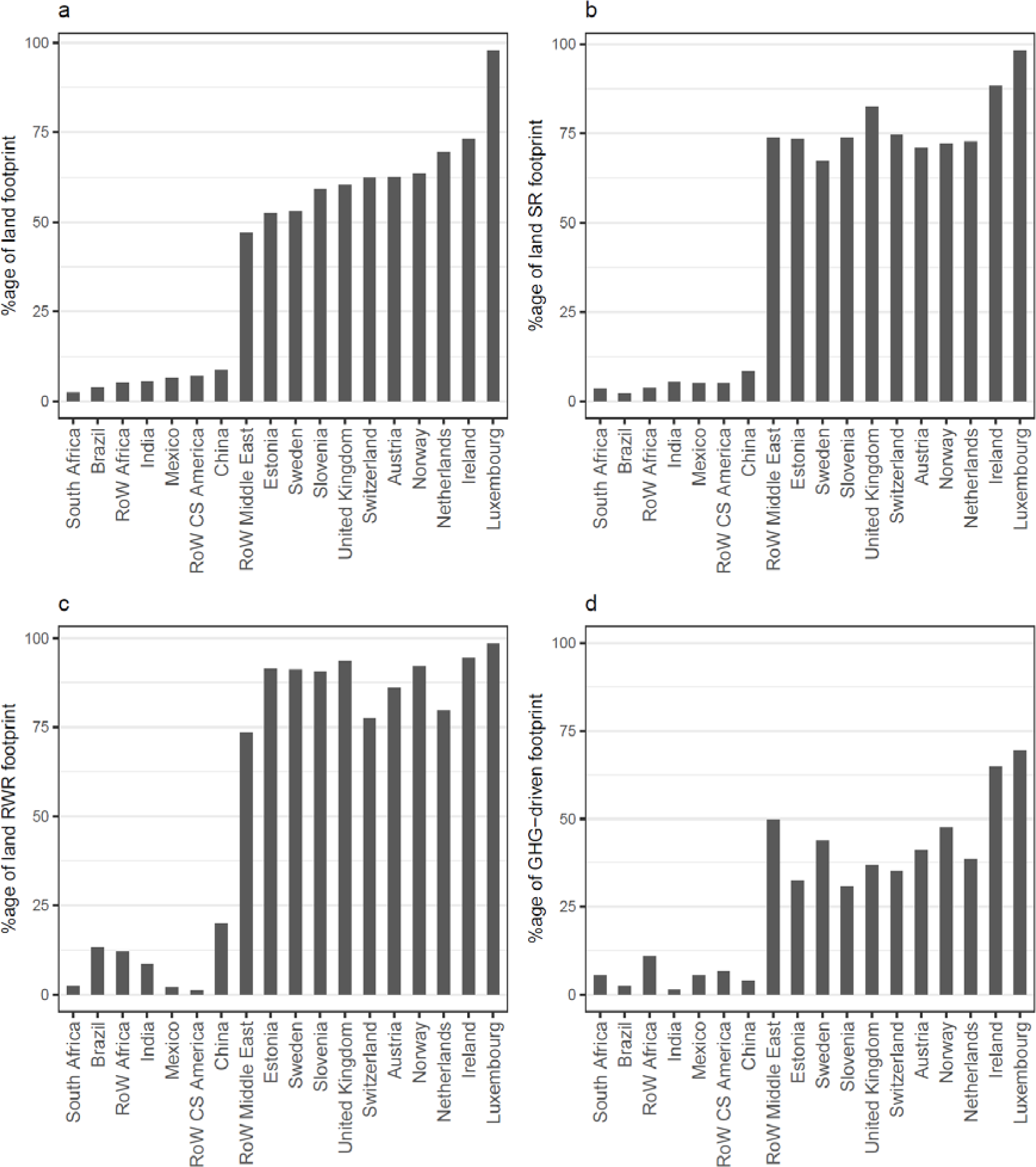
The percentage of a region’s food-related footprint that is imported. Footprints are in terms of a) land use, b) land-driven species richness (SR) loss, c) land-driven rarity-weighted richness (RWR) loss and d) GHG-driven richness loss. Regions which are in the 5 lowest or highest percentage for one or more footprints are shown.

We find that 10%, 15% and 8% respectively of land-driven species richness, land-driven rarity-weighted richness and GHG-driven richness footprints are embedded within trade between world regions (Supplementary Figure 11), with patterns differing between the footprint types, and between animal and plant-derived foods (Supplementary Figures 11-13). Although the total footprints of animal-derived products are higher than plant-derived, a greater percentage of the plant-derived footprint is traded between world regions: approximately 8%, 15%, 7% (animal-derived) versus 17%, 21%, 11% (plant-derived) of the total footprint for land-driven species richness, land-driven rarity-weighted richness and GHG-driven species richness respectively. These results are discussed in more detail in the Supplementary Information (Supplementary Figure 13).

## 4. Discussion

Our metric of land-driven species richness change captures the higher biodiversity footprint of products from more species-rich regions and from biomes in which biodiversity is more sensitive to land-use change. The metric thus provides a more accurate footprint of the biodiversity impact embedded within food than would be obtained using land area alone. Weighting richness by species’ range area further changes assessment of embedded biodiversity footprint. Rarity-weighted richness places a greater emphasis on the biodiversity costs in areas containing a higher proportion of narrow-ranged species, which are of greater conservation concern than wider-ranging species, notably parts of CS America. Our analysis also shows the need to consider GHG-driven biodiversity loss, particularly in assessing longer term future impacts of food production. We find that the climate contributions of food production are accumulating rapidly over time – in several regions, annual GHG-driven production footprints are as high as 2% of the total historic land-use footprint, meaning that just a decade’s worth of emissions (not including land conversion emissions) will add an additional 20% to the biodiversity loss that has already occurred due to wholesale conversion of land to agriculture. 70% of the total GHG-driven biodiversity footprint of food-related products stems from methane.

### 4.1 Comparison with other studies

Our study incorporated the novel combination of: (i) allowing sensitivity to land use to vary by land-use type and biome; (ii) allowing for the natural variation of species richness across political regions; and (iii) allowing for spatial variation in future temperature change and in species’ responses to that temperature change. The models we used to capture land-use and climate impacts output the same biodiversity metrics, allowing us to compare these two major drivers of biodiversity change. Comparing the results of our study with those of previous analyses of the embedded biodiversity footprints of food is not straightforward since studies generally differ in resolution with respect to regions and/or products. Where comparisons can be made, we find both common ground and divergence. In line with Marques et al.^26^, we show that production in Africa, CS America and Asia & Pacific regions has the highest overall biodiversity footprints, and that cattle products have a particularly high impact on biodiversity globally. Marques et al.^26^ use a biodiversity metric of number of impending extinctions of bird species and, in common with our land-driven species richness metric, identify almost equally high impacts of cattle in Africa as in CS America. However, it is only by using our rarity-weighted richness measure that the particularly high cost to narrow-ranged species from cattle products in CS America is revealed. Chaudhary and Kastner^15^ use a metric of the number of species committed to extinction. Their study had greater spatial disaggregation than ours meaning we cannot compare regions included within our RoW regions. Nevertheless, our results broadly support each other with respect to highlighting particularly high consumption footprints in India, China, Brazil and the US. However, Chaudhary and Kastner found Indonesia to have the second highest consumption footprint, whereas it does not appear in our top ten consumption footprints for any metric. The relative magnitude of the regions with the greatest consumption footprints in Chaudhary et al.’s^52^ study differs from our results. These discrepancies are likely to result from the biodiversity metric used as well as differences in trade models, and further support our finding that different biodiversity metrics lead to different conclusions. Sun et al.^53^ also use a metric of the number of species committed to extinction at a greater spatial disaggregation than our study. Their results show strong agreement with ours with respect to consumption per capita, both studies finding high footprints in Central America, Brazil, South Africa and Australia. Lenzen et al.^23^ look at the biodiversity footprint, as measured by number of threats to species, of all commodities, not just those associated with food. Both Lenzen et al.’s^23^ and our studies find the US, Germany, the UK and Japan to be among the greatest importers of embedded biodiversity but our study has Russia as a major importer of embedded biodiversity whereas Lenzen et al.^23^ find it to be one of the greatest net exporters.

Our estimates of the impacts of food-related GHGs on biodiversity are lower than those of Wilting et al.^27^ which is at least in part due to differences in methodology. We predicted the temperature rise in 2031 due to an emissions pulse in 2011 using the global temperature potential (GTP). Wilting et al.^27^ used the integrated GTP over a 100 year time horizon, thus summing the warming that occurred in every year following the emission pulse, as opposed to calculating the temperature of the hundredth year only. Wilting et al.^27^ analysed the land-driven and GHG-driven biodiversity loss from all economic sectors and estimated that GHGs contributed to an average of 18% of the biodiversity loss associated with food although contributions varied by region from 7% (Africa) to 45% (rest of Oceania). Given the different metric and time horizons of global warming used, we would expect our results to differ from Wilting et al.’s^27^, with our GHG-driven footprints estimated to contribute from the order of 0.1% – 6% of biodiversity loss for food products and up to 16% for fertiliser. Wilting et al.’s^27^ study further differed from ours since it used a different biodiversity metric and MRIO, and assumed agricultural land was evenly distributed across regions. Our studies support each other in suggesting current GHG-driven emissions from the food-related products will play a very significant role in future biodiversity loss. Reducing emissions associated with food is therefore a high priority, particularly in Europe, North America and Asia & Pacific.

### 4.2 Policy implications

Our study shows that measurements of trade-related impacts of biodiversity differ considerably depending on the biodiversity metric used. Whilst detailed discussion of the merits and relevance of different biodiversity metrics may be appropriate in the scientific literature, it is unlikely to be helpful to businesses and policy makers who require succinct and readily interpretable biodiversity footprints. This leads to the question of which measure(s) should be recommended to the public and private sectors to aid them in reducing their environmental footprints. What might the consequences be of one metric being chosen over others? Our study suggests that a metric of land area, whilst easy to measure and interpret, will over inflate the cost of biodiversity in biodiversity-poor areas and under-estimate it in biodiversity-rich areas, and thus is not recommended when the focus is biodiversity conservation. Species richness is preferable in this case, and is still relatively easy to interpret, particularly if used in relative rather than absolute terms. However, species richness has been criticised as a biodiversity metric, since it does not decline if rare species are replaced with common species (see Hillebrand et al.^31^). Weighting for species rarity overcomes this, and highlights the higher losses of smaller-ranged species from CS America. Using rarity-weighted richness over a simple species richness measure would likely lead to different decisions with respect to sustainable trading.

We must also consider whether GHG-driven biodiversity loss adds value or unnecessary complexity to footprint estimates. Our study has shown that it is fairly simple to calculate GHG-driven biodiversity loss and to aggregate it with the land-driven impact. The annual GHG-driven biodiversity impact of livestock sectors is equivalent to approximately 5% of land-driven biodiversity loss in Asia, the US and Western Europe. Importantly, this equates to 50% of the biodiversity impact of complete land conversion within the timespan of just one decade. For some individual products, the ratio of GHG to land-driven impacts is even higher. Including the GHG-driven losses in estimates of the food industry’s impact on biodiversity would therefore seem worthwhile, particularly since the bulk of emissions are from methane. Methane’s short lifetime means that (assuming methane sinks are constant), reducing methane emissions actually leads to global cooling but on the flipside, increased emissions lead to substantial rapid warming^54^. Reducing emissions from food in the immediate-term would therefore be an extremely effective route to reducing near-term warming. Our results add to the already weighty evidence, e.g.^55–57^, in support of policies that assist farmers to transition away from livestock and nudge consumers toward a more plant-based diet.

The biodiversity footprints that we present could be used to inform policies and trade agreements targeted at reducing biodiversity loss. Policies which encourage a reduction of high-income countries’ consumption of beef, for example, could be used to target biodiversity loss in Australia and CS America, regions which dominate exported biodiversity footprint embedded in cattle products. Trade deals which lead to an increase in imports from these regions (e.g. the 2021 UK-Australia trade deal, the European Union’s ‘New Green Deal’) will likely lead to an increase in biodiversity loss and offshoring of yet more of Europe’s environmental footprint^58^ unless strict and enforceable environmental regulations are included in the agreements. Decisions on future trade deals must take embedded environmental costs into account and use appropriate metrics of biodiversity. If impacts are based on land use alone they will severely underestimate biodiversity impacts in CS America.

The examination of production-based footprints per-area (which facilitate comparisons of production between different-sized regions) allows for an increased understanding of the implications of regional/national agricultural policies. Regions with high per-area production-based footprints are targets for policies that reduce the environmental impact of farming. There is considerable overlap of high per-area land-driven and GHG-driven production impacts and it is interesting that the Netherlands and Belgium, both less species-rich temperate countries, have some of the highest per-area production-based footprints, even for rarity-weighted richness (Figure 4). Taiwan is notable for all three of its per-area land-driven production-based footprints being extremely high (Figure 4), with impacts being driven by a variety of products including fruit/vegetables/nuts, other animal products (e.g., eggs, honey), fish, pork products, sugar, and other food products (e.g., pizza, soups, sauces) (Supplementary Figure 7). India stands out as having very high paddy rice production per area for all footprints (Supplementary Figure 7).

Per-capita consumption-based footprints (which facilitate comparisons of consumption between regions with different population sizes) also allow for targeting of agricultural and trade policies. We found China, India and RoW Africa had the highest overall GHG-driven biodiversity footprints, but once population size was considered, only RoW Africa remained in the top ten per-capita footprints. Identifying the products that lead to regions’ high per-capita footprints is the first step towards reducing those footprints. For example, extremely high per-capita consumption-based footprints from cattle, raw milk and dairy products contribute to Estonia’s place in the top ten for GHG-driven biodiversity loss (see Figure 5 and Supplementary Figure 8). Whilst we recognise that changing cultural norms can be very difficult, reducing Estonia’s per-capita footprint would be relatively straight forward from a policy perspective, since it involves only one food group and could perhaps be tackled via a combination of dietary shifts, source-shifting and change in dairy-farming practices. Australia, in contrast, has particularly high per-capita land-driven consumption-based footprints for a wide variety of products spanning fruit/vegetables/nuts, all products associated with cattle and other meat animals (e.g. sheep), rice, other food products and beverages. Reducing these footprints will likely require a more complex suite of strategies.

We found that some regions import over 90% of their embedded biodiversity footprint, further indicating the need for sustainable trade as a measure to halt biodiversity loss. Regions with a high proportion of imported footprint coupled with high per-capita consumption-based footprints are targets for reducing their imports and/or the biodiversity footprint embedded within those imports^16^; Estonia, Luxembourg, the Netherlands and Norway all fall into this category (Figures 5 and 7). Conversely, regions with a low percentage of imported footprint but high per-capita consumption such as Brazil, Mexico and South Africa should be lowering their domestic footprint in order to reduce their biodiversity impact.

The main net importers of land-driven biodiversity footprint tend to be net exporters of GHG-driven biodiversity footprint, and vice versa. The emissions associated with crop production largely arise from fertiliser use. Low fertiliser application will be associated with low emissions but lead to a smaller yield and therefore a larger land-use footprint. On the other hand, heavy fertiliser use will result in more emissions but smaller land use. This negative correlation is exacerbated since areas that use more fertiliser tend to be areas of lower species richness such as N America and W Europe^59^.

### 4.3 Limitations

There are several limitations of MRIOs which must be considered in interpreting our results: sectoral and regional aggregation^60,61^, an assumption of price homogeneity^62,63^, limitations in data reporting^64^, and the age of input data^65^. The original EXIOBASE 3 data end in 2011 and although more recent trade data are now available, land use is still limited to 2011 (https://zenodo.org/record/5589597#.YnkHvOjMK3A). Our results therefore reflect the biodiversity impact of food production over a decade ago, and will likely underestimate the current impact due to the expansion of agricultural land^66^ in the intervening period.

In common with other footprinting studies, e.g.^25,26^, our analysis only measures biodiversity impacts caused by direct land-use change, and does not explicitly consider the impacts of agricultural intensification. Neither does our study explicitly account for indirect biodiversity loss that might occur through loss of ecosystem function, e.g. soil impoverishment. Nevertheless, the PREDICTS database does sample gradients of agricultural intensity and ecosystem degradation, and so these factors should be captured implicitly^29^.

While our study highlights that the consumption of cattle products contributes considerably to land-driven biodiversity loss, it is important to be aware that there is high variability in pasture maps stemming from the different definitions, methods and underlying datasets used in their generation^67^. However, the Ramankutty et al.^39^ dataset that we used is one of the best available, aligning well with other datasets^67^.

### 4.4 Conclusions

We have presented new methods to estimate the land-driven and GHG-driven biodiversity impacts embedded within the production, consumption and trade of food-related commodities. We find that land-driven footprints differ depending on the metric used, and that GHG-driven biodiversity impacts are driven largely by methane emissions and contribute a higher proportion of the total footprint in regions of higher income and lower species-richness. The measures we present are simple to calculate and could be incorporated into decision making and environmental impact assessments by governments and businesses. Our consistent metric for biodiversity impacts allows us to present multiple aspects of land-driven and GHG-driven biodiversity footprints, enabling insight into priorities for reducing biodiversity costs via both global and regional production and consumption.

## Data availability

The EXIOBASE 3.8.1 pxp data that were used in our study are publicly available here: https://zenodo.org/record/4588235#.ZGT8NnbMK3B

## Supporting information

Supplementary Methods; Figures S1, S2, S3, S6 & S13; Legends for all Supplementary Figures; Tables S1-S5

## Notes

### Competing Interest Statement

The authors have declared no competing interest.

### Summary of Updates

The manuscript has been updated as a result of a bug in our code. Further detail has been added to the results section, splitting greenhouse gases into three components: carbon dioxide, methane and nitrous oxide. The abstract,introduction, methods, results and discussion have been updated where relevant to incorporate these changes. All figures and supplemental files have been updated to reflect the revised results.

## References

1 IPBES. Global assessment report on biodiversity and ecosystem services of the Intergovernmental Science-Policy Platform on Biodiversity and Ecosystem Services. (Bonn, Germany, 2019).

2 Maxwell, S. L., Fuller, R. A., Brooks, T. M. & Watson, J. E. M. The ravages of guns, nets and bulldozers. Nature 536, 143–145 (2016).

3 Schipper, A. M. et al. Projecting terrestrial biodiversity intactness with GLOBIO 4. Glob Chang Biol 26, 760–771 (2020). 10.1111/gcb.14848

4 Di Marco, M. et al. Projecting impacts of global climate and land-use scenarios on plant biodiversity using compositional-turnover modelling. Glob Chang Biol 25, 2763–2778 (2019). 10.1111/gcb.14663

5 Newbold, T. Future effects of climate and land-use change on terrestrial vertebrate community diversity under different scenarios. Proc Biol Sci 285 (2018). 10.1098/rspb.2018.0792

6 Ellis, E. C., Klein Goldewijk, K., Siebert, S., Lightman, D. & Ramankutty, N. Anthropogenic transformation of the biomes, 1700 to 2000. Global Ecology and Biogeography, no-no (2010). 10.1111/j.1466-8238.2010.00540.x

7 FAOStat. (License: CC BY-NC-SA 3.0 IGO., 2022). 8

8 Mbow, C. et al. Food Security. (2019).

9 Rosenzweig, C., N Tubiello, F., Sandalow, D., Benoit, P. & N Hayek, M. Finding and fixing food system emissions: the double helix of science and policy. Environmental Research Letters 16 (2021). 10.1088/1748-9326/ac0134

10 Crippa, M. et al. Food systems are responsible for a third of global anthropogenic GHG emissions. Nature Food 2, 198–209 (2021).

11 Leclere, D. et al. Bending the curve of terrestrial biodiversity needs an integrated strategy. Nature 585, 551–556 (2020). 10.1038/s41586-020-2705-y

12 Lynch, J., Cain, M., Frame, D. & Pierrehumbert, R. Agriculture’s contribution to climate change and role in mitigation is distinct from predominantly fossil CO2-emitting sectors. Frontiers in Sustainable Food Systems 4, 518039 (2021).

13 Kastner, T., Erb, K.-H. & Haberl, H. Rapid growth in agricultural trade: effects on global area efficiency and the role of management. Environmental Research Letters 9, 034015 (2014).

14 FAO. The State of Agricultural Commodity Markets 2020. Agricultural markets and sustainable development: Global value chains, smallholder farmers and digital innovations. (FAO, 2020).

15 Chaudhary, A. & Kastner, T. Land use biodiversity impacts embodied in international food trade. Global Environmental Change 38, 195–204 (2016). 10.1016/j.gloenvcha.2016.03.013

16 Kitzes, J. et al. Consumption-Based Conservation Targeting: Linking Biodiversity Loss to Upstream Demand through a Global Wildlife Footprint. Conserv Lett 10, 531–538 (2017). 10.1111/con4.12321

17 Balmford, A. et al. The environmental costs and benefits of high-yield farming. Nature Sustainability 1, 477–485 (2018). 10.1038/s41893-018-0138-5

18 Miller, R. E. & Blair, P. D. Input-output Analysis: Foundations and Extension. (Cambridge University Press, 2009).

19 Kitzes, J. An introduction to environmentally-extended input-output analysis. Resources 2, 489–503 (2013).

20 Roux, N., Kastner, T., Erb, K.-H. & Haberl, H. Does agricultural trade reduce pressure on land ecosystems? Decomposing drivers of the embodied human appropriation of net primary production. Ecological Economics 181 (2021). 10.1016/j.ecolecon.2020.106915

21 Peters, G. & Hertwich, E. G. CO_2_ embodied in international trade with implications for global climate policy. Environmental Science and Technology 42, 1401–1407 (2008).

22 Hoekstra, A. Y. & Mekonnen, M. M. The water footprint of humanity. Proceedings of the National Academy of Sciences 109, 3232–3237 (2012).

23 Lenzen, M. et al. International trade drives biodiversity threats in developing nations. Nature 486, 109–112 (2012).

24 Moran, D. & Kanemoto, K. Identifying species threat hotspots from global supply chains. Nature Ecology and Evolution 1, 0023 (2017).

25 Chaudhary, A., Verones, F., de Baan, L. & Hellweg, S. Quantifying Land Use Impacts on Biodiversity: Combining Species-Area Models and Vulnerability Indicators. Environ Sci Technol 49, 9987–9995 (2015). 10.1021/acs.est.5b02507

26 Marques, A. et al. Increasing impacts of land use on biodiversity and carbon sequestration driven by population and economic growth. Nat Ecol Evol 3, 628–637 (2019). 10.1038/s41559-019-0824-3

27 Wilting, H. C., Schipper, A. M., Bakkenes, M., Meijer, J. R. & Huijbregts, M. A. Quantifying Biodiversity Losses Due to Human Consumption: A Global-Scale Footprint Analysis. Environ Sci Technol 51, 3298–3306 (2017). 10.1021/acs.est.6b05296

28 Alkemade, R. et al. GLOBIO3: A Framework to Investigate Options for Reducing Global Terrestrial Biodiversity Loss. Ecosystems 12, 374–390 (2009). 10.1007/s10021-009-9229-5

29 Newbold, T., Oppenheimer, P., Etard, A. & Williams, J. J. Tropical and Mediterranean biodiversity is disproportionately sensitive to land-use and climate change. Nat Ecol Evol 4, 1630–1638 (2020). 10.1038/s41559-020-01303-0

30 Martins, I. S. & Pereira, H. M. Improving extinction projections across scales and habitats using the countryside species-area relationship. Sci Rep 7, 12899 (2017). 10.1038/s41598-017-13059-y

31 Hillebrand, H. et al. Biodiversity change is uncoupled from species richness trends: Consequences for conservation and monitoring. Journal of Applied Ecology 55, 169–184 (2018). 10.1111/1365-2664.12959

32 Chichorro, F., Juslén, A. & Cardoso, P. A review of the relation between species traits and extinction risk. Biological Conservation 237, 220–229 (2019).

33 Stadler, K., et al. EXIOBASE 3: Developing a time series of detailed environmentally extended multi-regional input-output tables. Journal of Industrial Ecology 10.1111/jiec.12715 (2018).

34 Smith, P. & Gregory, P. J. Climate change and sustainable food production. Proc Nutr Soc 72, 21–28 (2013). 10.1017/S0029665112002832

35 ESRI. (2015).

36 The Nature Conservancy. (2019).

37 International Food Policy Research Institute. MapSPAM (ed International Food Policy Research Institute) (Harvard Dataverse, 2019).

38 Monfreda, C., Ramankutty, N. & Foley, J. A. Farming the planet: 2. Geographic distribution of crop areas, yields, physiological types, and net primary production in the year 2000. Global Biogeochemical Cycles 22, GB1022 (2008). doi: 10.1029/2007GB002947

39 Ramankutty, N., Evan, A. T., Monfreda, C. & Foley, J. A. Farming the planet: 1. Geographic distribution of global agricultural lands in the year 2000. Global Biogeochemical Cycles 22, GB1003 (2008). doi:10.1029/2007GB002952

40 IUCN, I. U. f. C. o. N. (2020).

41 BirdLife International. (2020).

42 Meiri, S. et al. (2017).

43 Roll, U. et al. The global distribution of tetrapods reveals a need for targeted reptile conservation. Nat Ecol Evol 1, 1677–1682 (2017). 10.1038/s41559-017-0332-2

44 Etard, A., Morrill, S. & Newbold, T. Global gaps in trait data for terrestrial vertebrates. Global Ecology and Biogeography 29, 2143–2158 (2020). 10.1111/geb.13184

45 Hudson, L. N. et al. The database of the PREDICTS (Projecting Responses of Ecological Diversity In Changing Terrestrial Systems) project. Ecol Evol 7, 145–188 (2017). 10.1002/ece3.2579

46 Hudson, L. N. et al. (ed Natural History Museum) (10.5519/0066354, 2016).

47 Shine, K. P., Fuglestvedt, J. S., Hailemariam, K. & Stuber, N. Alternatives to the global warming potential for comparing climate impacts of emissions of greenhouse gases. Climate Change 68, 281-302 (2005).

48 Myhre, G. et al. Anthropogenic and Natural Radiative Forcing. (Cambridge University Press, Cambridge, United Kingdom and New York, NY, USA, 2013).

49 Hijmans, R., Cameron, S. E., Parra, J. L., Jones, P. G. & Jarvis, A. Very high resolution interpolated climate surfaces for global land areas. International Journal of Climatology 25, 1965–1978 (2005). 10.1002/joc.1276

50 CountryEconomy.com. https://countryeconomy.com/demography/population/taiwan?year=2011 2021).

51 World Bank. (2021).

52 Chaudhary, A., Carrasco, L. R. & Kastner, T. Linking national wood consumption with global biodiversity and ecosystem service losses. Sci Total Environ 586, 985–994 (2017). 10.1016/j.scitotenv.2017.02.078

53 Sun, Z., Behrens, P., Tukker, A., Bruckner, M. & Scherer, L. Shared and environmentally just responsibility for global biodiversity loss. Ecological Economics 194 (2022). 10.1016/j.ecolecon.2022.107339

54 Cain, M. et al. Improved calculation of warming-equivalent emissions for short-lived climate pollutants. NPJ climate and atmospheric science 2, 29 (2019).

55 Kozicka, M. et al. Feeding climate and biodiversity goals with novel plant-based meat and milk alternatives. Nature Communications 14, 5316 (2023).

56 Dimbleby, H. National Food Strategy Independent Review: The Plan. (2021).

57 Ivanovich, C. C., Sun, T., Gordon, D. R. & Ocko, I. B. Future warming from global food consumption. Nature Climate Change 13, 297–302 (2023).

58 Fuchs, R., Brown, C. & Rounsevell, M. Europe’s Green Deal offshores environmental damage to other nations. Nature 586, 671–673 (2020).

59 Mueller, N. D. et al. Closing yield gaps through nutrient and water management. Nature 490, 254–257 (2012). 10.1038/nature11420.490:254-257

60 Steen-Olsen, K., Owen, A., Hertwich, E. G. & Lenzen, M. Effects of sector aggregation on CO_2_ multipliers in multiregional input-output analyses. Economic Systems Research 26, 284–302 (2014).

61 Lenzen, M. Aggregation versus disaggregation in input-output analysis of the environment. Economic Systems Research 23, 73–89 (2011).

62 Merciai, S. & Heijungs, R. Balance issues in monetary input-output tables. Ecological Economics 102, 69–74 (2014).

63 de Koning, A. et al. Effect of aggregation and disaggregation on embodied material use of products in input-output analysis. Ecological Economics 116, 289–299 (2015).

64 Tukker, A. et al. Towards robust, authoratitive assessments of environmental impacts embodied in trade: Current state and recommendations. Journal of Industrial Ecology 22 (2018).

65 Chandrakumar, C., McLaren, S., Malik, A., Ramilan, T. & Lenzen, M. Understanding New Zealand’s consumption-based greenhouse gas emissions: an application of multi-regional input-output analysis. The International Journal of Life Cycle Assessment 25, 1323–1332 (2020).

66 FAO. How to Feed the World in 2050. (Rome, Italy, 2009).

67 Oliveira, J. et al. Choosing pasture maps: An assessment of pasture land classification definitions and a case study of Brazil. International Journal of Applied Earth Observation and Geoinformation 93 (2020). 10.1016/j.jag.2020.102205

