## Supplementary Methods; Figures S1, S2, S3, S6 & S13; Legends for all Supplementary Figures; Tables S1-S5 for "Impacts of global food supply on biodiversity via land use and climate change"

*Details of the cropland data sources*

SPAM^1^ presents gridded land-use estimates for 42 individual crops at a 5-arc-minute spatial resolution, derived from satellite-based land-cover data from around the year 2010; Monfreda et al. ^2^ present similar estimates for 175 crop types, also at a 5-arc-minute spatial resolution, derived from agricultural census data and satellite-based land-cover data from around the year 2000. Where possible, we chose SPAM data over Monfreda et al. ^2^ since it is closer in time to our 2011 input-output table. However, any disjoint between the datasets should have only a minor impact on the final results, since these estimates were used only to estimate the average natural species richness across farmed areas (at a 10-km resolution), which is not likely to have changed markedly between 2000 and 2010. We aggregated SPAM’s 42 crop layers to align with the EXIOBASE land-use categories of paddy rice, wheat, cereal grains nec (‘not elsewhere classified’), vegetables/fruit/nuts, oil seeds, sugar cane/sugar beet and crops nec. Since SPAM does not separate fodder from non-fodder crops, we obtained spatial estimates of fodder crops from Monfreda et al. ^2^. Monfreda et al. ^2^ do not disaggregate fodder crops into the different livestock categories given by EXIOBASE so, following Marques et al. ^3^, Monfreda et al.’s ^2^ 15 fodder crop layers were aggregated and then replicated for each EXIOBASE cropland-fodder category (fodder-cattle, fodder-pigs, fodder-poultry, fodder-meat nec, fodder-raw milk). In other words, we assumed that all 5 EXIOBASE fodder crop categories were present wherever any Monfreda et al. ^2^ fodder crop occurred. Ramankutty et al. ^4^ present pasture data at 5-arc-minute spatial resolution, also derived from census data and satellite-based land-cover data from around the year 2000. Again, separating pasture into the different livestock categories given by EXIOBASE (Permanent pasture: grazing-cattle, grazing-meat animals, grazing-raw milk) was not possible so the same pasture layer was allocated to each EXIOBASE pasture category. Cells which contained less than 2.5% of each agricultural land-use type were discarded. Supplementary Table 3 shows the allocation of crops and pasture to EXIOBASE’s land-use categories. The methods were implemented using the R packages raster v 3.4-5^5^ and rgdal v 1.5-23^6^.

*Details of the PREDICTS model*

The PREDICTS database^7^ contains 3.25 million snapshot samples of biodiversity in different land uses from 666 different studies, collected from 26,114 locations in all of the world’s terrestrial biomes (see Supplementary Figure 1). The PREDICTS data represent 47,044 different species, including plants, fungi, invertebrates and vertebrates. Sites in the PREDICTS database are classified into one of six different land-use types, based on the description of the habitat given in the original source publication or following consultation with the authors of the original study: primary vegetation (natural habitat, with no record of prior destruction by human activities or extreme natural events), secondary vegetation (destroyed in the past, but now recovering toward a natural state), plantation forest (areas used to cultivate woody crops), cropland (areas used to cultivate herbaceous crops, including for use as livestock fodder), pasture (areas used directly for grazing livestock, including both managed pasture and rangeland), and urban (areas of human settlement or civic amenity).

For the purposes of this study, we considered just four PREDICTS land-use classes: primary vegetation, secondary vegetation, agriculture (including both plantations and herbaceous cropland) and pasture. As in a recent study^8^, we allowed the response of biodiversity to these different land-use categories to vary among six broad biome groupings (Supplementary Table 2). Unlike the estimates of biodiversity responses to climate change, which because of data availability could be made only for vertebrates, we consider all taxonomic groups in estimating land-use responses. This broader taxonomic scope allowed us to include more data, and thus to consider how responses varied among biomes. We measured biodiversity at each site in the PREDICTS database in terms of species richness and rarity-weighted species richness. Species richness was measured, as in previous studies using the PREDICTS database e.g. ^8^, as the count of the unique species sampled at a site. To account for differences in the species composition of communities, and to consider potential shifts in the representation of narrow- versus wide-ranged species within communities, previous studies have used a measure of abundance-weighted community-average (log_­10_-transformed) species range size as an additional measure of biodiversity (e.g. ^8^). Species’ range sizes were estimated as the number of 110-km grid cells with a record for that species in the Global Biodiversity Information Facility database e.g.^8^. Using GBIF data, rather than the expert-derived range maps provided for vertebrates by the IUCN and others (see main text), allowed us to obtain range-size estimates for non-vertebrate species. Here, we modified this measure to calculate rarity-weighted species richness. Specifically, we calculated species weights based on the inverse of estimated range size, and calculated rarity-weighted species richness as the sum of weights of all species sampled at a given site. Within each study in the PREDICTS database, these weights were rescaled linearly, such that the highest weight (i.e., for the species with the smallest range) had a value of 1. We performed this rescaling because range sizes vary enormously among regions and species, and thus among PREDICTS studies, which prevents model convergence. Given the hierarchical structure of the models (see below), the rescaling should not affect estimates of the impact of land use on relative rarity-weighted species richness.

We fitted the response of species richness to land use using a generalized linear mixed-effects model with a Poisson distribution of errors (Supplementary Figure 2, Supplementary Tables 4, 5). Since rarity-weighted species richness had many non-integer values, and many values less than one, a Poisson model was not appropriate. Instead, we log_e_-transformed rarity-weighted species richness values, adding 1% of the mean (0.014) to all values to deal with zeroes, and then fitted responses to land use using a linear mixed-effects model. Fixed effects for both models were land use, biome, and their interaction. Random effects were study identity (to account for variation in biodiversity among studies caused by differences in sampling methods and effort, the taxonomic groups sampled, and broad geographic differences), and for the species richness model also site nested within study (to account for the over-dispersion present) ^9^.

*Methods for calculating the sensitivity of biodiversity to climate change*

Projected changes in the distributions of species’ (and thus changes in local species richness across terrestrial areas) as a result of climate change were derived from^10^. The response of species’ distributions to climate was captured using five different species distribution modelling algorithms (BIOCLIM, DOMAIN, Maxent, Random Forests and Generalised Linear Models). These models each related species’ observed distributions according to the IUCN Red List^11^ and Birdlife International^12^, to four climatic variables shown in previous studies to be strong correlates of animal distributions: minimum temperature of the coldest month, total annual precipitation, growing degree days and water balance, derived from the Worldclim Version 1.4 database^13^, which captures average climatic conditions for the period 1961-1990. BIOCLIM fits relationships between distribution records and climatic variables using a bounding-box approach in niche space, DOMAIN by comparing the climatic similarity between observed occurrence points and potentially inhabitable areas, random forests using a machine-learning approach to identify climatic patterns in species’ occurrence records, while generalized linear models and Maxent use classical parametric statistics or a maximum-entropy approach, respectively, to fit linear and quadratic relationships between species’ occurrences and the climatic variables. The resulting distribution models were projected onto the Worldclim Version 1.4 estimates of future climatic conditions under the RCP scenarios, averaged across the years 2061-2080, and averaged across outputs from an ensemble of different Global Climate Models^13^. Species were assumed to be limited in their ability to disperse in response to climate change, such that mammals and birds could move 3 km year^-1^, and reptiles and amphibians 0.5 km year^-1^ from conditions currently estimated to be suitable for the species^10^. Species were further considered unable to move outside the combinations of biogeographic realm and biome estimated to be occupied in the present day according to the original range maps from the IUCN and Birdlife International^10^. All calculations (both distribution models and their projections) were performed on a grid with 10-km spatial resolution^10^. Newbold (2018) ^10^ estimated a single metric of biodiversity, local species richness change. We also calculated rarity-weighted species richness change. To do so, each species was given a weight equal to the inverse of the number of 10-km grid cells estimated to be climatically suitable in the baseline period (1961-1990). We then calculated rarity-weighted species richness for both the baseline and future periods by summing these weights for all species for which each grid cell was estimated to be climatically suitable, and reachable given dispersal constraints. To calculate climate sensitivity, we divided projected change in species richness or rarity-weighted species richness between the baseline period and 2061-2080 by the projected change in temperature over the same time period (according to the same climate models used to drive the biodiversity projections). We focused here on projections of biodiversity and temperature under the RCP 8.5 climate scenario. This gave estimates of the sensitivity of biodiversity to climate change for every 10-km grid cell on the land surface of the world.

**Supplementary Figures**

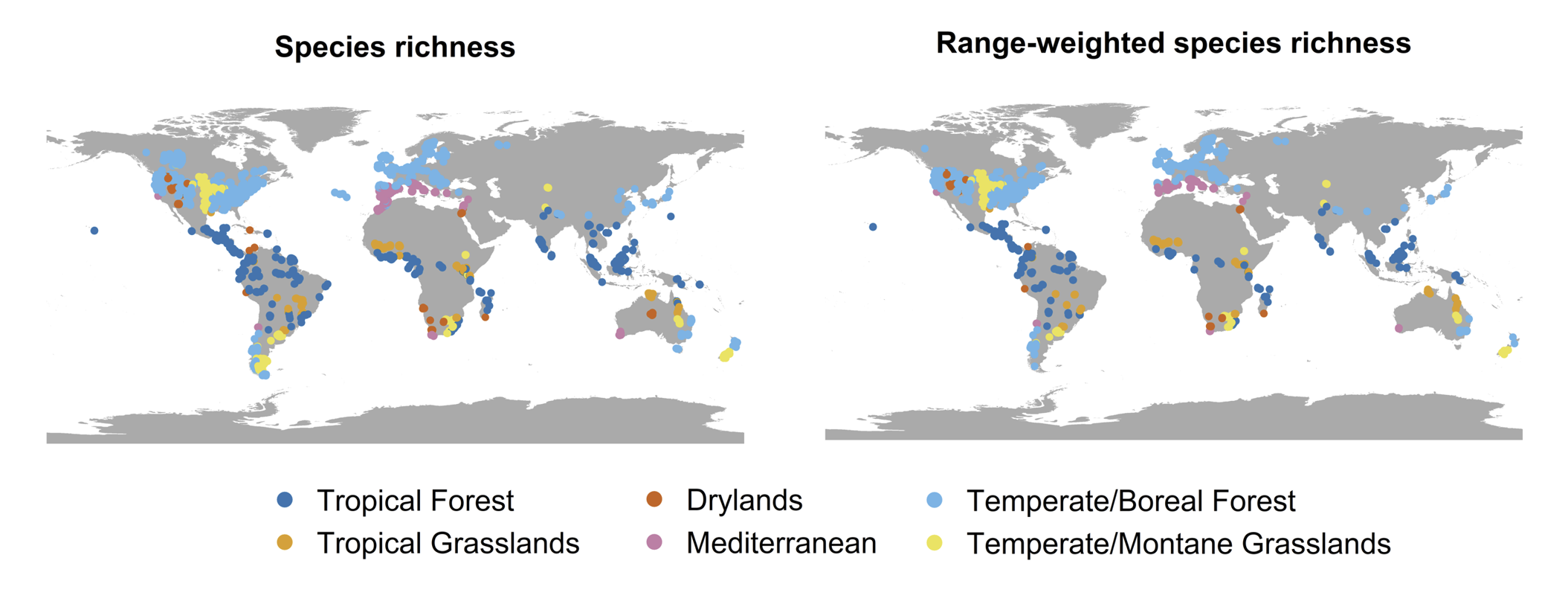
 Supplementary Figure 1. The locations (colour-coded by biome) of the studies in the PREDICTS database.

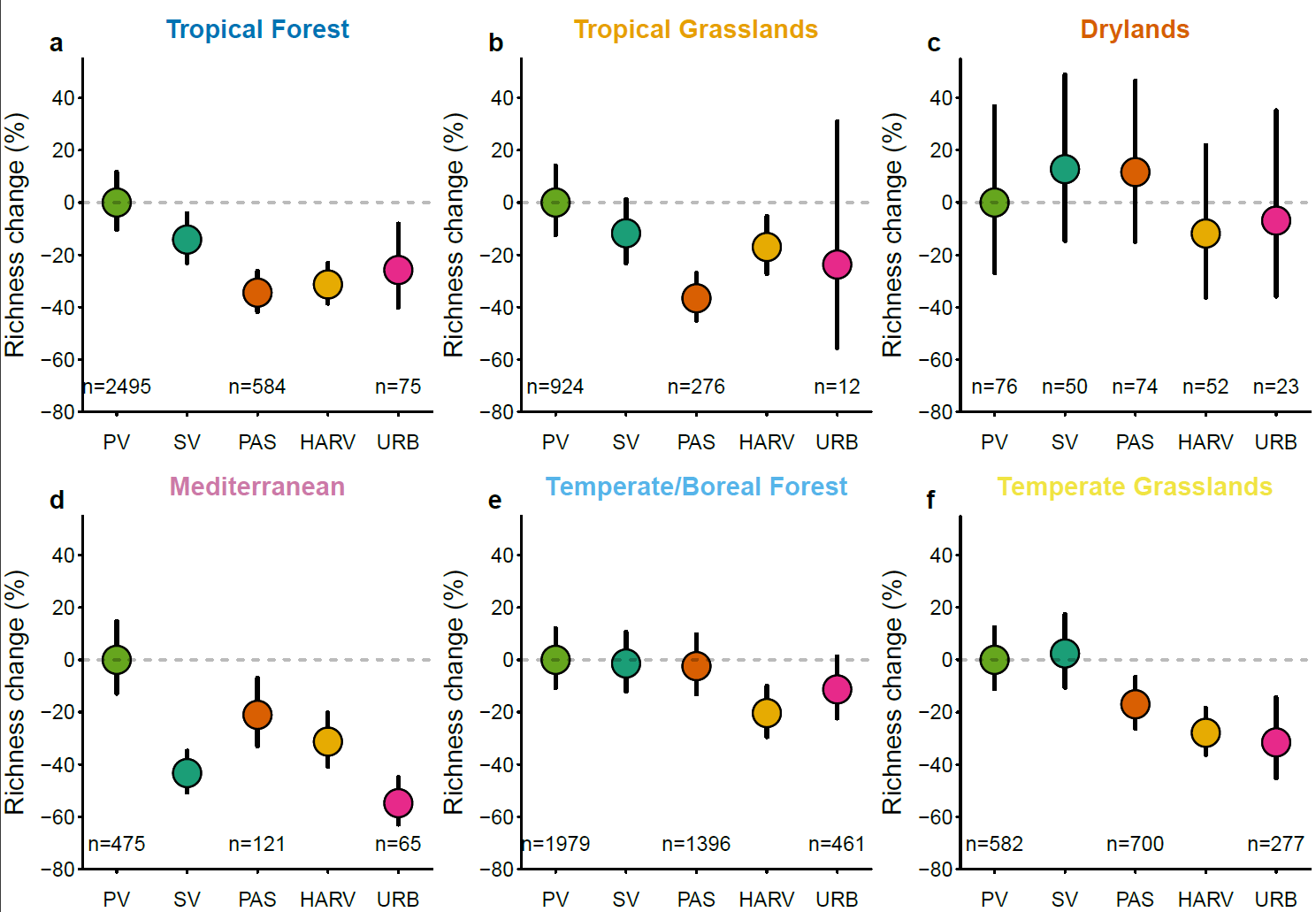

**Supplementary Figure 2.** Differences in total sampled species richness among land use types across different biomes. a) Tropical forest; b) Tropical grasslands; c) Drylands; d) Mediterranean; e) Temperate/Boreal forest; and f) Temperate/Montane grasslands. Plots show the percentage change in species richness compared to primary vegetation (PV), in secondary vegetation (SV), pasture (PAS), areas of harvested agriculture (woody plantations and herbaceous croplands) (HARV) and urban areas (URB). Error bars show 95% confidence intervals. Sample sizes at the bottom of each panel refer to the number of sites in each combination of land use and biome. In total 21986 sites from 637 studies were used. The final model plotted here had an R^2^_conditional_ of 0.61 and an R^2^_marginal_ of 0.024.

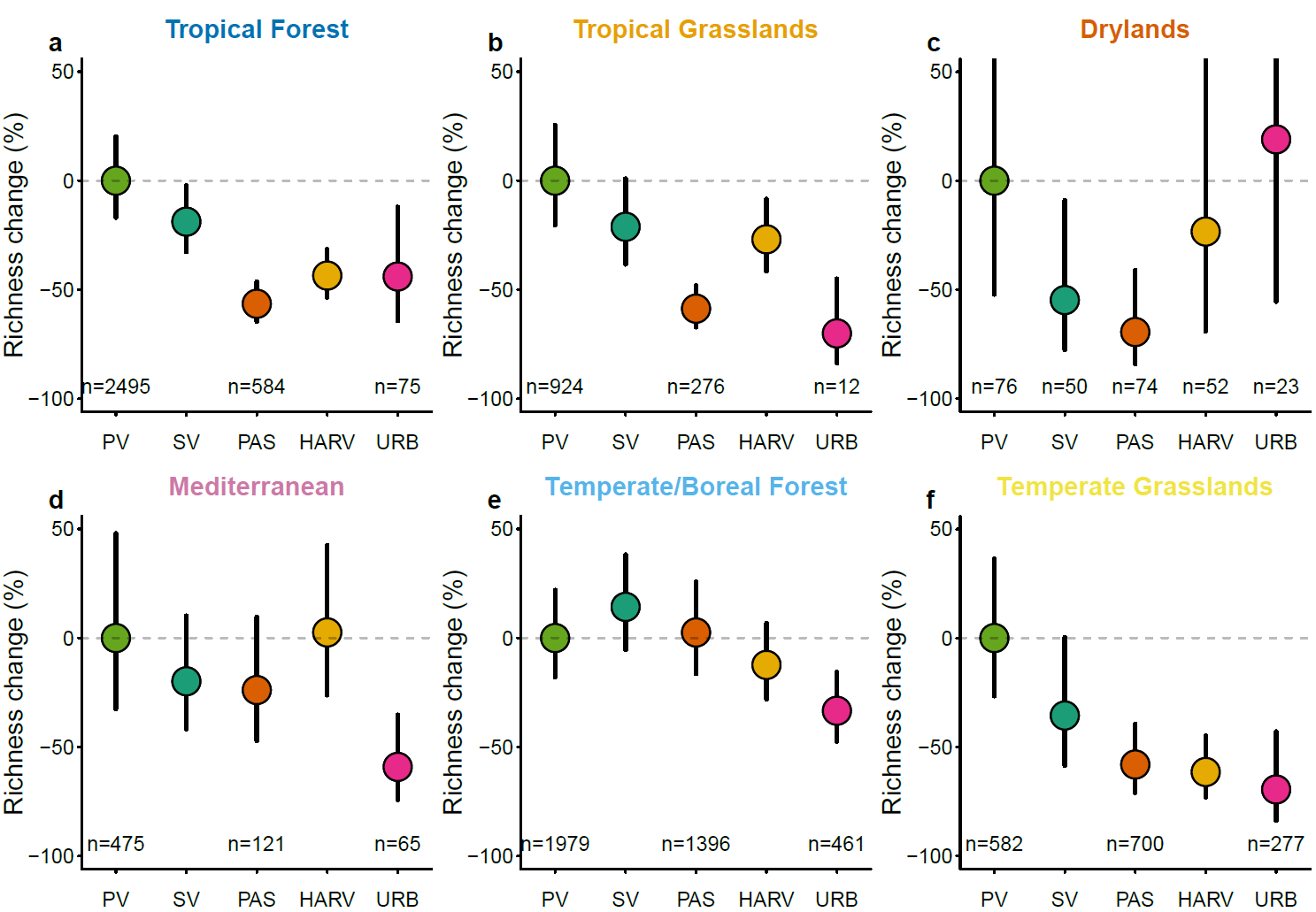

**Supplementary Figure 3.** Differences in total sampled rarity-weighted species richness among land use types across different biomes. a) Tropical forest; b) Tropical grasslands; c) Drylands; d)Mediterranean ; e) Temperate/Boreal forest; and f) Temperate/Montane grasslands. Plots show the percentage change in species richness compared to primary vegetation (PV), in secondary vegetation (SV), pasture (PAS), areas of harvested agriculture (woody plantations and herbaceous croplands) (HARV) and urban areas (URB). Error bars show 95% confidence intervals. Sample sizes at the bottom of each panel refer to the number of sites in each combination of land use and biome. In total 15198 sites from 403 studies were used. The final model plotted here had an R^2^_conditional_ of 0.63 and an R^2^_marginal_ of 0.043

**Supplementary Figure 4.** Proportional species richness (a, b, e) and range weighted richness change (c, d, e) that would occur within a cell under crop use (a, c), pasture use (b, d), or 1° of climate warming (e). The distributions of cells containing cropland (a, c) are taken from SPAM^1^ (cropland with exception of fodder crops) and EarthStat^2^(fodder crops). The distributions of cells containing pasture (b,c) are taken from Ramankutty, et al. ^4^. (See separate file “SupplementaryFigure4.pdf”.)

**Supplementary Figure 5.** The production (blue) and consumption (red) footprints of each individual food-related product in terms of a) land area, b) land-driven species richness loss, c) land-driven rarity-weighted richness loss, d) GHG-driven biodiversity loss split by emissions type: carbon dioxide (dark blue/red), methane (mid blue/red), nitrous oxide (light blue/red) (right-hand axis – species richness; left hand axis – rarity-weighted richness), e) the ratio of land-driven species richness loss to GHG-driven species richness loss and f) the ratio of land-driven rarity-weighted richness loss to GHG-driven rarity-weighted richness loss. AT = Austria, AU = Australia, BE = Belgium, BG = Bulgaria, BR = Brazil, CA = Canada, CH = Switzerland, CN = China, CY = Cyprus, CZ = Czech Republic, DE = Germany, DK = Denmark, EE = Estonia, ES = Spain, FI = Finland, GB = United Kingdom, HR = Croatia, HU = Hungary, ID = Indonesia, IE = Ireland, IN = India, IT = Italy, JP = Japan, KR = South Korea, LT = Lithuania, LV = Latvia, MT = Malta, MX = Mexico, NL = Netherlands, NO = Norway, PL = Poland, PT = Portugal, RO = Romania, RU = Russia, SE = Sweden, SI = Slovenia, SK = Slovakia, TR = Turkey, TW = Taiwan, US = United States, WA = RoW Asia & Pacific, WE = RoW Europe, WF = RoW Africa, WL = RoW CS America, WM = RoW Middle East, ZA = South Africa. (See separate file “SupplementaryFigure5.pdf”.)

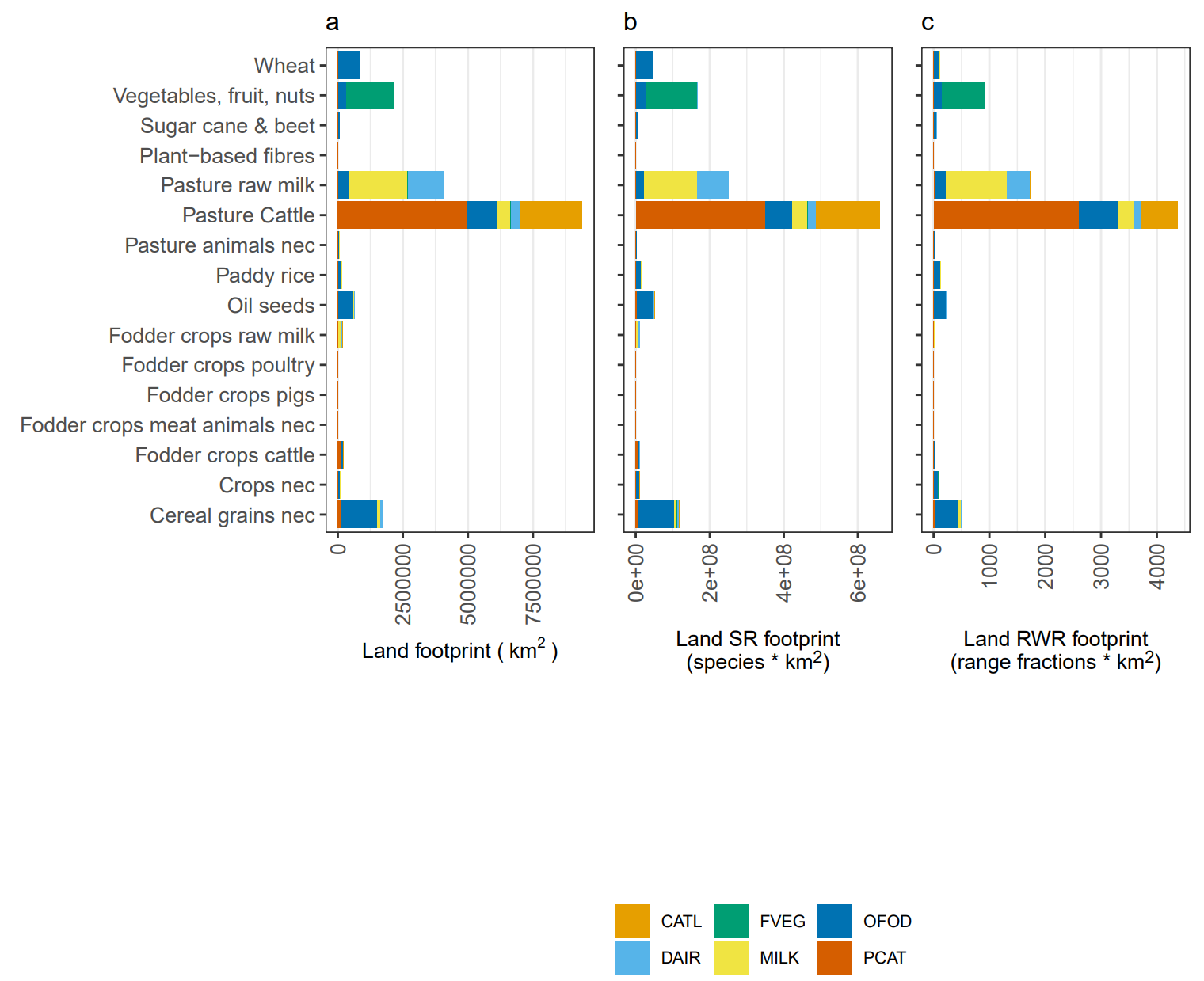

**Supplementary Figure 6.** The distribution of production-based footprints of a) land-use, b) land-driven species richness and c) land-driven rarity weighted richness across the EXIOBASE land-use categories for 6 high-footprint food sectors (CATL = cattle, DAIR = dairy products, FVEG = fruit/vegetables/nuts, MILK = raw milk, OFOD = food products nec, PCAT = products of cattle).

**Supplementary Figure 7.** Individual products’ production footprints per km^2^ for each EXIOBASE region for a) land use, b) land-driven species richness loss, c) land-driven rarity-weighted richness loss and d) GHG-driven biodiversity loss split by emissions type: carbon dioxide (black), methane (dark grey), nitrous oxide (light grey) (right-hand axis – species richness; left-hand axis – rarity-weighted richness). AT = Austria, AU = Australia, BE = Belgium, BG = Bulgaria, BR = Brazil, CA = Canada, CH = Switzerland, CN = China, CY = Cyprus, CZ = Czech Republic, DE = Germany, DK = Denmark, EE = Estonia, ES = Spain, FI = Finland, GB = United Kingdom, HR = Croatia, HU = Hungary, ID = Indonesia, IE = Ireland, IN = India, IT = Italy, JP = Japan, KR = South Korea, LT = Lithuania, LV = Latvia, MT = Malta, MX = Mexico, NL = Netherlands, NO = Norway, PL = Poland, PT = Portugal, RO = Romania, RU = Russia, SE = Sweden, SI = Slovenia, SK = Slovakia, TR = Turkey, TW = Taiwan, US = United States, WA = RoW Asia & Pacific, WE = RoW Europe, WF = RoW Africa, WL = RoW CS America, WM = RoW Middle East, ZA = South Africa. (See separate file “SupplementaryFigure7.pdf”.)

**Supplementary Figure 8.** Consumption-based footprints for individual products for each EXIOBASE region for a) land use, b) land-driven species richness loss, c) land-driven rarity-weighted richness loss and d) GHG-driven biodiversity loss (right-hand axis – species richness; left-hand axis – rarity-weighted richness). AT = Austria, AU = Australia, BE = Belgium, BG = Bulgaria, BR = Brazil, CA = Canada, CH = Switzerland, CN = China, CY = Cyprus, CZ = Czech Republic, DE = Germany, DK = Denmark, EE = Estonia, ES = Spain, FI = Finland, GB = United Kingdom, HR = Croatia, HU = Hungary, ID = Indonesia, IE = Ireland, IN = India, IT = Italy, JP = Japan, KR = South Korea, LT = Lithuania, LV = Latvia, MT = Malta, MX = Mexico, NL = Netherlands, NO = Norway, PL = Poland, PT = Portugal, RO = Romania, RU = Russia, SE = Sweden, SI = Slovenia, SK = Slovakia, TR = Turkey, TW = Taiwan, US = United States, WA = RoW Asia & Pacific, WE = RoW Europe, WF = RoW Africa, WL = RoW CS America, WM = RoW Middle East, ZA = South Africa. (See separate file “SupplementaryFigure8.pdf”.)

**Supplementary Figure 9.** Regions’ net import/export footprints for a) land use, b) land-driven species richness loss, c) land-driven rarity-weighted richness loss and d) GHG-driven richness loss. AT = Austria, AU = Australia, BE = Belgium, BG = Bulgaria, BR = Brazil, CA = Canada, CH = Switzerland, CN = China, CY = Cyprus, CZ = Czech Republic, DE = Germany, DK = Denmark, EE = Estonia, ES = Spain, FI = Finland, GB = United Kingdom, HR = Croatia, HU = Hungary, ID = Indonesia, IE = Ireland, IN = India, IT = Italy, JP = Japan, KR = South Korea, LT = Lithuania, LV = Latvia, MT = Malta, MX = Mexico, NL = Netherlands, NO = Norway, PL = Poland, PT = Portugal, RO = Romania, RU = Russia, SE = Sweden, SI = Slovenia, SK = Slovakia, TR = Turkey, TW = Taiwan, US = United States, WA = RoW Asia & Pacific, WE = RoW Europe, WF = RoW Africa, WL = RoW CS America, WM = RoW Middle East, ZA = South Africa. (See separate file “SupplementaryFigure9.pdf”.)

**Supplementary Figure 10.** The percentage of a region’s footprint that is imported for a) land use, b) land-driven species richness loss, c) land-driven rarity-weighted richness loss and d) GHG-driven richness loss. AT = Austria, AU = Australia, BE = Belgium, BG = Bulgaria, BR = Brazil, CA = Canada, CH = Switzerland, CN = China, CY = Cyprus, CZ = Czech Republic, DE = Germany, DK = Denmark, EE = Estonia, ES = Spain, FI = Finland, GB = United Kingdom, HR = Croatia, HU = Hungary, ID = Indonesia, IE = Ireland, IN = India, IT = Italy, JP = Japan, KR = South Korea, LT = Lithuania, LV = Latvia, MT = Malta, MX = Mexico, NL = Netherlands, NO = Norway, PL = Poland, PT = Portugal, RO = Romania, RU = Russia, SE = Sweden, SI = Slovenia, SK = Slovakia, TR = Turkey, TW = Taiwan, US = United States, WA = RoW Asia & Pacific, WE = RoW Europe, WF = RoW Africa, WL = RoW CS America, WM = RoW Middle East, ZA = South Africa. (See separate file “SupplementaryFigure10.pdf”.)

**Supplementary Figure 11.** Biodiversity impacts embodied within all food-related products for three footprint types: a) land-driven species richness, b) land-driven rarity-weighted richness, c) GHG-driven species richness. The left axis shows the trade region where the impacts are produced and the right axis shows the trade region in which they are consumed. The width of the flows shows the magnitude of the footprints. Domestic production and consumption impacts are included. (See separate file “SupplementaryFigure11.pdf”.)

**Supplementary Figure 12.** Biodiversity impacts embodied within trade between continental regions for three footprint types: a, b) land-driven species richness, c, d) land-driven rarity-weighted richness, e, f) GHG-driven species richness for a, c, e) animal-derived food products and b, d, f) plant-derived food products. The left axis shows the trade region where the impacts are produced and the right axis shows the trade region in which they are consumed. The width of the flows shows the magnitude of the footprints. The impacts shown here represent 8%, 17%, 15%, 21%, 6%, 12% respectively of the total impact. It can be seen that CS America dominates the export of animal-derived, land-driven biodiversity footprints whereas the traded land-driven biodiversity footprint of plant-derived products is more evenly distributed across the tropics. Western Europe is the greatest importer of biodiversity loss embedded in crops, with the largest fraction of its species richness footprint coming from Africa but the largest fraction of its rarity-weighted footprint from CS America. The total animal-derived import/export biodiversity footprint is higher than its plant-derived counterpart for land-driven footprints but lower for the GHG-driven biodiversity footprint. (See separate file “SupplementaryFigure12.pdf”.)

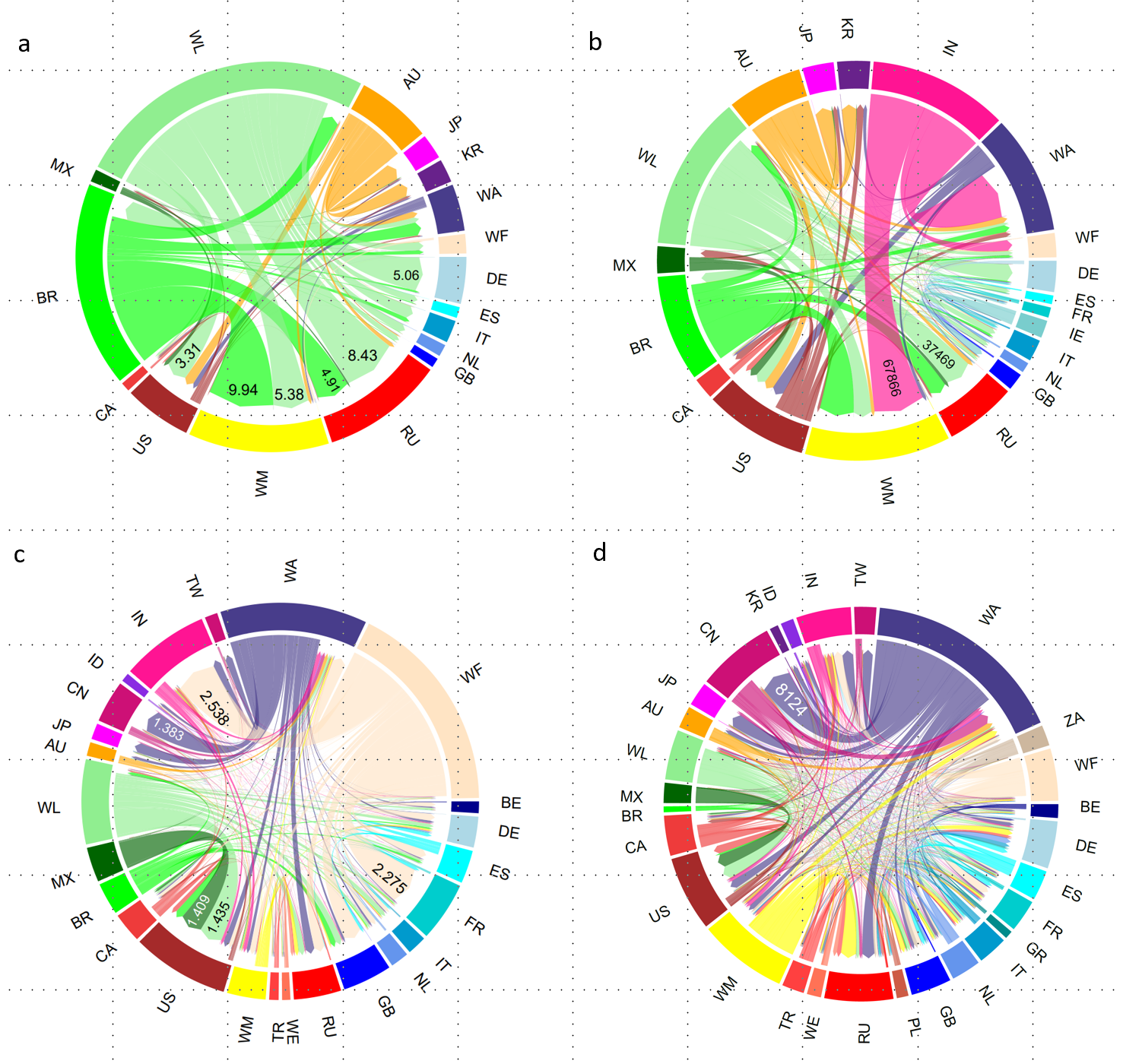

**Supplementary Figure 13.** Embedded land-driven species richness loss (a, c) and GHG-driven species richness loss (b, d) footprint in the international trade of processed cattle products (a, b) and fruit/vegetables/nuts (c, d), two of the products with the highest footprints for both land-driven and GHG-driven biodiversity loss. Land-driven footprints are in units of species × km^2^ x 10^6^; GHG-driven footprints are in units of species x km^2^. Regions with footprints < 0.5% of the circumference of the plot are not shown. AU = Australia; BE = Belgium, BR = Brazil, CA = Canada, CN = China, DE = Germany, DK = Denmark, ES = Spain, FR = France, ID = Indonesia, IE = Ireland, IN = India, IT = Italy, JP = Japan, MX = Mexico, NL = the Netherlands, GB = United Kingdom, GR = Greece, KR = South Korea, PL = Poland, RU = Russia, SE = Sweden, TR = Turkey, TW = Taiwan, US = United States, WA = RoW Asia, WE = RoW Europe, WF = RoW Africa, WL = RoW Central & South America, WM = RoW Middle East, ZA = South Africa. The trade flows show two striking differences. (1) The origin of the land-driven species richness footprint differs between cattle products and fruit/vegetables/nuts. Almost half of the land-driven species richness loss from cattle products is exported from only three regions: RoW CS America, Brazil and Australia. In contrast, the land-driven biodiversity footprint of fruit/vegetables/nuts comes from a wider spread of regions, with the largest share originating from RoW Africa. (2) Cattle products’ land-driven and GHG-driven species richness footprint patterns differ markedly, with GHG-driven export footprints being more evenly distributed across regions than land-driven. The United States, India and W European regions, for example, have very low land-driven export footprints but sizable GHG-driven export footprints. The difference between India’s land-driven and GHG-driven export footprints are particularly striking.

A similar theme is seen for fruit/vegetables/nuts with W European regions mostly importing land-driven species richness footprint, but both importing and exporting GHG-driven richness footprint.

**Supplementary Tables**

**Supplementary Table 1.** The food-related products included in the analysis. The EXIOBASE product code is given in brackets in the first column. Examples of products are given in the second column.

| EXIOBASE product name | Examples of products |
| --- | --- |
| Paddy rice (PARI) | Paddy rice |
| Wheat (WHEA) | Wheat |
| Cereal grains nec (OCER) | Maize, barley, rye |
| Vegetables, fruit, nuts (FVEG) | Onions, avocados, cashews in shell |
| Oil seeds (OILS) | Soy beans, ground nuts, palm nuts and kernels |
| Sugar cane, sugar beet (SUGB) | Sugar cane, sugar beet |
| Crops nec (OTCR) | Potatoes, yams, coffee, tea, cocoa, spices |
| Cattle (CATL) | Live cattle, buffalo, other bovine animals |
| Pigs (PIGS) | Live pigs |
| Poultry (PLTR) | Live chickens, turkeys, geese, ducks |
| Meat animals nec (OMEA) | Live goats, sheep, horses |
| Animal products nec (OANP) | Eggs, honey, snails |
| Raw milk (MILK) | Raw milk of cattle, sheep, goats, camels |
| Fish and other fishing products; services incidental of fishing (FISH) | Live fish not for human consumption, fish live fresh or chilled for human consumption, crustaceans, marine molluscs, aquatic invertebrates |
| Chemical and fertilizer minerals, salt and other mining and quarrying products nec (CHMF) | Natural calcium phosphate, sodium chloride in sea water |
| Products of meat cattle (PCAT) | Meat of bovine animals - fresh, chilled, frozen, smoked |
| Products of meat pigs (PPIG) | Meat of pigs - fresh, chilled, frozen, smoked |
| Products of meat poultry (PPLT) | Meat of poultry - fresh, chilled, frozen, smoked |
| Meat products nec (POME) | Meat of sheep, goats, horses - fresh, chilled, frozen, smoked |
| Products of vegetable oils and fats (VOIL) | Lard, fish-liver oil, palm oil, olive oil, coconut oil |
| Dairy products (DAIR) | Processed liquid milk, cheese, yoghurt, butter, ice-cream |
| Processed rice (RICE) | Semi or wholly milled rice, husked rice |
| Sugar (SUGR) | Cane sugar, beet sugar, refined sugar |
| Food products nec (OFOD) | Frozen vegetables, fruit juices, wheat flour, soups, sandwiches, pizza, plant-based meat substitutes, chocolate, pasta, roasted coffee, processed spices |
| Beverages (BEVR) | Soft drinks, bottled water, wine, beer |
| Fish products (FSHP) | Frozen or dried fish, crustaceans and molluscs |
| N-fertiliser (NFER) | Urea, ammonium sulphate, ammonium nitrate |
| P- and other fertilizer (PFER) | Superphosphate, potassium chloride, ammonium chloride |
| Food waste for treatment: incineration (INCF) |  |
| Food waste for treatment: biogasification and land application (BIOF) |  |
| Food waste for treatment: composting and land application (COMF) |  |
| Food waste for treatment: waste water treatment (WASF) |  |
| Food waste for treatment: landfill (LANF) |  |

**Supplementary Table 2.** The aggregation of The Nature Conservancy’s biomes^14^ into the broad biome groupings used in the models of biodiversity responses to land use.

| Biome – The Nature Conservancy | Biome – PREDICTS |
| --- | --- |
| Tropical and subtropical moist broadleaf forests | Tropical forest |
| Tropical and subtropical dry broadleaf forests | Tropical forest |
| Tropical and subtropical coniferous forests | Tropical forest |
| Temperate broadleaf and mixed forests | Temperate forest |
| Temperate conifer forests | Temperate forest |
| Boreal forests/taiga | Temperate forest |
| Tropical and subtropical grasslands, savannas and shrublands | Tropical grasslands |
| Temperate grasslands, savannas and shrublands | Temperate grasslands |
| Montane grasslands and shrublands | Temperate grasslands |
| Mediterranean forests, woodlands and scrub | Mediterranean |
| Deserts and xeric shrublands | Drylands |
| Tundra | Excluded |
| Mangroves | Excluded |
| Flooded grasslands and savannas | Excluded |
| Inland water | Excluded |
| Rock and ice | Excluded |

**Supplementary Table 3.** The allocation of crops and pasture to EXIOBASE’s and PREDICTS’ land-use categories. Maps were taken from SPAM^1^ with the exception of those with an asterisk which came from EarthStat^2^. In some instances, crops are allocated to more than one EXIOBASE land-use category.

| Crop/pasture | Land use category - EXIOBASE | Land use category - PREDICTS |
| --- | --- | --- |
| Wheat | Cropland - Paddy rice | Harvested agriculture |
| Rice | Cropland - Wheat | Harvested agriculture |
| Maize | Cropland - Other cereals nec | Harvested agriculture |
| Barley | Cropland - Other cereals nec | Harvested agriculture |
| Pearl millet | Cropland - Other cereals nec | Harvested agriculture |
| Small millet | Cropland - Other cereals nec | Harvested agriculture |
| Sorghum | Cropland - Other cereals nec | Harvested agriculture |
| Other cereals | Cropland - Other cereals nec | Harvested agriculture |
| Potato | Cropland - Other crops nec | Harvested agriculture |
| Sweet potato | Cropland - Other crops nec | Harvested agriculture |
| Yams | Cropland - Other crops nec | Harvested agriculture |
| Cassava | Cropland - Other crops nec | Harvested agriculture |
| Other roots | Cropland – Other crops nec | Harvested agriculture |
| Bean | Cropland - Vegetables, fruit, nuts; Cropland - Other crops nec | Harvested agriculture |
| Chickpea | Cropland - Vegetables, fruit, nuts | Harvested agriculture |
| Cow pea | Cropland - Vegetables, fruit, nuts | Harvested agriculture |
| Pigeon pea | Cropland - Vegetables, fruit, nuts | Harvested agriculture |
| Lentil | Cropland - Vegetables, fruit, nuts | Harvested agriculture |
| Other pulses | Cropland - Vegetables, fruit, nuts | Harvested agriculture |
| Soy bean | Cropland - Oil seeds | Harvested agriculture |
| Groundnut | Cropland - Oil seeds | Harvested agriculture |
| Coconut | Cropland - Oil seeds | Harvested agriculture |
| Banana | Cropland - Vegetables, fruit, nuts | Harvested agriculture |
| Plantain | Cropland - Vegetables, fruit, nuts | Harvested agriculture |
| Tropical fruit | Cropland - Vegetables, fruit, nuts | Harvested agriculture |
| Temperate fruit | Cropland - Vegetables, fruit, nuts | Harvested agriculture |
| Vegetables | Cropland - Vegetables, fruit, nuts | Harvested agriculture |
| Oil palm | Cropland - Oil seeds | Harvested agriculture |
| Sunflower | Cropland - Oil seeds | Harvested agriculture |
| Rape seed | Cropland - Oil seeds | Harvested agriculture |
| Sesame seed | Cropland - Oil seeds | Harvested agriculture |
| Other oil crops | Cropland - Oil seeds | Harvested agriculture |
| Sugar cane | Cropland - Sugar cane, sugar beet | Harvested agriculture |
| Sugar beet | Cropland - Sugar cane, sugar beet | Harvested agriculture |
| Cotton | Cropland - Oil seeds; Cropland - Plant-based fibers | Harvested agriculture |
| Other fibre crops | Cropland - Plant-based fibers | Harvested agriculture |
| Arabica coffee | Cropland - Other crops nec | Harvested agriculture |
| Robusta coffee | Cropland - Other crops nec | Harvested agriculture |
| Cocoa | Cropland - Other crops nec | Harvested agriculture |
| Tea | Cropland - Other crops nec | Harvested agriculture |
| Rest of Crops | Cropland - Other crops nec | Harvested agriculture |
| Alfalfa* | Cropland – Fodder crops cattle; Cropland – Fodder crops meat animals; Cropland – fodder crops pigs; Cropland – fodder crops poultry; Cropland – fodder crops raw milk | Harvested agriculture |
| Beet for forage* | Cropland – Fodder crops cattle; Cropland – Fodder crops meat animals; Cropland – fodder crops pigs; Cropland – fodder crops poultry; Cropland – fodder crops raw milk | Harvested agriculture |
| Cabbage for forage* | Cropland – Fodder crops cattle; Cropland – Fodder crops meat animals; Cropland – fodder crops pigs; Cropland – fodder crops poultry; Cropland – fodder crops raw milk | Harvested agriculture |
| Carrot for forage* | Cropland – Fodder crops cattle; Cropland – Fodder crops meat animals; Cropland – fodder crops pigs; Cropland – fodder crops poultry; Cropland – fodder crops raw milk | Harvested agriculture |
| Clover* | Cropland – Fodder crops cattle; Cropland – Fodder crops meat animals; Cropland – fodder crops pigs; Cropland – fodder crops poultry; Cropland – fodder crops raw milk | Harvested agriculture |
| Forage nec* | Cropland – Fodder crops cattle; Cropland – Fodder crops meat animals; Cropland – fodder crops pigs; Cropland – fodder crops poultry; Cropland – fodder crops raw milk | Harvested agriculture |
| Grass nec* | Cropland – Fodder crops cattle; Cropland – Fodder crops meat animals; Cropland – fodder crops pigs; Cropland – fodder crops poultry; Cropland – fodder crops raw milk | Harvested agriculture |
| Maize for forage* | Cropland – Fodder crops cattle; Cropland – Fodder crops meat animals; Cropland – fodder crops pigs; Cropland – fodder crops poultry; Cropland – fodder crops raw milk | Harvested agriculture |
| Mixed grass* | Cropland – Fodder crops cattle; Cropland – Fodder crops meat animals; Cropland – fodder crops pigs; Cropland – fodder crops poultry; Cropland – fodder crops raw milk | Harvested agriculture |
| Oil seed for forage* | Cropland – Fodder crops cattle; Cropland – Fodder crops meat animals; Cropland – fodder crops pigs; Cropland – fodder crops poultry; Cropland – fodder crops raw milk | Harvested agriculture |
| Rye for forage* | Cropland – Fodder crops cattle; Cropland – Fodder crops meat animals; Cropland – fodder crops pigs; Cropland – fodder crops poultry; Cropland – fodder crops raw milk | Harvested agriculture |
| Sorghum for forage* | Cropland – Fodder crops cattle; Cropland – Fodder crops meat animals; Cropland – fodder crops pigs; Cropland – fodder crops poultry; Cropland – fodder crops raw milk | Harvested agriculture |
| Swede for forage* | Cropland – Fodder crops cattle; Cropland – Fodder crops meat animals; Cropland – fodder crops pigs; Cropland – fodder crops poultry; Cropland – fodder crops raw milk | Harvested agriculture |
| Turnip for forage* | Cropland – Fodder crops cattle; Cropland – Fodder crops meat animals; Cropland – fodder crops pigs; Cropland – fodder crops poultry; Cropland – fodder crops raw milk | Harvested agriculture |
| Vegetable for forage* | Cropland – Fodder crops cattle; Cropland – Fodder crops meat animals; Cropland – fodder crops pigs; Cropland – fodder crops poultry; Cropland – fodder crops raw milk | Harvested agriculture |
| Pasture* | Permanent pastures – grazing cattle; Permanent pastures – grazing meat animals nec; Permanent pastures – grazing raw milk | Livestock pasture |

**Supplementary Table 4.** Modelled differences in total sampled species richness among land-use types. All values are expressed as a percentage change relative to primary vegetation as the baseline. Numbers in parentheses are the lower and upper bounds of the 95% confidence limits

| **Biome** | **Primary vegetation** | **Secondary vegetation** | **Harvested agriculture** | **Livestock pasture** | **Urban area** |
| --- | --- | --- | --- | --- | --- |
| **Tropical forest** | 0  (-10.3\|11.5) | -14.2  (-23.1\|-4.2) | -31.1  (-38.5\|-23.2) | -34.3  (-41.7\|-26.2) | -25.8  (-40.0\|-8.2) |
| **Tropical grassland** | 0  (-12.3\|14.0) | -11.7  (-23.1\|1.3) | -17.0  (-27.2\|-5.3) | -36.6  (-45.0\|-26.9) | -23.7  (-55.5\|30.9) |
| **Drylands** | 0  (-26.9\|36.8) | 12.8  (-14.6\|48.9) | -11.8  (-36.2\|21.9) | 11.6  (-14.9\|46.5) | -6.9  (-35.9\|35.1) |
| **Mediterranean** | 0  (-12.8\|14.7) | -43.2  (-50.6\|-34.9) | -31.3  (-40.8\|-20.3) | -21.0  (-33.0\|-6.9) | -54.8  (-62.9\|-44.9) |
| **Temperate/boreal forest** | 0  (-10.7\|12.0) | -1.4  (-12.1\|10.7) | -20.3  (-29.4\|-10.1) | -2.4  (-13.1\|9.6) | -11.3  (-22.3\|1.2) |
| **Temperate/montane grasslands** | 0  (-11.0\|12.4) | 2.4  (-10.5\|17.3) | -27.9  (-36.2\|-18.4) | -17.0  (-26.1\|-6.7) | -31.6  (-45.2\|-14.4) |

**Supplementary Table 5.** Modelled differences in total sampled rarity-weighted species richness among land-use types. All values are expressed as a percentage change relative to primary vegetation as the baseline. Numbers in parentheses are the lower and upper bounds of the 95% confidence limits

| **Biome** | **Primary vegetation** | **Secondary vegetation** | **Harvested agriculture** | **Livestock pasture** | **Urban area** |
| --- | --- | --- | --- | --- | --- |
| **Tropical forest** | 0  (-16.9\|20.3) | -18.8  (-32.6\|-2.2) | -43.5  (-53.4\|-31.4) | -56.4  (-64.6\|-46.4) | -43.9  (-64.3\|-12.1) |
| **Tropical grassland** | 0  (-20.3\|25.5) | -21.1  (-38.4\|0.9) | -26.9  (-41.6\|-8.5) | -58.7  (-67.1\|-48.0) | -70.0  (-83.7\|44.0) |
| **Drylands** | 0  (-52.2\|109.1) | -54.7  (-77.5\|-8.9) | -23.3  (-69.0\|89.8) | -69.4  (-84.1\|-41.0) | 19.0  (-55.4\|217.3) |
| **Mediterranean** | 0  (-32.4\|48.0) | -19.8  (-41.7\|10.2) | 2.5  (-26.2\|42.6) | -23.9  (-47.1\|9.7) | -59.1  (-74.1\|-35.3) |
| **Temperate/boreal forest** | 0  (-18.0\|22.0) | 14.3  (-5.5\|38.2) | -12.3  (-28.0\|6.7) | 2.5  (-16.4\|25.8) | -33.3  (-47.4\|-15.6) |
| **Temperate/montane grasslands** | 0  (-26.6\|36.3) | -35.3  (-58.6\|0.3) | -61.5  (-73.0\|-44.9) | -58.0  (-70.9\|-39.4) | -69.5  (-83.7\|-42.9) |

**Supplementary Table 6.** Characterisation factors for land-driven species richness change (species). (See Excel file “SupplementaryTables6-13.xlsx”.)

**Supplementary Table 7.** Characterisation factors for land-driven rarity-weighted richness change (range fractions). (See Excel file “SupplementaryTables6-13.xlsx”.)

**Supplementary Table 8.** Characterisation factors for GHG-driven species richness change (species). For the purposes of the analyses, the 19 GHG-driven factors listed were summed into a single characterization factor or a single factor for each gas (carbon dioxide, methane and nitrous oxide). (See Excel file “SupplementaryTables6-13.xlsx”.)

**Supplementary Table 9.** Characterisation factors for GHG-driven rarity-weighted richness change (range fractions). For the purposes of the analyses, the 19 GHG-driven factors listed were summed into a single characterization factor or a single factor for each gas (carbon dioxide, methane and nitrous oxide). (See Excel file “SupplementaryTables6-13.xlsx”.)

**Supplementary Table 10.** Areas of the EXIOBASE trade regions (km²) as calculated from ESRI country shapefiles^15^. (See Excel file “SupplementaryTables6-13.xlsx”.)

**Supplementary Table 11:** The 2011 populations of the EXIOBASE trade regions as estimated by CountryEconomy.com^16^ (Taiwan) and the World Bank^17^ (all other regions). (See Excel file “SupplementaryTables6-13.xlsx”.)

**Supplementary Table 12.** Production and consumption footprints of land use (km²), land-driven species richness (species × km²), land-driven rarity-weighted richness (range fractions × km²), GHG-driven species richness (species × km²) (total, CO2, CH4 and N2O) and GHG-driven rarity-weighted richness (range fractions × km²) (total, CO2, CH4 and N2O) for all regions and all food-related products. (See Excel file “SupplementaryTables6-13.xlsx”.)

**Supplementary Table 13.** Production footprints of land use (km²), land-driven species richness (species × km²), land-driven rarity-weighted richness (range fractions × km²), GHG-driven species richness (species × km²) (total, CO2, CH4 and N2O) and GHG-driven rarity-weighted richness (range fractions × km²) (total, CO2, CH4 and N2O) for world regions and all food-related products. (See Excel file “SupplementaryTables6-13.xlsx”.)

Supplementary Table 14: Production per km2 footprints of land use (km²), land-driven species richness (species x km²), land-driven rarity-weighted richness (range fractions x km²), GHG-driven species richness (species x km²) (total, CO2, CH4 and N2O) and GHG-driven rarity-weighted richness (range fractions x km²) (total, CO2, CH4 and N2O) for world regions and all food-related products

Supplementary Table 15: Consumption per capita footprints of land use (km²), land-driven species richness (species x km²), land-driven rarity-weighted richness (range fractions x km²), GHG-driven species richness (species x km²) (total, CO2, CH4 and N2O) and GHG-driven rarity-weighted richness (range fractions x km²) (total, CO2, CH4 and N2O) for world regions and all food-related products

**Supplementary Table 16.** The ten highest per-capita consumption-based footprints at a product-region level for the different footprint types. (The values of GHG-driven rarity-weighted richness are not given since they are directly proportional to the values of the GHG-driven species richness footprints). Each cell shows the consumption-based footprint per capita, the food product category and the region of consumption. CATL = cattle, DAIR = dairy products, FVEG = fruit/vegetables/nuts, MILK = raw milk, OFOD = food product nec (e.g. soups, pizza, sauces), OILS = oil seeds, OTCR = crops nec (e.g. coffee, cocoa, spices), PCAT = products of meat cattle.

| Land use (km^2^) | Land-driven species richness (species x km^2^) | Land-driven rarity-weighted richness (range fractions x km^2^) | GHG-driven species richness (species x km^2^) |
| --- | --- | --- | --- |
| 4.69E-03, PCAT, US | 1.15, PCAT, Australia | 4.26E-06, PCAT, Australia | 1.02E-02, PCAT, Australia |
| 4.50E-03, DAIR, Australia | 6.97E-01, PCAT, Brazil | 2.71E-06, PCAT, RoW Central & South America | 4.50E-03, DAIR, Australia |
| 4.24E-03, OFOD, Australia | 3.72E-01, CATL, Brazil | 2.11E-06, PCAT, Brazil | 3.21E-03, PCAT, Brazil |
| 3.74E-03, PCAT, Brazil | 2.47E-01, FVEG, Luxembourg | 2.05E-06, OFOD, RoW Central & South America | 2.57E-03, OFOD, RoW Central & South America |
| 3.54E-03, MILK, South Africa | 2.36E-01, OTCR, Luxembourg | 2.03E-06, OFOD, Mexico | 2.55E-03, CATL, Luxembourg |
| 3.43E-03, OFOD, Russia | 2.16E-01, OFOD, Australia | 2.02E-06, MILK, RoW Central & South America | 2.36E-03, PCAT, RoW Central & South America |
| 3.09E-03, PCAT, South Africa | 2.05E-01, OFOD, Taiwan | 1.99E-06, MILK, South Africa | 2.07E-03, OFOD, Netherlands |
| 3.04E-03, FVEG, Luxembourg | 2.03E-01, DAIR, Brazil | 1.89E-06, FVEG, Luxembourg | 2.02E-03, OFOD, Australia |
| 2.85E-03, OILS, Luxembourg | 1.85E-01, OFOD, Russia | 1.78E-06, OTCR, Luxembourg | 1.91E-03, OFOD, Belgium |
| 2.85E-03, CATL, Australia | 1.81E-01, DAIR, Australia | 1.74E-06, PCAT, South Africa | 1.89E-03, OTCR, Luxembourg |

**References**

1	International Food Policy Research Institute. *MapSPAM* (ed International Food Policy Research Institute) (Harvard Dataverse, 2019).

2	Monfreda, C., Ramankutty, N. & Foley, J. A. Farming the planet: 2. Geographic distribution of crop areas, yields, physiological types, and net primary production in the year 2000. *Global Biogeochemical Cycles* **22**, GB1022 (2008). https://doi.org:doi: 10.1029/2007GB002947

3	Marques, A. *et al.* Increasing impacts of land use on biodiversity and carbon sequestration driven by population and economic growth. *Nat Ecol Evol* **3**, 628-637 (2019). https://doi.org:10.1038/s41559-019-0824-3

4	Ramankutty, N., Evan, A. T., Monfreda, C. & Foley, J. A. Farming the planet: 1. Geographic distribution of global agricultural lands in the year 2000. *Global Biogeochemical Cycles* **22**, GB1003 (2008). https://doi.org:doi:10.1029/2007GB002952

5	raster: Geographic analysis and modeling with raster data. (R package version 2.0-12, 2012).

6	rgdal: Bindings for the 'Geospatial' Data Abstraction Library (R package version 1.5-23, 2021).

7	Hudson, L. N. *et al.* The database of the PREDICTS (Projecting Responses of Ecological Diversity In Changing Terrestrial Systems) project. *Ecol Evol* **7**, 145-188 (2017). https://doi.org:10.1002/ece3.2579

8	Newbold, T., Oppenheimer, P., Etard, A. & Williams, J. J. Tropical and Mediterranean biodiversity is disproportionately sensitive to land-use and climate change. *Nat Ecol Evol* **4**, 1630-1638 (2020). https://doi.org:10.1038/s41559-020-01303-0

9	Newbold, T. *et al.* Global effects of land use on local terrestrial biodiversity. *Nature* **520**, 45-50 (2015). https://doi.org:10.1038/nature14324

10	Newbold, T. Future effects of climate and land-use change on terrestrial vertebrate community diversity under different scenarios. *Proc Biol Sci* **285** (2018). https://doi.org:10.1098/rspb.2018.0792

11	IUCN. The IUCN Red List of Threatened Species. Version 2013.7 (2013).

12	BirdLife International. Bird species distribution maps of the world. Version 2.0	(http://www.birdlife.org/datazone/info/spcdownload, 2012).

13	Hijmans, R., Cameron, S. E., Parra, J. L., Jones, P. G. & Jarvis, A. Very high resolution interpolated climate surfaces for global land areas. *International Journal of Climatology* **25**, 1965-1978 (2005). https://doi.org:doi.org/10.1002/joc.1276

14	The Nature Conservancy. Terrestrial Ecoregions (2019). Date accessed: 11.11.2021

15	ESRI. Countries WGS84. https://hub.arcgis.com/datasets/a21fdb46d23e4ef896f31475217cbb08_1/data	(2015).

16	CountryEconomy.com. *https://countryeconomy.com/demography/population/taiwan?year=2011* 2021).

17	World Bank. Data Bank, https://databank.worldbank.org, accessed 26 October 2021	(2021).
