## Supplementary figures and images for "Impacts of global food supply on biodiversity via land use and climate change"

### Supplementary Figure 4

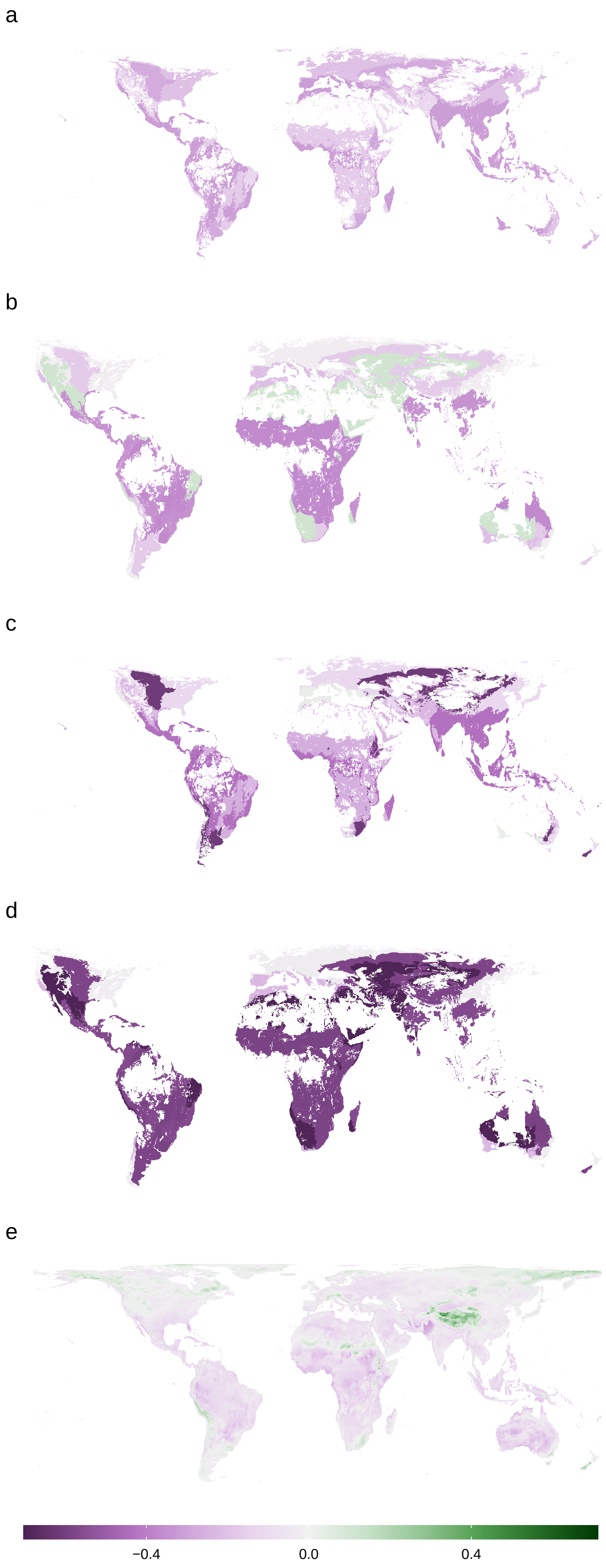

### Supplementary Figure 9

Net imported footprint: total food−related products

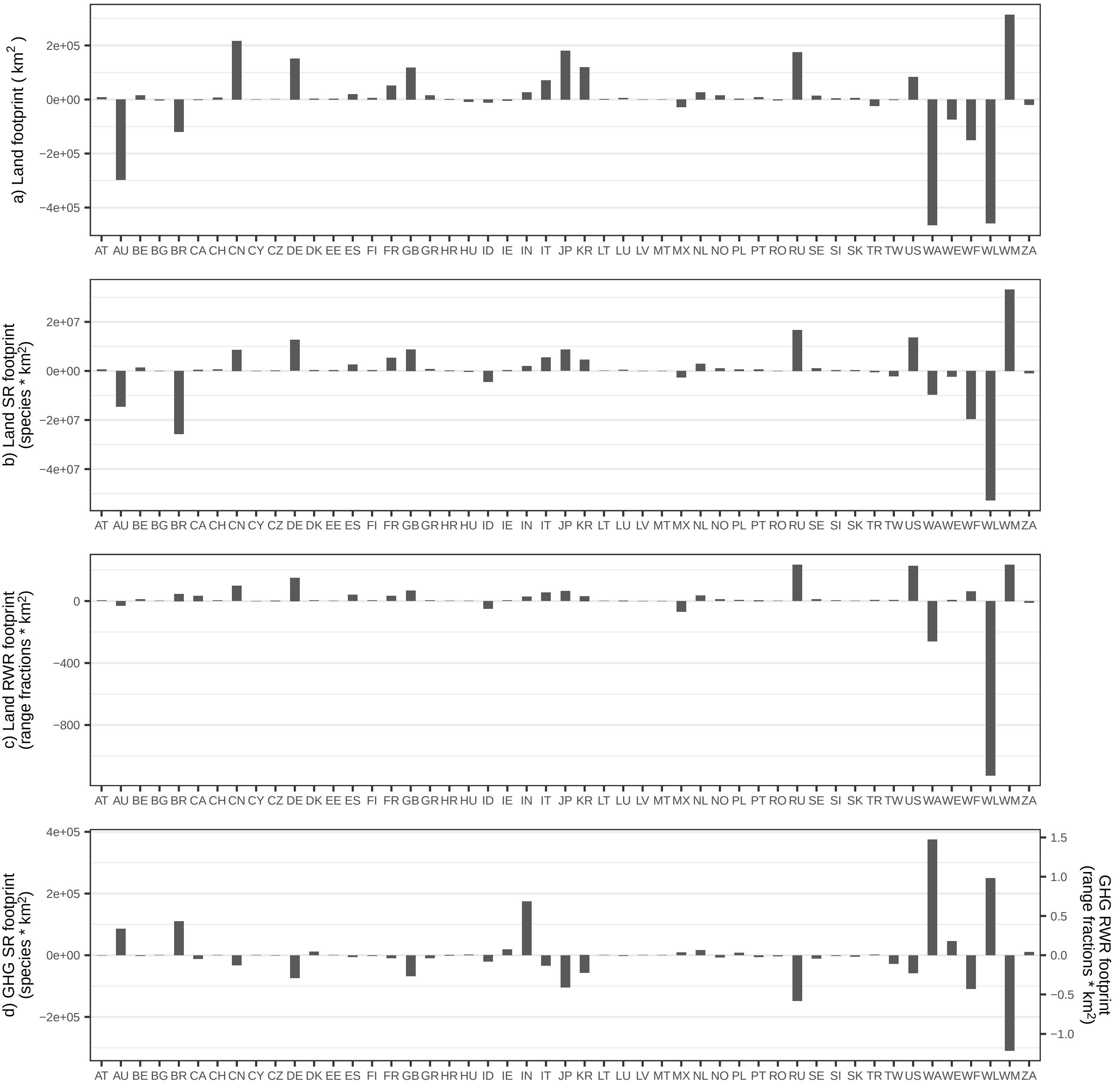

### Supplementary Figure 10

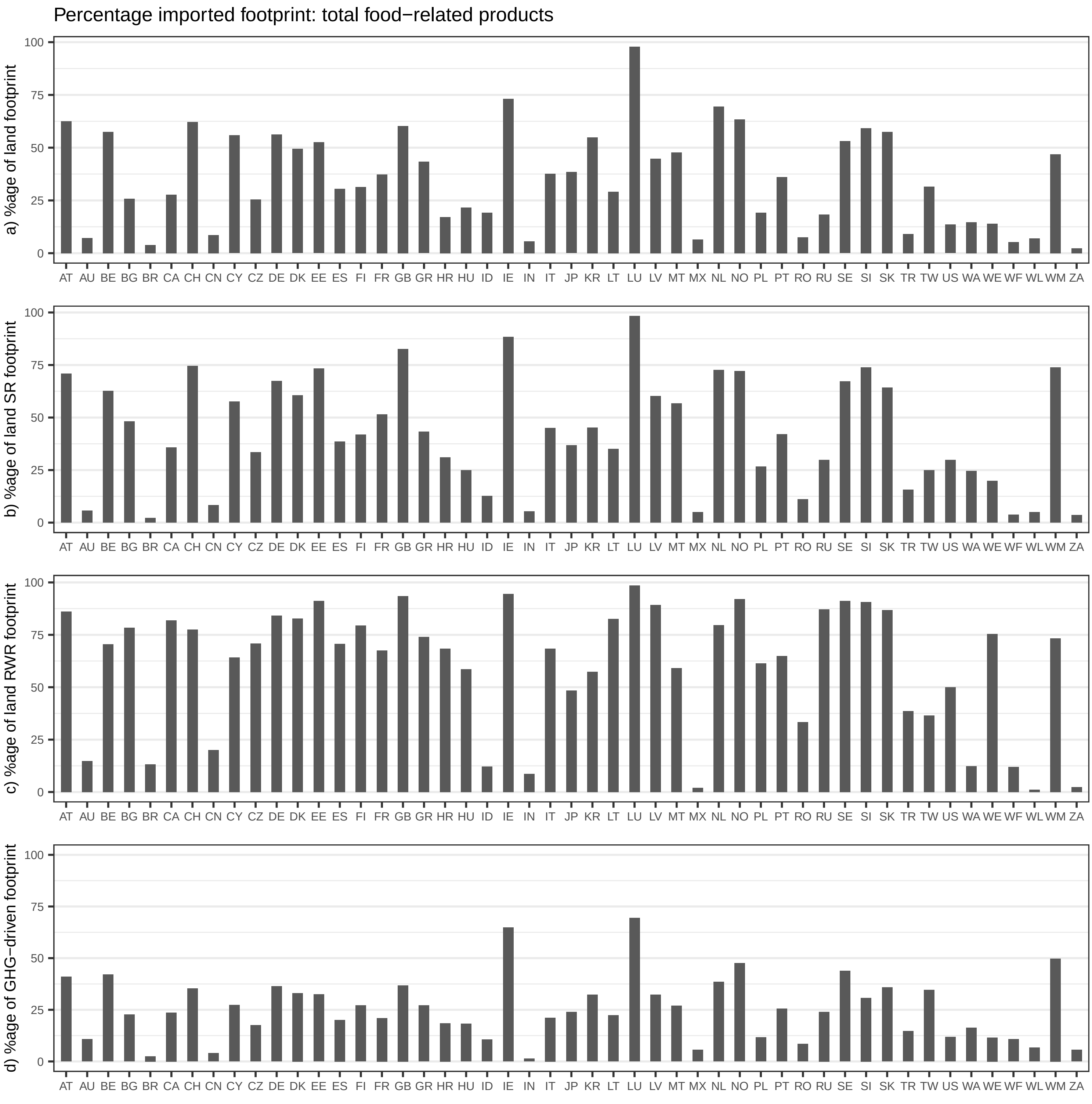

### Supplementary Figure 11

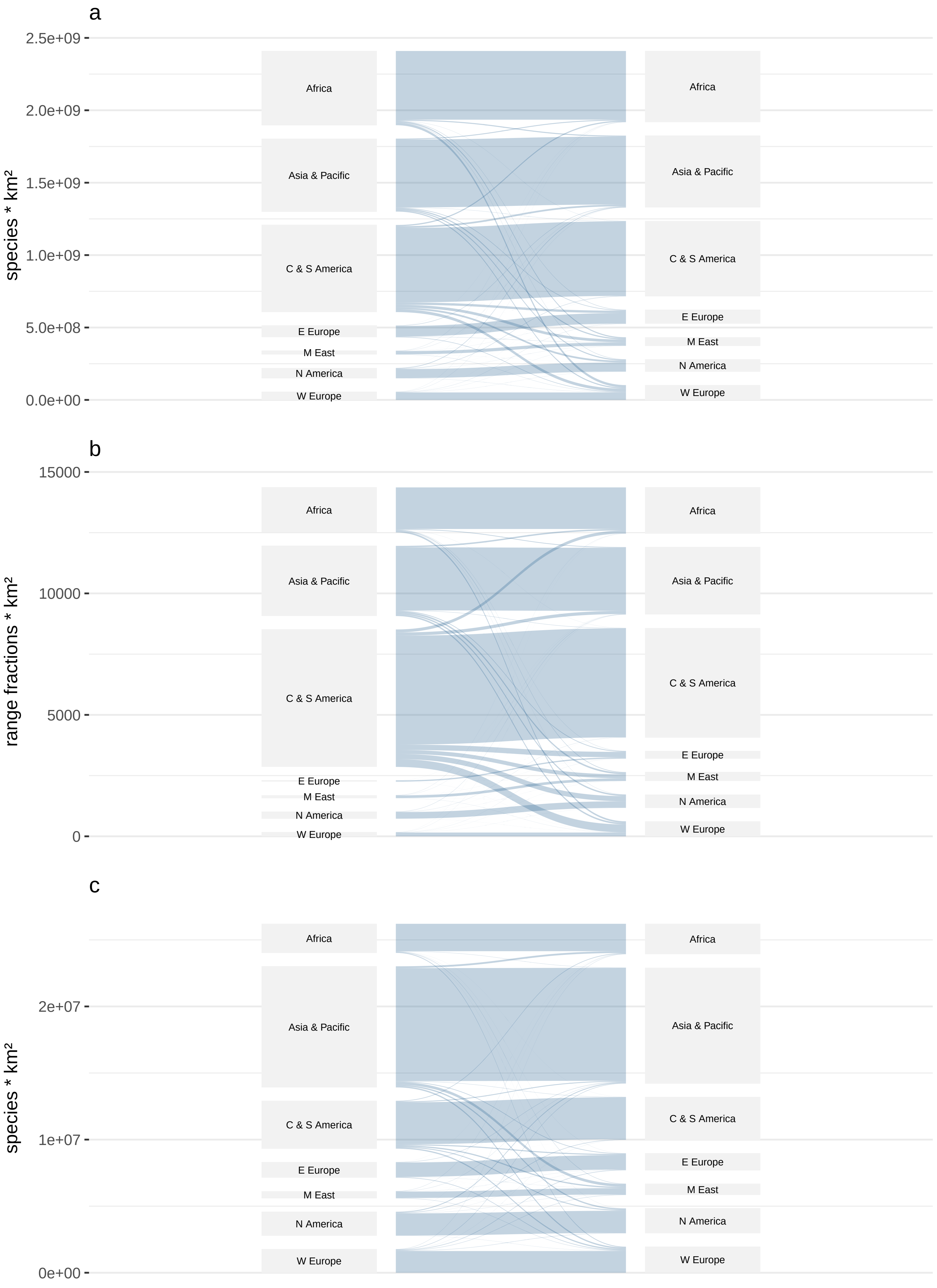

### Supplementary Figure 12

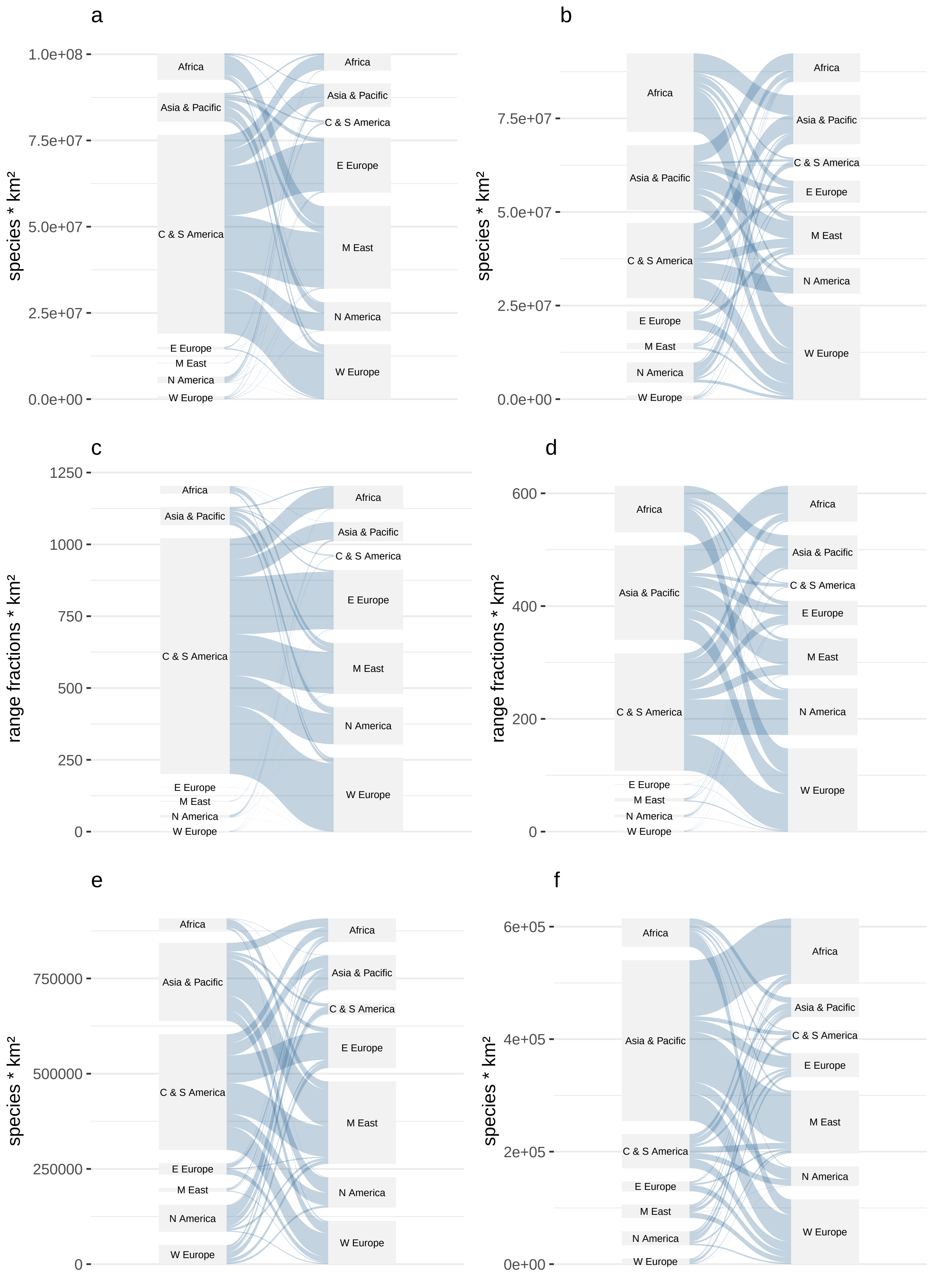
