## Supplementary Figure 5 for "Impacts of global food supply on biodiversity via land use and climate change"

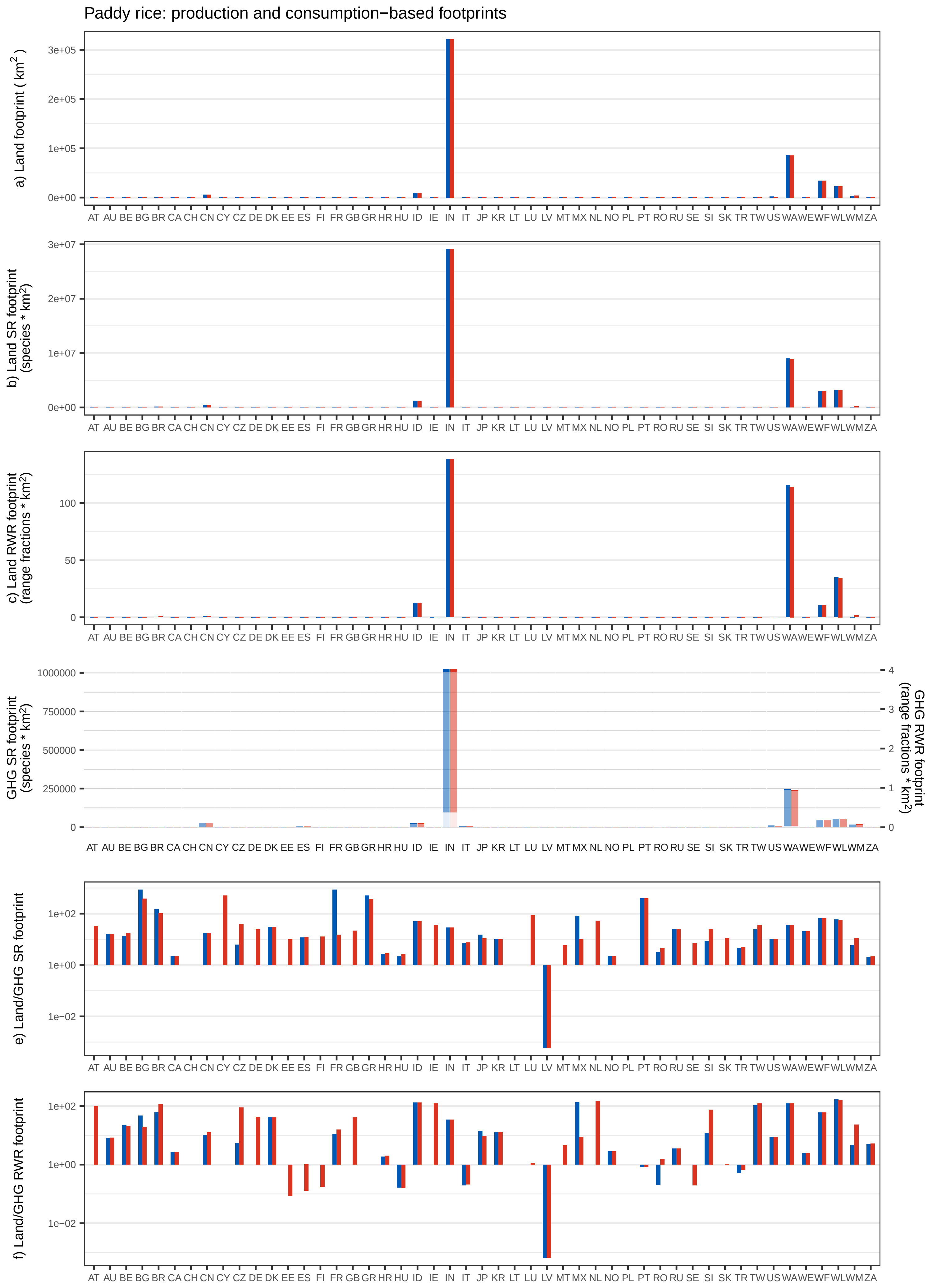

Paddy rice: production and consumption−based footprints
a) Land footprint ( km2 )
b) Land SR footprint (species * km2)
c) Land RWR footprint (range fractions * km2)
GHG SR footprint (species * km2)
GHG RWR footprint (range fractions * km2)
e) Land/GHG SR footprint
f) Land/GHG RWR footprint
AT AU BE BG BR CA CH CN CY CZ DE DK EE ES FI FR GB GR HR HU ID IE IN IT JP KR LT LU LV MT MX NL NO PL PT RO RU SE SI SK TR TW US WA WE WF WL WM ZA

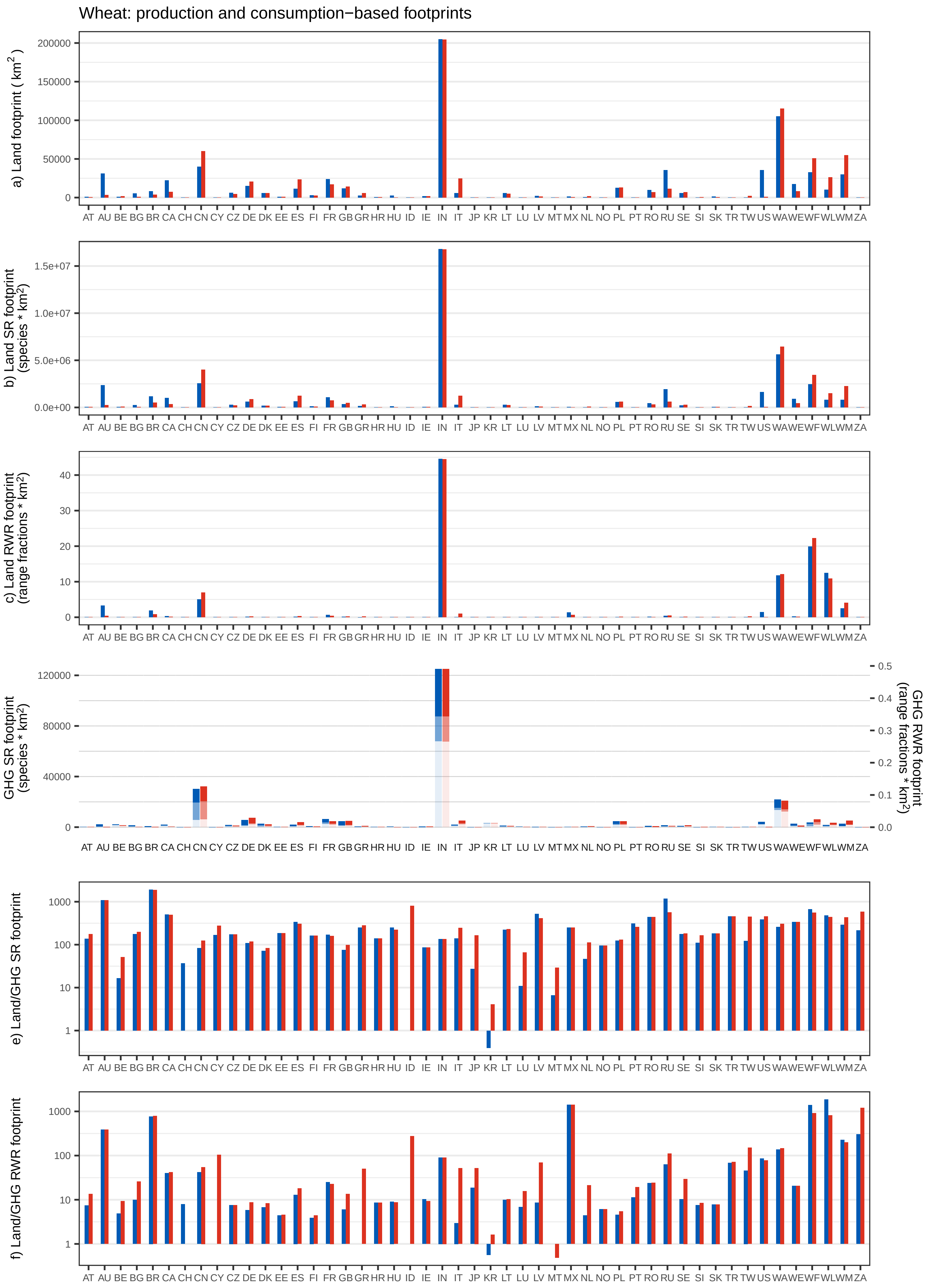

Wheat: production and consumption−based footprints
a) Land footprint ( km2 )
200000
150000
100000
50000
0
b) Land SR footprint (species * km2)
1.5e+07
1.0e+07
5.0e+06
0.0e+00
c) Land RWR footprint (range fractions * km2)
40
30
20
10
0
GHG SR footprint (species * km2)
120000
80000
40000
0
GHG RWR footprint (range fractions * km2)
0.5
0.4
0.3
0.2
0.1
0.0
e) Land/GHG SR footprint
1000
100
10
1
f) Land/GHG RWR footprint
1000
100
10
1
AT AU BE BG BR CA CH CN CY CZ DE DK EE ES FI FR GB GR HR HU ID IE IN IT JP KR LT LU LV MT MX NL NO PL PT RO RU SE SI SK TR TW US WA WE WF WL WM ZA

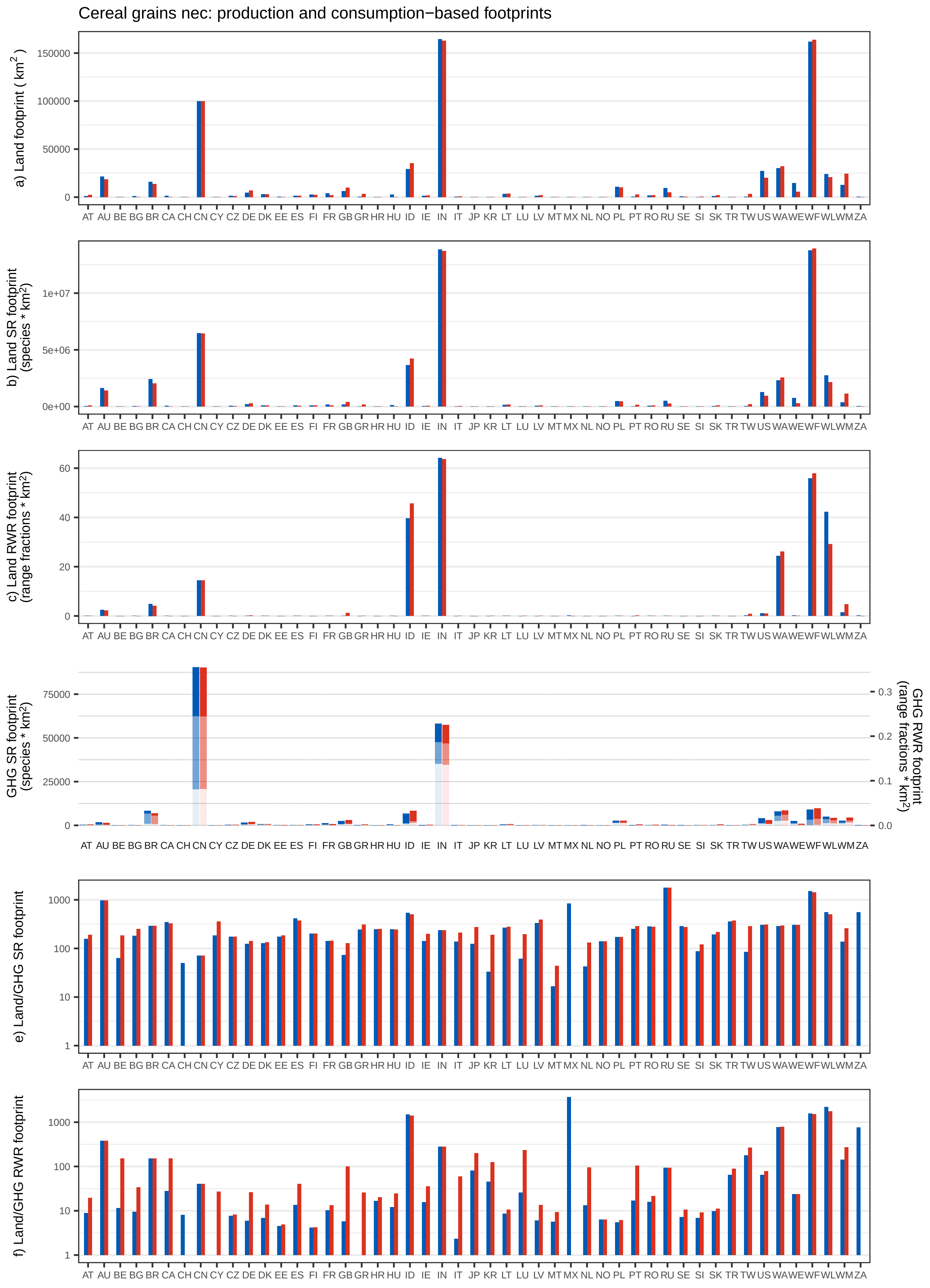

Cereal grains nec: production and consumption−based footprints
a) Land footprint ( km$^2$ )
b) Land SR footprint (species * km$^2$)
c) Land RWR footprint (range fractions * km$^2$)
GHG SR footprint (species * km$^2$)
GHG RWR footprint (range fractions * km$^2$)
e) Land/GHG SR footprint
f) Land/GHG RWR footprint
AT AU BE BG BR CA CH CN CY CZ DE DK EE ES FI FR GB GR HR HU ID IE IN IT JP KR LT LU LV MT MX NL NO PL PT RO RU SE SI SK TR TW US WA WE WF WL WM ZA

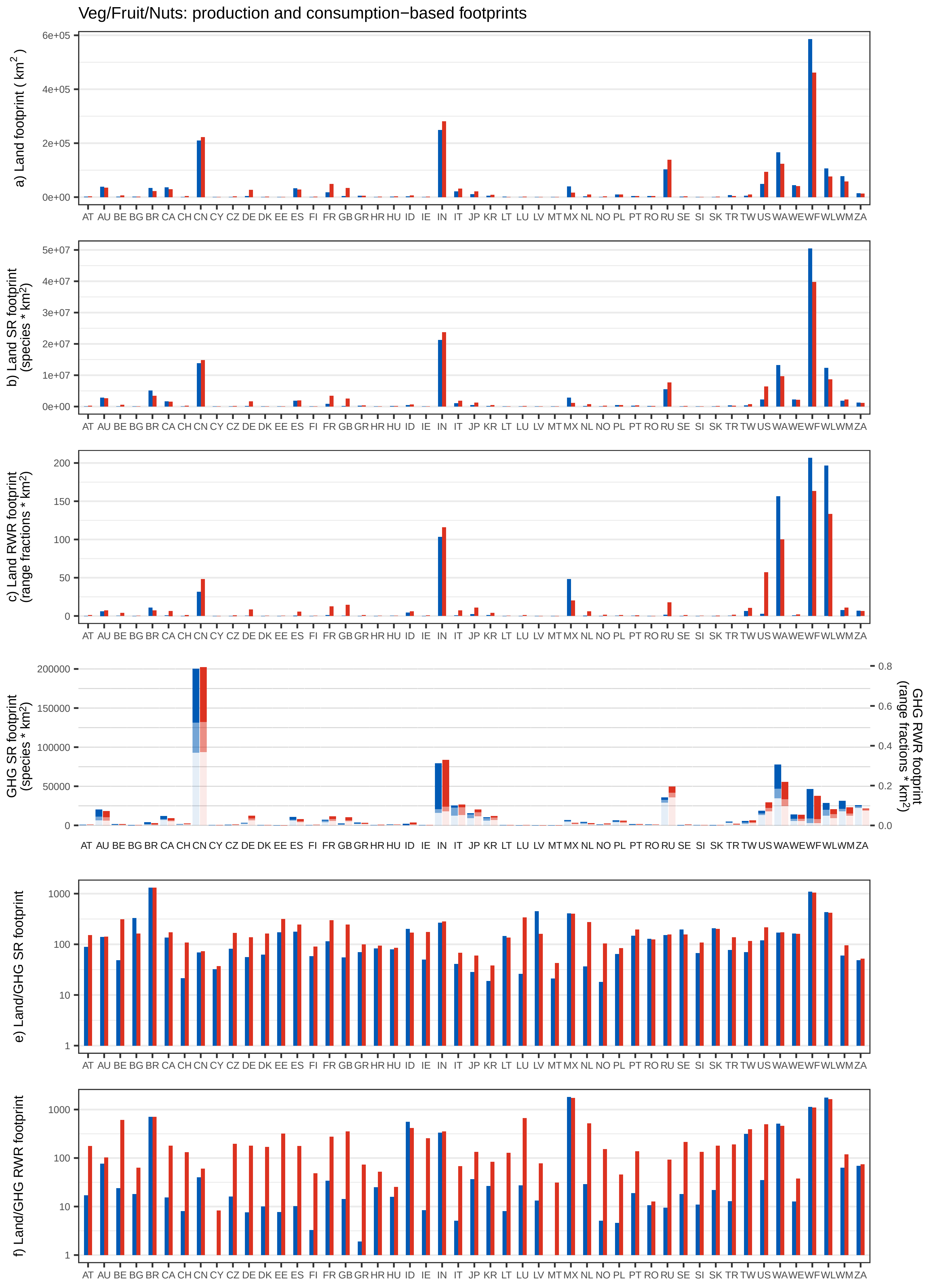

Veg/Fruit/Nuts: production and consumption−based footprints
a) Land footprint ( km2 )
b) Land SR footprint (species * km2)
c) Land RWR footprint (range fractions * km2)
GHG SR footprint (species * km2)
GHG RWR footprint (range fractions * km2)
e) Land/GHG SR footprint
f) Land/GHG RWR footprint
AT AU BE BG BR CA CH CN CY CZ DE DK EE ES FI FR GB GR HR HU ID IE IN IT JP KR LT LU LV MT MX NL NO PL PT RO RU SE SI SK TR TW US WA WE WF WL WM ZA

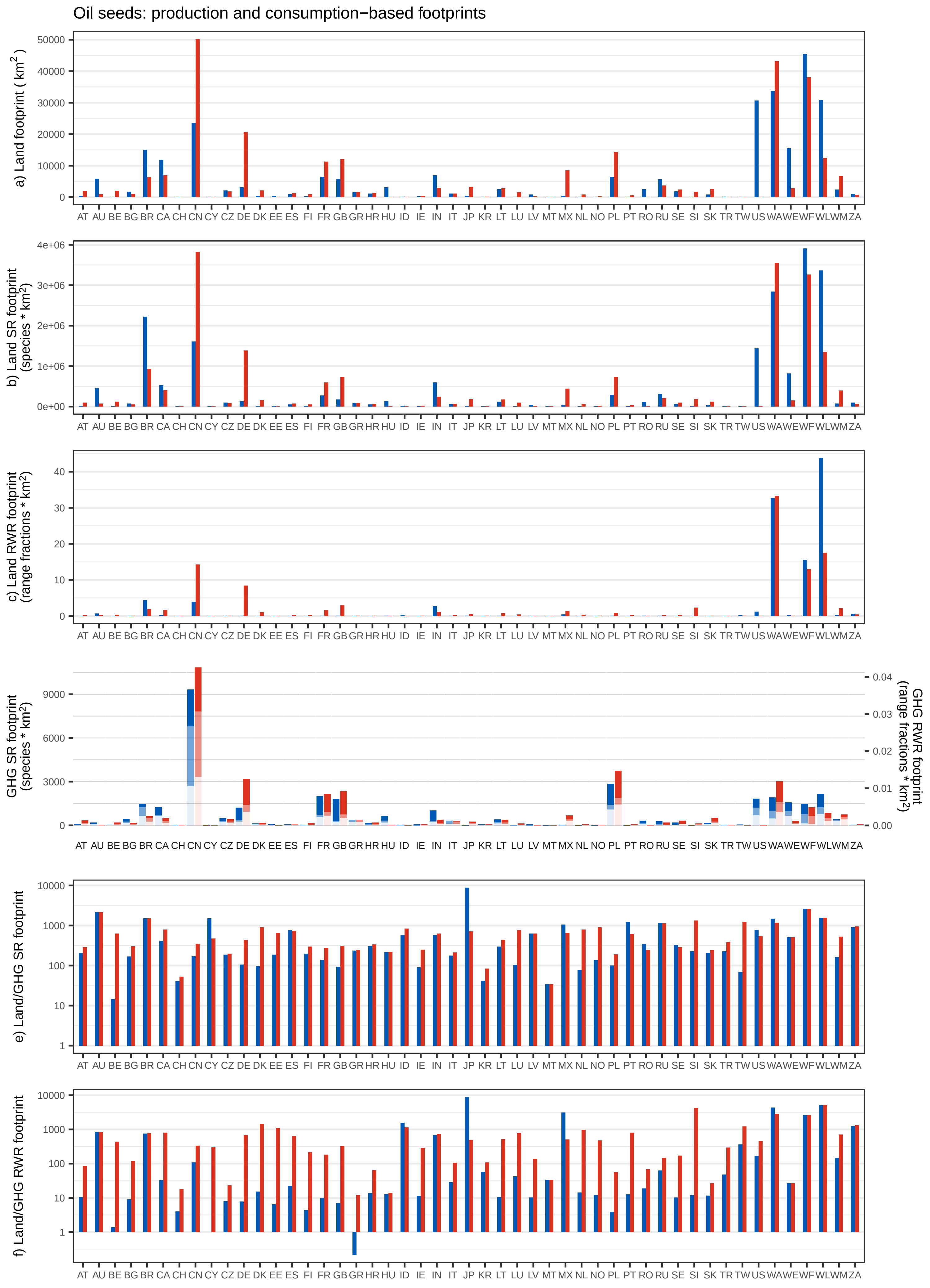

Oil seeds: production and consumption−based footprints
a) Land footprint ( km2 )
b) Land SR footprint (species * km2)
c) Land RWR footprint (range fractions * km2)
GHG SR footprint (species * km2)
GHG RWR footprint (range fractions * km2)
e) Land/GHG SR footprint
f) Land/GHG RWR footprint
AT AU BE BG BR CA CH CN CY CZ DE DK EE ES FI FR GB GR HR HU ID IE IN IT JP KR LT LU LV MT MX NL NO PL PT RO RU SE SI SK TR TW US WA WE WF WL WM ZA

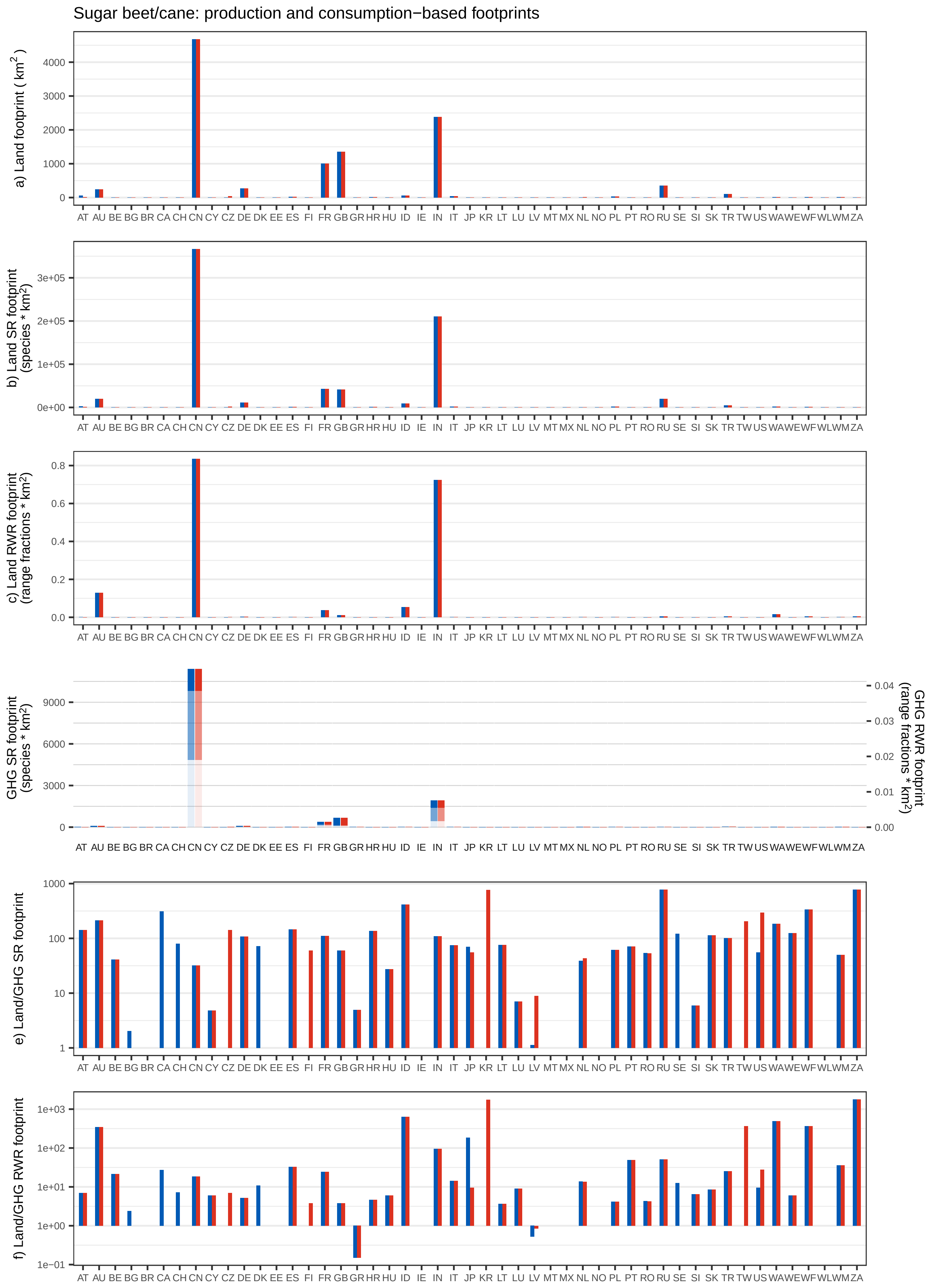

Sugar beet/cane: production and consumption−based footprints
a) Land footprint ( km$^2$ )
b) Land SR footprint (species * km$^2$)
c) Land RWR footprint (range fractions * km$^2$)
GHG SR footprint (species * km$^2$)
GHG RWR footprint (range fractions * km$^2$)
e) Land/GHG SR footprint
f) Land/GHG RWR footprint
AT AU BE BG BR CA CH CN CY CZ DE DK EE ES FI FR GB GR HR HU ID IE IN IT JP KR LT LU LV MT MX NL NO PL PT RO RU SE SI SK TR TW US WA WE WF WL WM ZA

Crops nec: production and consumption−based footprints

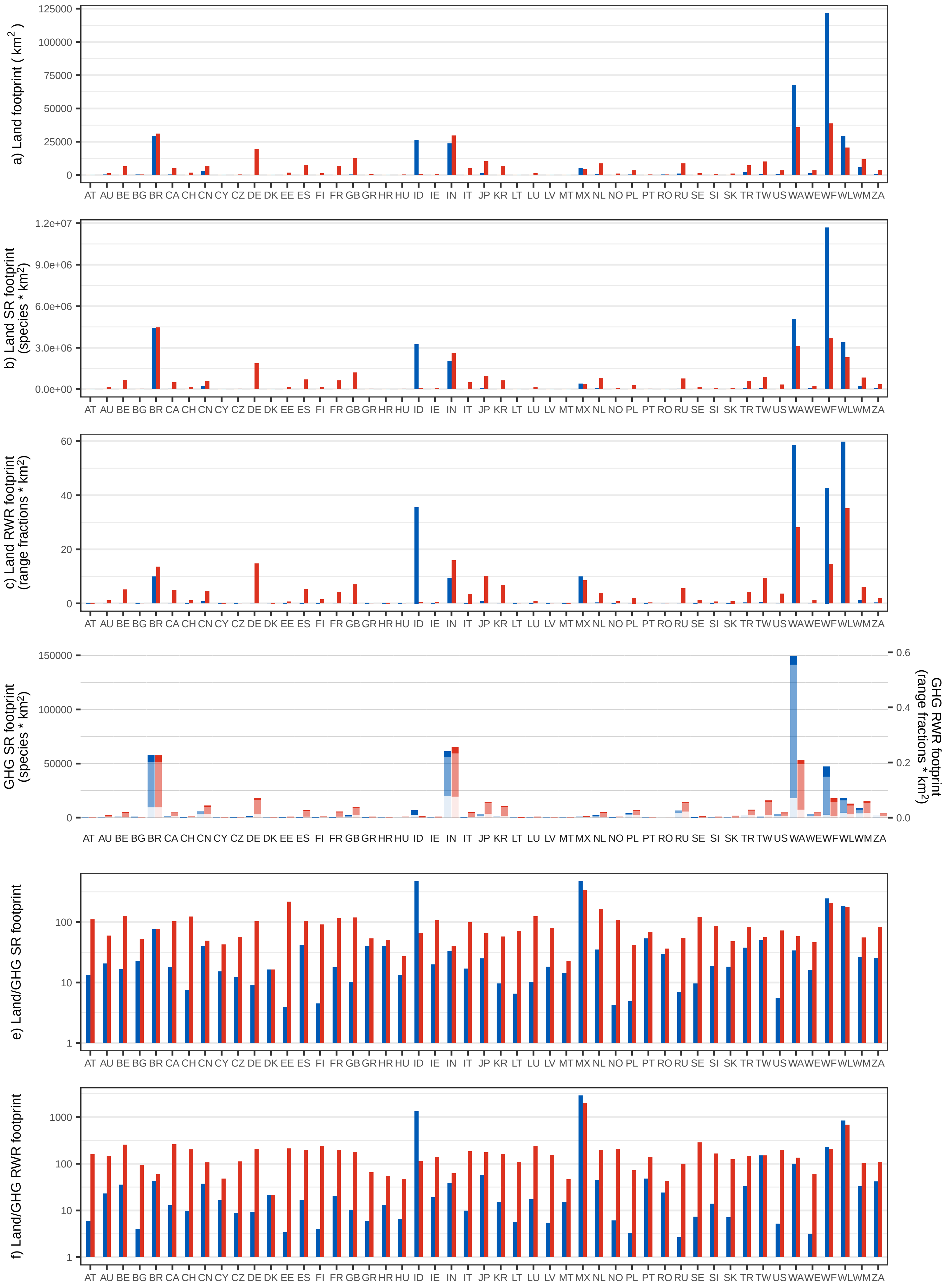

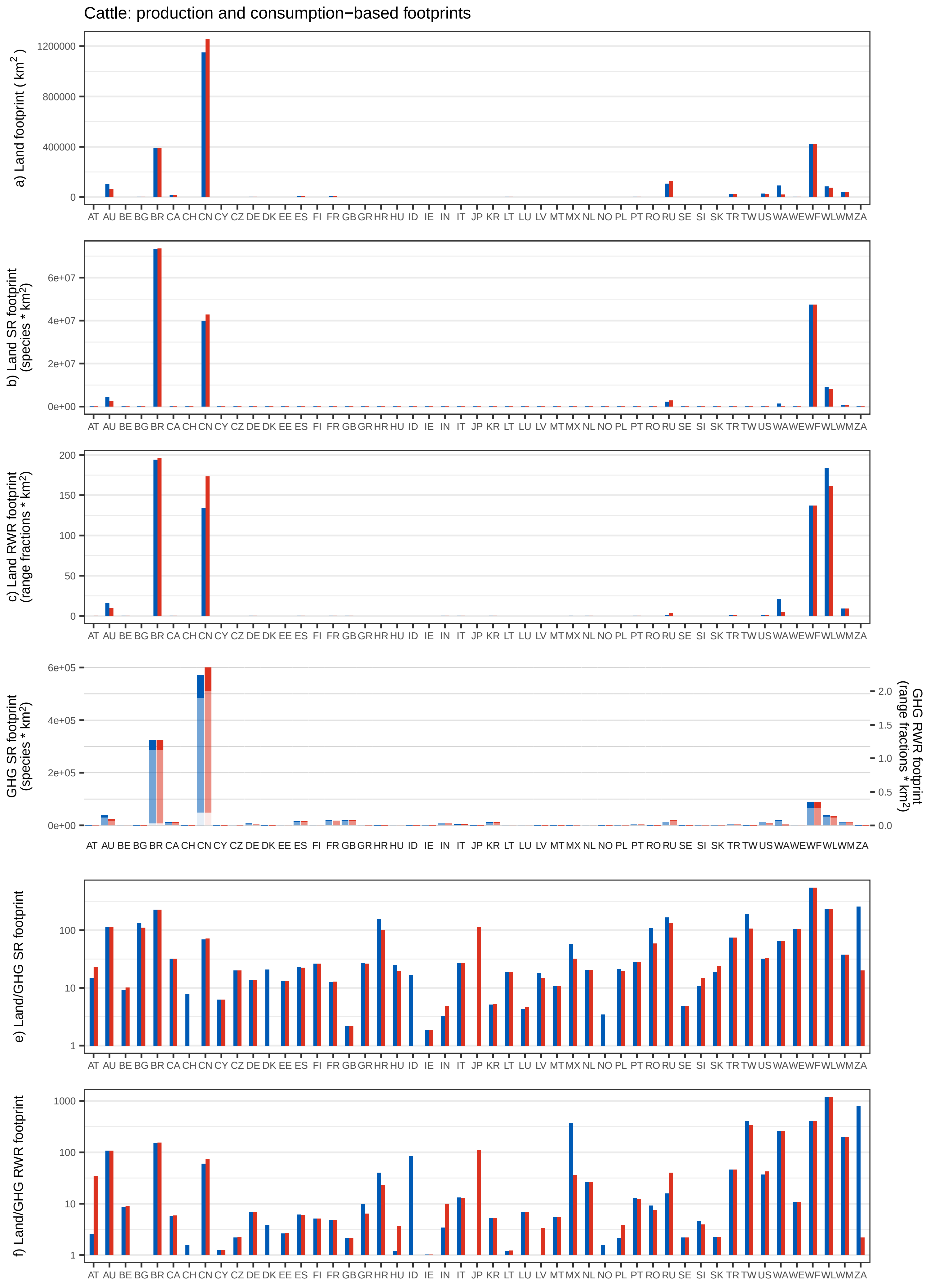

Cattle: production and consumption−based footprints
a) Land footprint ( km$^2$ )
b) Land SR footprint (species * km$^2$)
c) Land RWR footprint (range fractions * km$^2$)
GHG SR footprint (species * km$^2$)
GHG RWR footprint (range fractions * km$^2$)
e) Land/GHG SR footprint
f) Land/GHG RWR footprint
AT AU BE BG BR CA CH CN CY CZ DE DK EE ES FI FR GB GR HR HU ID IE IN IT JP KR LT LU LV MT MX NL NO PL PT RO RU SE SI SK TR TW US WA WE WF WL WM ZA

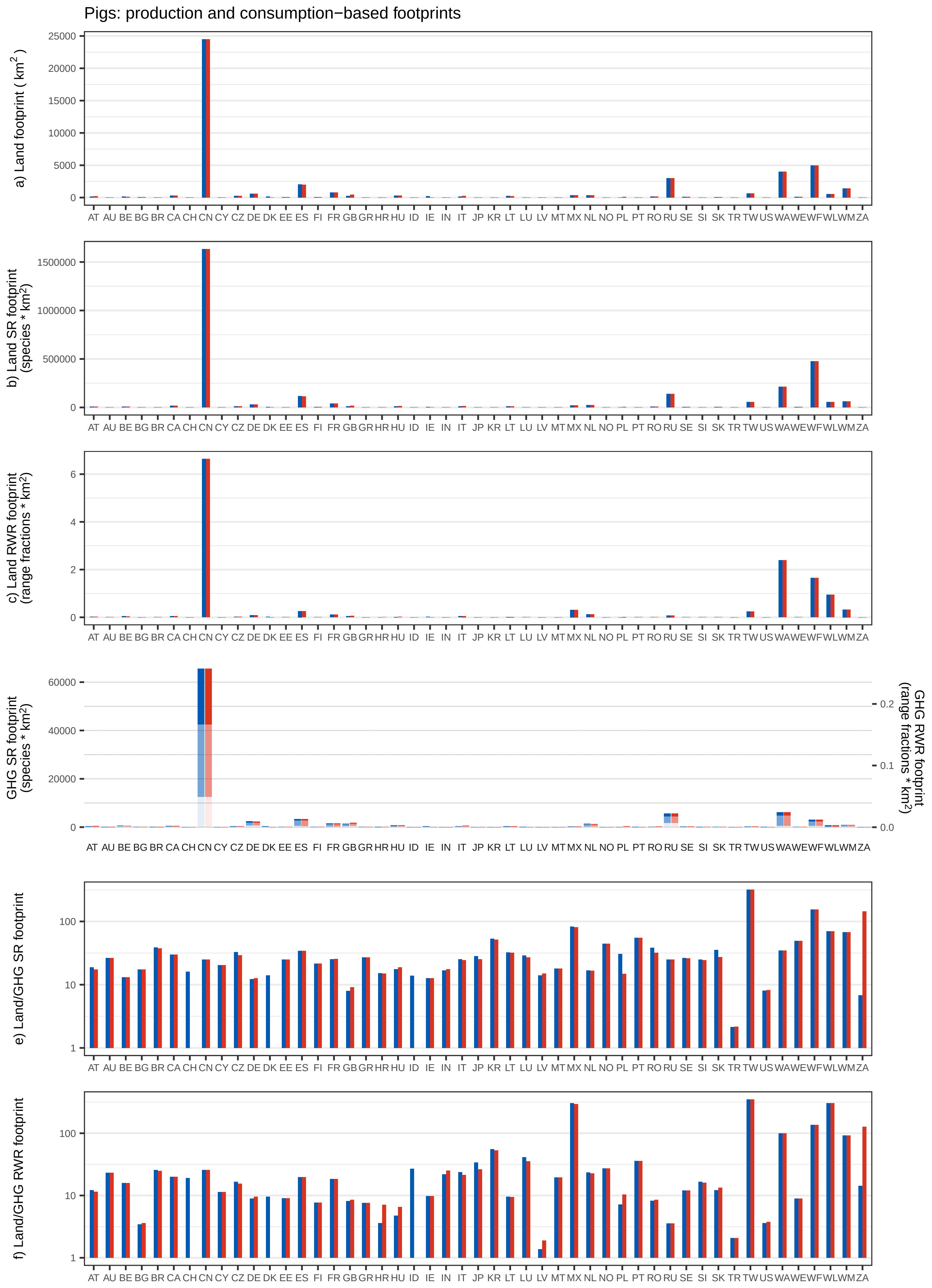

Pigs: production and consumption−based footprints
a) Land footprint ( km$^2$ )
b) Land SR footprint (species * km$^2$)
c) Land RWR footprint (range fractions * km$^2$)
GHG SR footprint (species * km$^2$)
GHG RWR footprint (range fractions * km$^2$)
e) Land/GHG SR footprint
f) Land/GHG RWR footprint
AT AU BE BG BR CA CH CN CY CZ DE DK EE ES FI FR GB GR HR HU ID IE IN IT JP KR LT LU LV MT MX NL NO PL PT RO RU SE SI SK TR TW US WA WE WF WL WM ZA

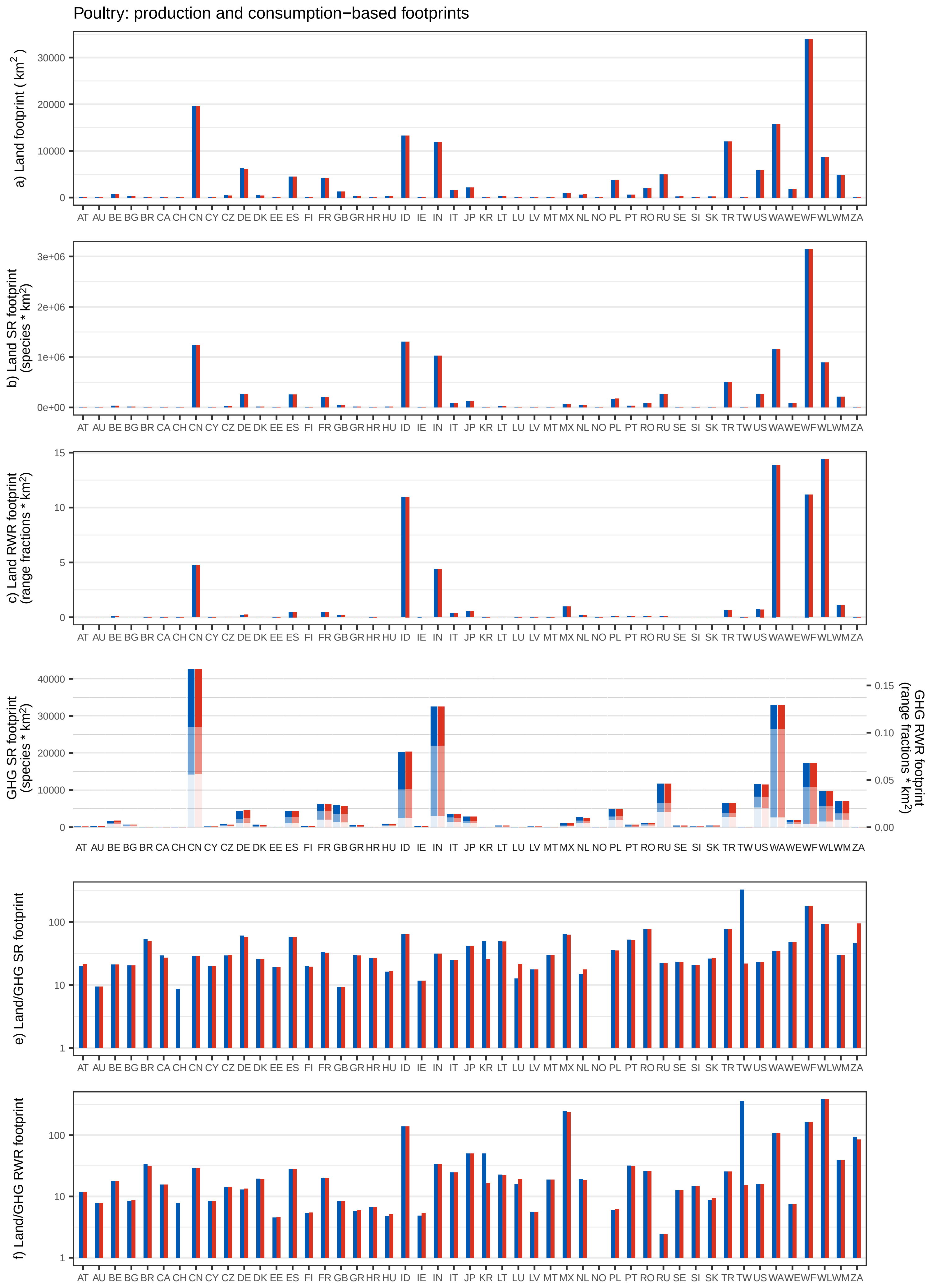

Poultry: production and consumption−based footprints
a) Land footprint ( km$^2$ )
b) Land SR footprint (species * km$^2$)
c) Land RWR footprint (range fractions * km$^2$)
GHG SR footprint (species * km$^2$)
GHG RWR footprint (range fractions * km$^2$)
e) Land/GHG SR footprint
f) Land/GHG RWR footprint
AT AU BE BG BR CA CH CN CY CZ DE DK EE ES FI FR GB GR HR HU ID IE IN IT JP KR LT LU LV MT MX NL NO PL PT RO RU SE SI SK TR TW US WA WE WF WL WM ZA

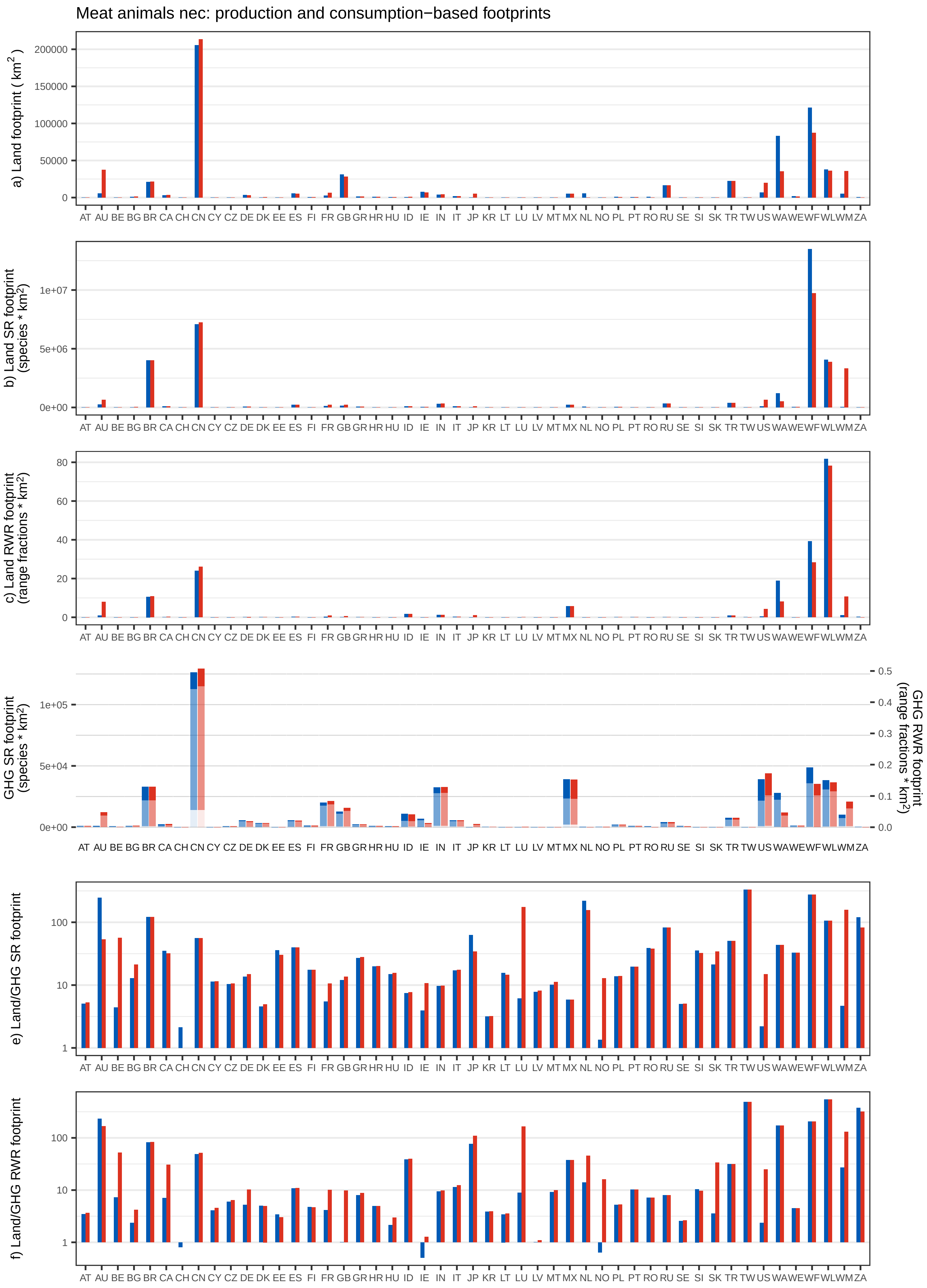

Meat animals nec: production and consumption−based footprints
a) Land footprint ( km$^2$ )
b) Land SR footprint (species * km$^2$)
c) Land RWR footprint (range fractions * km$^2$)
GHG SR footprint (species * km$^2$)
GHG RWR footprint (range fractions * km$^2$)
e) Land/GHG SR footprint
f) Land/GHG RWR footprint
AT AU BE BG BR CA CH CN CY CZ DE DK EE ES FI FR GB GR HR HU ID IE IN IT JP KR LT LU LV MT MX NL NO PL PT RO RU SE SI SK TR TW US WA WE WF WL WM ZA

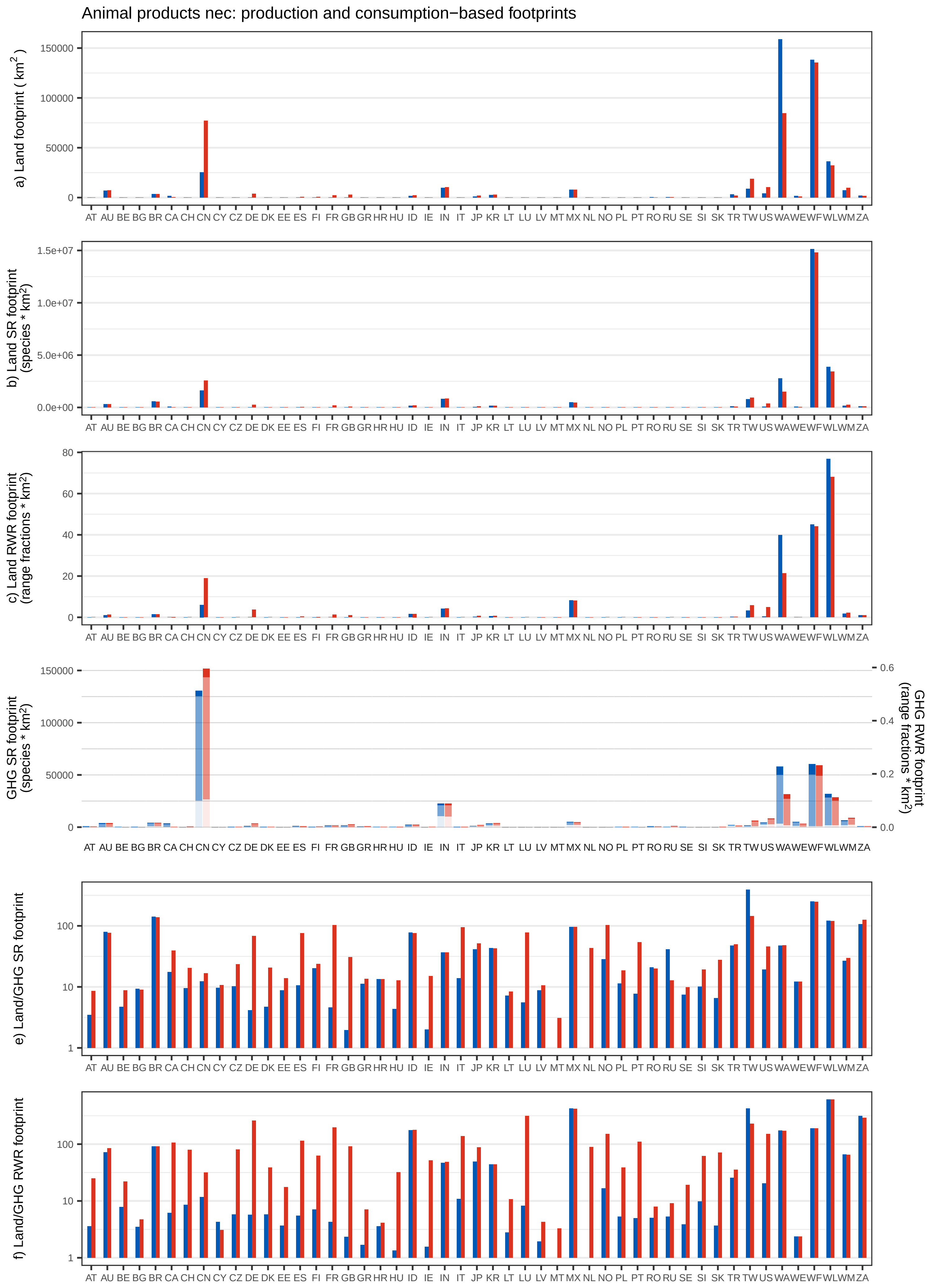

Animal products nec: production and consumption−based footprints
a) Land footprint ( km2 )
b) Land SR footprint (species * km2)
c) Land RWR footprint (range fractions * km2)
GHG SR footprint (species * km2)
GHG RWR footprint (range fractions * km2)
e) Land/GHG SR footprint
f) Land/GHG RWR footprint
AT AU BE BG BR CA CH CN CY CZ DE DK EE ES FI FR GB GR HR HU ID IE IN IT JP KR LT LU LV MT MX NL NO PL PT RO RU SE SI SK TR TW US WA WE WF WL WM ZA

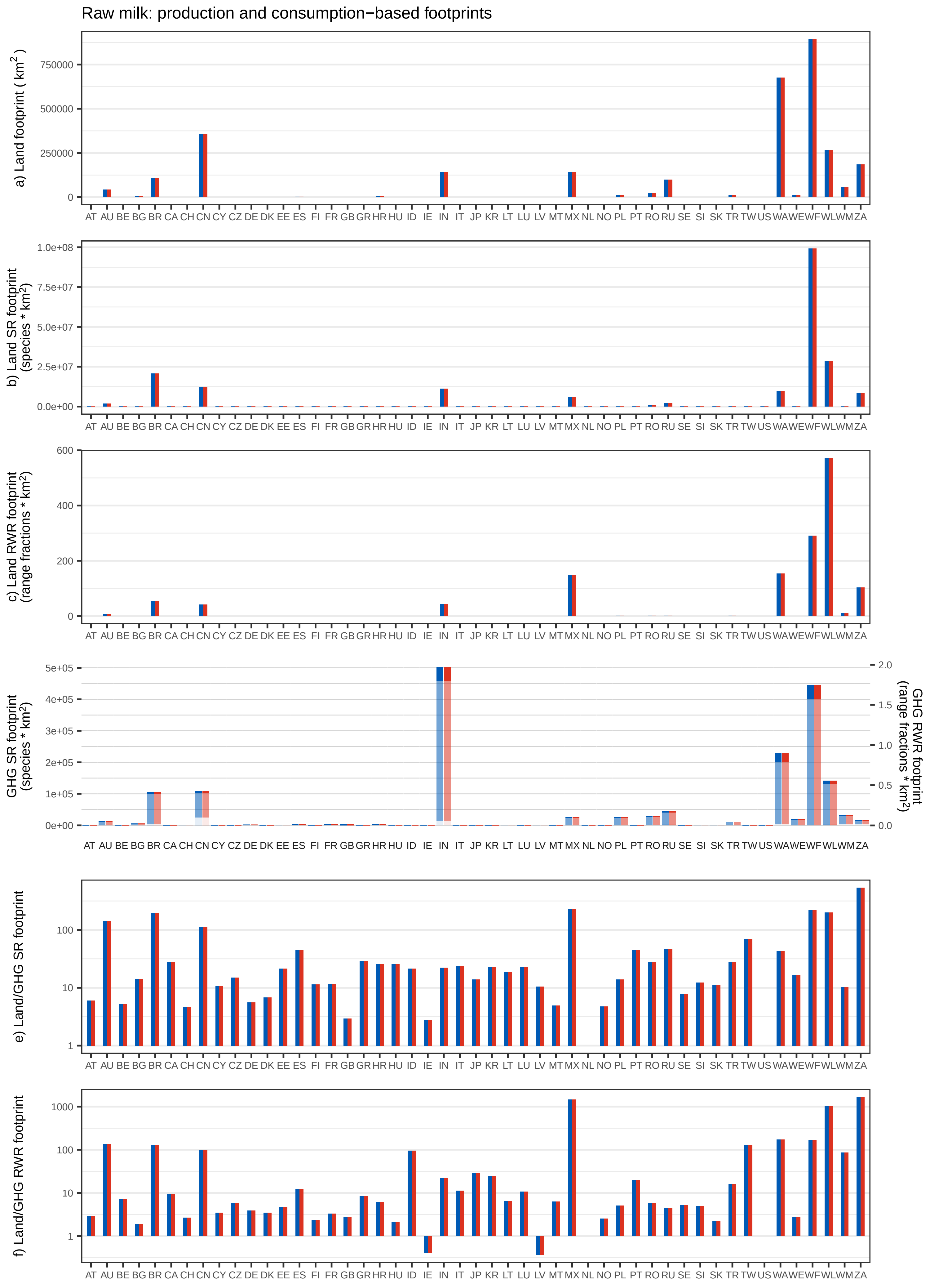
Raw milk: production and consumption−based footprints
a) Land footprint ( km2 )
b) Land SR footprint (species * km2)
c) Land RWR footprint (range fractions * km2)
GHG SR footprint (species * km2)
GHG RWR footprint (range fractions * km2)
e) Land/GHG SR footprint
f) Land/GHG RWR footprint
AT AU BE BG BR CA CH CN CY CZ DE DK EE ES FI FR GB GR HR HU ID IE IN IT JP KR LT LU LV MT MX NL NO PL PT RO RU SE SI SK TR TW US WA WE WF WL WM ZA

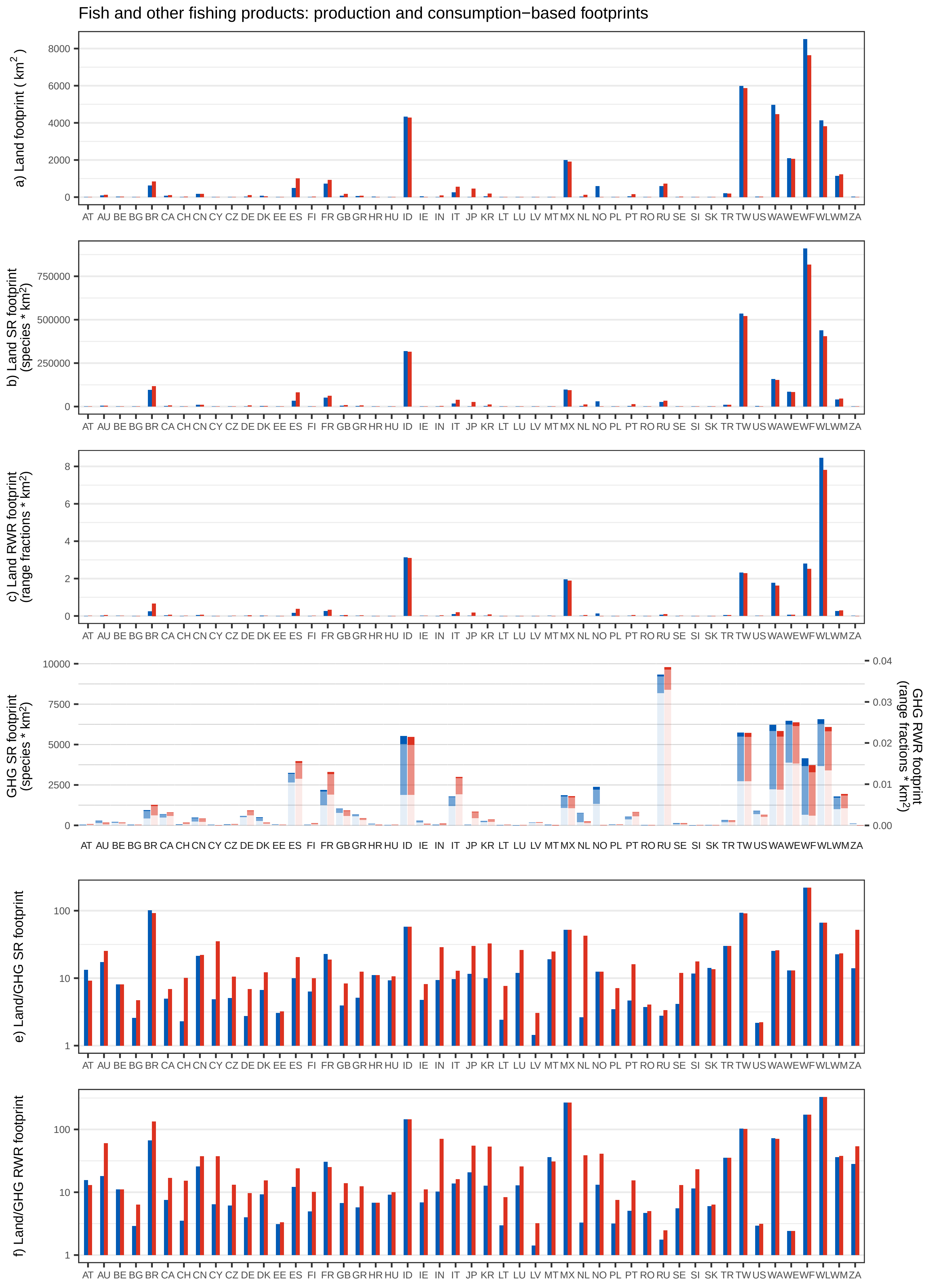
Fish and other fishing products: production and consumption−based footprints
a) Land footprint ( km$^2$ )
b) Land SR footprint (species * km$^2$)
c) Land RWR footprint (range fractions * km$^2$)
GHG SR footprint (species * km$^2$)
GHG RWR footprint (range fractions * km$^2$)
e) Land/GHG SR footprint
f) Land/GHG RWR footprint
AT AU BE BG BR CA CH CN CY CZ DE DK EE ES FI FR GB GR HR HU ID IE IN IT JP KR LT LU LV MT MX NL NO PL PT RO RU SE SI SK TR TW US WA WE WF WL WM ZA

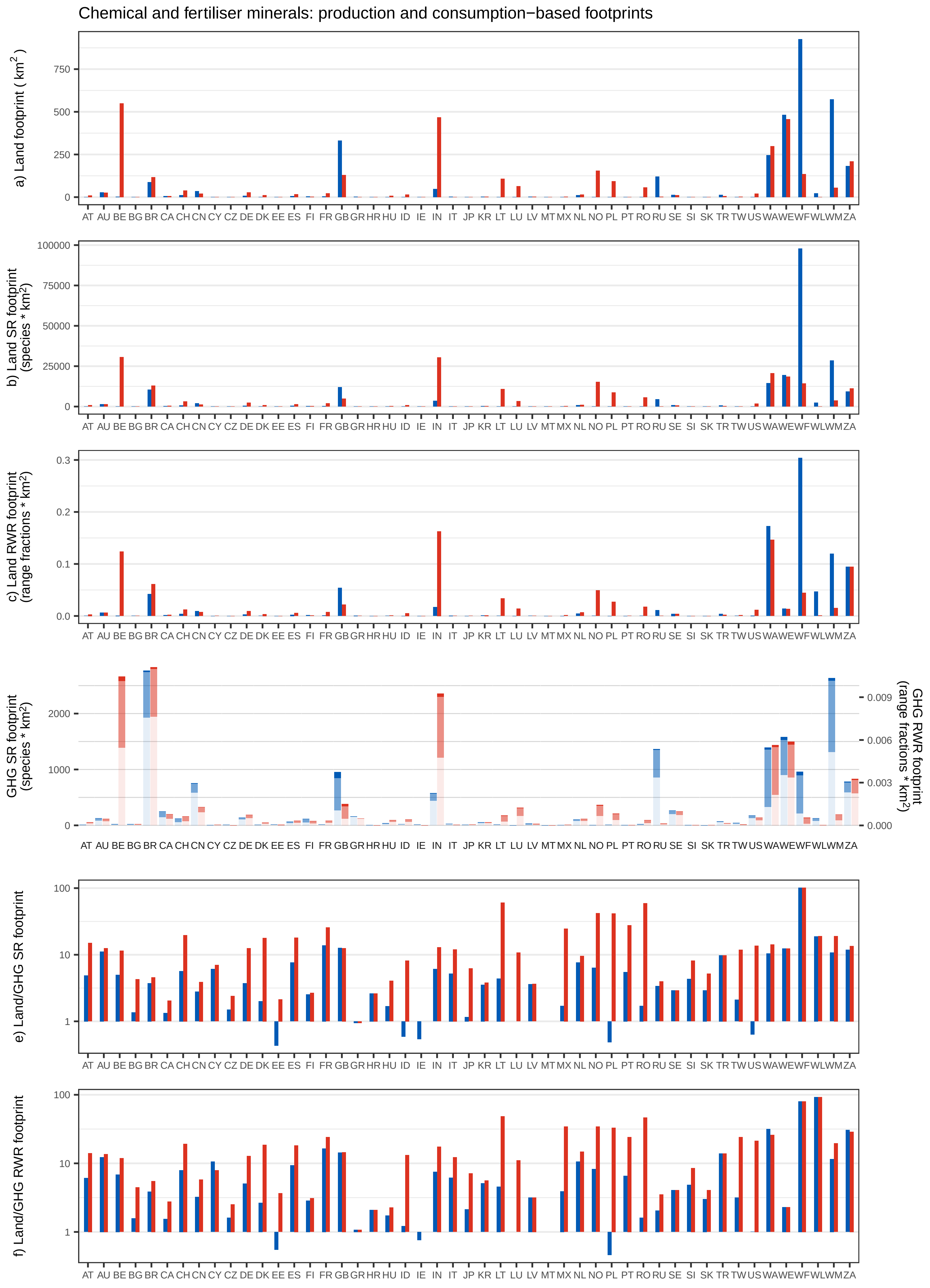
Chemical and fertiliser minerals: production and consumption−based footprints
a) Land footprint ( km² )
b) Land SR footprint (species * km²)
c) Land RWR footprint (range fractions * km²)
GHG SR footprint (species * km²)
GHG RWR footprint (range fractions * km²)
e) Land/GHG SR footprint
f) Land/GHG RWR footprint
AT AU BE BG BR CA CH CN CY CZ DE DK EE ES FI FR GB GR HR HU ID IE IN IT JP KR LT LU LV MT MX NL NO PL PT RO RU SE SI SK TR TW US WA WE WF WL WM ZA

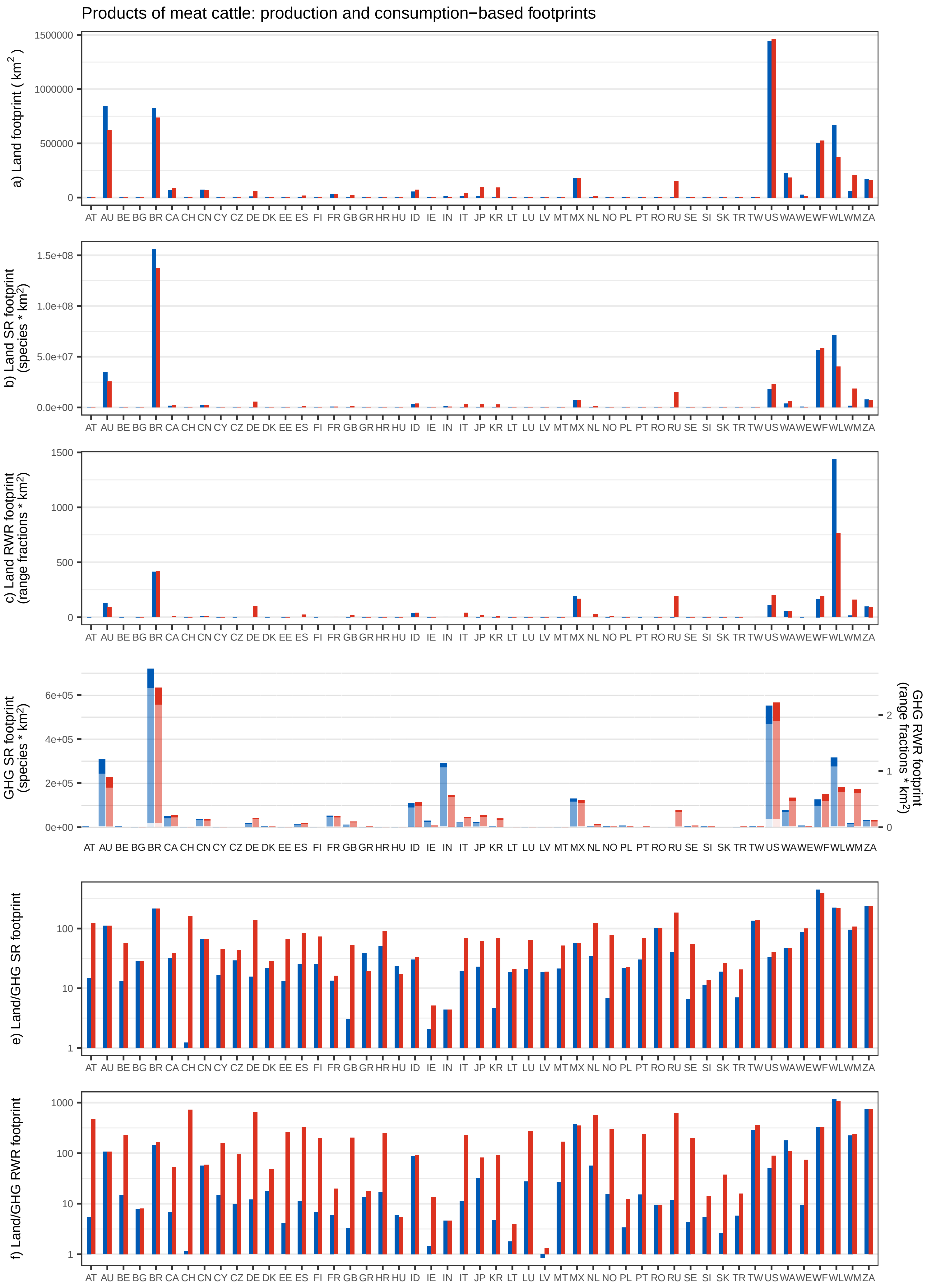
Products of meat cattle: production and consumption−based footprints
a) Land footprint ( km²)
b) Land SR footprint (species * km²)
c) Land RWR footprint (range fractions * km²)
GHG SR footprint (species * km²)
GHG RWR footprint (range fractions * km²)
e) Land/GHG SR footprint
f) Land/GHG RWR footprint
AT AU BE BG BR CA CH CN CY CZ DE DK EE ES FI FR GB GR HR HU ID IE IN IT JP KR LT LU LV MT MX NL NO PL PT RO RU SE SI SK TR TW US WA WE WF WL WM ZA

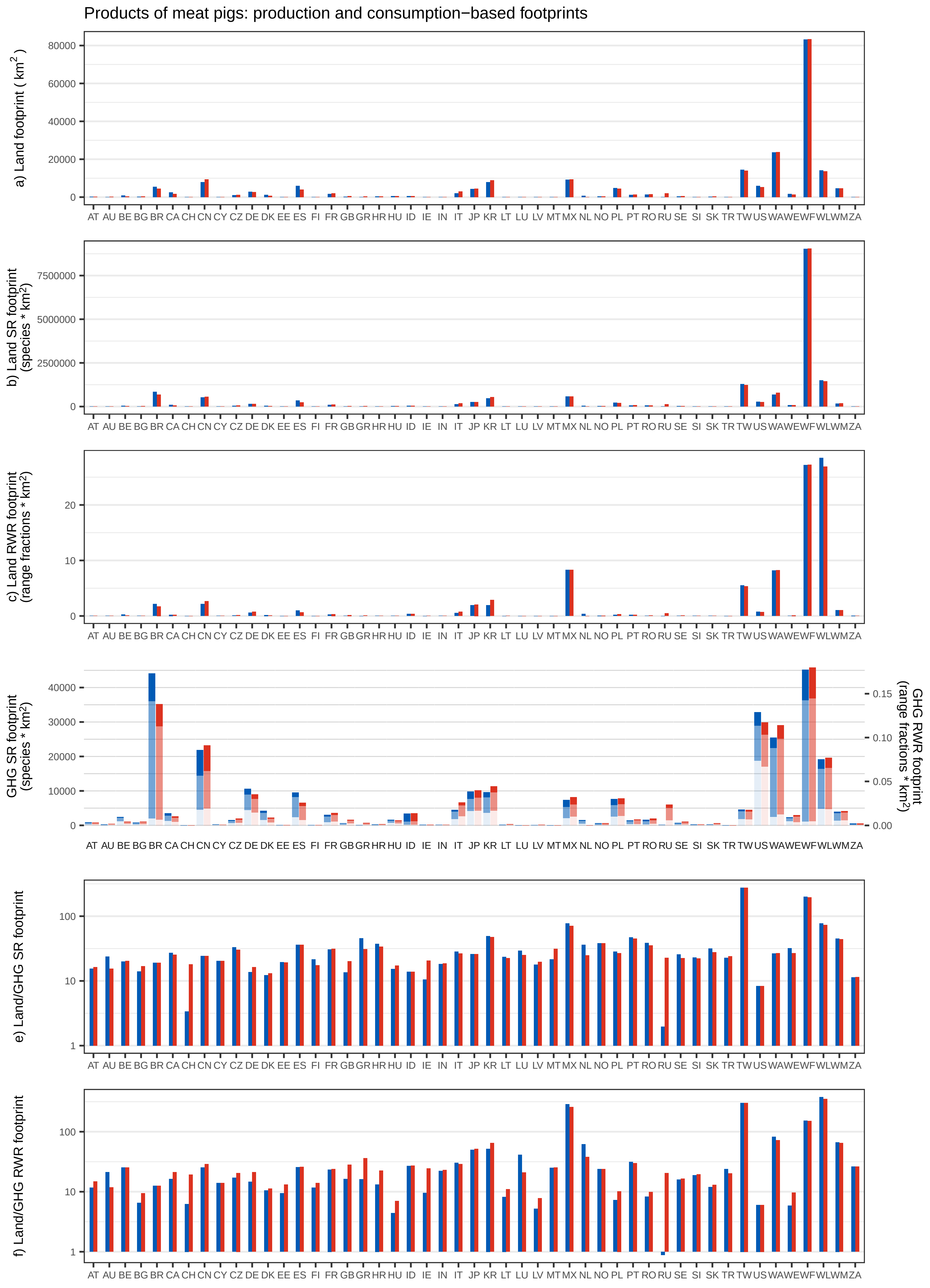

Products of meat pigs: production and consumption−based footprints
a) Land footprint ( km2 )
b) Land SR footprint (species * km2)
c) Land RWR footprint (range fractions * km2)
GHG SR footprint (species * km2)
GHG RWR footprint (range fractions * km2)
e) Land/GHG SR footprint
f) Land/GHG RWR footprint
AT AU BE BG BR CA CH CN CY CZ DE DK EE ES FI FR GB GR HR HU ID IE IN IT JP KR LT LU LV MT MX NL NO PL PT RO RU SE SI SK TR TW US WA WE WF WL WM ZA

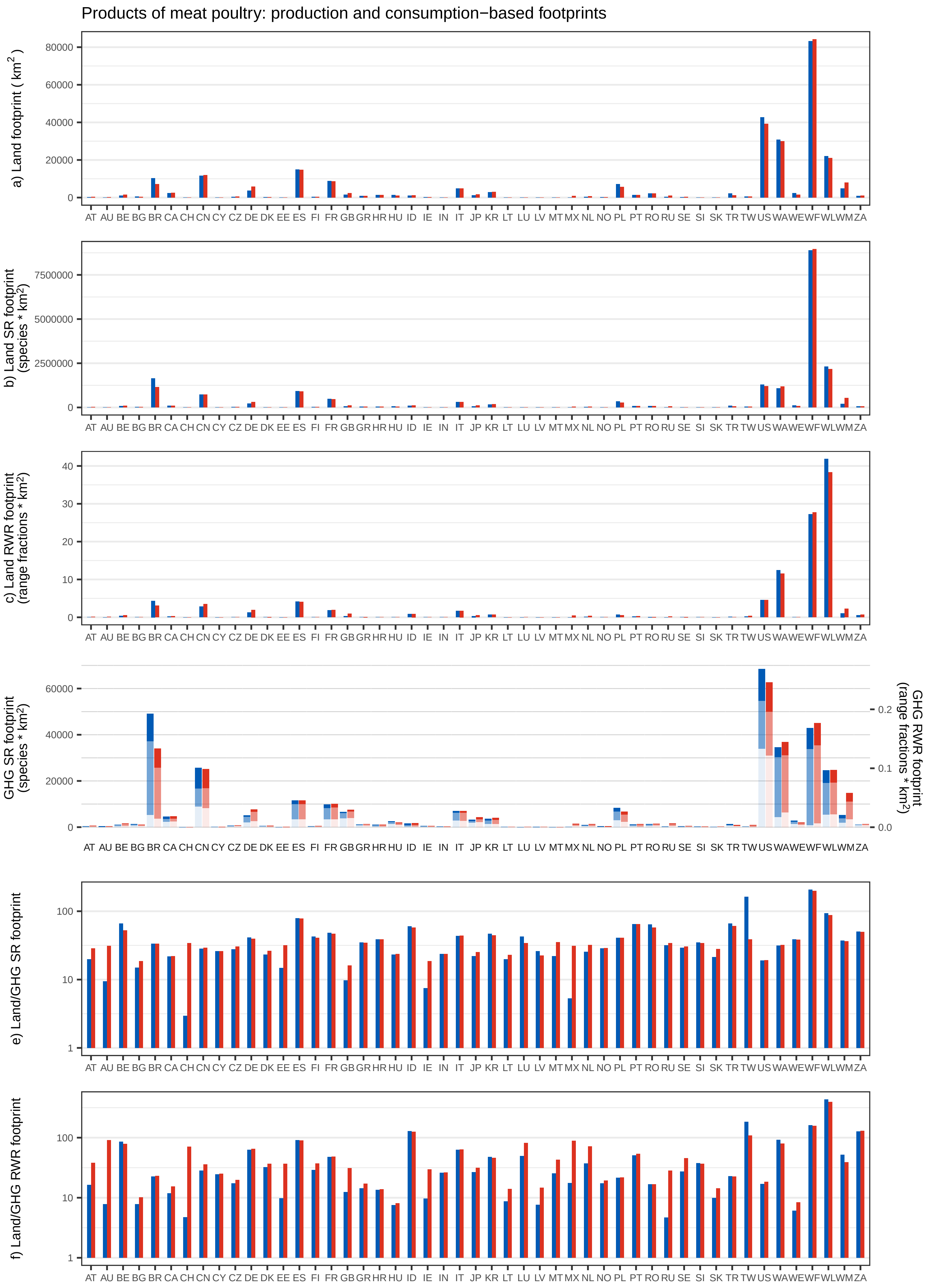

Products of meat poultry: production and consumption−based footprints
a) Land footprint ( km²)
b) Land SR footprint (species * km²)
c) Land RWR footprint (range fractions * km²)
GHG SR footprint (species * km²)
GHG RWR footprint (range fractions * km²)
e) Land/GHG SR footprint
f) Land/GHG RWR footprint
AT AU BE BG BR CA CH CN CY CZ DE DK EE ES FI FR GB GR HR HU ID IE IN IT JP KR LT LU LV MT MX NL NO PL PT RO RU SE SI SK TR TW US WA WE WF WL WM ZA

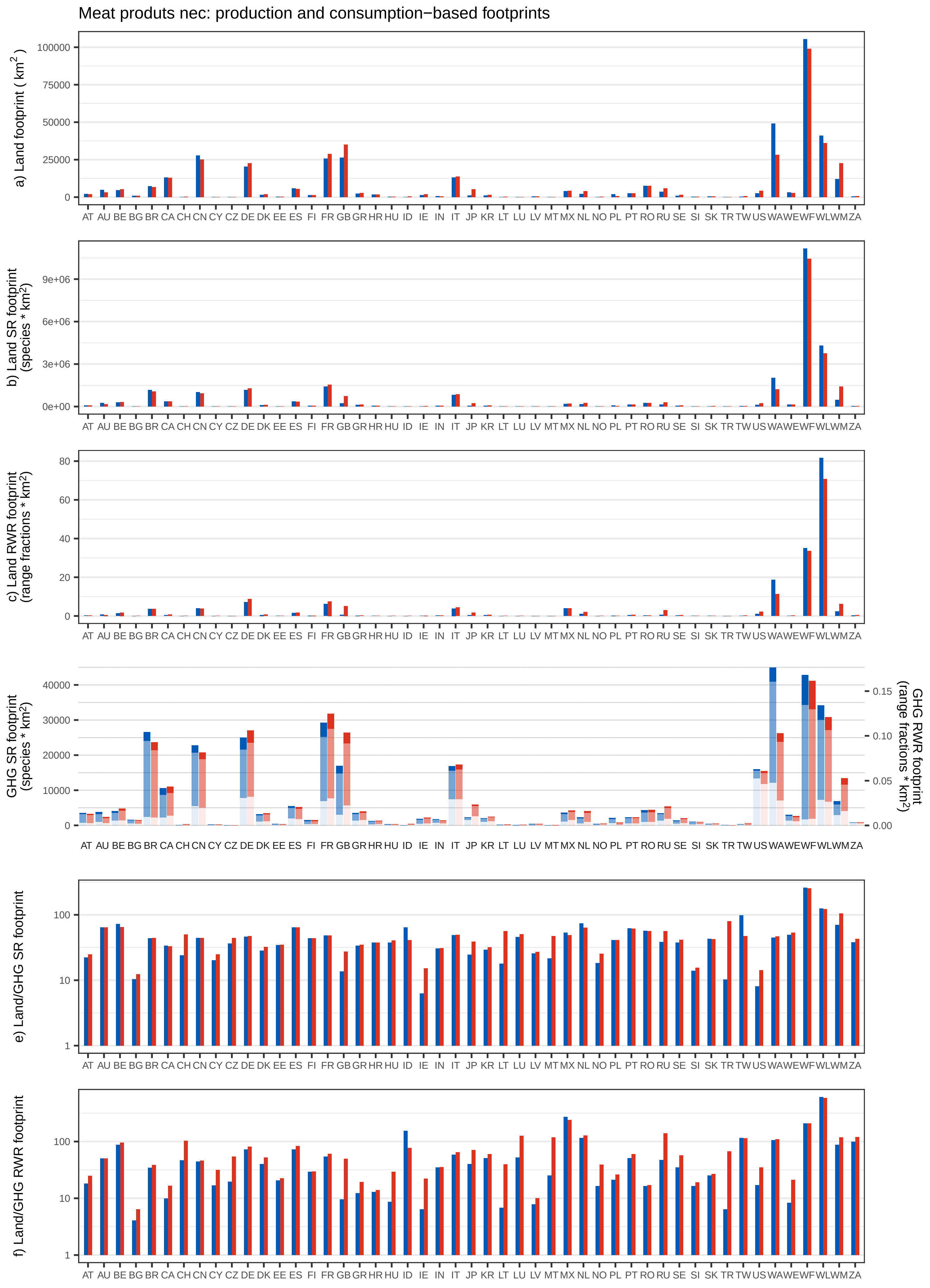

Meat produts nec: production and consumption−based footprints
a) Land footprint ( km2 )
b) Land SR footprint (species * km2)
c) Land RWR footprint (range fractions * km2)
GHG SR footprint (species * km2)
GHG RWR footprint (range fractions * km2)
e) Land/GHG SR footprint
f) Land/GHG RWR footprint
AT AU BE BG BR CA CH CN CY CZ DE DK EE ES FI FR GB GR HR HU ID IE IN IT JP KR LT LU LV MT MX NL NO PL PT RO RU SE SI SK TR TW US WA WE WF WL WM ZA

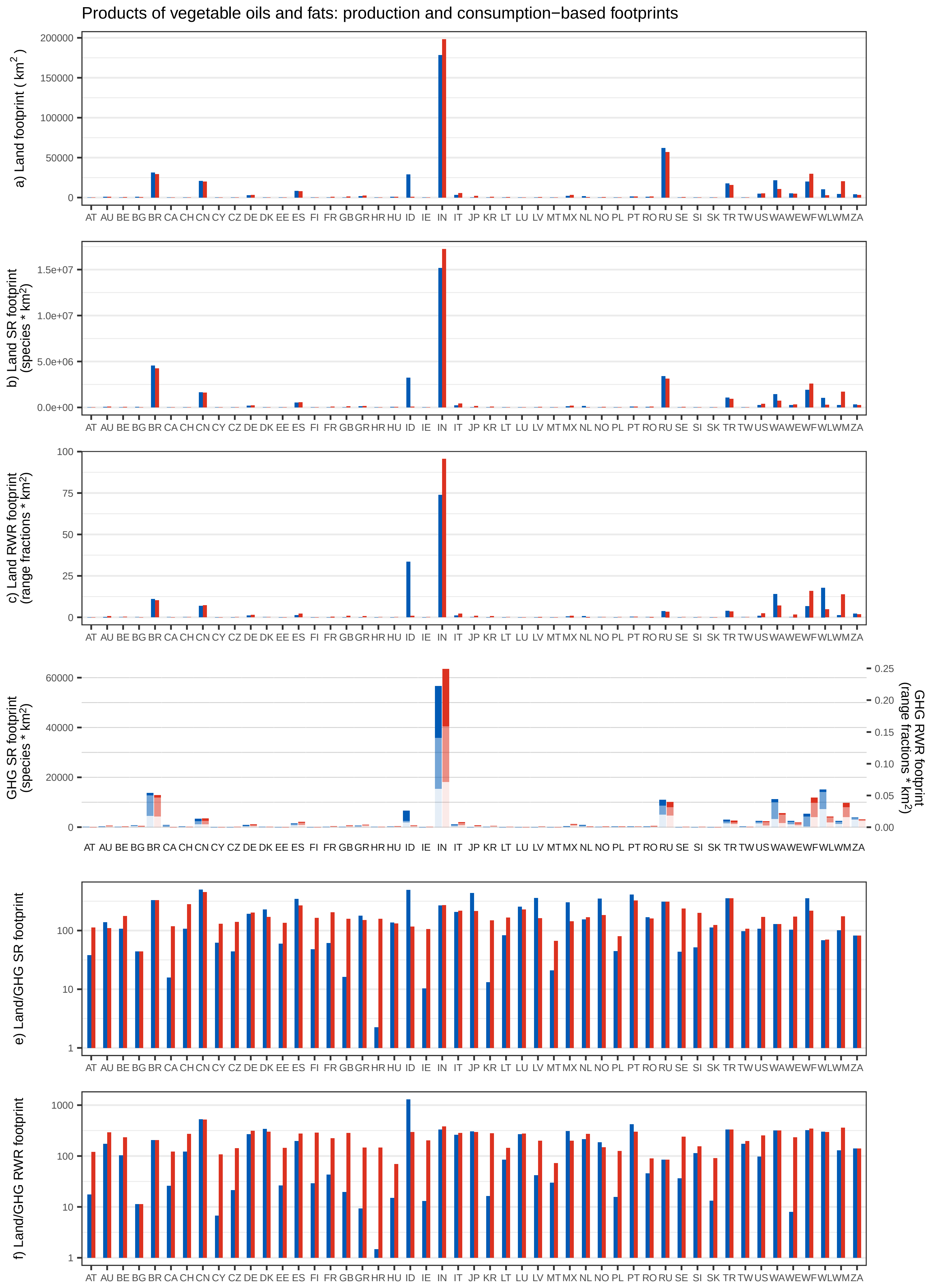

Products of vegetable oils and fats: production and consumption−based footprints
a) Land footprint ( km$^2$ )
b) Land SR footprint (species * km$^2$)
c) Land RWR footprint (range fractions * km$^2$)
GHG SR footprint (species * km$^2$)
GHG RWR footprint (range fractions * km$^2$)
e) Land/GHG SR footprint
f) Land/GHG RWR footprint
AT AU BE BG BR CA CH CN CY CZ DE DK EE ES FI FR GB GR HR HU ID IE IN IT JP KR LT LU LV MT MX NL NO PL PT RO RU SE SI SK TR TW US WA WE WF WL WM ZA

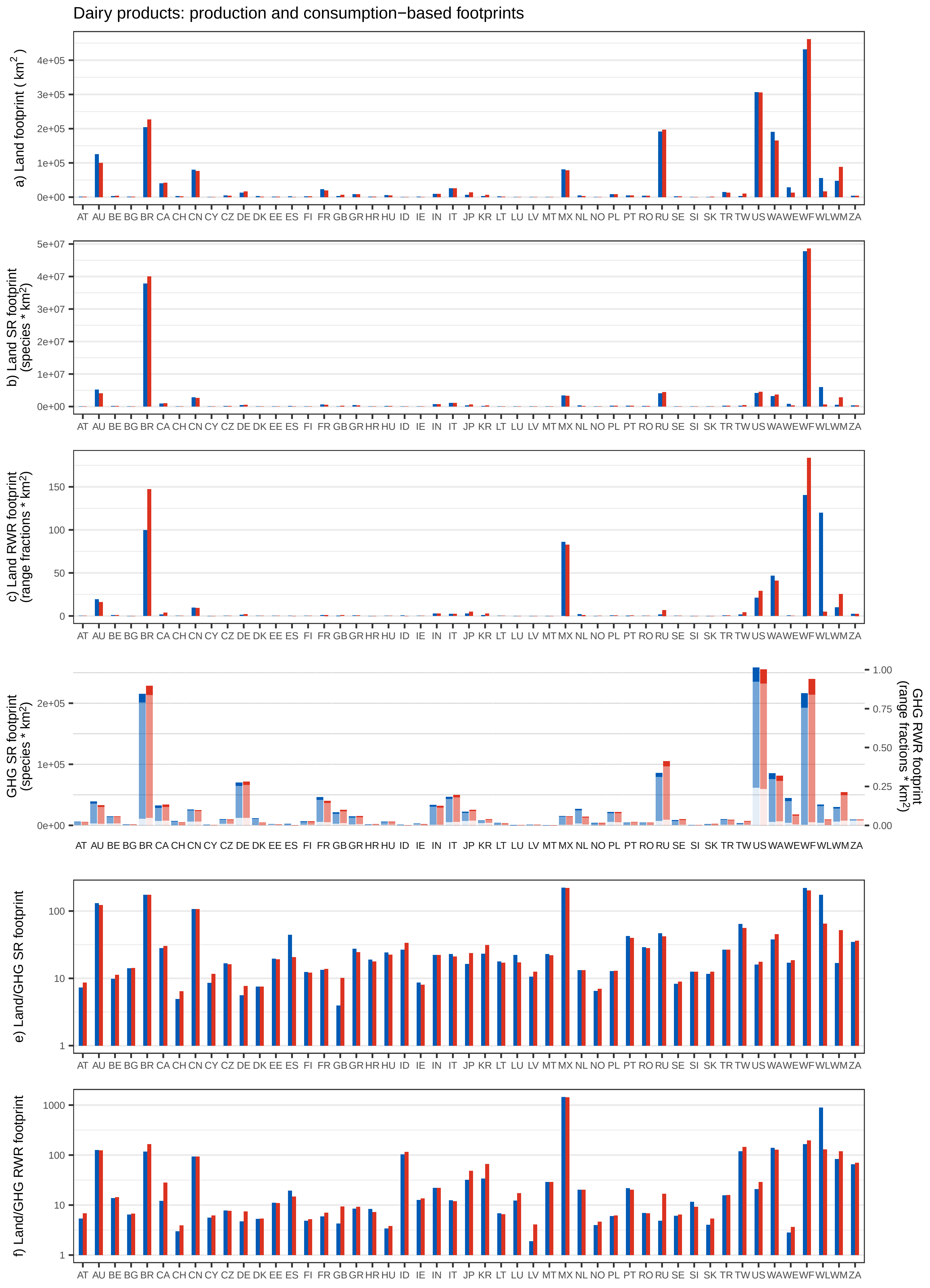
Dairy products: production and consumption−based footprints
a) Land footprint ( km$^2$ )
b) Land SR footprint (species * km$^2$)
c) Land RWR footprint (range fractions * km$^2$)
GHG SR footprint (species * km$^2$)
GHG RWR footprint (range fractions * km$^2$)
e) Land/GHG SR footprint
f) Land/GHG RWR footprint
AT AU BE BG BR CA CH CN CY CZ DE DK EE ES FI FR GB GR HR HU ID IE IN IT JP KR LT LU LV MT MX NL NO PL PT RO RU SE SI SK TR TW US WA WE WF WL WM ZA

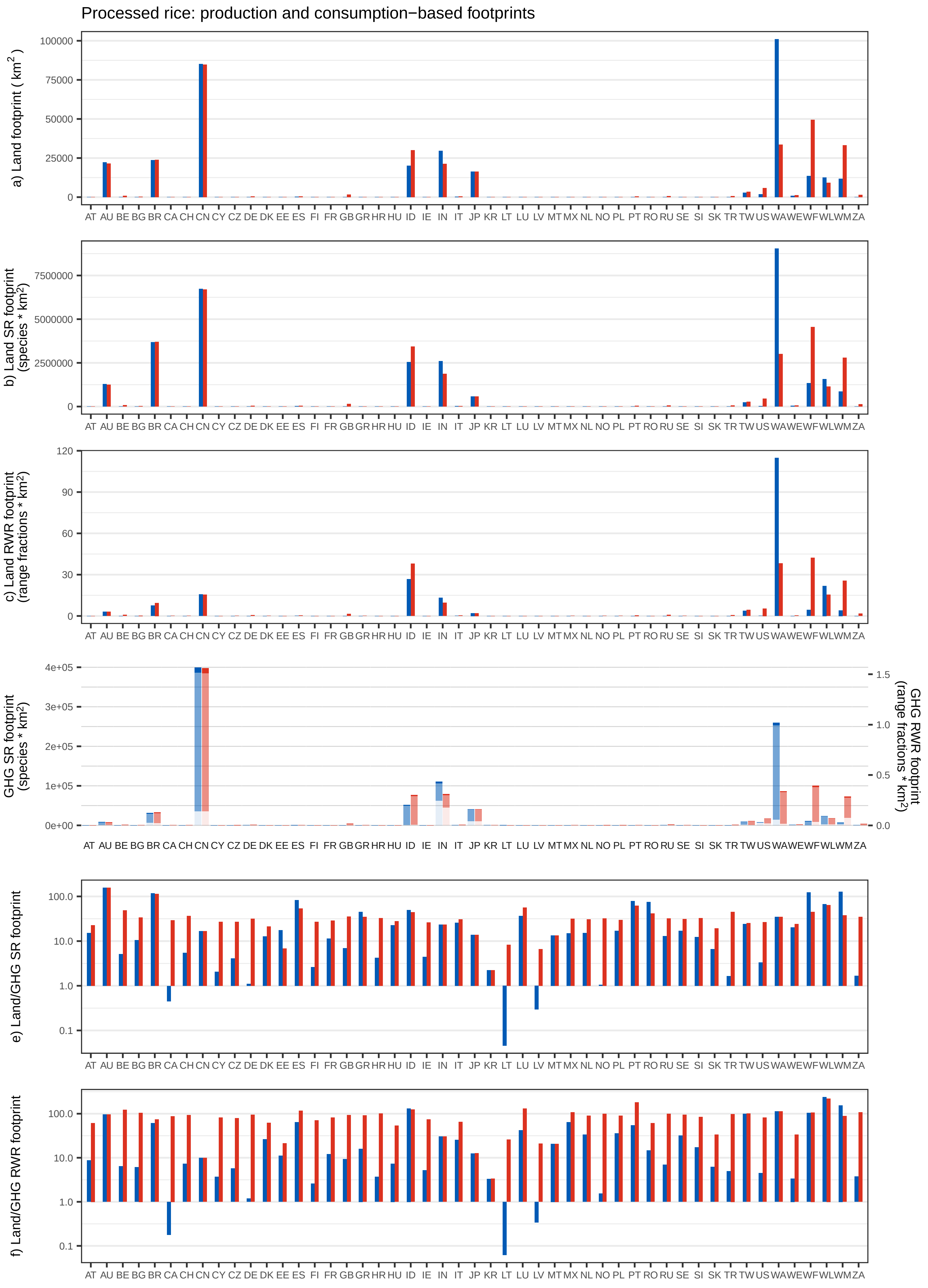
Processed rice: production and consumption−based footprints
a) Land footprint ( km$^2$ )
b) Land SR footprint (species * km$^2$)
c) Land RWR footprint (range fractions * km$^2$)
GHG SR footprint (species * km$^2$)
GHG RWR footprint (range fractions * km$^2$)
e) Land/GHG SR footprint
f) Land/GHG RWR footprint
AT AU BE BG BR CA CH CN CY CZ DE DK EE ES FI FR GB GR HR HU ID IE IN IT JP KR LT LU LV MT MX NL NO PL PT RO RU SE SI SK TR TW US WA WE WF WL WM ZA

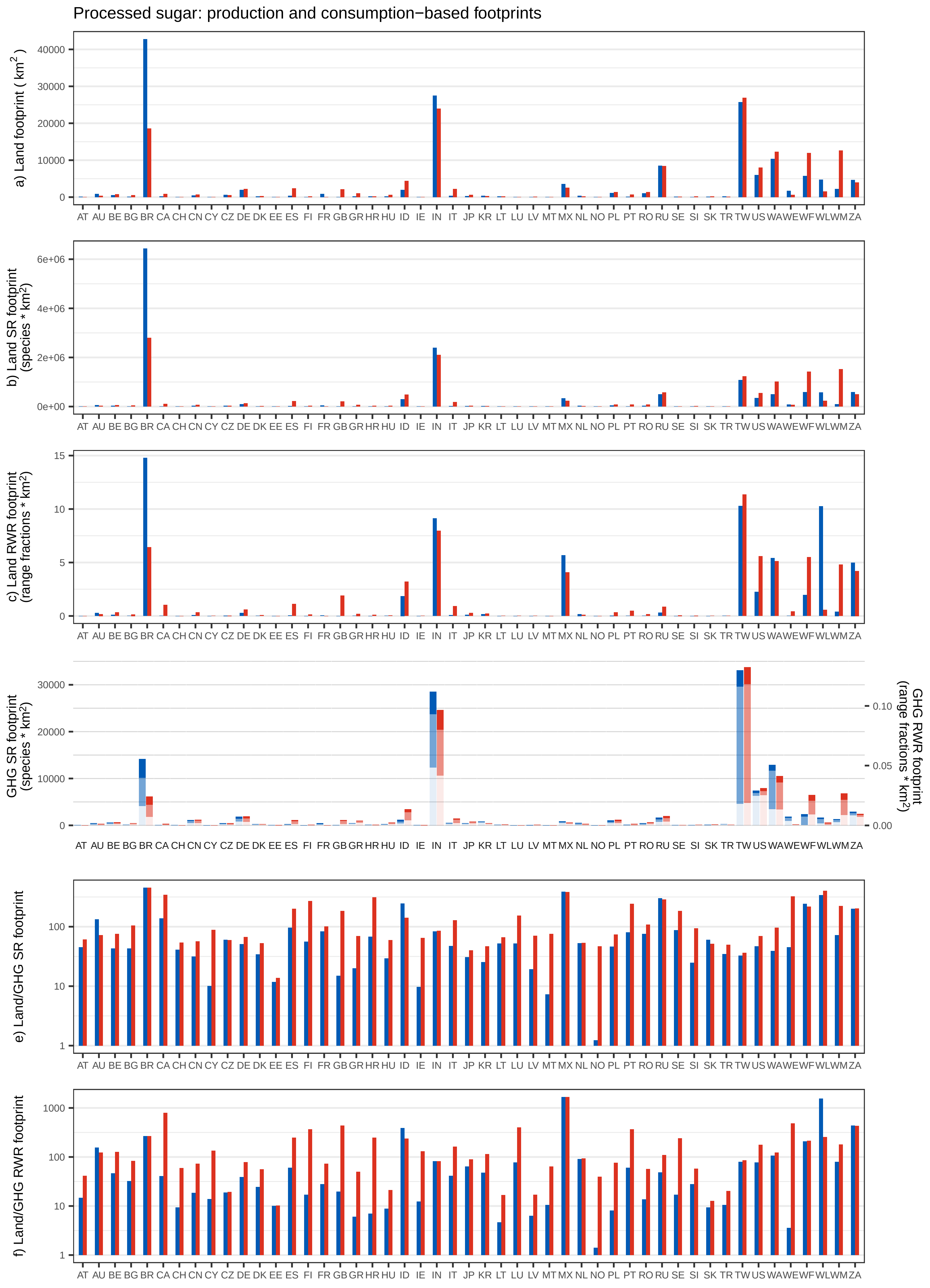

Processed sugar: production and consumption−based footprints
a) Land footprint ( km²)
b) Land SR footprint (species * km²)
c) Land RWR footprint (range fractions * km²)
GHG SR footprint (species * km²)
GHG RWR footprint (range fractions * km²)
e) Land/GHG SR footprint
f) Land/GHG RWR footprint
AT AU BE BG BR CA CH CN CY CZ DE DK EE ES FI FR GB GR HR HU ID IE IN IT JP KR LT LU LV MT MX NL NO PL PT RO RU SE SI SK TR TW US WA WE WF WL WM ZA

Food products nec: production and consumption−based footprints

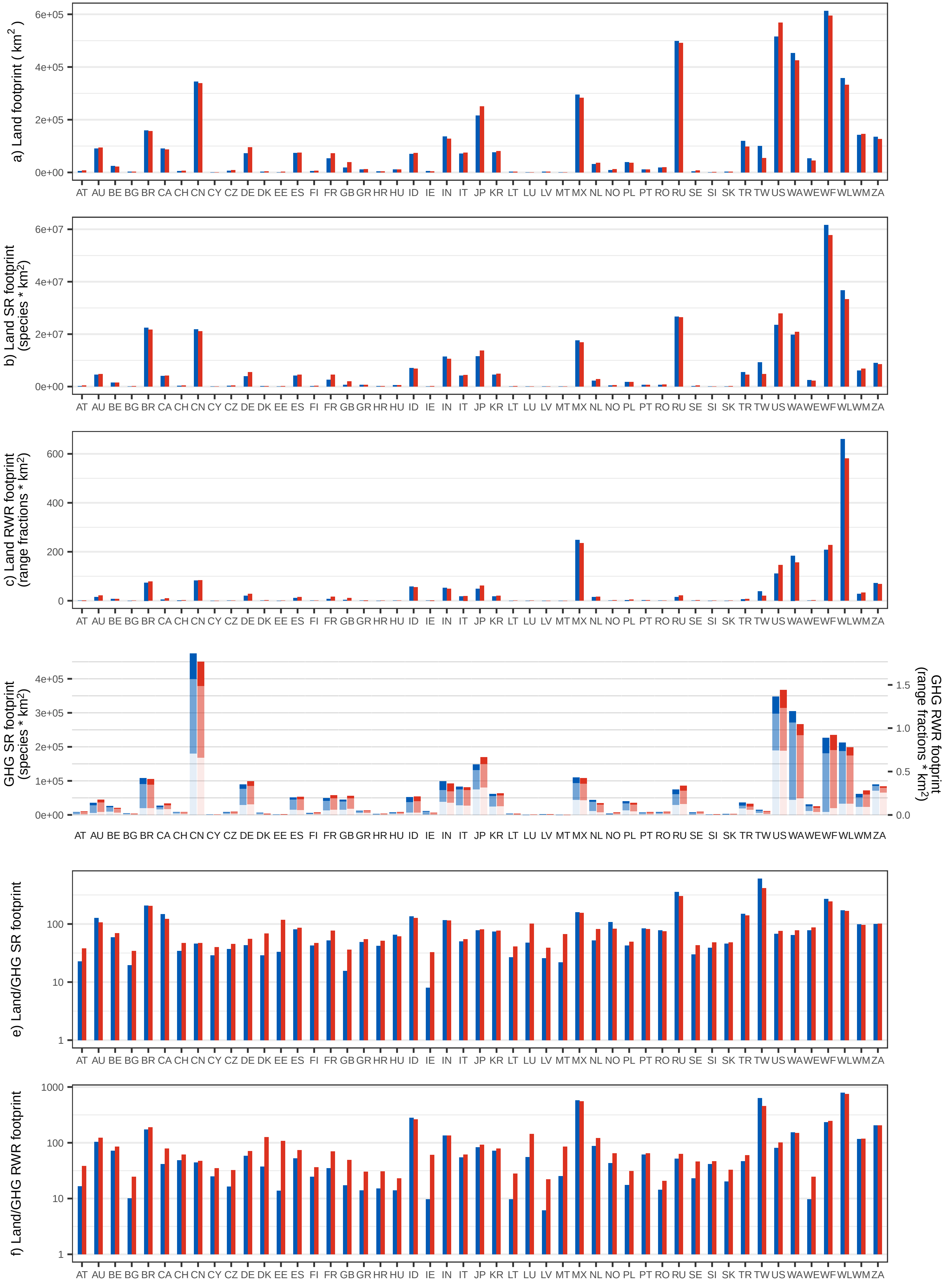

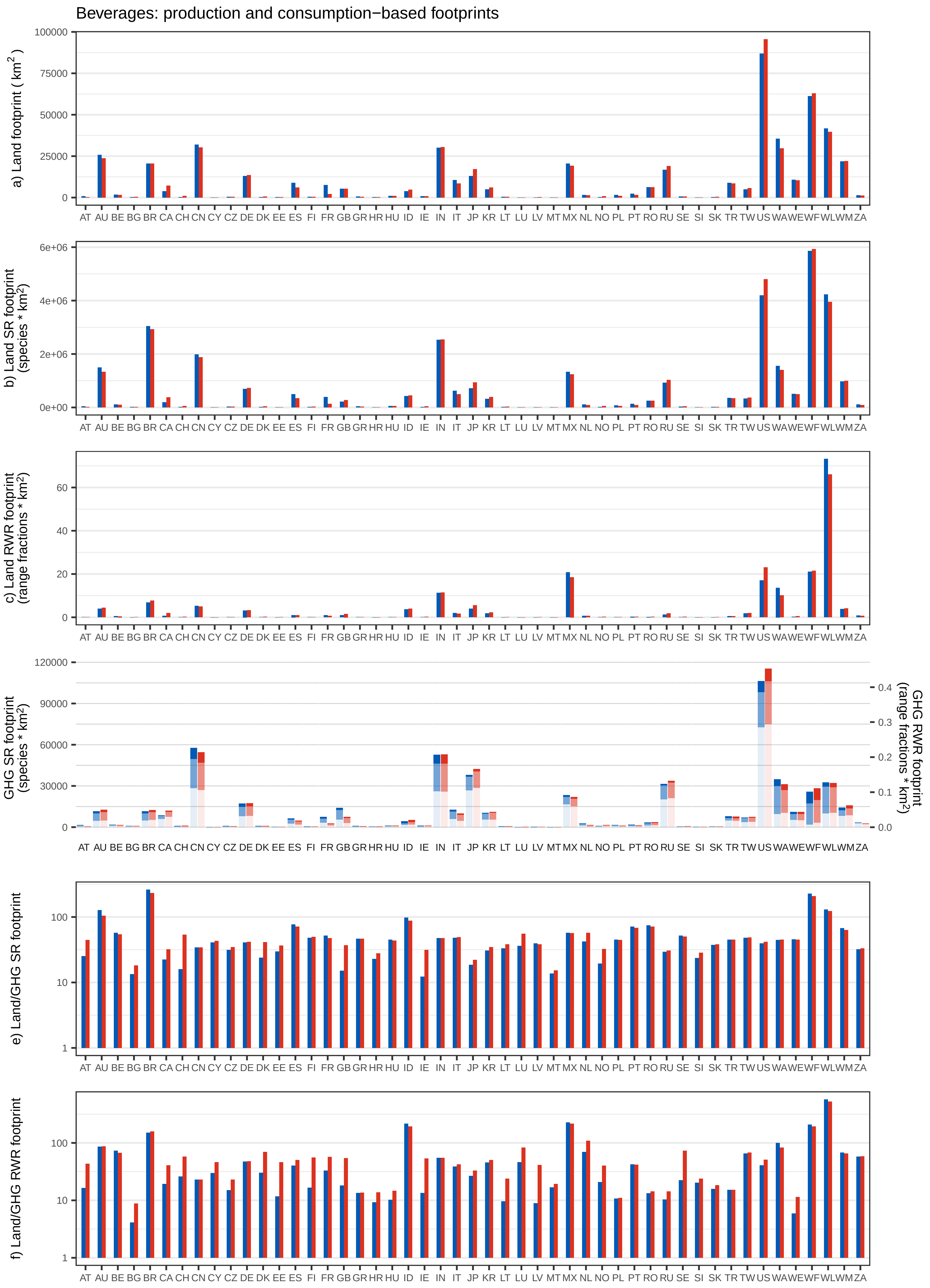
Beverages: production and consumption−based footprints
a) Land footprint ( km2 )
b) Land SR footprint (species * km2)
c) Land RWR footprint (range fractions * km2)
GHG SR footprint (species * km2)
GHG RWR footprint (range fractions * km2)
e) Land/GHG SR footprint
f) Land/GHG RWR footprint
AT AU BE BG BR CA CH CN CY CZ DE DK EE ES FI FR GB GR HR HU ID IE IN IT JP KR LT LU LV MT MX NL NO PL PT RO RU SE SI SK TR TW US WA WE WF WL WM ZA

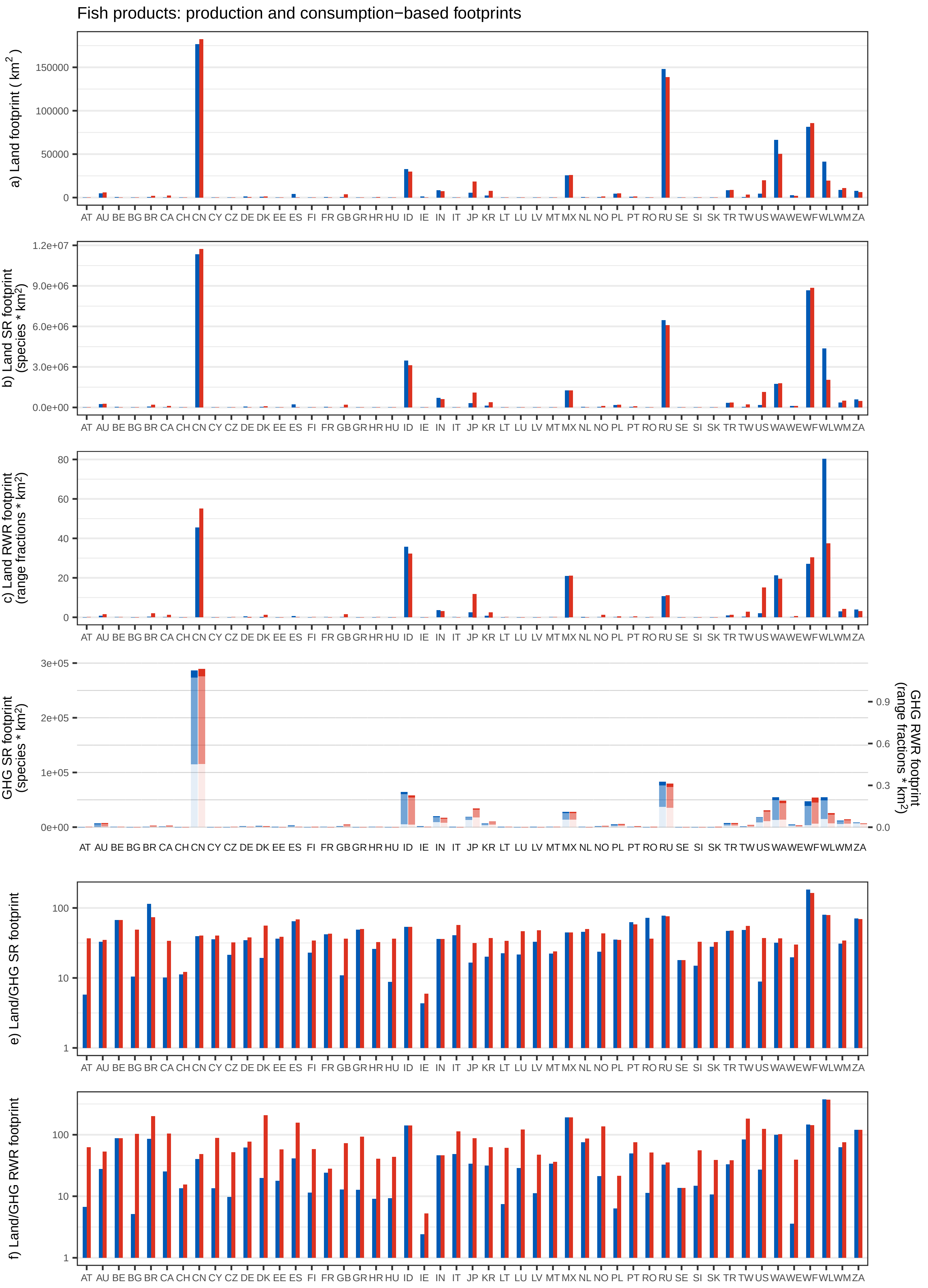
Fish products: production and consumption−based footprints
a) Land footprint ( km²)
150000
100000
50000
0
b) Land SR footprint (species * km²)
1.2e+07
9.0e+06
6.0e+06
3.0e+06
0.0e+00
c) Land RWR footprint (range fractions * km²)
80
60
40
20
0
GHG SR footprint (species * km²)
3e+05
2e+05
1e+05
0e+00
GHG RWR footprint (range fractions * km²)
0.9
0.6
0.3
0.0
e) Land/GHG SR footprint
100
10
1
f) Land/GHG RWR footprint
100
10
1
AT AU BE BG BR CA CH CN CY CZ DE DK EE ES FI FR GB GR HR HU ID IE IN IT JP KR LT LU LV MT MX NL NO PL PT RO RU SE SI SK TR TW US WA WE WF WL WM ZA

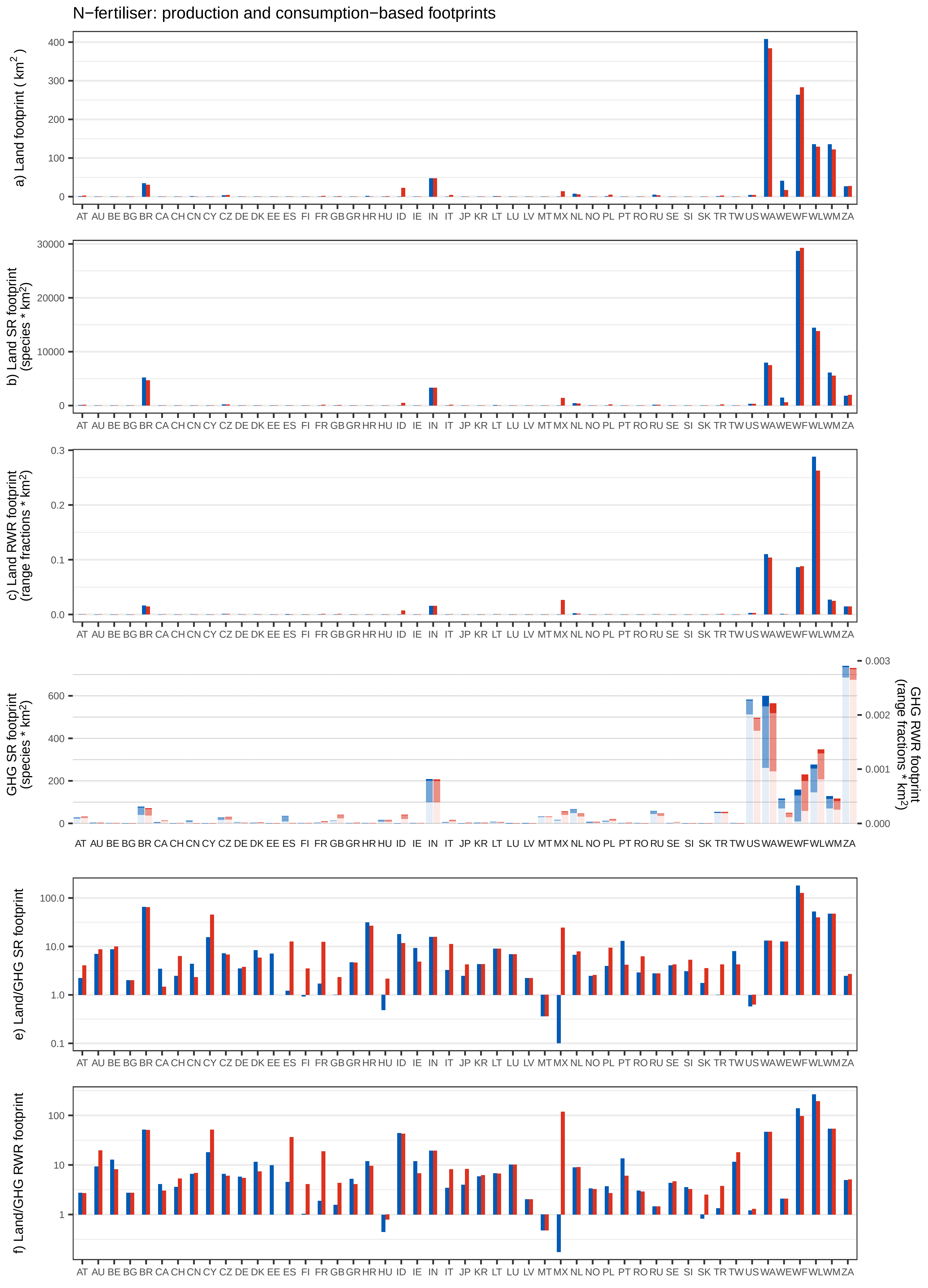
N−fertiliser: production and consumption−based footprints
a) Land footprint ( km$^2$ )
0
100
200
300
400
b) Land SR footprint (species * km$^2$)
0
10000
20000
30000
c) Land RWR footprint (range fractions * km$^2$)
0.0
0.1
0.2
0.3
GHG SR footprint (species * km$^2$)
0
200
400
600
GHG RWR footprint (range fractions * km$^2$)
0.000
0.001
0.002
0.003
e) Land/GHG SR footprint
0.1
1.0
10.0
100.0
f) Land/GHG RWR footprint
1
10
100
AT AU BE BG BR CA CH CN CY CZ DE DK EE ES FI FR GB GR HR HU ID IE IN IT JP KR LT LU LV MT MX NL NO PL PT RO RU SE SI SK TR TW US WA WE WF WL WM ZA

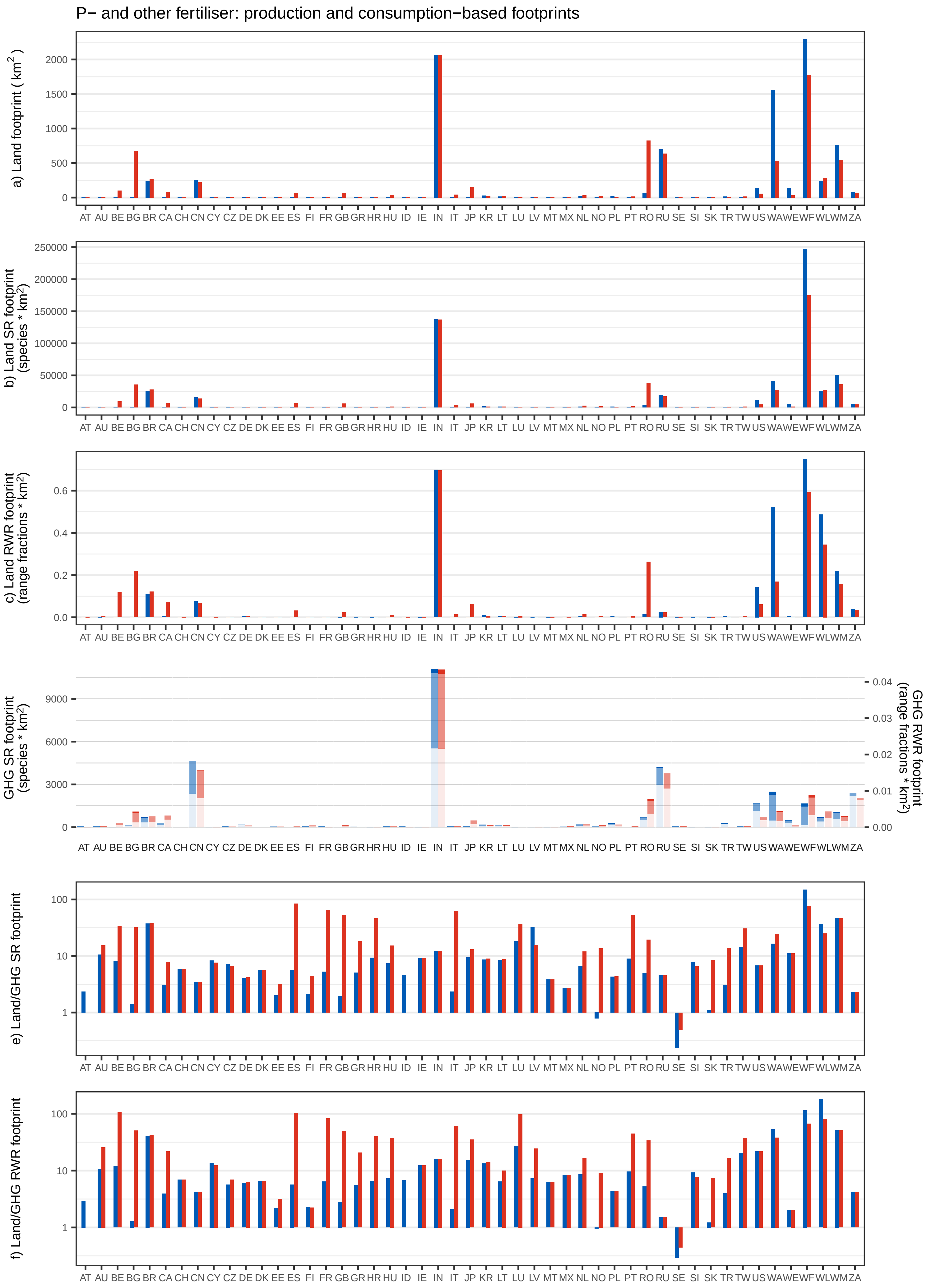
P− and other fertiliser: production and consumption−based footprints
a) Land footprint ( km²)
b) Land SR footprint (species * km²)
c) Land RWR footprint (range fractions * km²)
GHG SR footprint (species * km²)
GHG RWR footprint (range fractions * km²)
e) Land/GHG SR footprint
f) Land/GHG RWR footprint
AT AU BE BG BR CA CH CN CY CZ DE DK EE ES FI FR GB GR HR HU ID IE IN IT JP KR LT LU LV MT MX NL NO PL PT RO RU SE SI SK TR TW US WA WE WF WL WM ZA

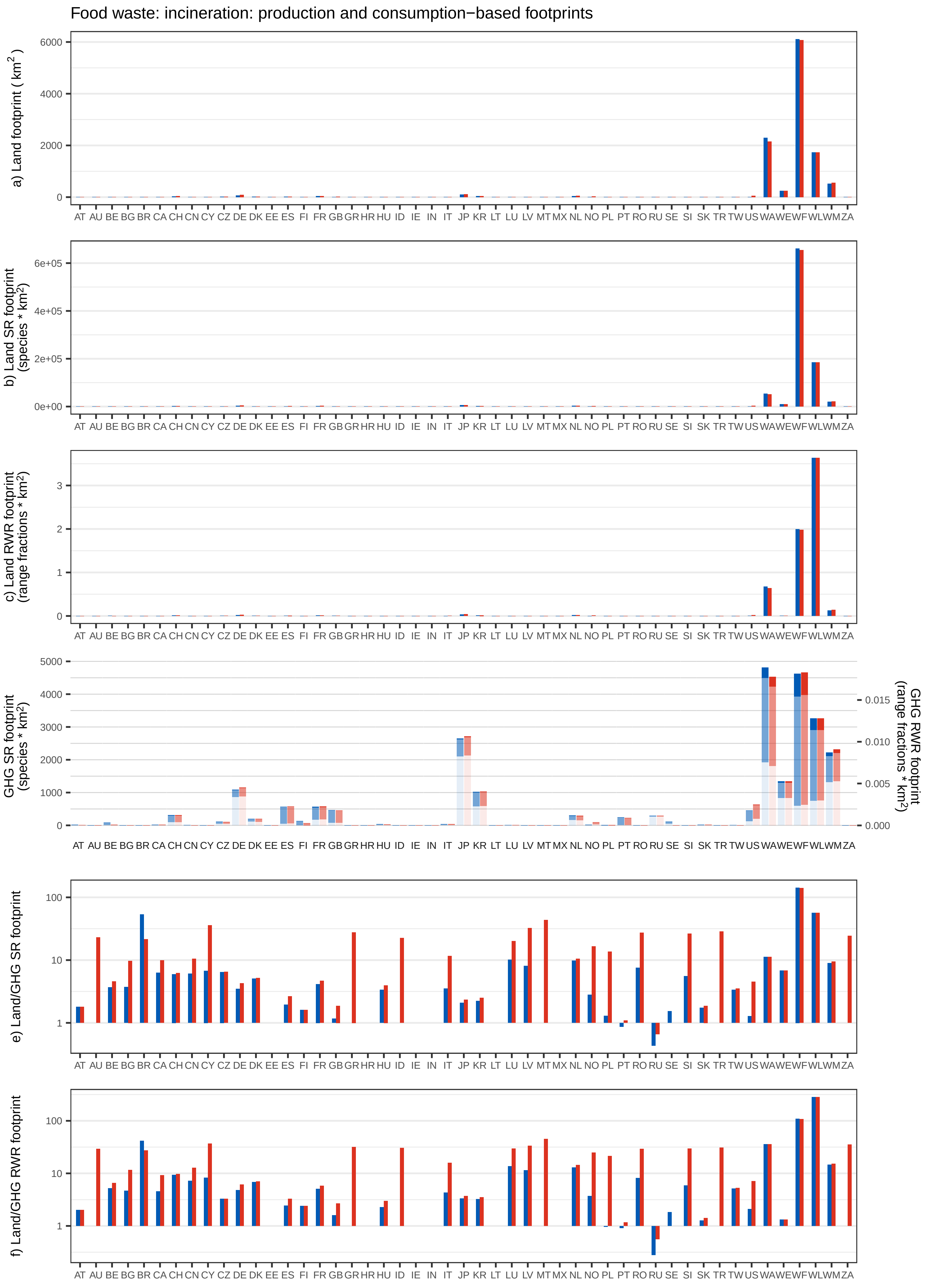
Food waste: incineration: production and consumption−based footprints
a) Land footprint ( km$^2$ )
0
2000
4000
6000
b) Land SR footprint (species * km$^2$)
0e+00
2e+05
4e+05
6e+05
c) Land RWR footprint (range fractions * km$^2$)
0
1
2
3
GHG SR footprint (species * km$^2$)
0
1000
2000
3000
4000
5000
GHG RWR footprint (range fractions * km$^2$)
0.000
0.005
0.010
0.015
e) Land/GHG SR footprint
1
10
100
f) Land/GHG RWR footprint
1
10
100
AT AU BE BG BR CA CH CN CY CZ DE DK EE ES FI FR GB GR HR HU ID IE IN IT JP KR LT LU LV MT MX NL NO PL PT RO RU SE SI SK TR TW US WA WE WF WL WM ZA

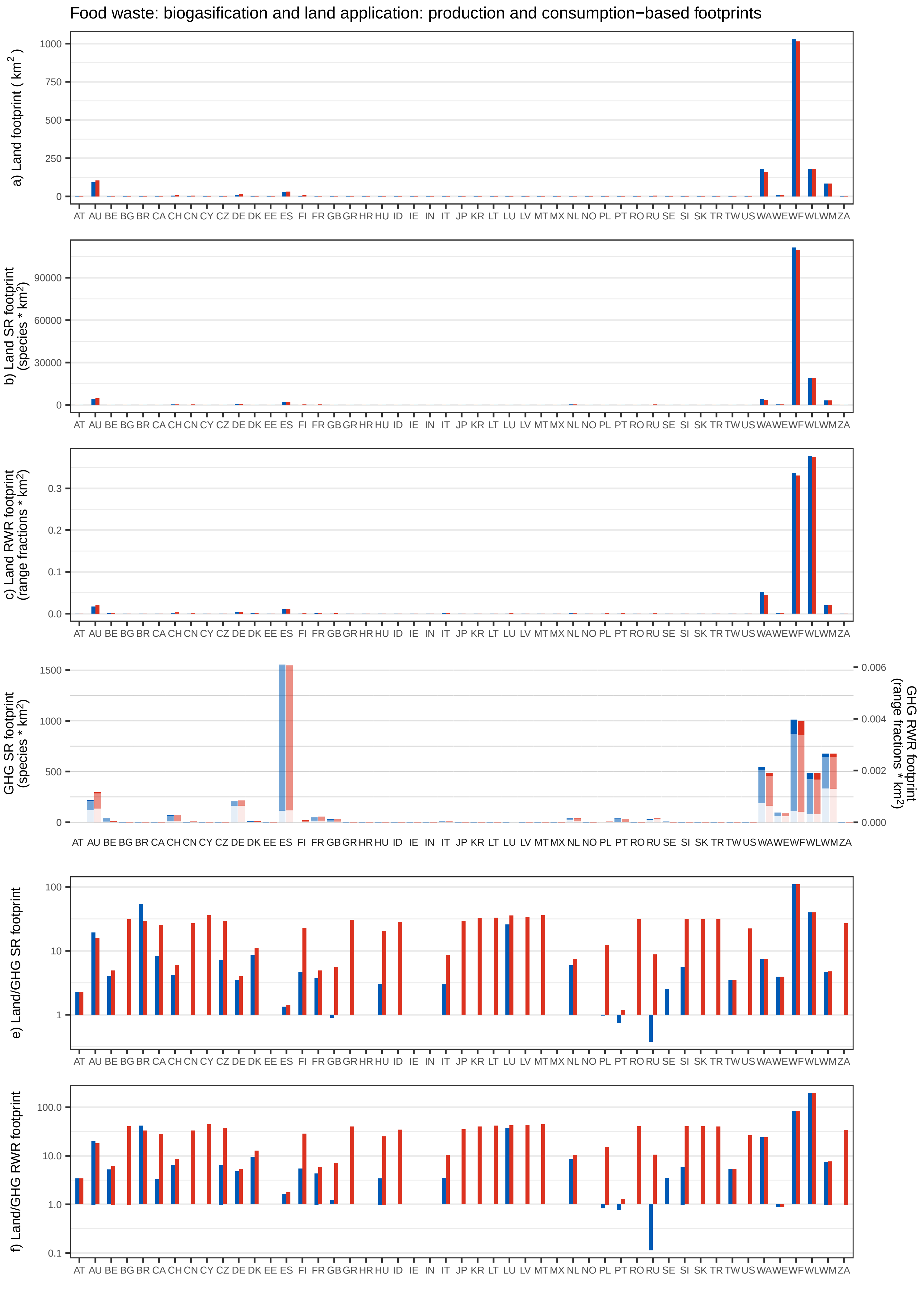

Food waste: biogasification and land application: production and consumption−based footprints
a) Land footprint ( km$^2$ )
b) Land SR footprint (species * km$^2$)
c) Land RWR footprint (range fractions * km$^2$)
GHG SR footprint (species * km$^2$)
GHG RWR footprint (range fractions * km$^2$)
e) Land/GHG SR footprint
f) Land/GHG RWR footprint
AT AU BE BG BR CA CH CN CY CZ DE DK EE ES FI FR GB GR HR HU ID IE IN IT JP KR LT LU LV MT MX NL NO PL PT RO RU SE SI SK TR TW US WA WE WF WL WM ZA

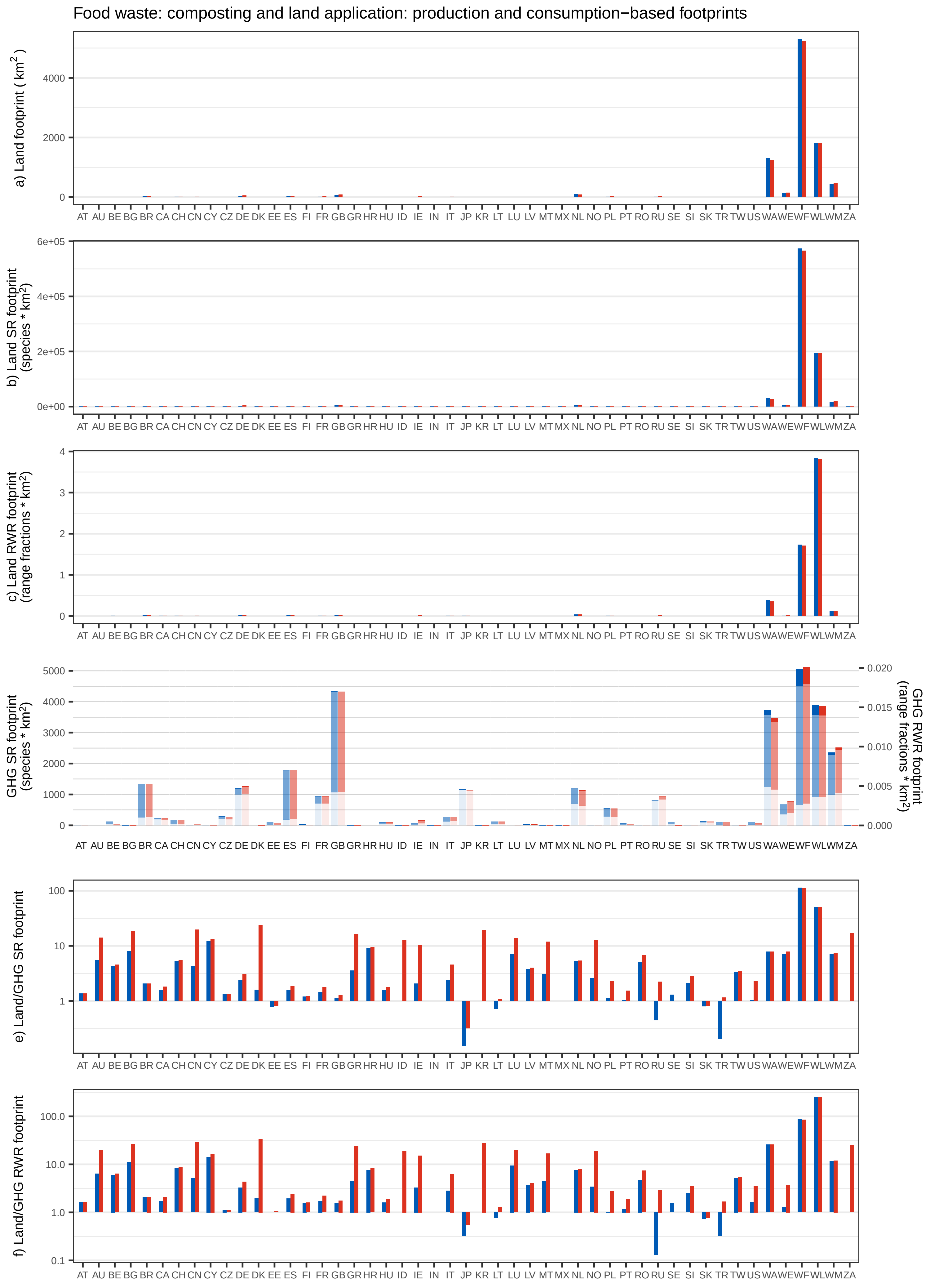

Food waste: composting and land application: production and consumption−based footprints
a) Land footprint ( km2 )
b) Land SR footprint (species * km2)
c) Land RWR footprint (range fractions * km2)
GHG SR footprint (species * km2)
GHG RWR footprint (range fractions * km2)
e) Land/GHG SR footprint
f) Land/GHG RWR footprint
AT AU BE BG BR CA CH CN CY CZ DE DK EE ES FI FR GB GR HR HU ID IE IN IT JP KR LT LU LV MT MX NL NO PL PT RO RU SE SI SK TR TW US WA WE WF WL WM ZA

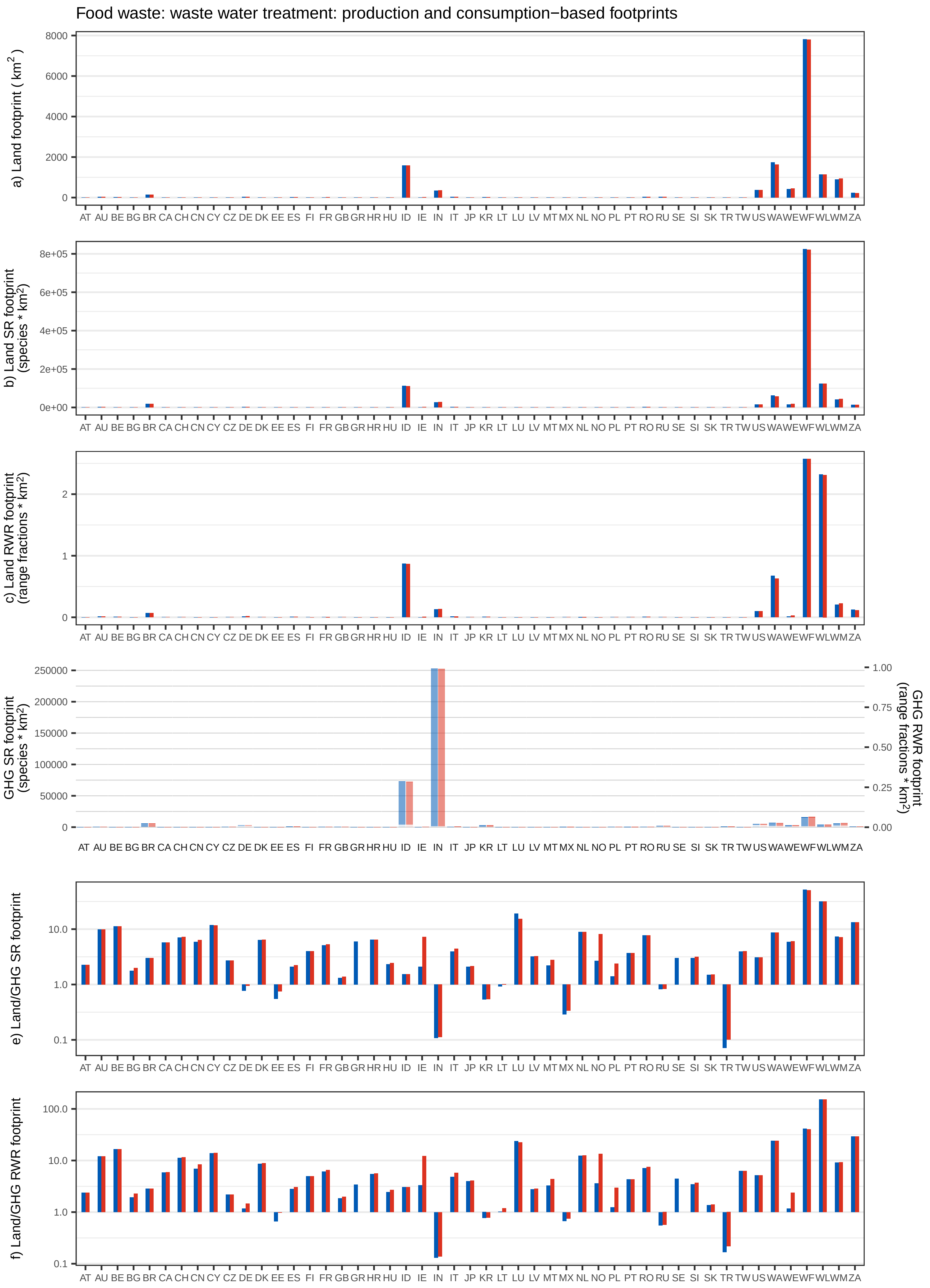

Food waste: waste water treatment: production and consumption−based footprints
a) Land footprint ( km²)
b) Land SR footprint (species * km²)
c) Land RWR footprint (range fractions * km²)
GHG SR footprint (species * km²)
GHG RWR footprint (range fractions * km²)
e) Land/GHG SR footprint
f) Land/GHG RWR footprint
AT AU BE BG BR CA CH CN CY CZ DE DK EE ES FI FR GB GR HR HU ID IE IN IT JP KR LT LU LV MT MX NL NO PL PT RO RU SE SI SK TR TW US WA WE WF WL WM ZA

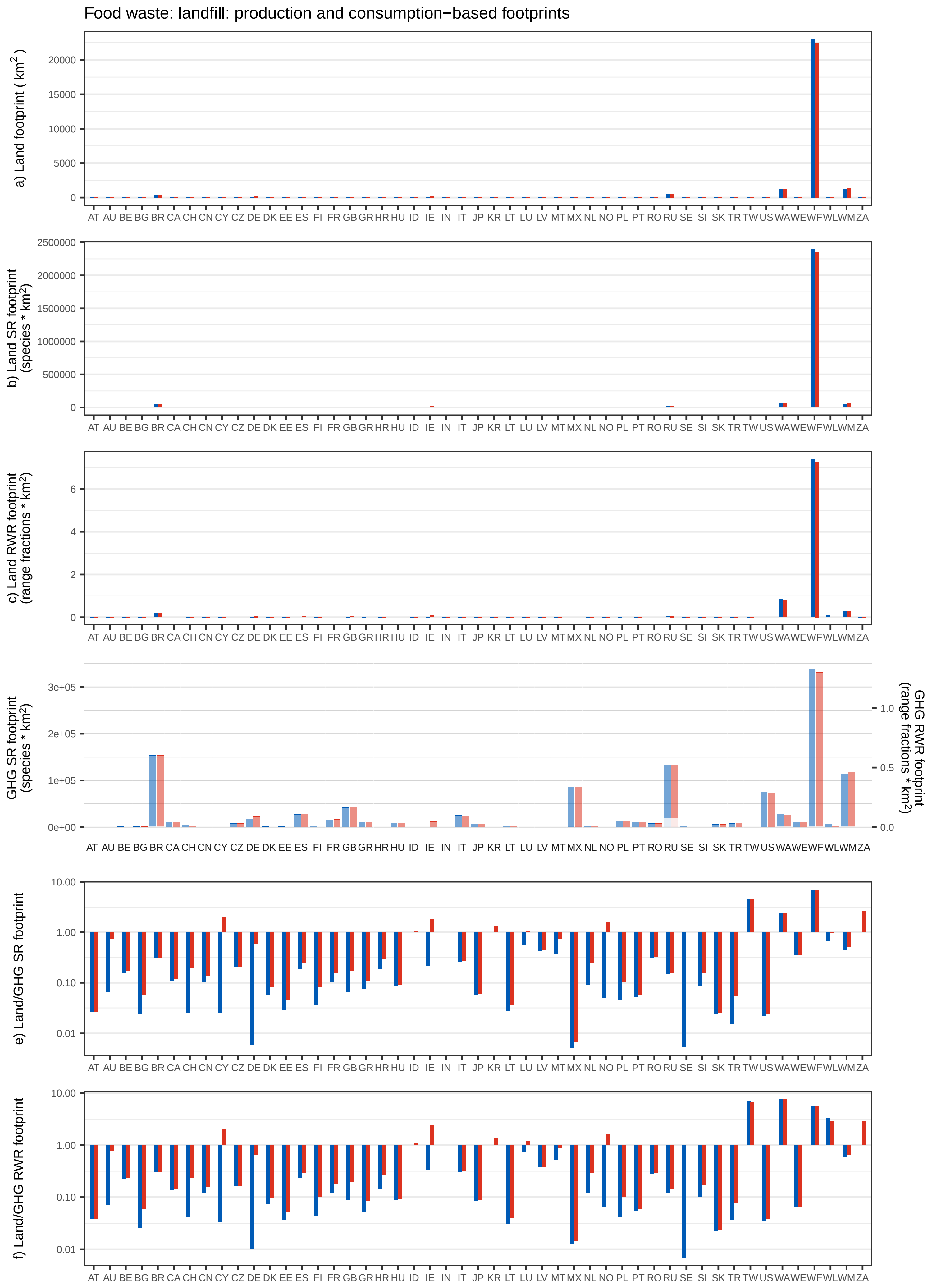
Food waste: landfill: production and consumption−based footprints
a) Land footprint ( km$^2$ )
b) Land SR footprint (species * km$^2$)
c) Land RWR footprint (range fractions * km$^2$)
GHG SR footprint (species * km$^2$)
GHG RWR footprint (range fractions * km$^2$)
e) Land/GHG SR footprint
f) Land/GHG RWR footprint
AT AU BE BG BR CA CH CN CY CZ DE DK EE ES FI FR GB GR HR HU ID IE IN IT JP KR LT LU LV MT MX NL NO PL PT RO RU SE SI SK TR TW US WA WE WF WL WM ZA
