## Supplementary Figure 7 for "Impacts of global food supply on biodiversity via land use and climate change"

Paddy rice: production−based footprints per square km

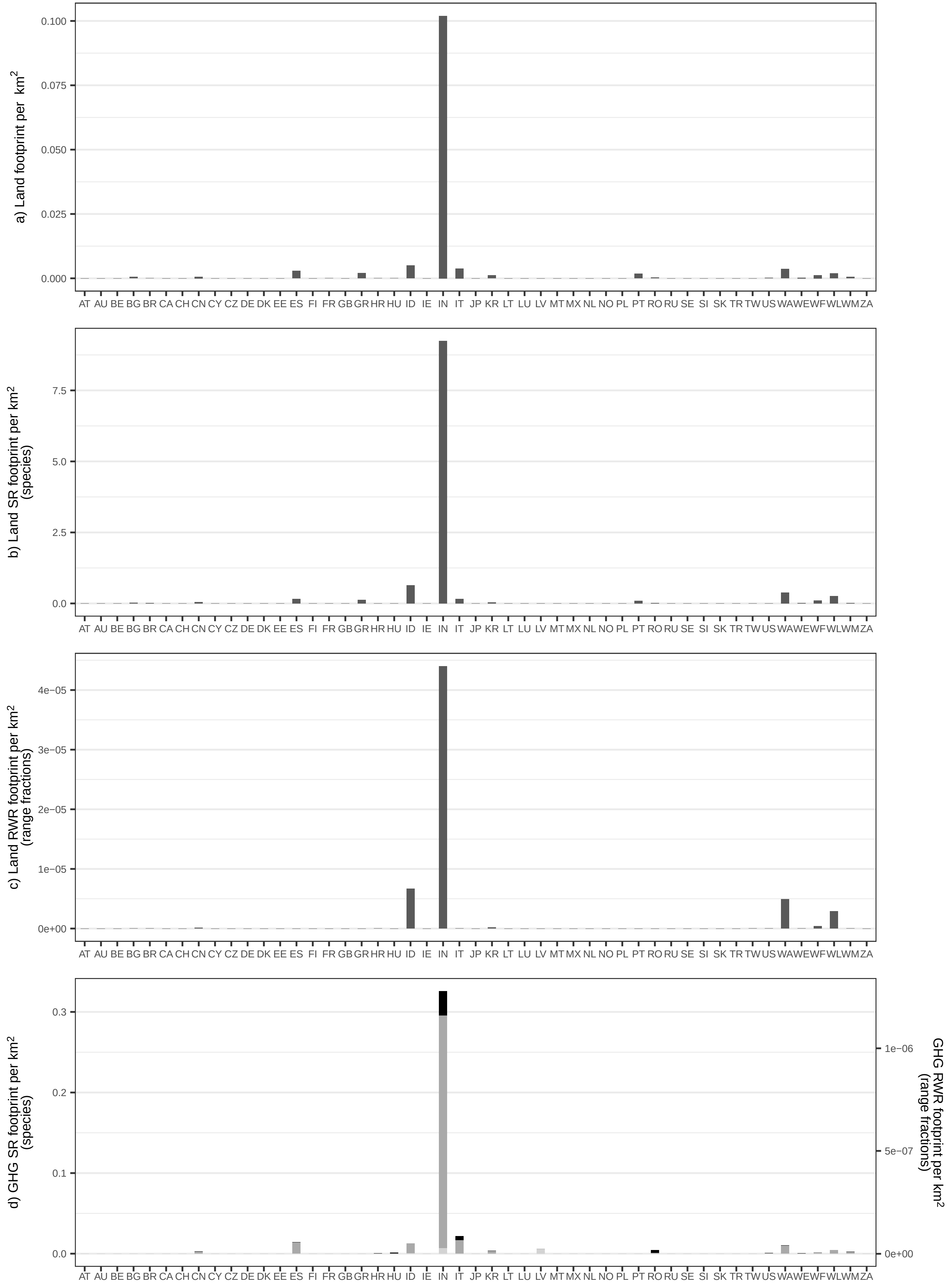

Wheat: production−based footprints per square km

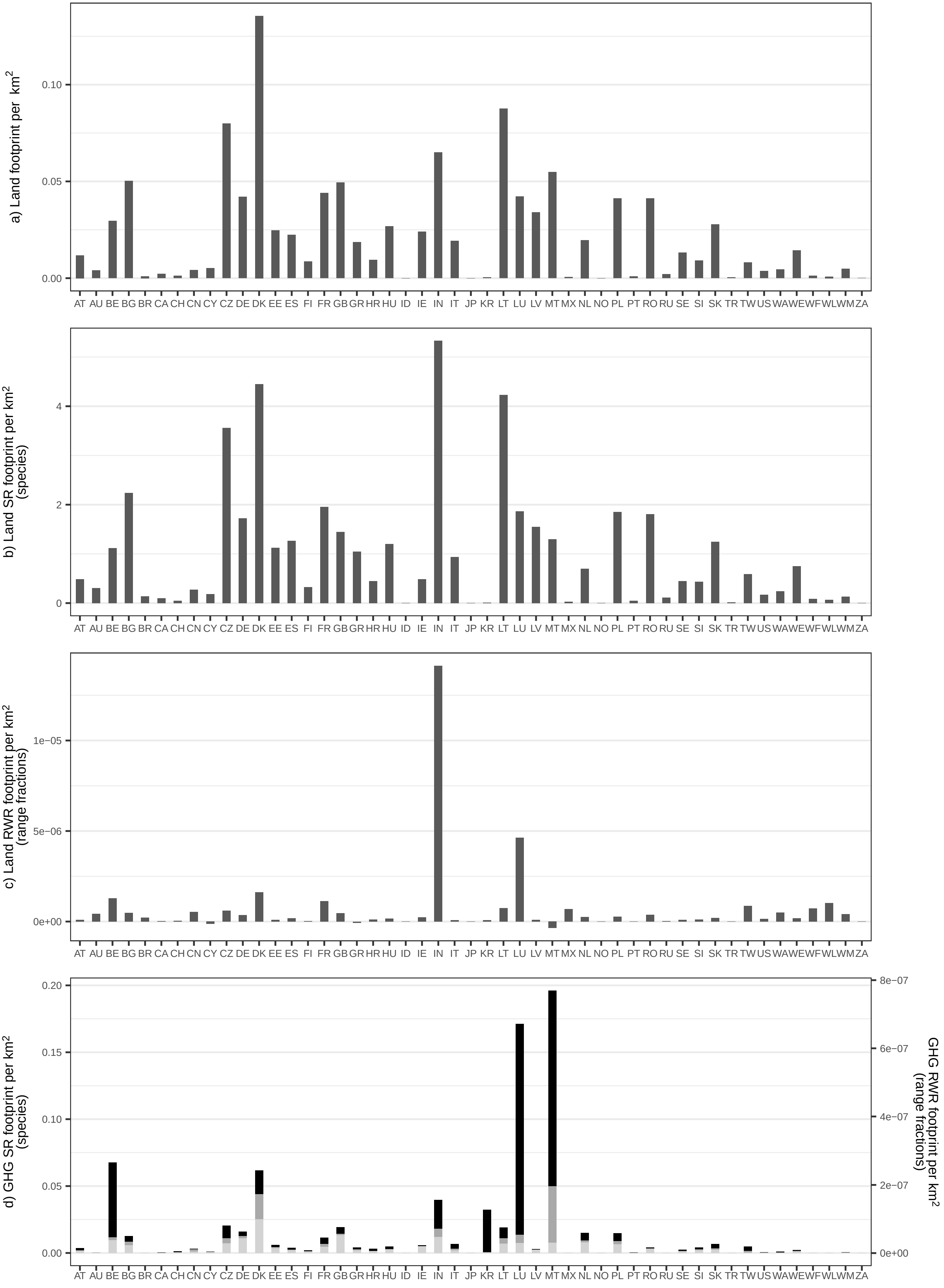

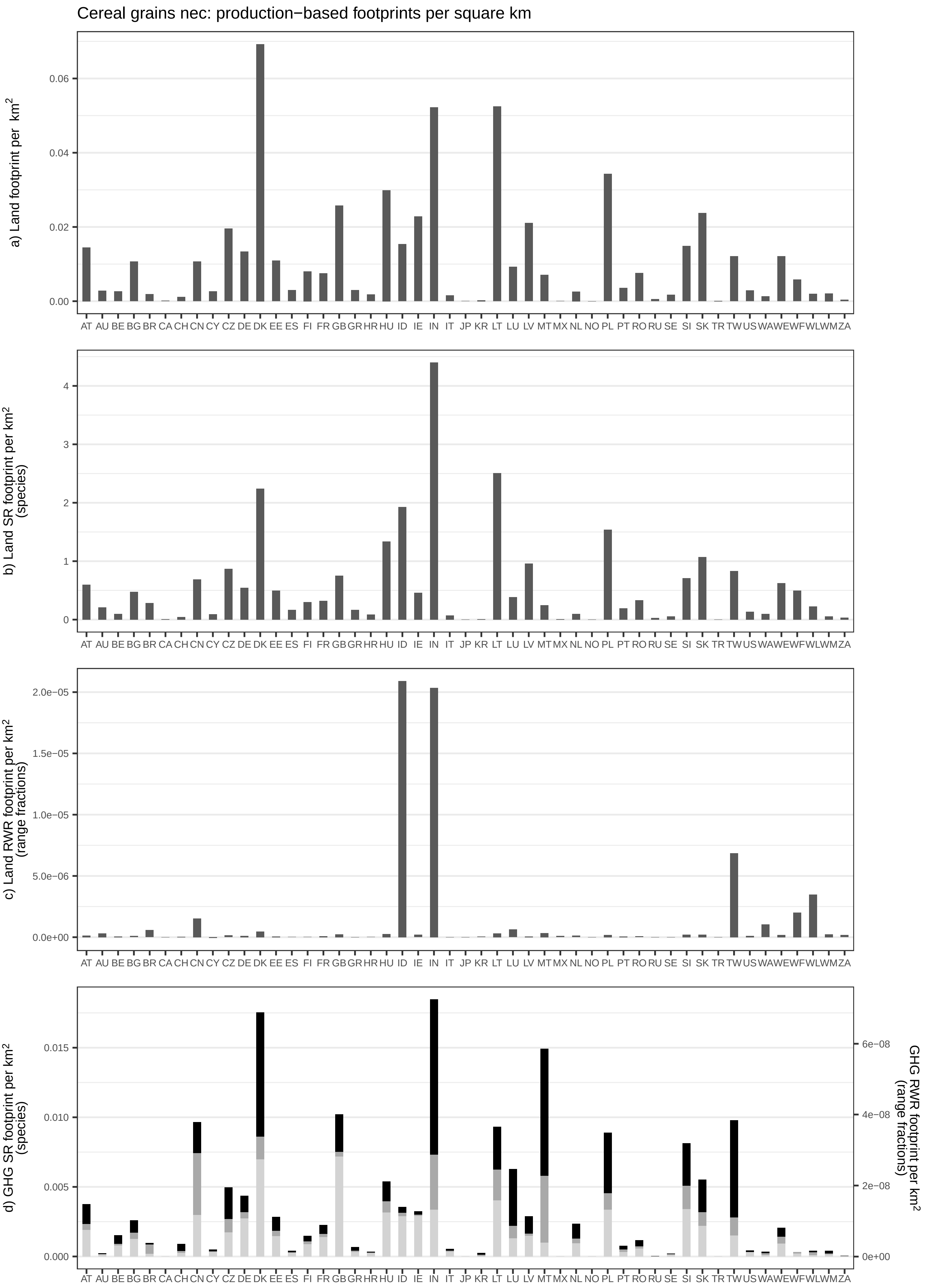

Cereal grains nec: production−based footprints per square km
a) Land footprint per km²
0.00
0.02
0.04
0.06
b) Land SR footprint per km² (species)
0
1
2
3
4
c) Land RWR footprint per km² (range fractions)
0.0e+00
5.0e−06
1.0e−05
1.5e−05
2.0e−05
d) GHG SR footprint per km² (species)
0.000
0.005
0.010
0.015
GHG RWR footprint per km² (range fractions)
0e+00
2e−08
4e−08
6e−08
AT AU BE BG BR CA CH CN CY CZ DE DK EE ES FI FR GB GR HR HU ID IE IN IT JP KR LT LU LV MT MX NL NO PL PT RO RU SE SI SK TR TW US WA WE WF WL WM ZA

Veg/Fruit/Nuts: production−based footprints per square km

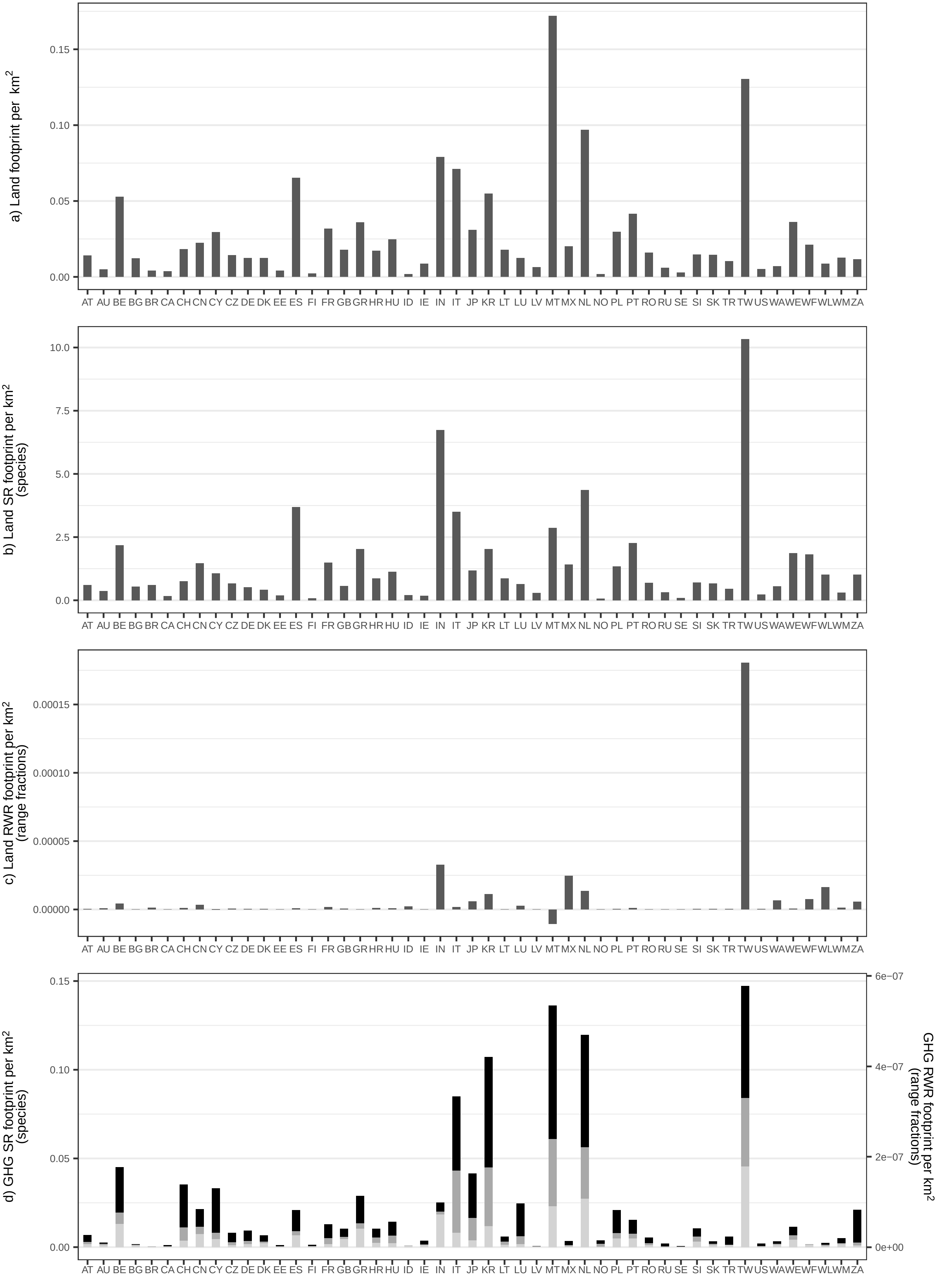

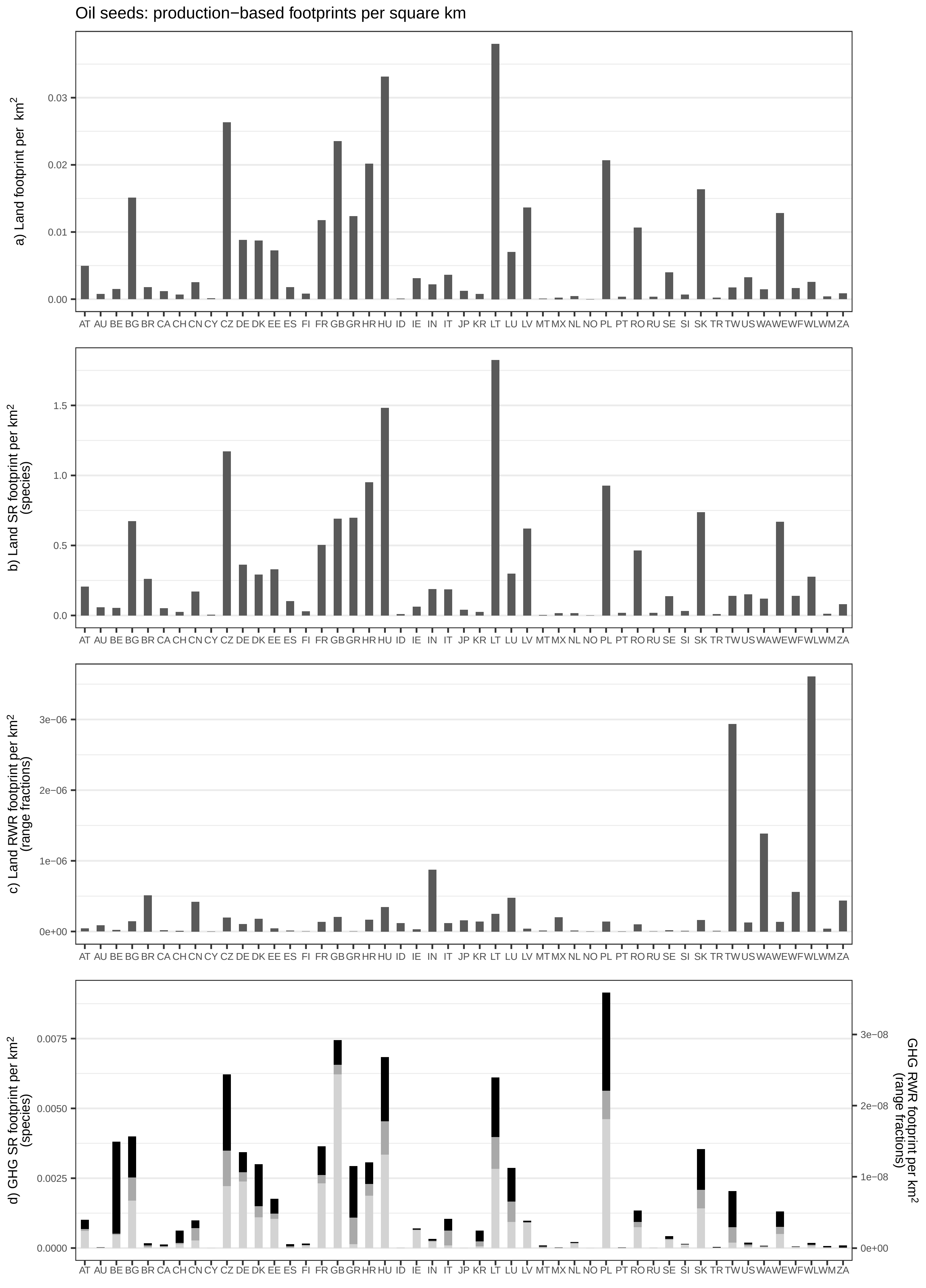
Oil seeds: production−based footprints per square km
a) Land footprint per km²
b) Land SR footprint per km² (species)
c) Land RWR footprint per km² (range fractions)
d) GHG SR footprint per km² (species)
GHG RWR footprint per km² (range fractions)
AT AU BE BG BR CA CH CN CY CZ DE DK EE ES FI FR GB GR HR HU ID IE IN IT JP KR LT LU LV MT MX NL NO PL PT RO RU SE SI SK TR TW US WA WE WF WL WM ZA

Sugar beet/cane: production−based footprints per square km

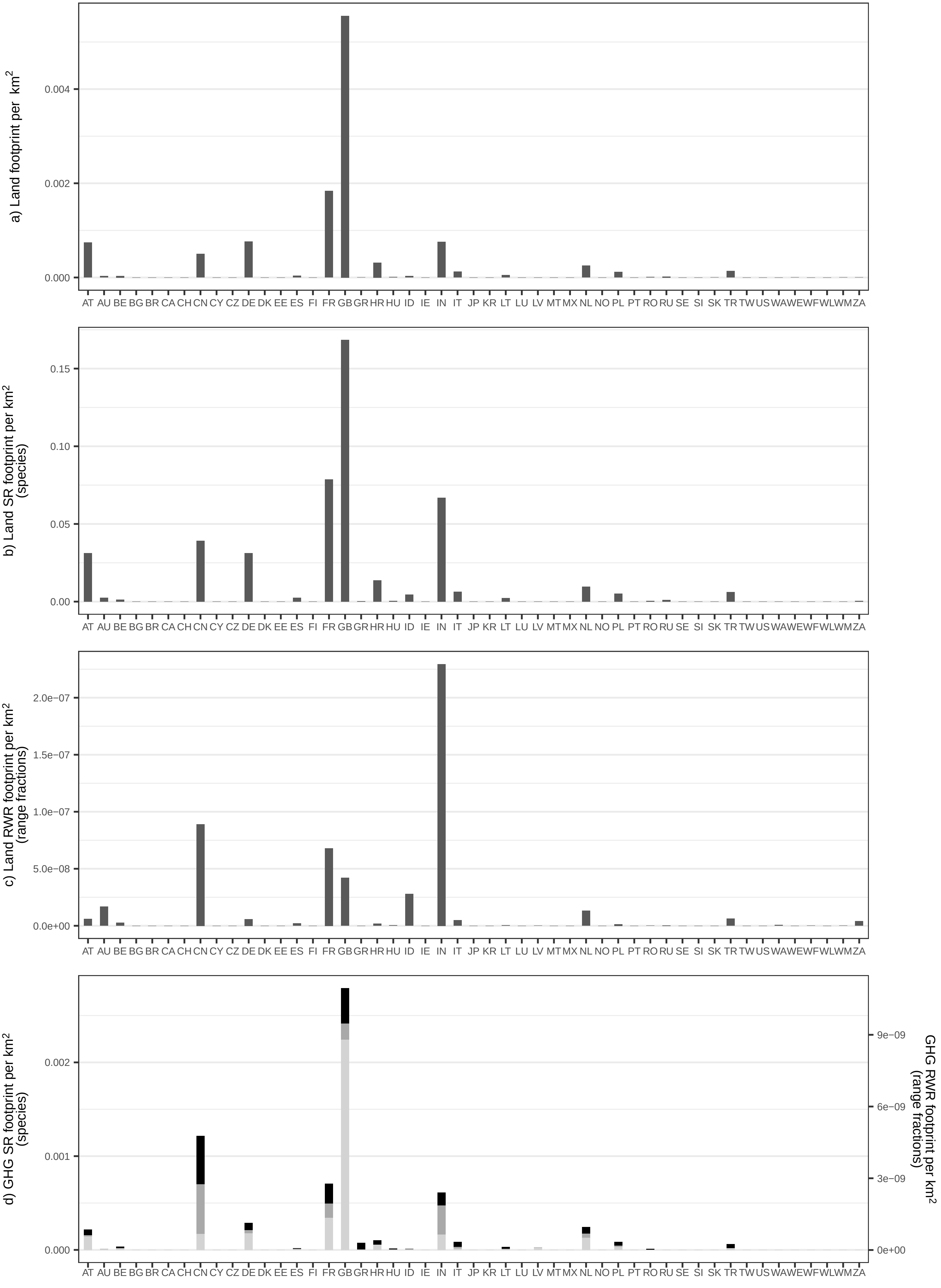

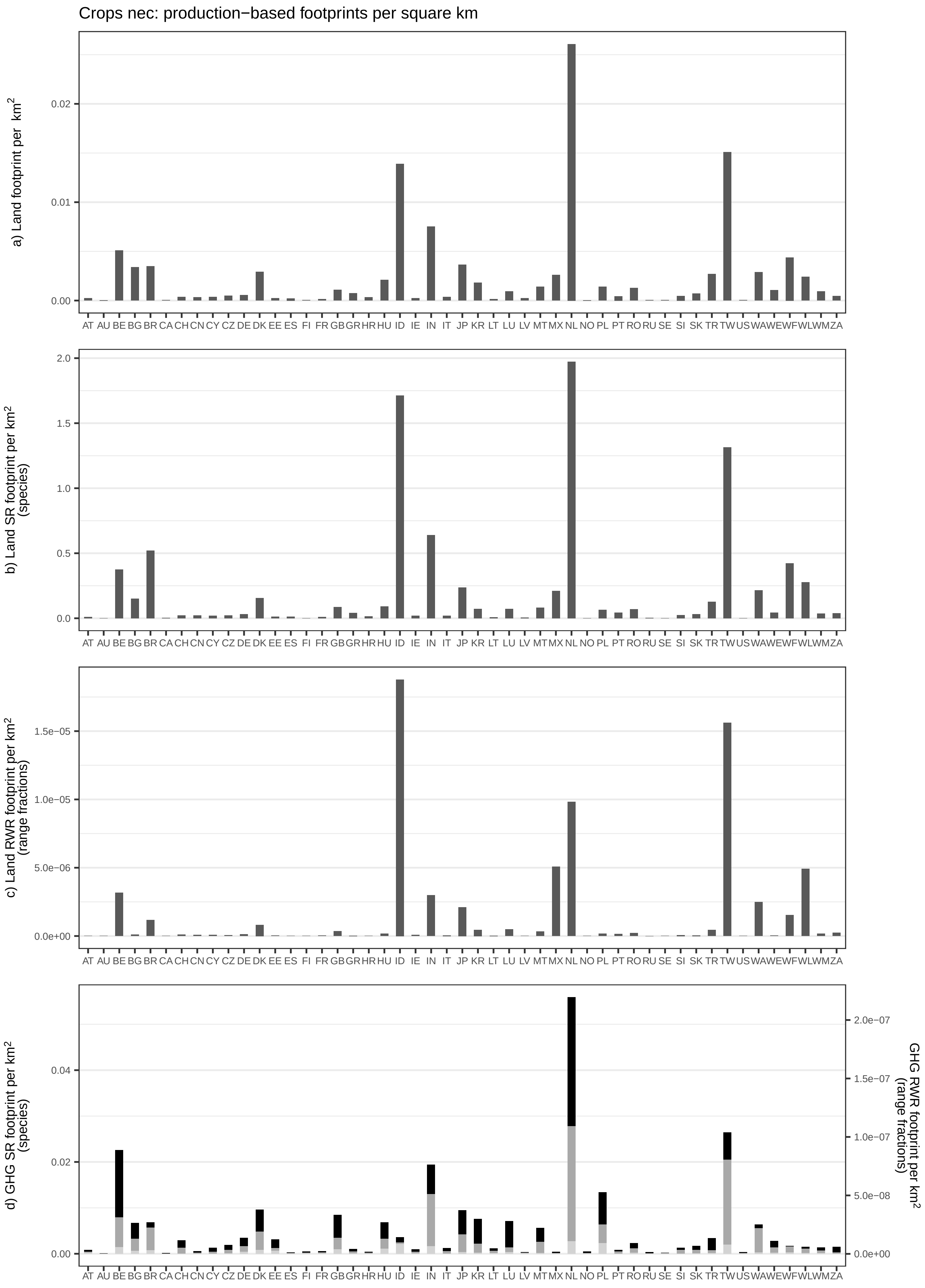

Crops nec: production−based footprints per square km
a) Land footprint per km²
b) Land SR footprint per km² (species)
c) Land RWR footprint per km² (range fractions)
d) GHG SR footprint per km² (species)
GHG RWR footprint per km² (range fractions)
AT AU BE BG BR CA CH CN CY CZ DE DK EE ES FI FR GB GR HR HU ID IE IN IT JP KR LT LU LV MT MX NL NO PL PT RO RU SE SI SK TR TW US WA WE WF WL WM ZA

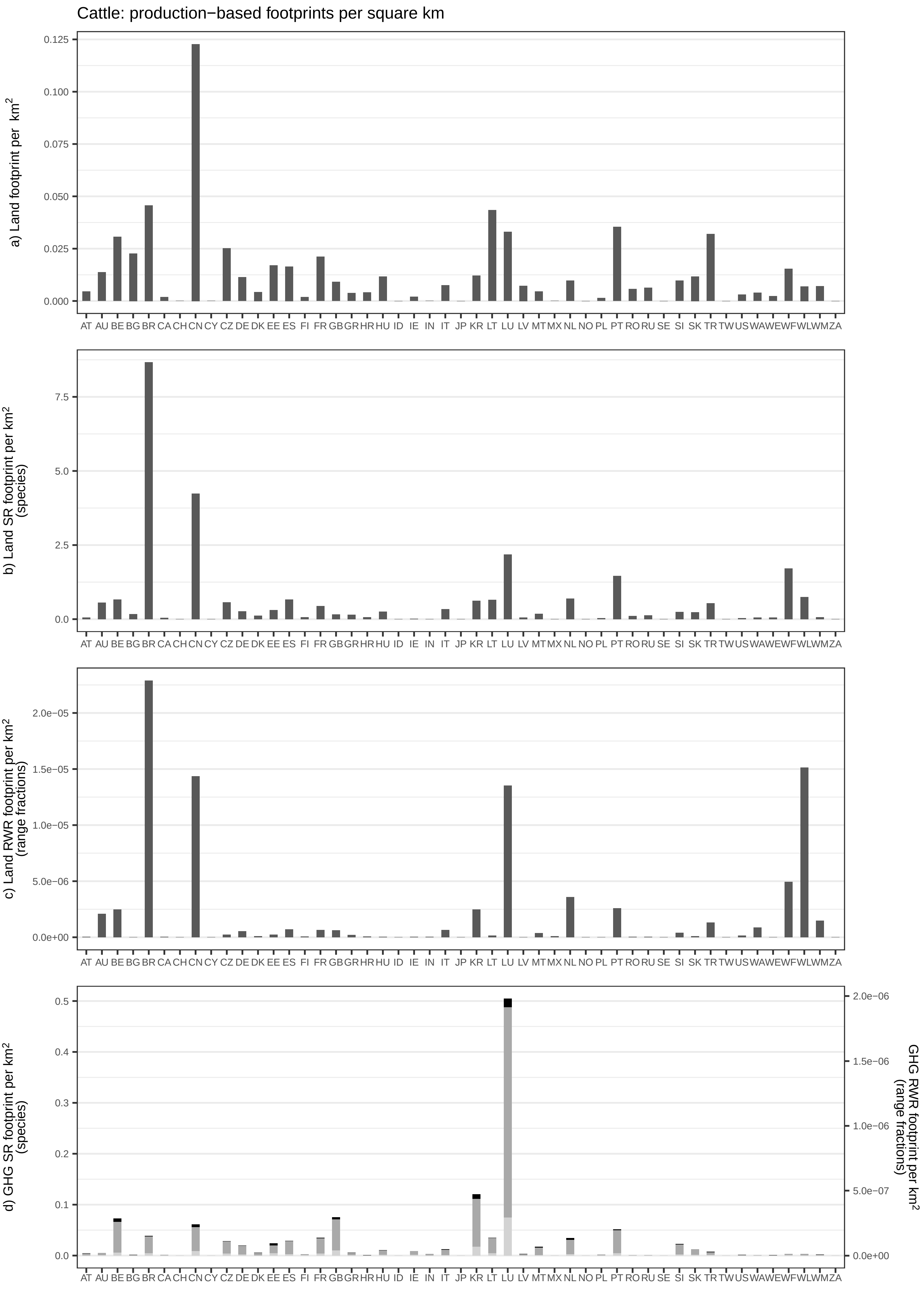

Cattle: production−based footprints per square km
a) Land footprint per km²
b) Land SR footprint per km² (species)
c) Land RWR footprint per km² (range fractions)
d) GHG SR footprint per km² (species)
GHG RWR footprint per km² (range fractions)
AT AU BE BG BR CA CH CN CY CZ DE DK EE ES FI FR GB GR HR HU ID IE IN IT JP KR LT LU LV MT MX NL NO PL PT RO RU SE SI SK TR TW US WA WE WF WL WM ZA

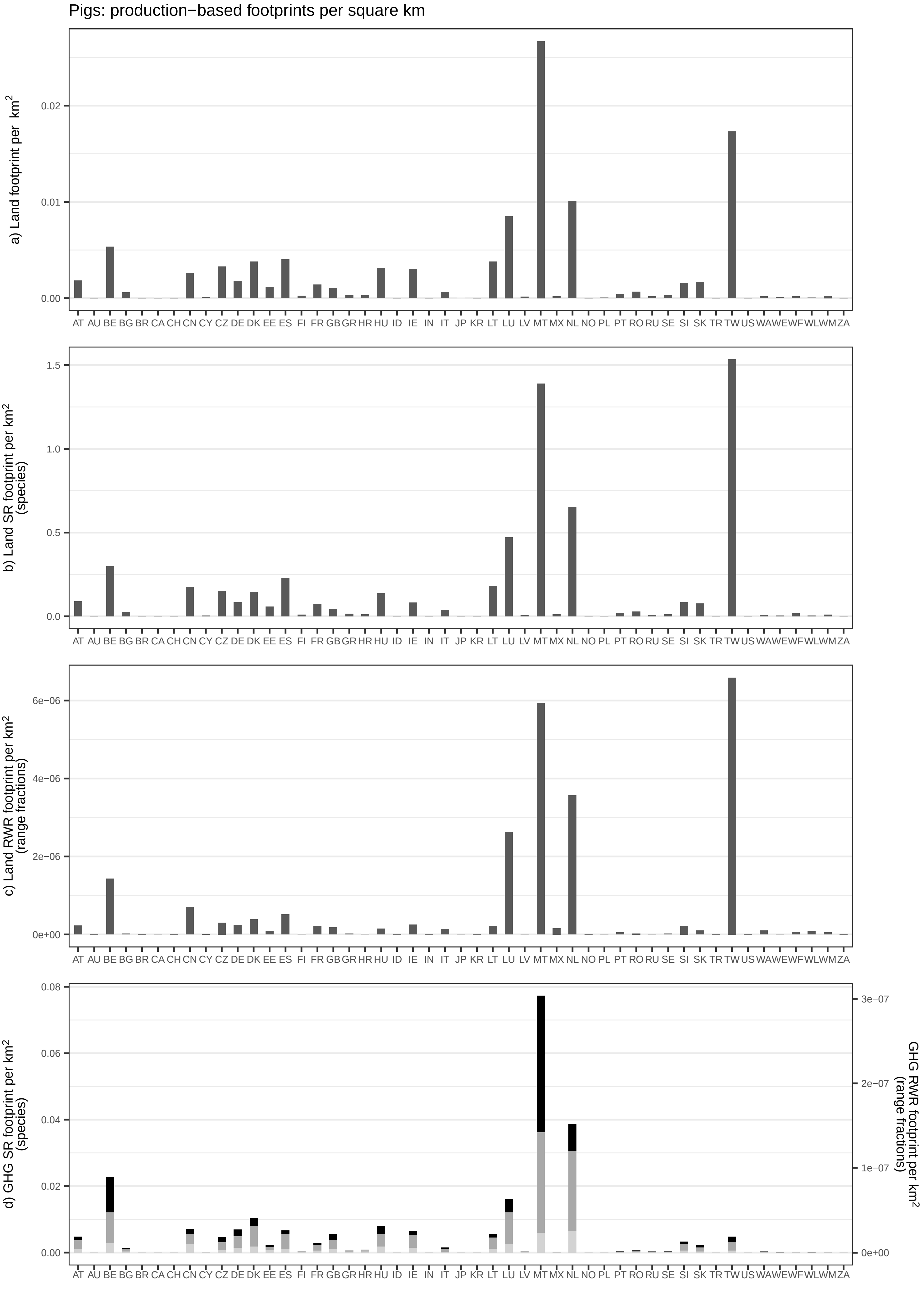
Pigs: production−based footprints per square km
a) Land footprint per km²
0.00
0.01
0.02
b) Land SR footprint per km² (species)
0.0
0.5
1.0
1.5
c) Land RWR footprint per km² (range fractions)
0e+00
2e−06
4e−06
6e−06
d) GHG SR footprint per km² (species)
0.00
0.02
0.04
0.06
0.08
GHG RWR footprint per km² (range fractions)
0e+00
1e−07
2e−07
3e−07
AT AU BE BG BR CA CH CN CY CZ DE DK EE ES FI FR GB GR HR HU ID IE IN IT JP KR LT LU LV MT MX NL NO PL PT RO RU SE SI SK TR TW US WA WE WF WL WM ZA

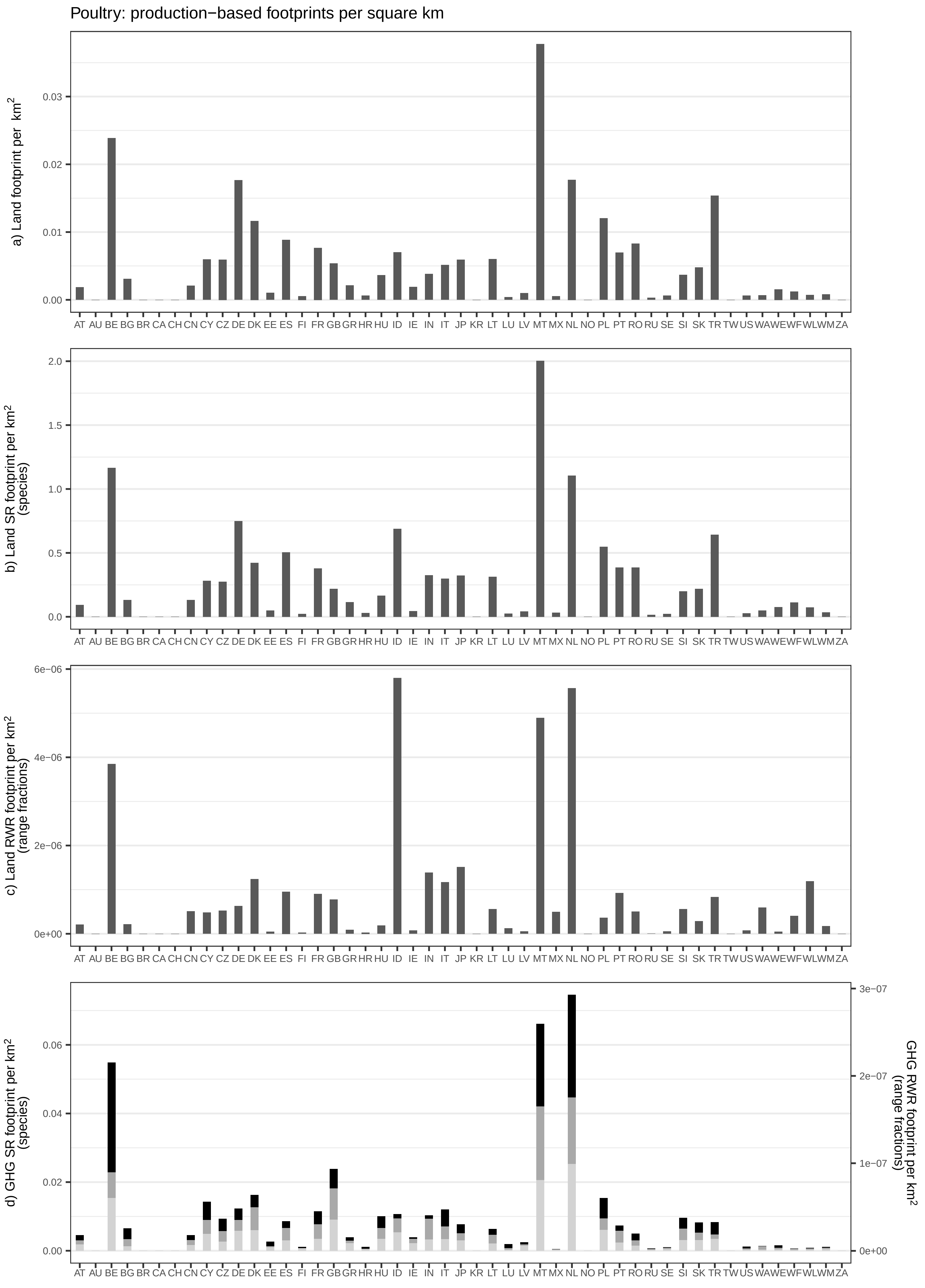
Poultry: production−based footprints per square km
a) Land footprint per km²
0.00
0.01
0.02
0.03
b) Land SR footprint per km² (species)
0.0
0.5
1.0
1.5
2.0
c) Land RWR footprint per km² (range fractions)
0e+00
2e−06
4e−06
6e−06
d) GHG SR footprint per km² (species)
0.00
0.02
0.04
0.06
GHG RWR footprint per km² (range fractions)
0e+00
1e−07
2e−07
3e−07
AT AU BE BG BR CA CH CN CY CZ DE DK EE ES FI FR GB GR HR HU ID IE IN IT JP KR LT LU LV MT MX NL NO PL PT RO RU SE SI SK TR TW US WA WE WF WL WM ZA

Meat animals nec: production−based footprints per square km

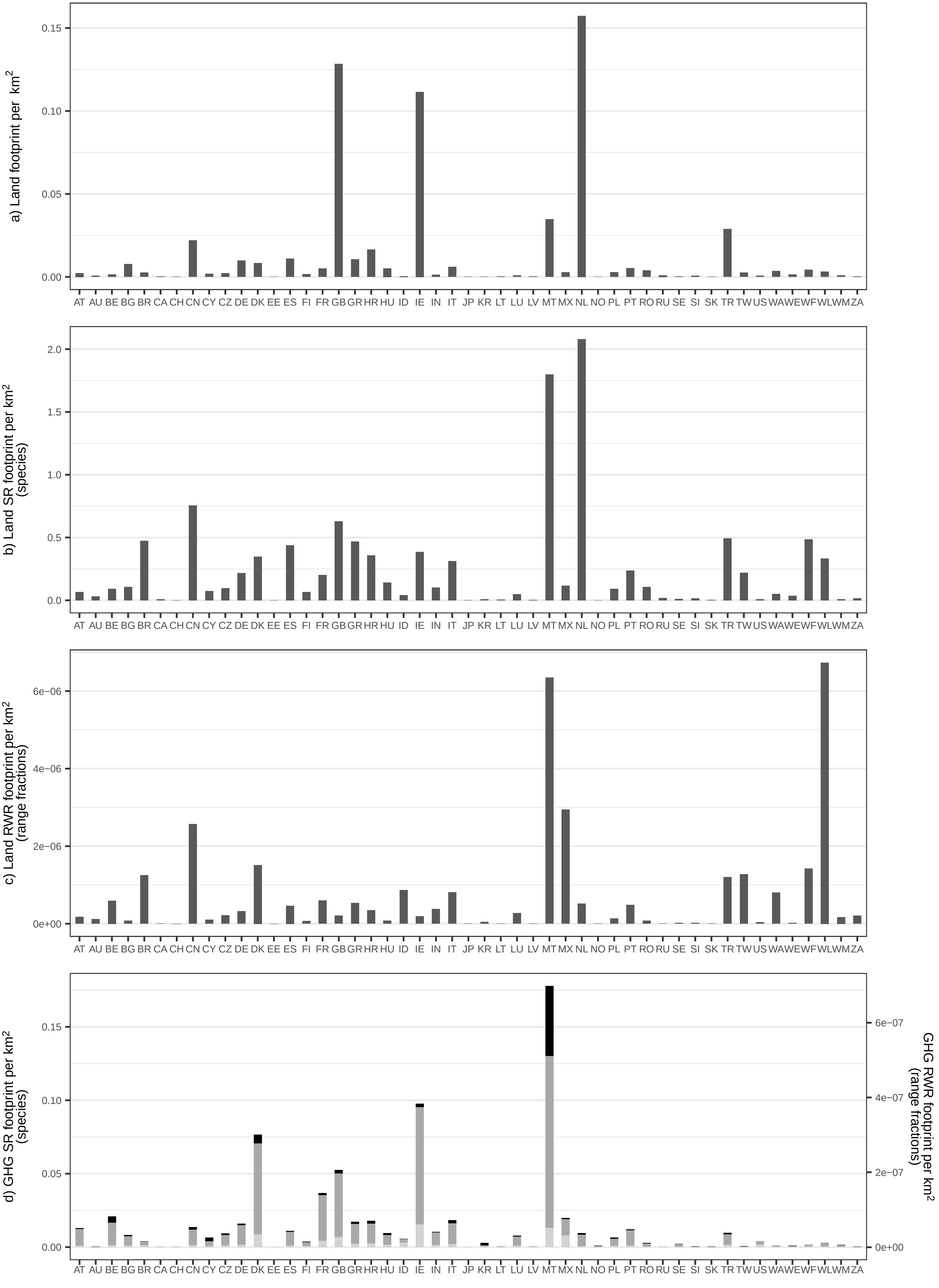

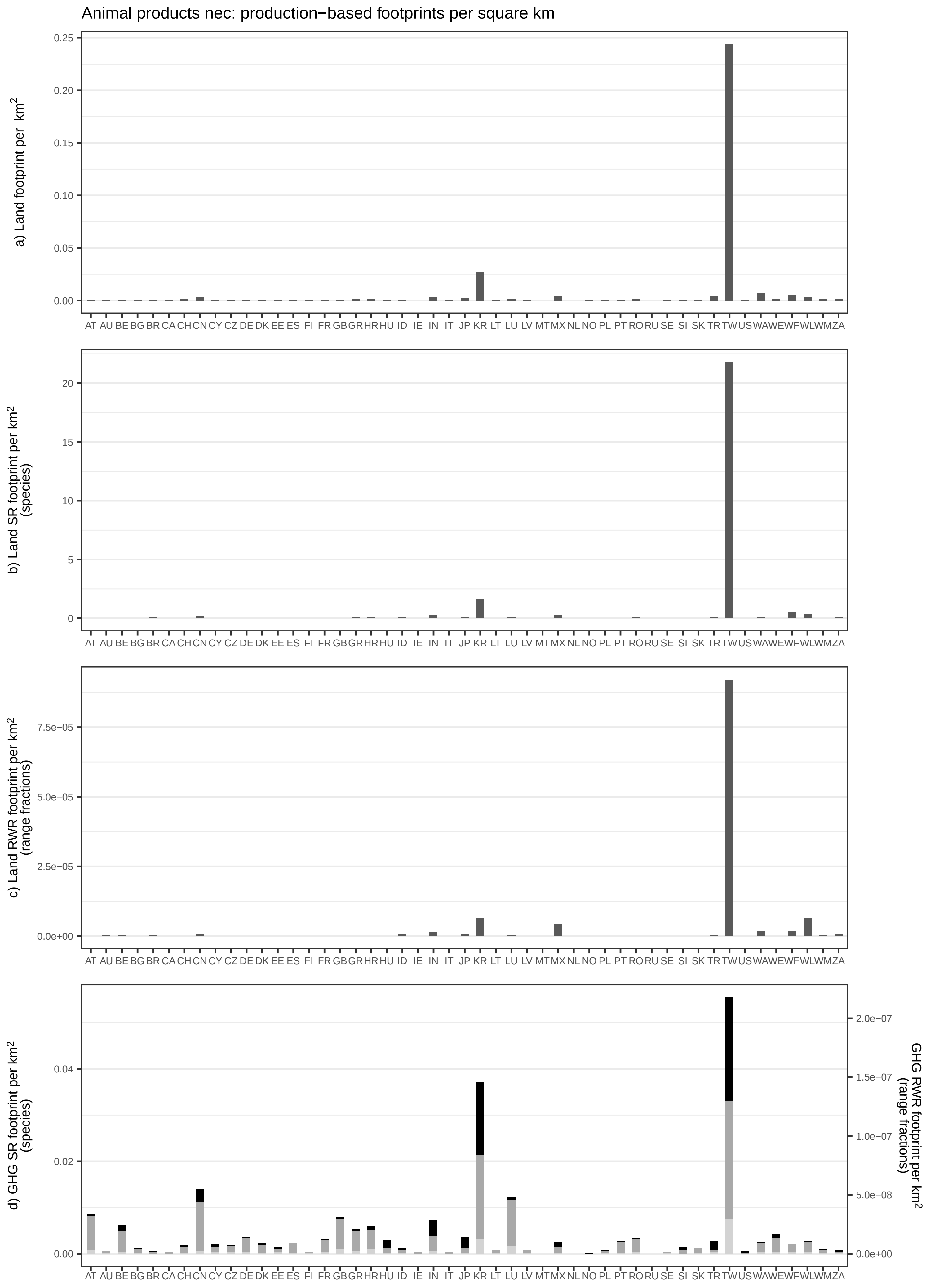
Animal products nec: production−based footprints per square km
a) Land footprint per km²
b) Land SR footprint per km² (species)
c) Land RWR footprint per km² (range fractions)
d) GHG SR footprint per km² (species)
GHG RWR footprint per km² (range fractions)
AT AU BE BG BR CA CH CN CY CZ DE DK EE ES FI FR GB GR HR HU ID IE IN IT JP KR LT LU LV MT MX NL NO PL PT RO RU SE SI SK TR TW US WA WE WF WL WM ZA

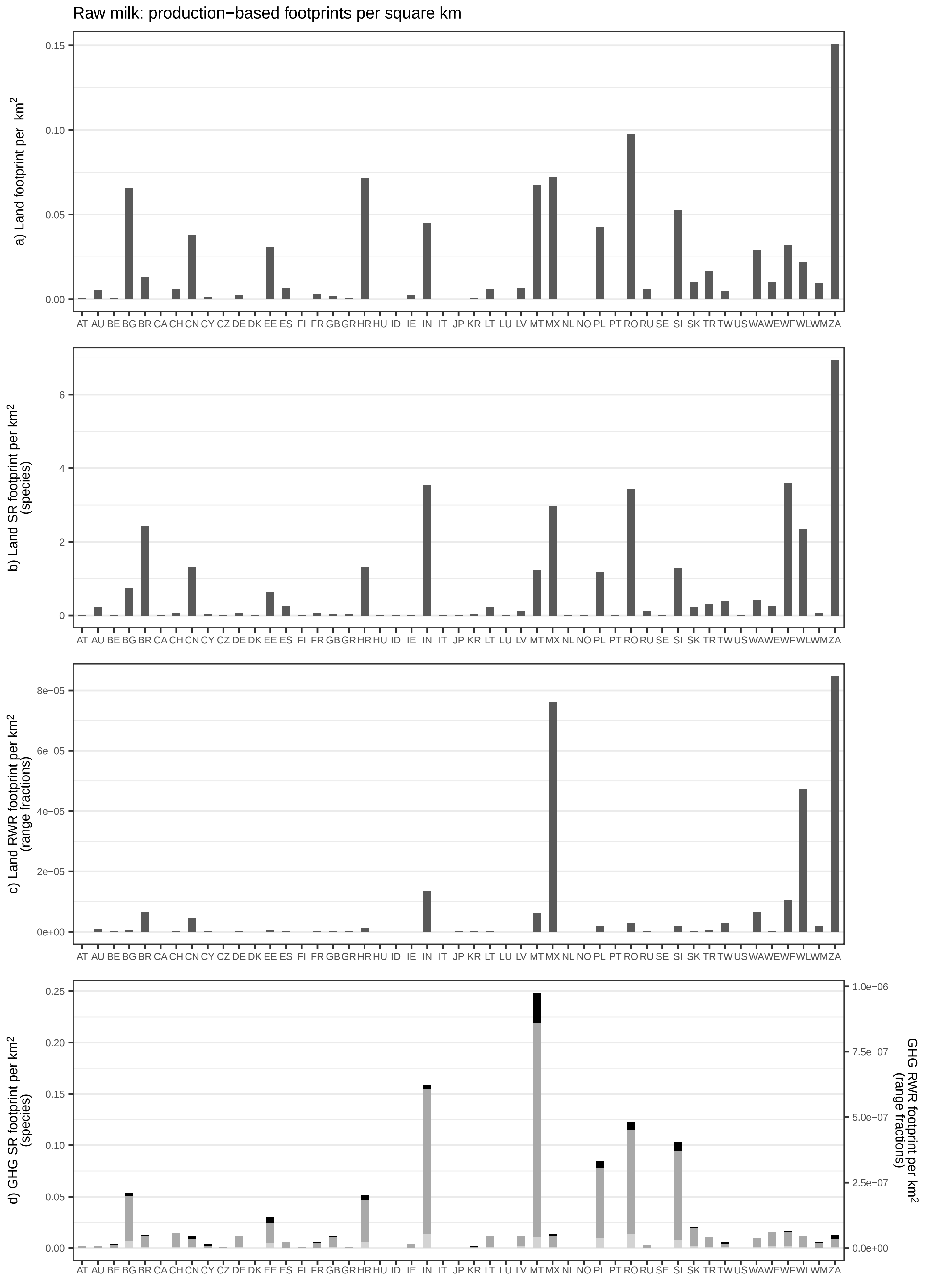

Raw milk: production−based footprints per square km
a) Land footprint per km²
b) Land SR footprint per km² (species)
c) Land RWR footprint per km² (range fractions)
d) GHG SR footprint per km² (species)
GHG RWR footprint per km² (range fractions)
AT AU BE BG BR CA CH CN CY CZ DE DK EE ES FI FR GB GR HR HU ID IE IN IT JP KR LT LU LV MT MX NL NO PL PT RO RU SE SI SK TR TW US WA WE WF WL WM ZA

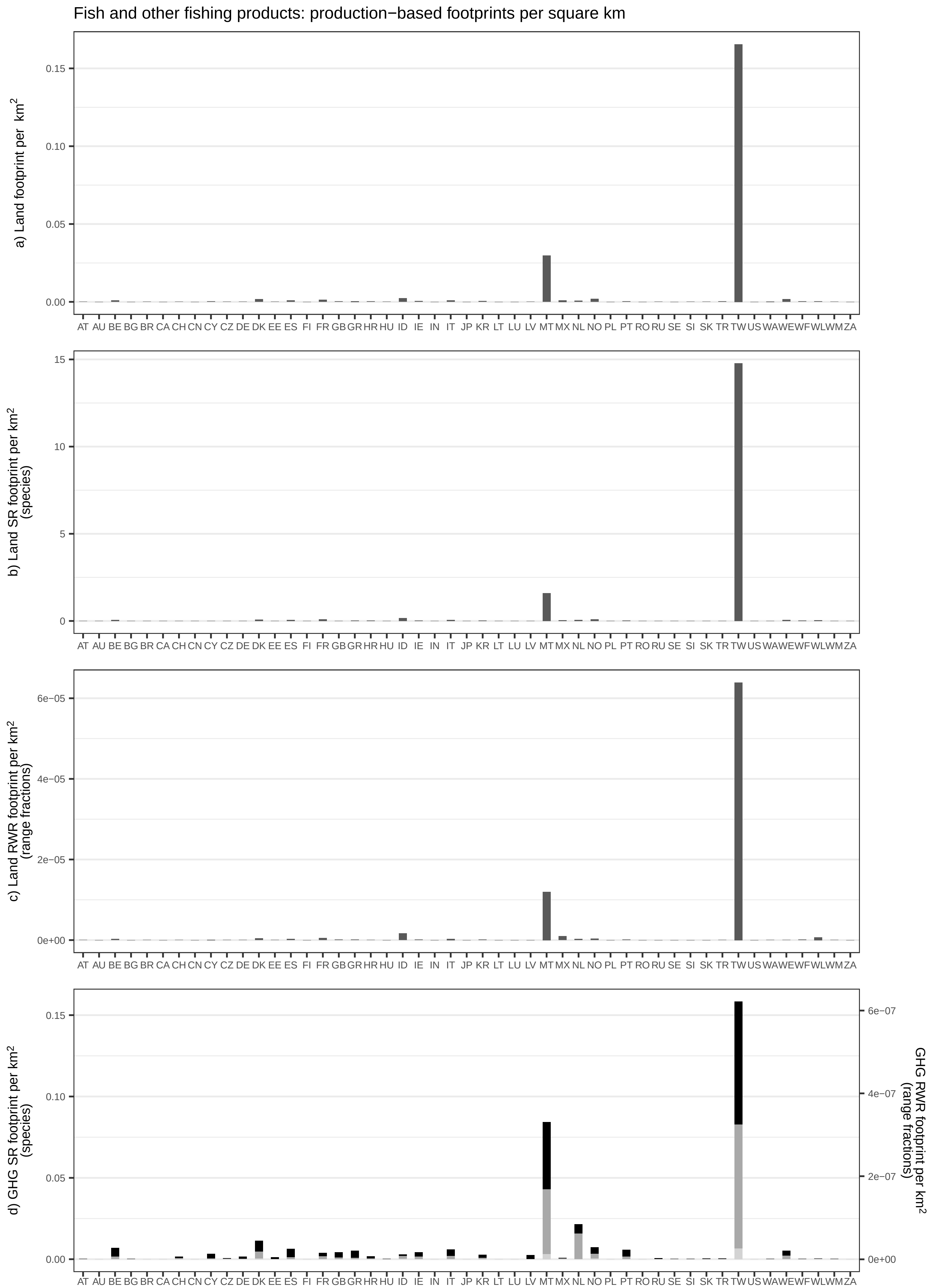
Fish and other fishing products: production−based footprints per square km
a) Land footprint per km²
0.00
0.05
0.10
0.15
b) Land SR footprint per km² (species)
0
5
10
15
c) Land RWR footprint per km² (range fractions)
0e+00
2e−05
4e−05
6e−05
d) GHG SR footprint per km² (species)
0.00
0.05
0.10
0.15
GHG RWR footprint per km² (range fractions)
0e+00
2e−07
4e−07
6e−07
AT AU BE BG BR CA CH CN CY CZ DE DK EE ES FI FR GB GR HR HU ID IE IN IT JP KR LT LU LV MT MX NL NO PL PT RO RU SE SI SK TR TW US WA WE WF WL WM ZA

Chemical and fertiliser minerals: production−based footprints per square km

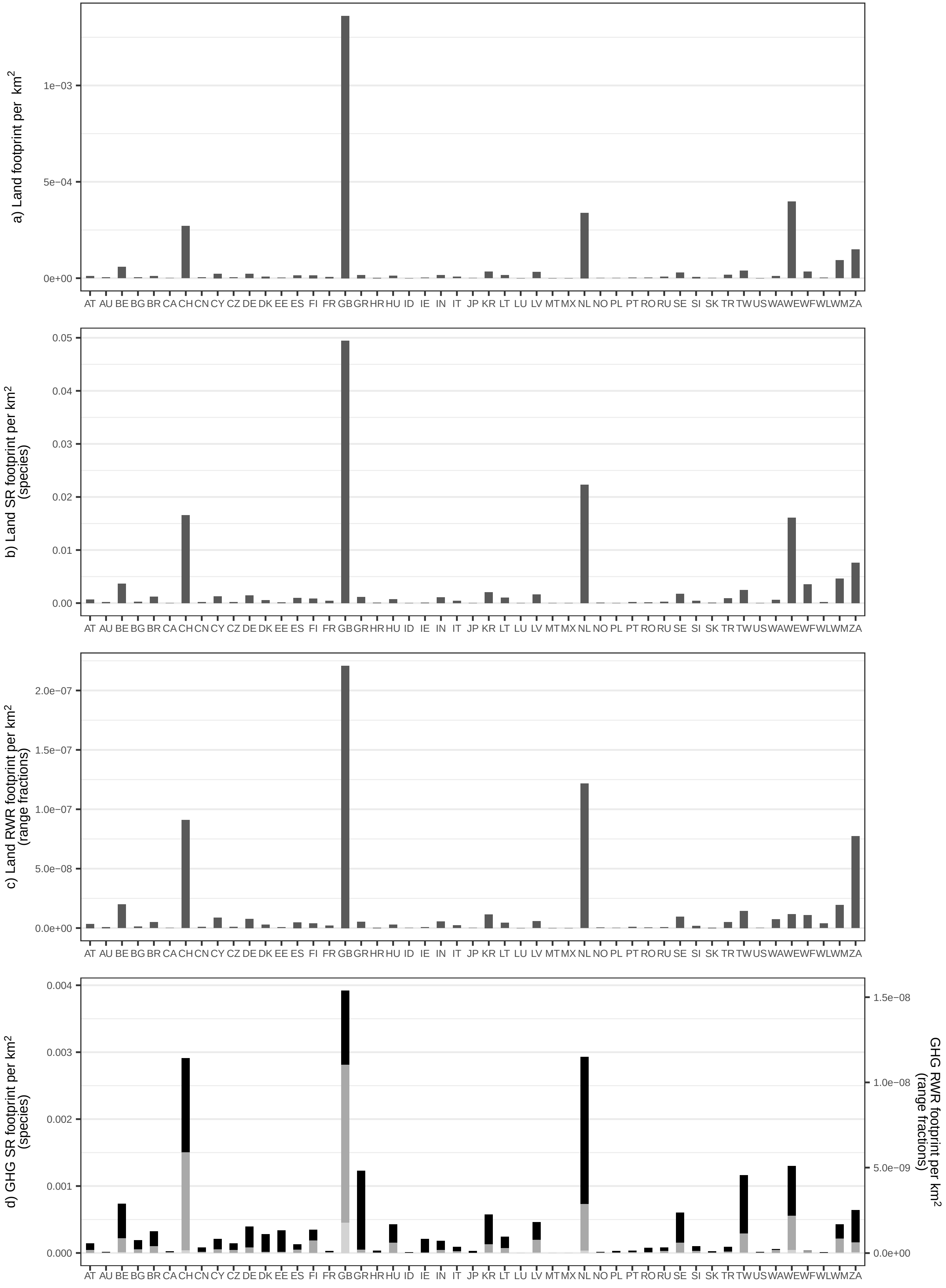

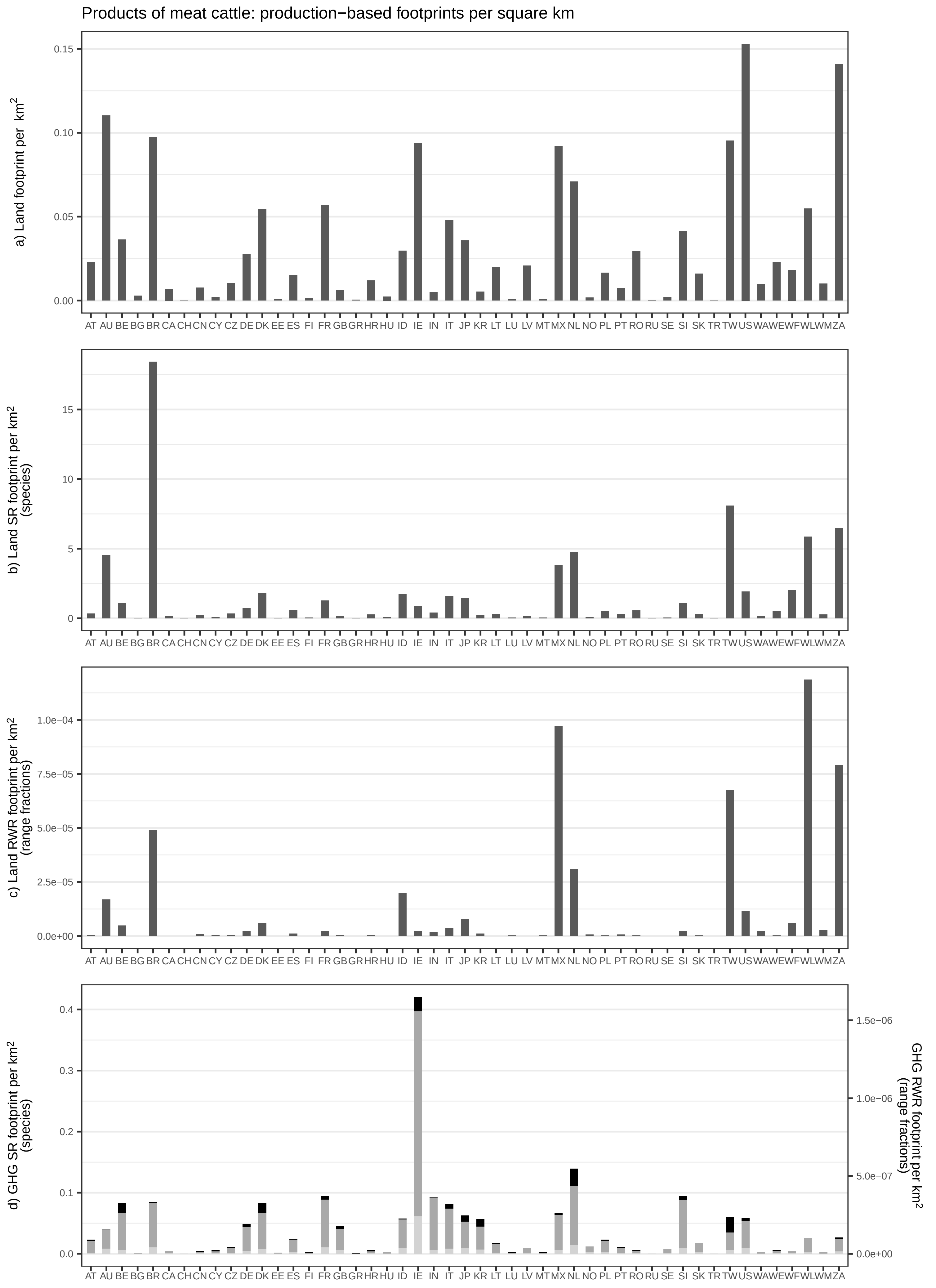

Products of meat cattle: production−based footprints per square km
a) Land footprint per km²
b) Land SR footprint per km² (species)
c) Land RWR footprint per km² (range fractions)
d) GHG SR footprint per km² (species)
GHG RWR footprint per km² (range fractions)
AT AU BE BG BR CA CH CN CY CZ DE DK EE ES FI FR GB GR HR HU ID IE IN IT JP KR LT LU LV MT MX NL NO PL PT RO RU SE SI SK TR TW US WA WE WF WL WM ZA

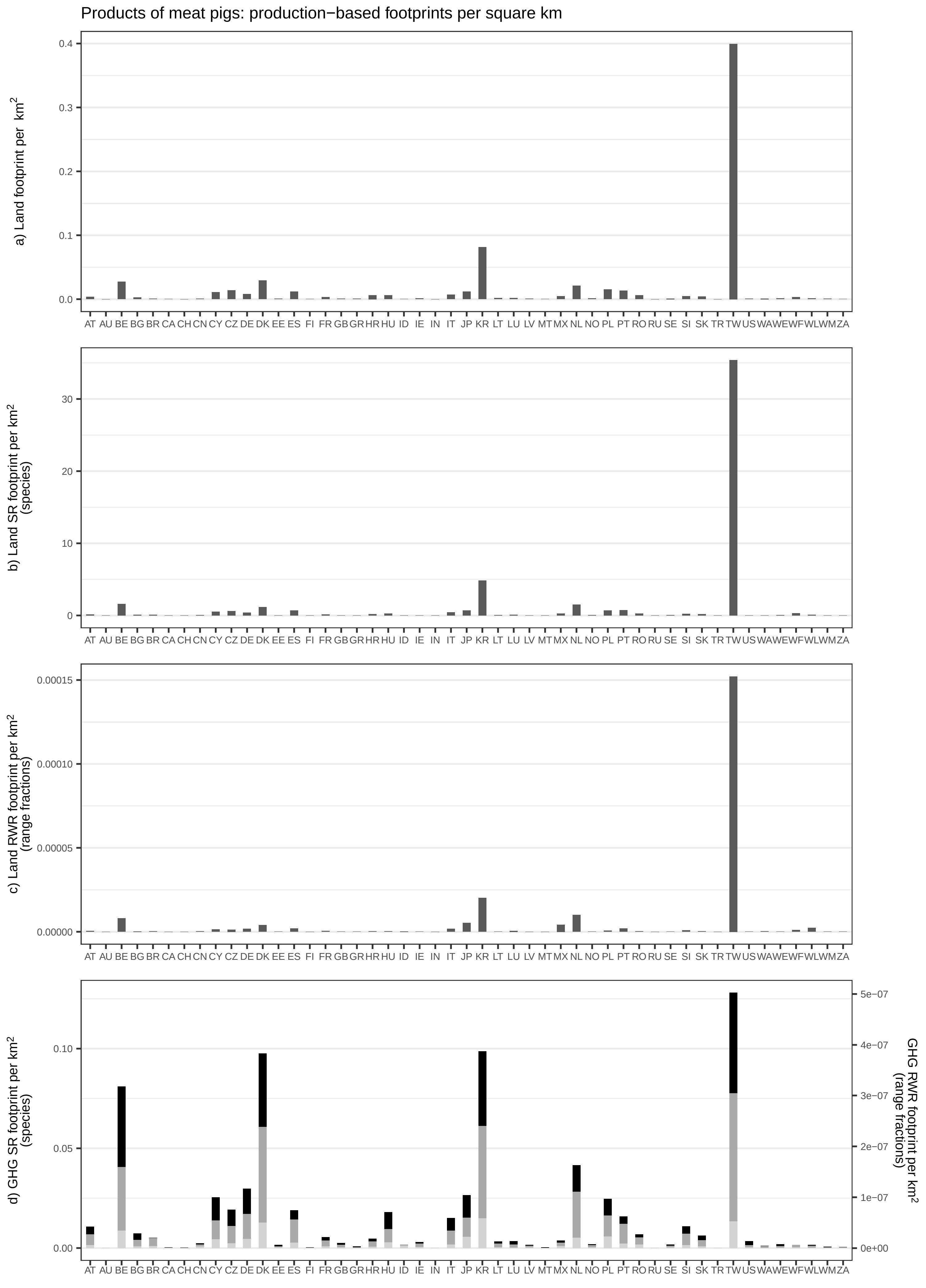

Products of meat pigs: production−based footprints per square km
a) Land footprint per km²
b) Land SR footprint per km² (species)
c) Land RWR footprint per km² (range fractions)
d) GHG SR footprint per km² (species)
GHG RWR footprint per km² (range fractions)
AT AU BE BG BR CA CH CN CY CZ DE DK EE ES FI FR GB GR HR HU ID IE IN IT JP KR LT LU LV MT MX NL NO PL PT RO RU SE SI SK TR TW US WA WE WF WL WM ZA

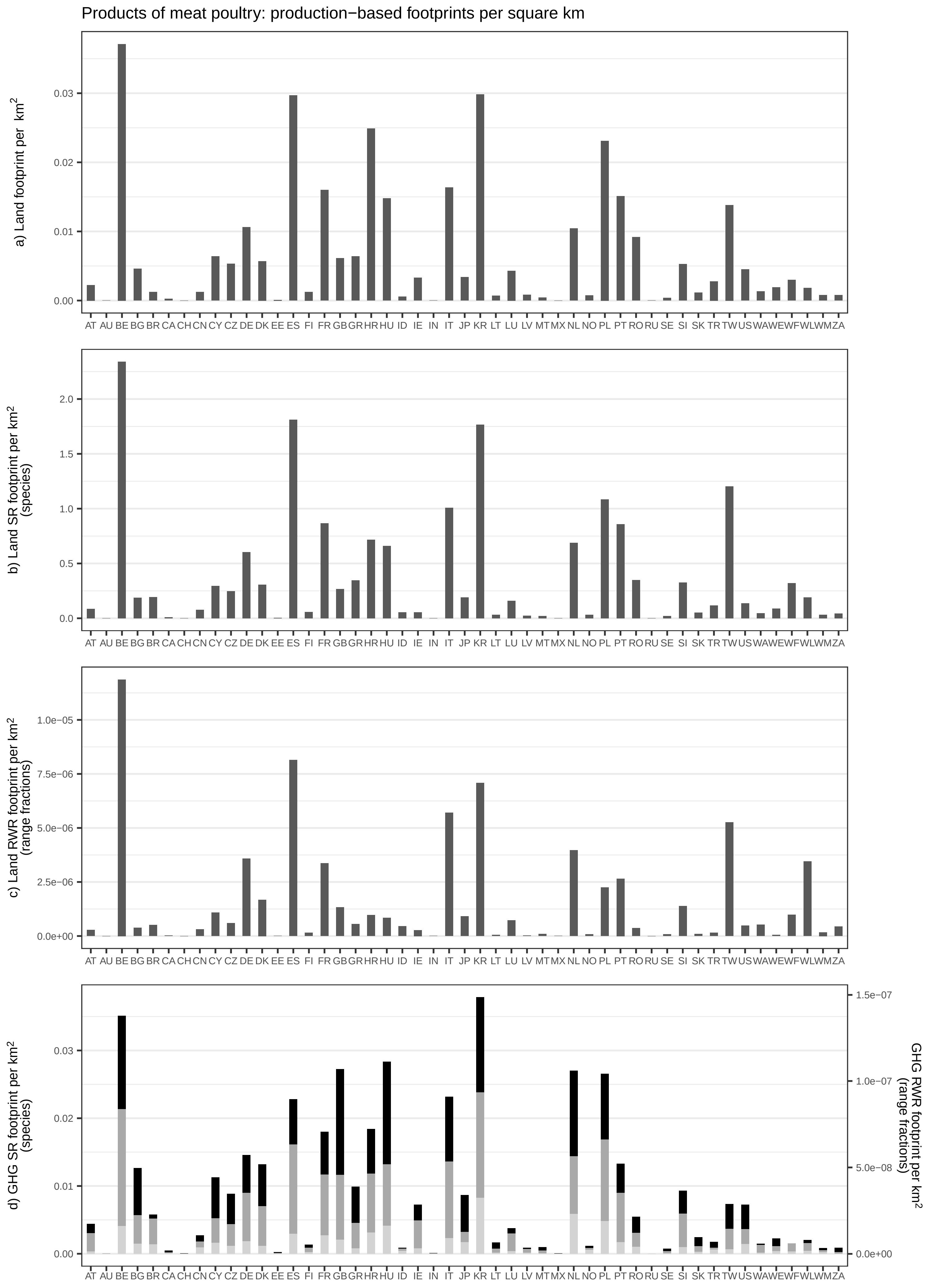

Products of meat poultry: production−based footprints per square km
a) Land footprint per km²
b) Land SR footprint per km² (species)
c) Land RWR footprint per km² (range fractions)
d) GHG SR footprint per km² (species)
GHG RWR footprint per km² (range fractions)
AT AU BE BG BR CA CH CN CY CZ DE DK EE ES FI FR GB GR HR HU ID IE IN IT JP KR LT LU LV MT MX NL NO PL PT RO RU SE SI SK TR TW US WA WE WF WL WM ZA

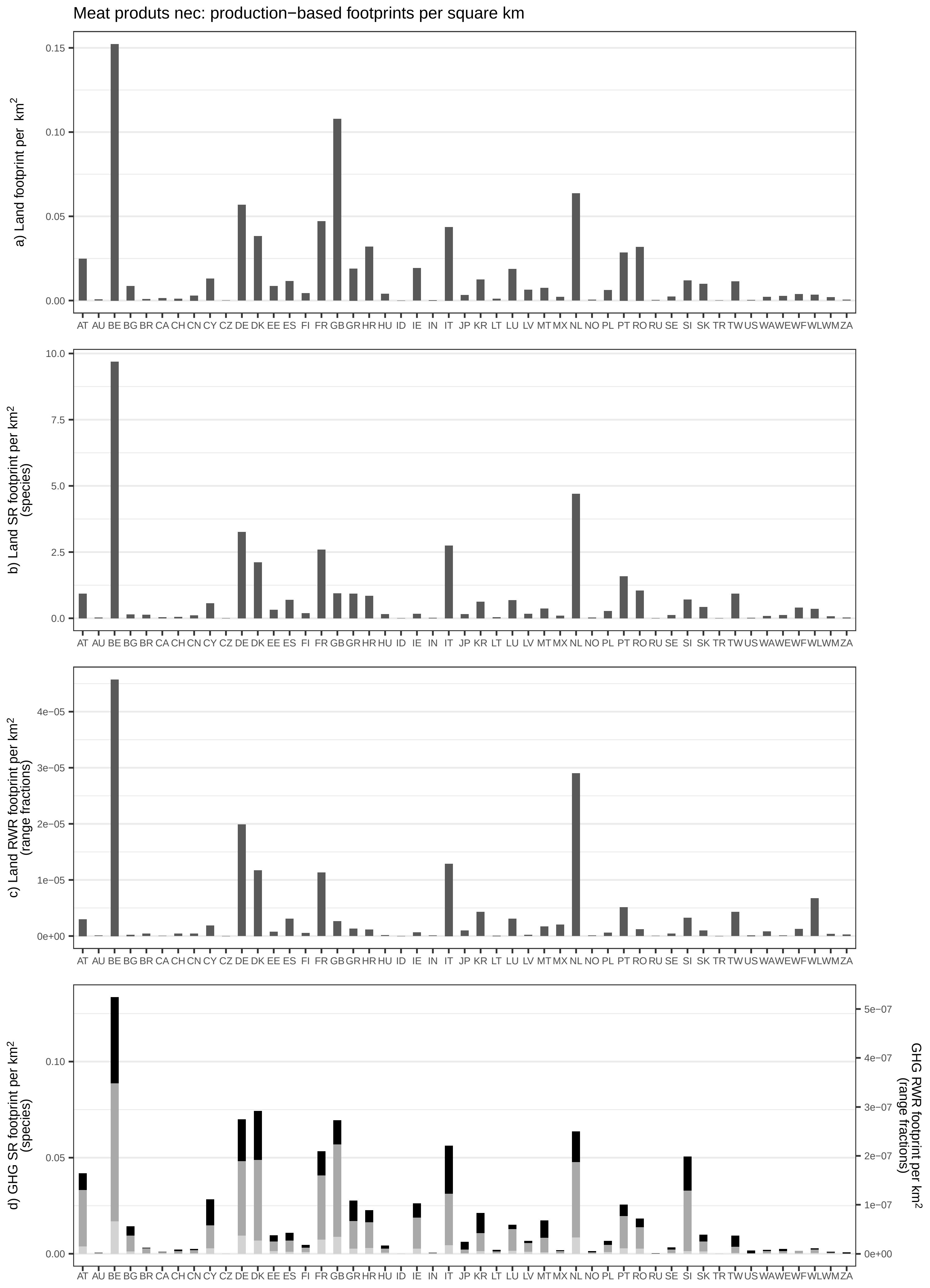

Meat produts nec: production−based footprints per square km
a) Land footprint per km²
b) Land SR footprint per km² (species)
c) Land RWR footprint per km² (range fractions)
d) GHG SR footprint per km² (species)
GHG RWR footprint per km² (range fractions)
AT AU BE BG BR CA CH CN CY CZ DE DK EE ES FI FR GB GR HR HU ID IE IN IT JP KR LT LU LV MT MX NL NO PL PT RO RU SE SI SK TR TW US WA WE WF WL WM ZA

Products of vegetable oils and fats: production−based footprints per square km

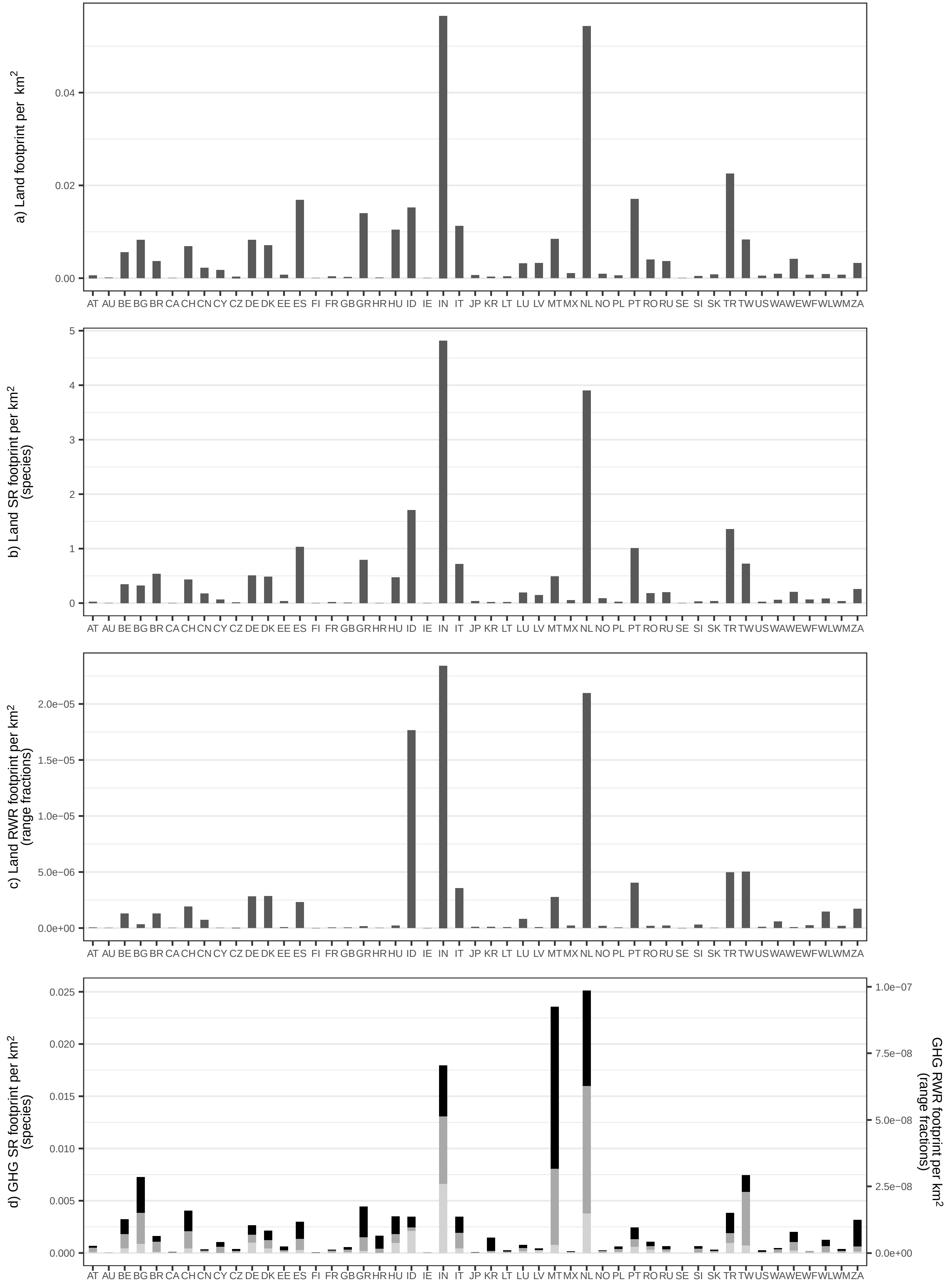

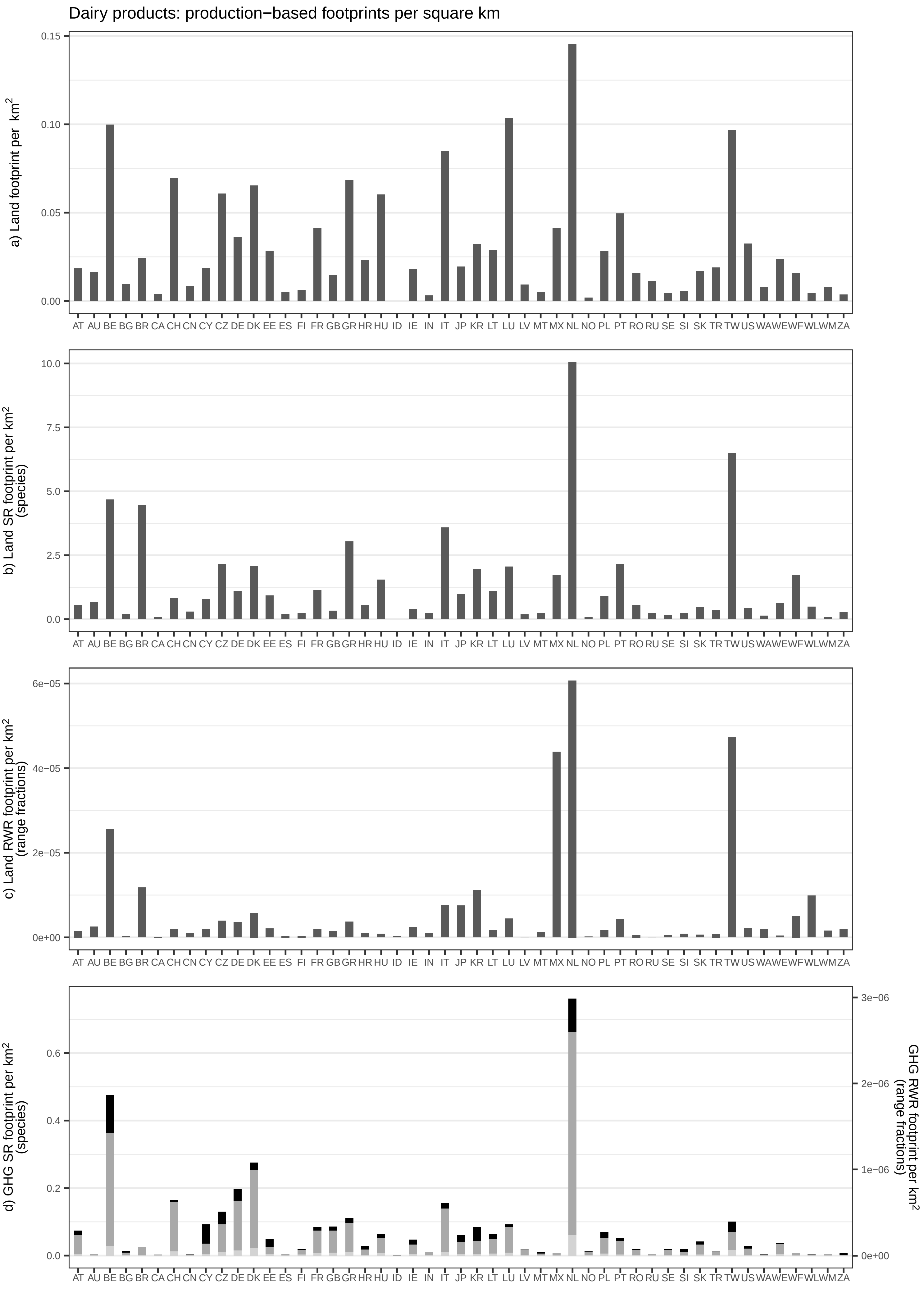

Dairy products: production−based footprints per square km
a) Land footprint per km²
b) Land SR footprint per km² (species)
c) Land RWR footprint per km² (range fractions)
d) GHG SR footprint per km² (species)
GHG RWR footprint per km² (range fractions)
AT AU BE BG BR CA CH CN CY CZ DE DK EE ES FI FR GB GR HR HU ID IE IN IT JP KR LT LU LV MT MX NL NO PL PT RO RU SE SI SK TR TW US WA WE WF WL WM ZA

Processed rice: production−based footprints per square km

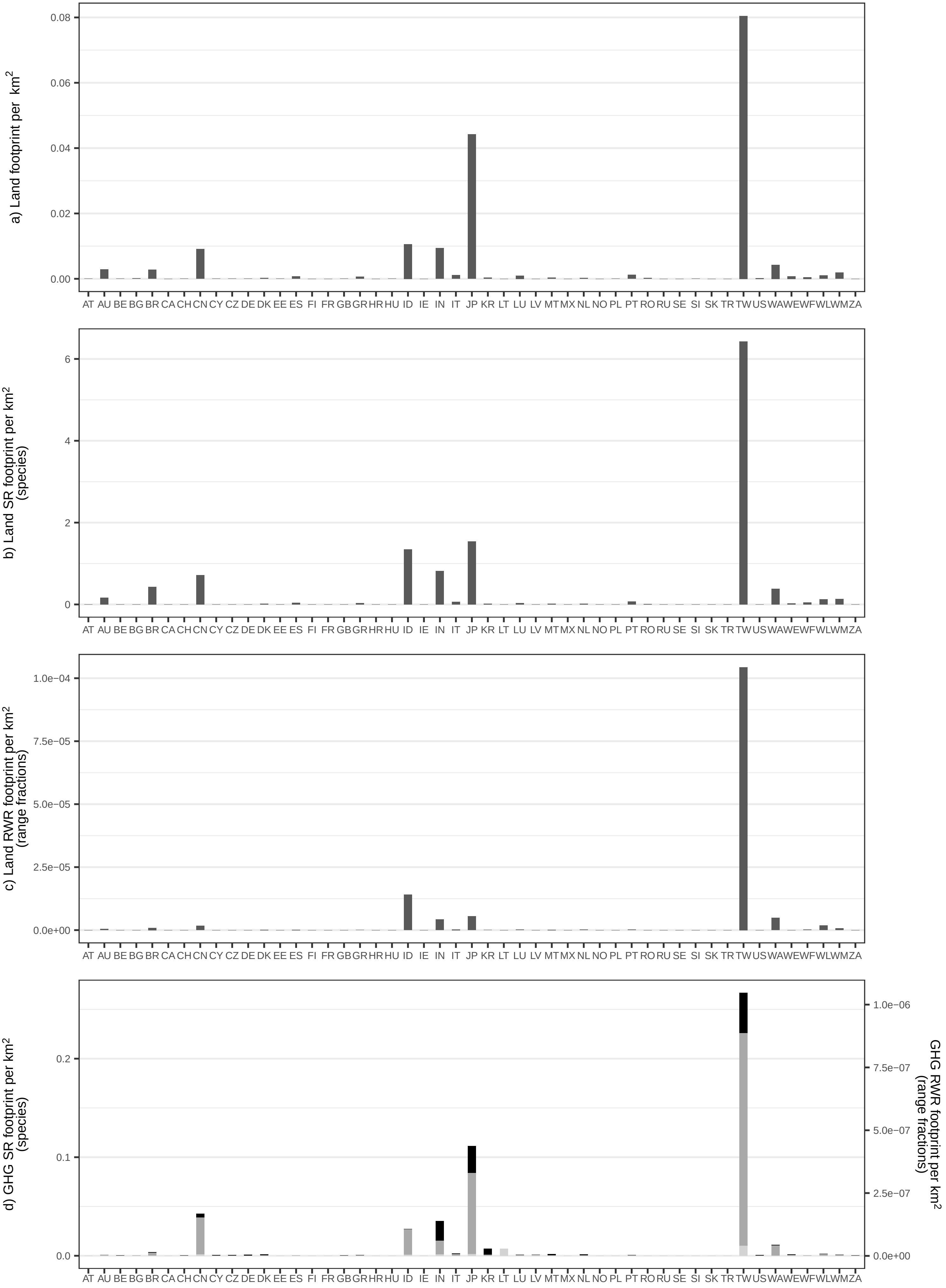

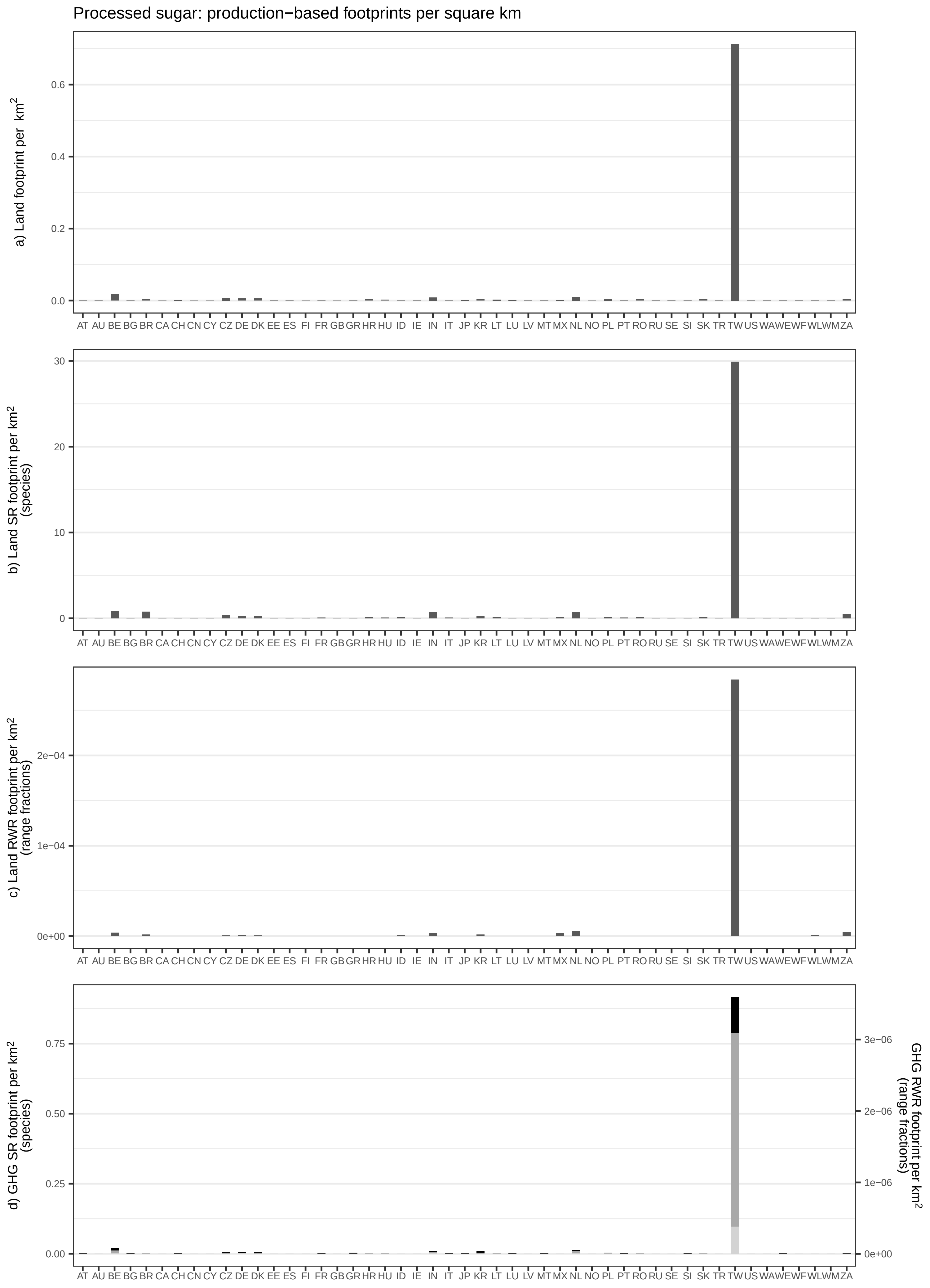
Processed sugar: production−based footprints per square km
a) Land footprint per km²
b) Land SR footprint per km² (species)
c) Land RWR footprint per km² (range fractions)
d) GHG SR footprint per km² (species)
GHG RWR footprint per km² (range fractions)
AT AU BE BG BR CA CH CN CY CZ DE DK EE ES FI FR GB GR HR HU ID IE IN IT JP KR LT LU LV MT MX NL NO PL PT RO RU SE SI SK TR TW US WA WE WF WL WM ZA

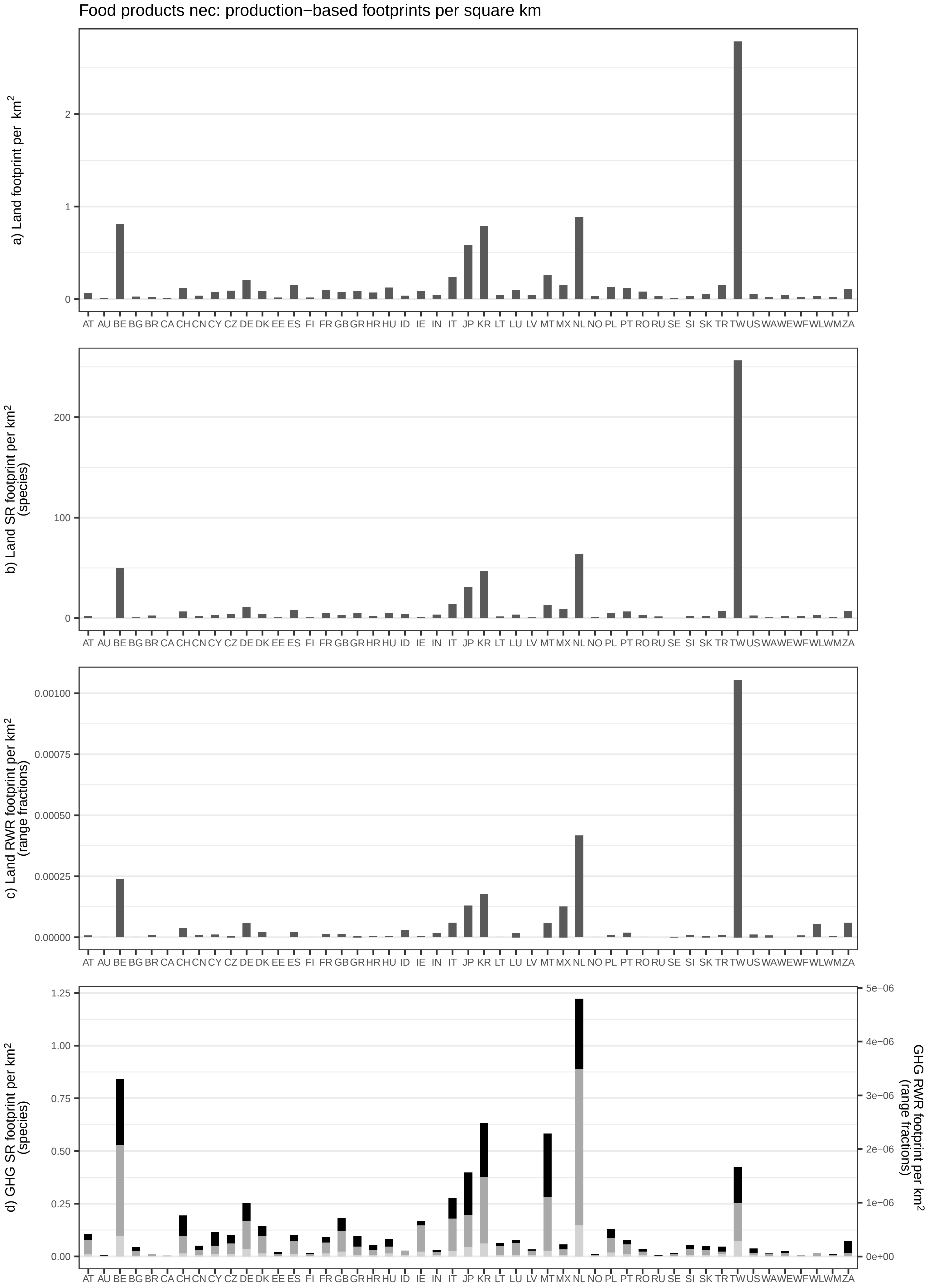

Food products nec: production−based footprints per square km
a) Land footprint per km²
b) Land SR footprint per km² (species)
c) Land RWR footprint per km² (range fractions)
d) GHG SR footprint per km² (species)
GHG RWR footprint per km² (range fractions)
AT AU BE BG BR CA CH CN CY CZ DE DK EE ES FI FR GB GR HR HU ID IE IN IT JP KR LT LU LV MT MX NL NO PL PT RO RU SE SI SK TR TW US WA WE WF WL WM ZA

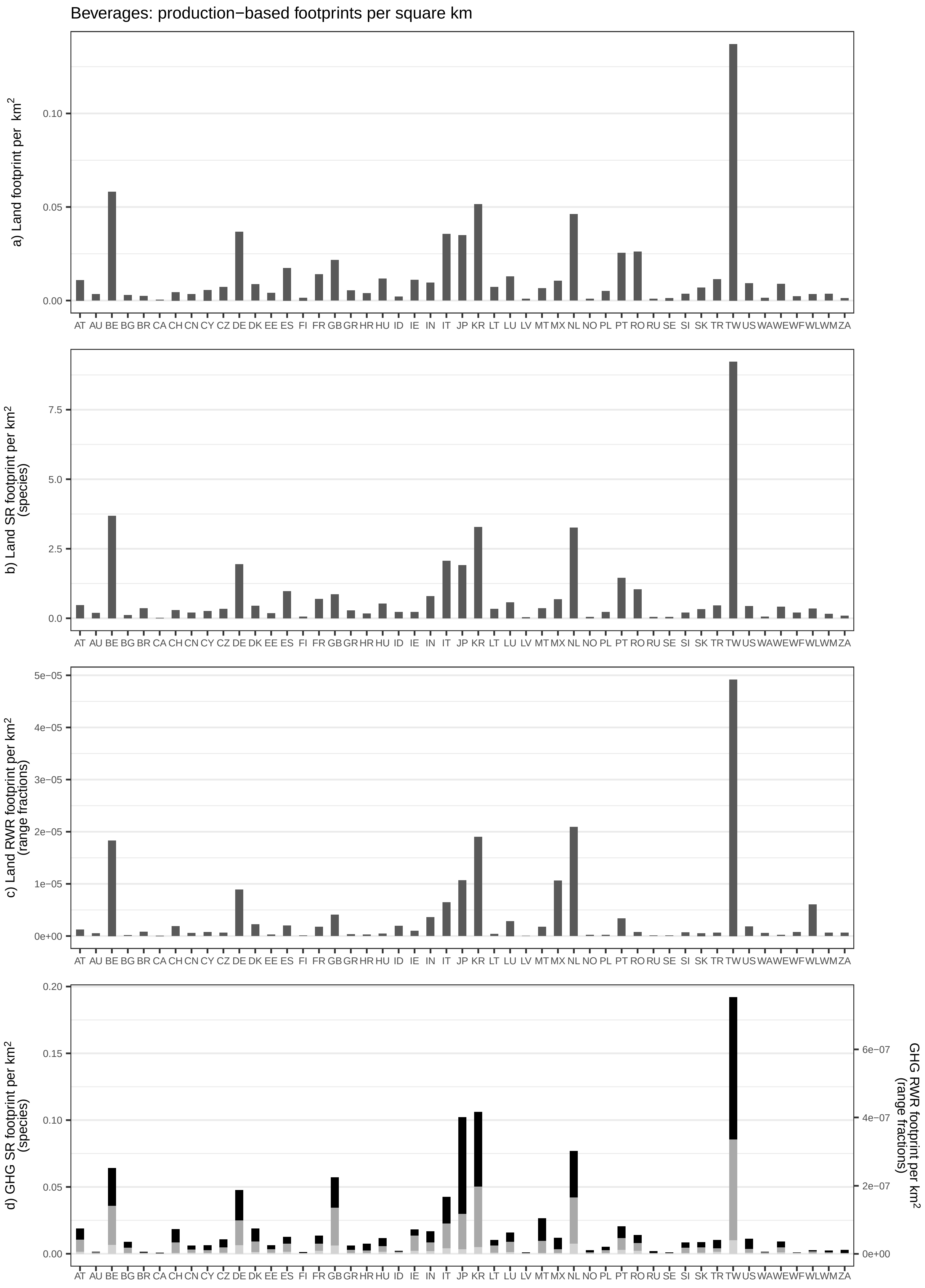
Beverages: production−based footprints per square km
a) Land footprint per km²
b) Land SR footprint per km² (species)
c) Land RWR footprint per km² (range fractions)
d) GHG SR footprint per km² (species)
GHG RWR footprint per km² (range fractions)
AT AU BE BG BR CA CH CN CY CZ DE DK EE ES FI FR GB GR HR HU ID IE IN IT JP KR LT LU LV MT MX NL NO PL PT RO RU SE SI SK TR TW US WA WE WF WL WM ZA

Fish products: production−based footprints per square km

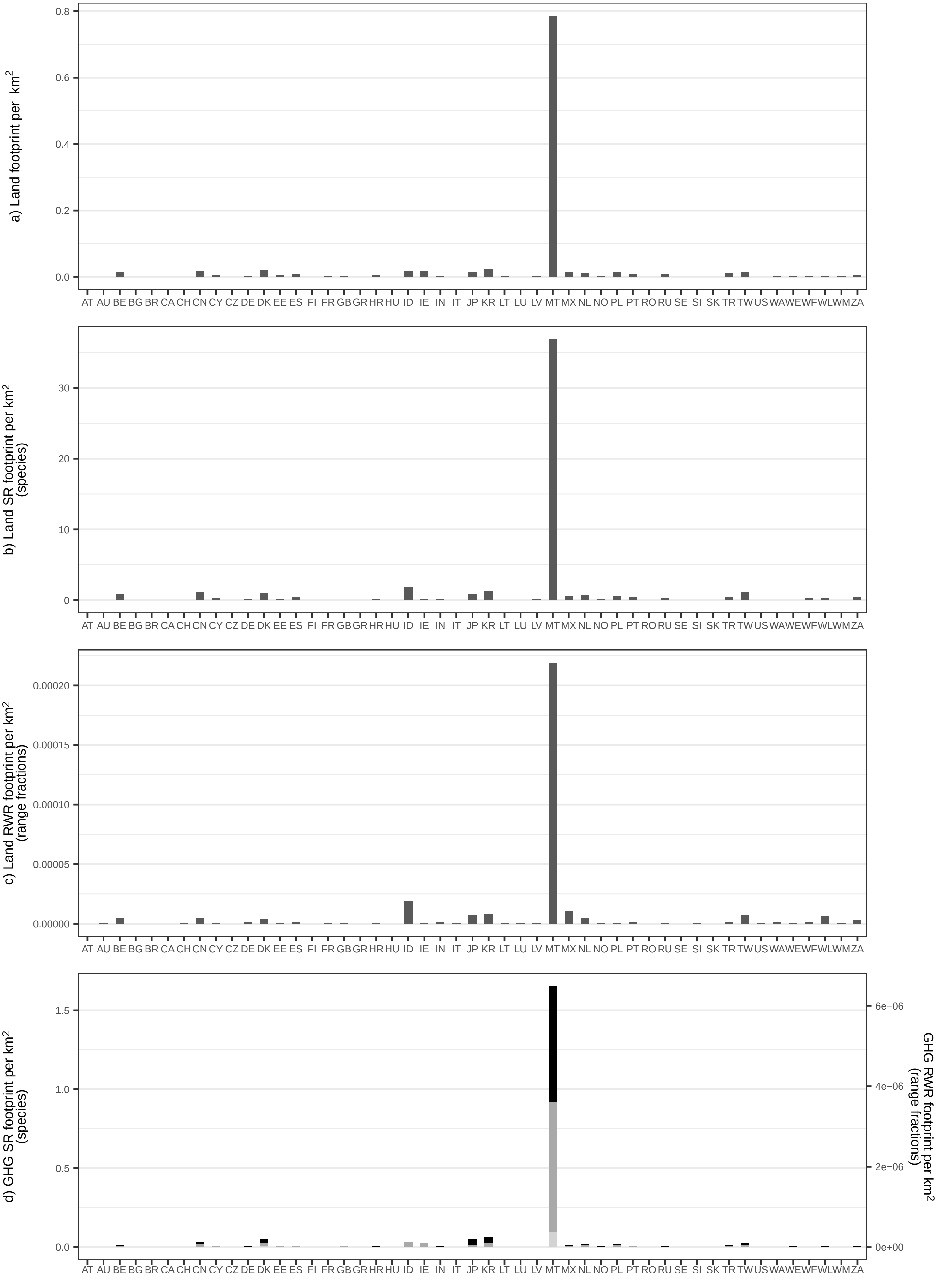

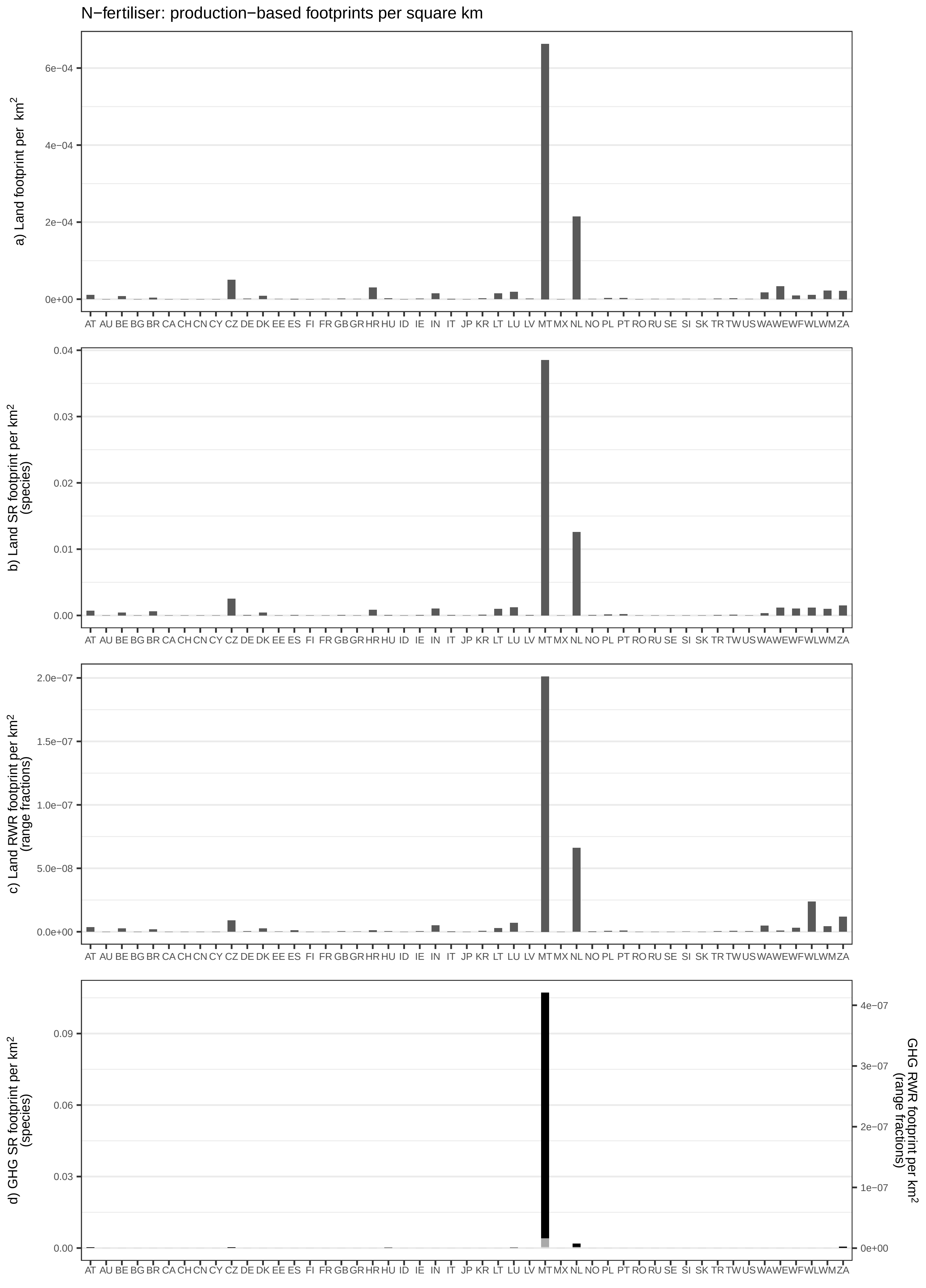
N−fertiliser: production−based footprints per square km
a) Land footprint per km²
6e−04
4e−04
2e−04
0e+00
b) Land SR footprint per km² (species)
0.04
0.03
0.02
0.01
0.00
c) Land RWR footprint per km² (range fractions)
2.0e−07
1.5e−07
1.0e−07
5.0e−08
0.0e+00
d) GHG SR footprint per km² (species)
0.09
0.06
0.03
0.00
GHG RWR footprint per km² (range fractions)
4e−07
3e−07
2e−07
1e−07
0e+00
AT AU BE BG BR CA CH CN CY CZ DE DK EE ES FI FR GB GR HR HU ID IE IN IT JP KR LT LU LV MT MX NL NO PL PT RO RU SE SI SK TR TW US WA WE WF WL WM ZA

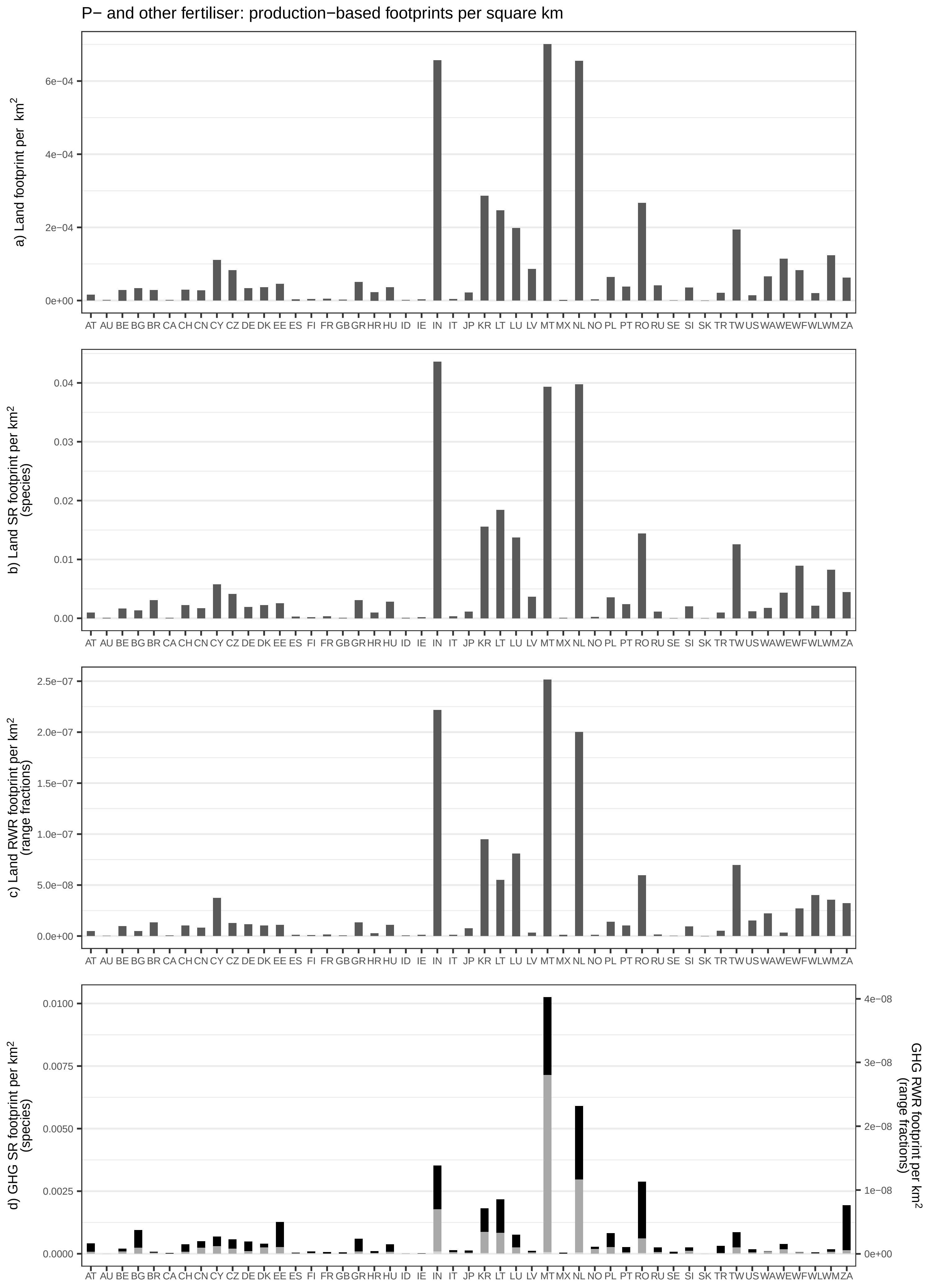
P− and other fertiliser: production−based footprints per square km
a) Land footprint per km²
b) Land SR footprint per km² (species)
c) Land RWR footprint per km² (range fractions)
d) GHG SR footprint per km² (species)
GHG RWR footprint per km² (range fractions)
0e+00
2e−04
4e−04
6e−04
0.00
0.01
0.02
0.03
0.04
0.0e+00
5.0e−08
1.0e−07
1.5e−07
2.0e−07
2.5e−07
0.0000
0.0025
0.0050
0.0075
0.0100
1e−08
2e−08
3e−08
4e−08
AT AU BE BG BR CA CH CN CY CZ DE DK EE ES FI FR GB GR HR HU ID IE IN IT JP KR LT LU LV MT MX NL NO PL PT RO RU SE SI SK TR TW US WA WE WF WL WM ZA

Food waste: incineration: production−based footprints per square km

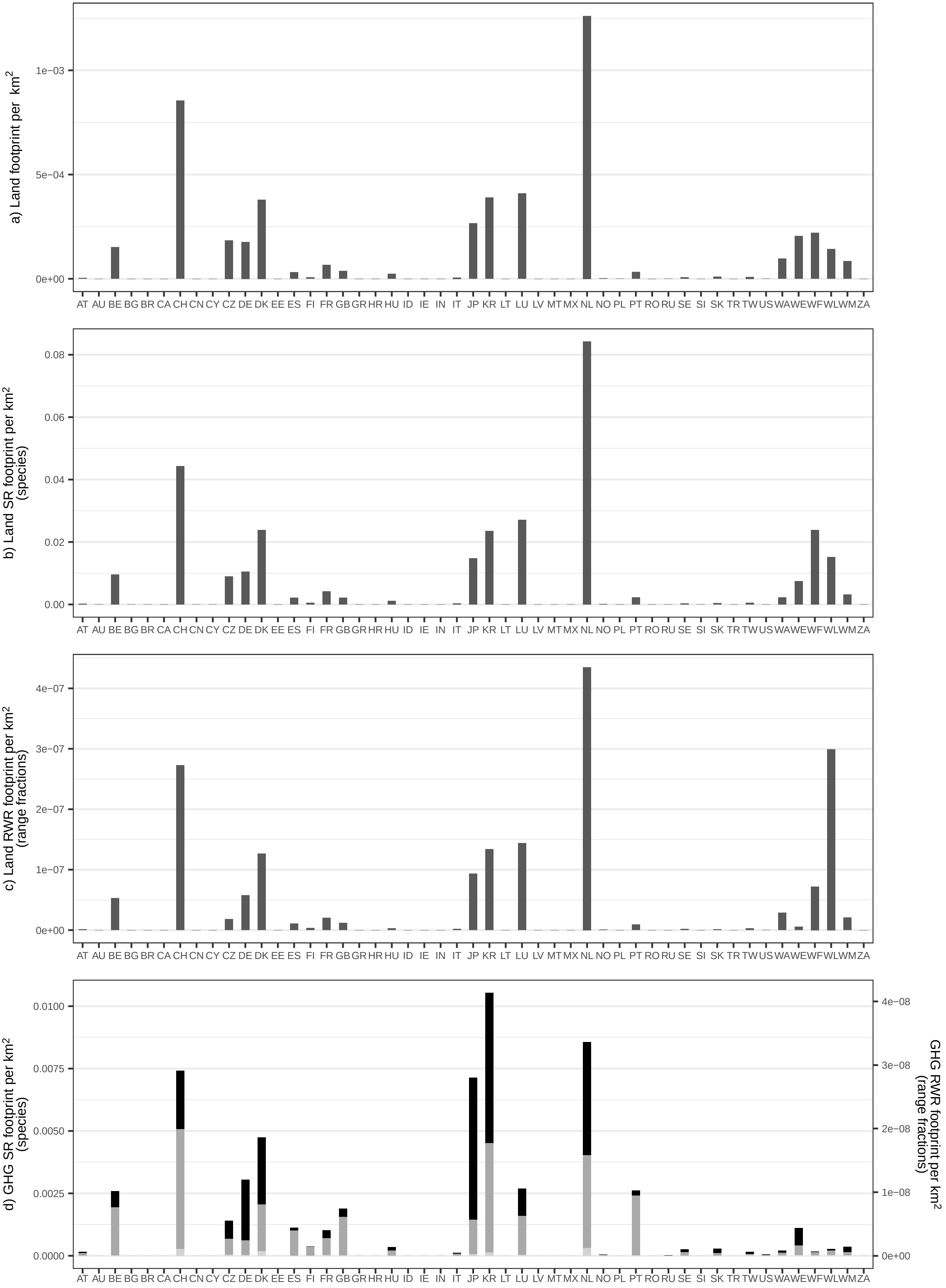

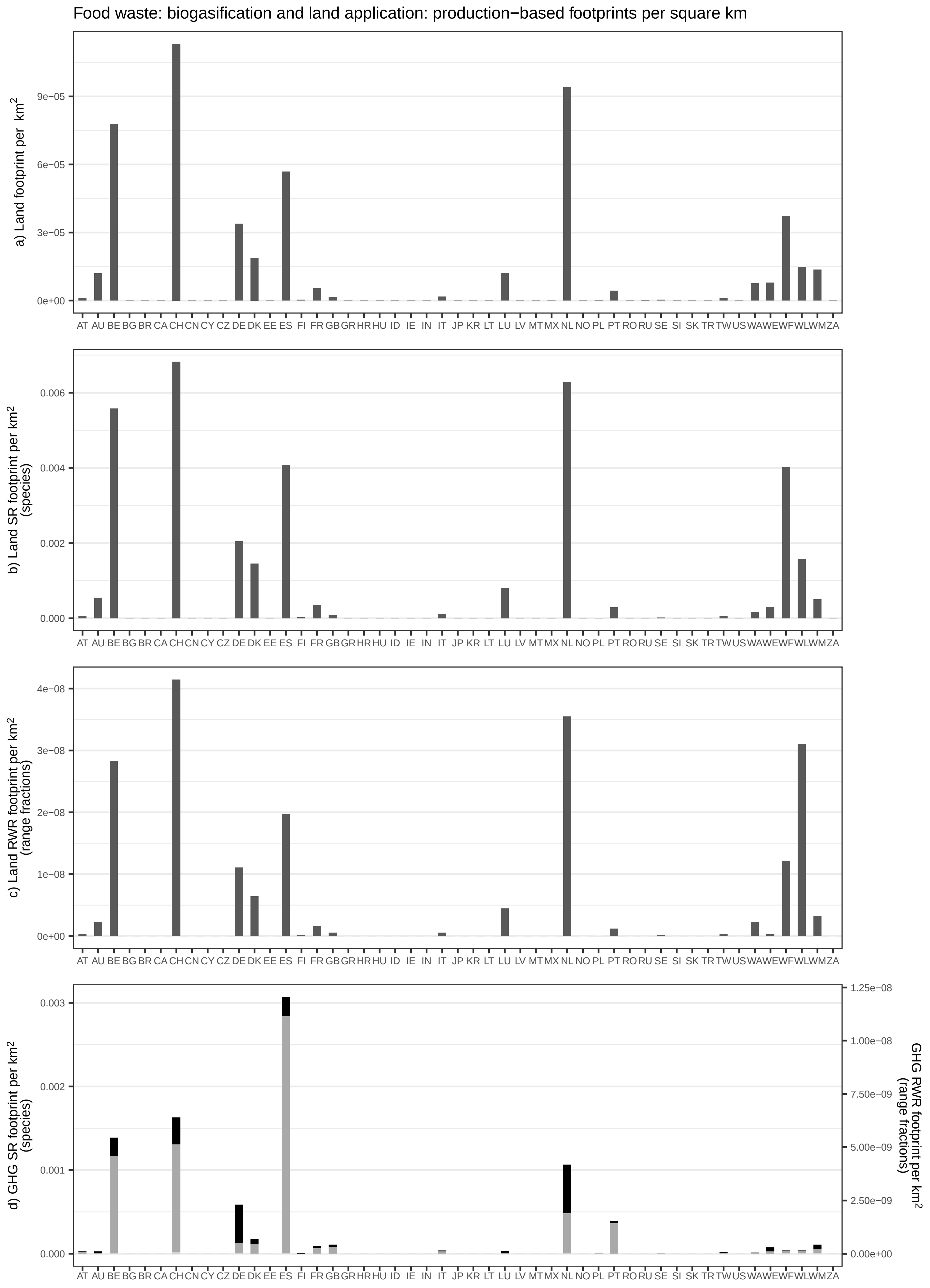
Food waste: biogasification and land application: production−based footprints per square km
a) Land footprint per km²
b) Land SR footprint per km² (species)
c) Land RWR footprint per km² (range fractions)
d) GHG SR footprint per km² (species)
GHG RWR footprint per km² (range fractions)
AT AU BE BG BR CA CH CN CY CZ DE DK EE ES FI FR GB GR HR HU ID IE IN IT JP KR LT LU LV MT MX NL NO PL PT RO RU SE SI SK TR TW US WA WE WF WL WM ZA
0e+00
3e−05
6e−05
9e−05
0.000
0.002
0.004
0.006
1e−08
2e−08
3e−08
4e−08
0.001
0.002
0.003
0.00e+00
2.50e−09
5.00e−09
7.50e−09
1.00e−08
1.25e−08

Food waste: composting and land application: production−based footprints per square km

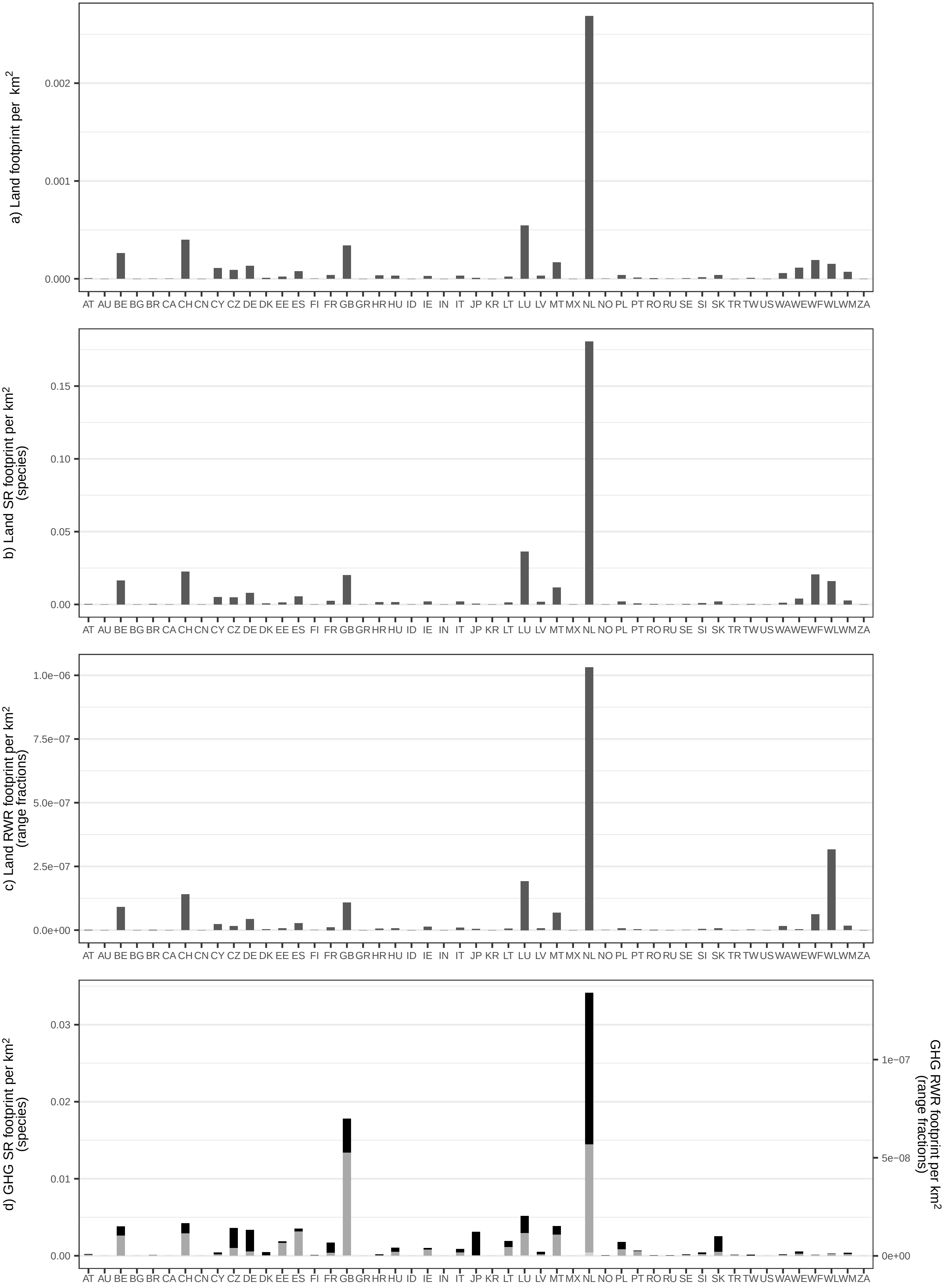

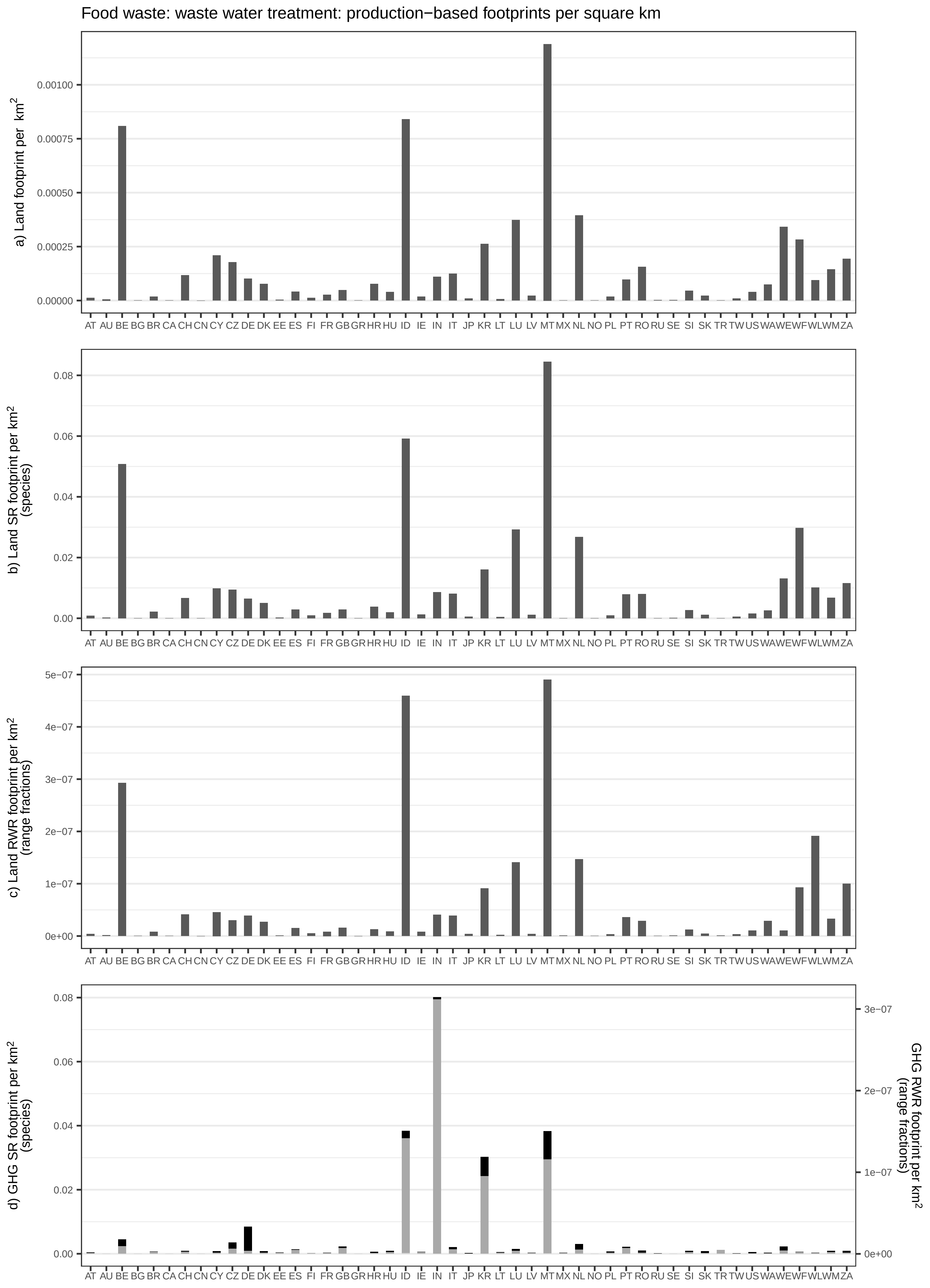

Food waste: waste water treatment: production−based footprints per square km
a) Land footprint per km²
b) Land SR footprint per km² (species)
c) Land RWR footprint per km² (range fractions)
d) GHG SR footprint per km² (species)
GHG RWR footprint per km² (range fractions)
AT AU BE BG BR CA CH CN CY CZ DE DK EE ES FI FR GB GR HR HU ID IE IN IT JP KR LT LU LV MT MX NL NO PL PT RO RU SE SI SK TR TW US WA WE WF WL WM ZA

Food waste: landfill: production−based footprints per square km

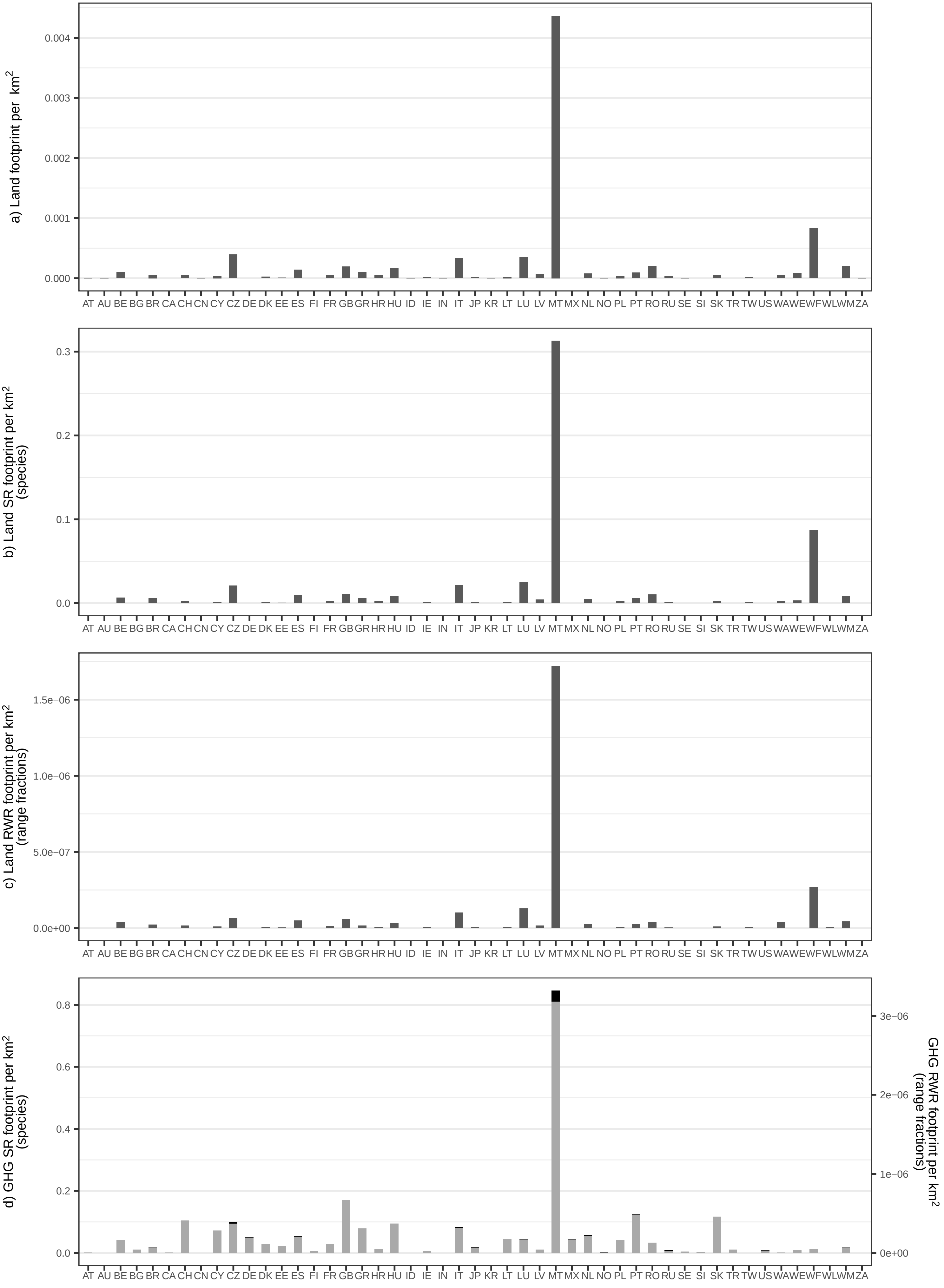
