## Supplementary Figure 8 for "Impacts of global food supply on biodiversity via land use and climate change"

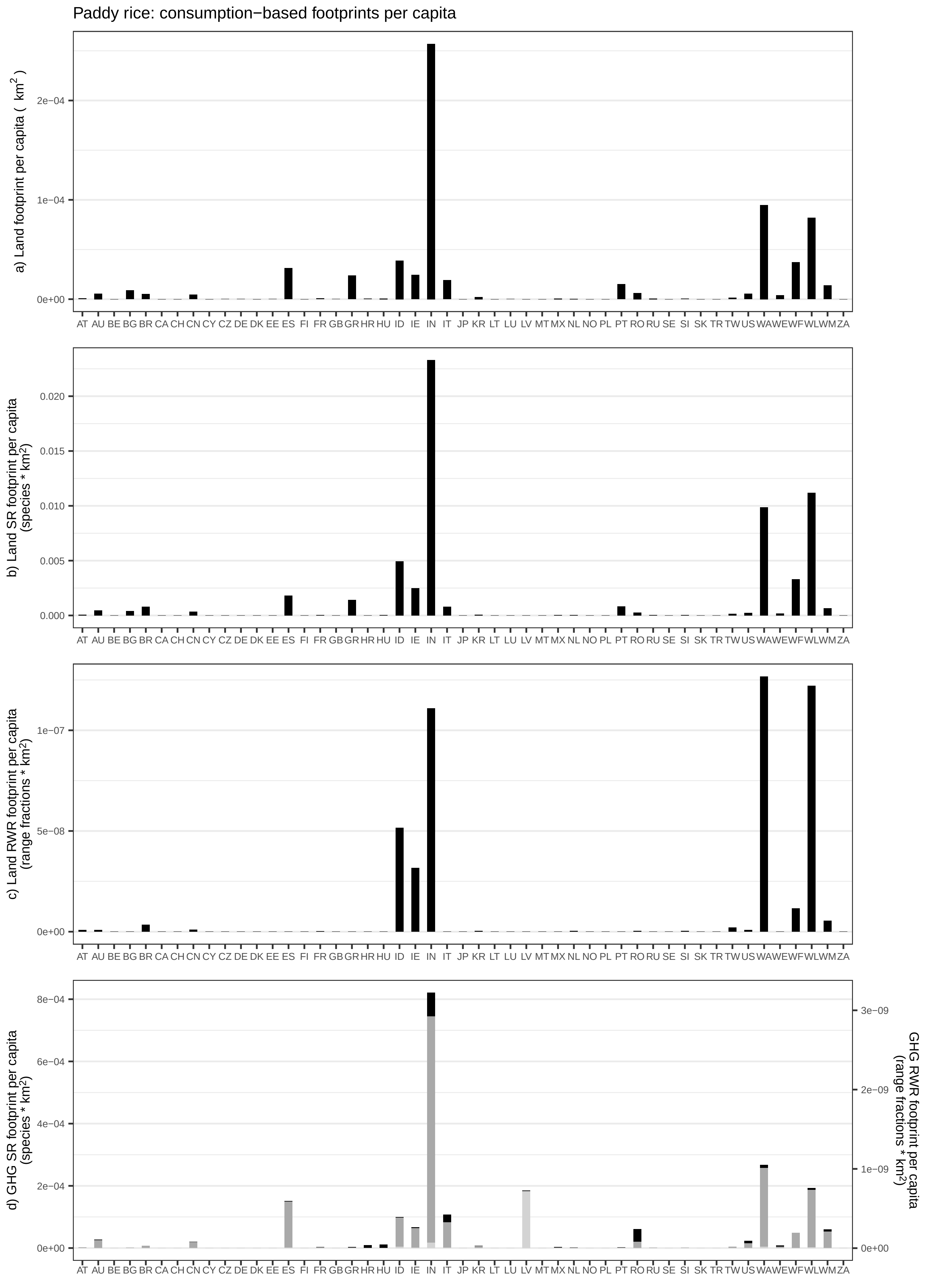

Paddy rice: consumption−based footprints per capita
a) Land footprint per capita ( km²)
b) Land SR footprint per capita (species * km²)
c) Land RWR footprint per capita (range fractions * km²)
d) GHG SR footprint per capita (species * km²)
GHG RWR footprint per capita (range fractions * km²)
AT AU BE BG BR CA CH CN CY CZ DE DK EE ES FI FR GB GR HR HU ID IE IN IT JP KR LT LU LV MT MX NL NO PL PT RO RU SE SI SK TR TW US WA WE WF WL WM ZA

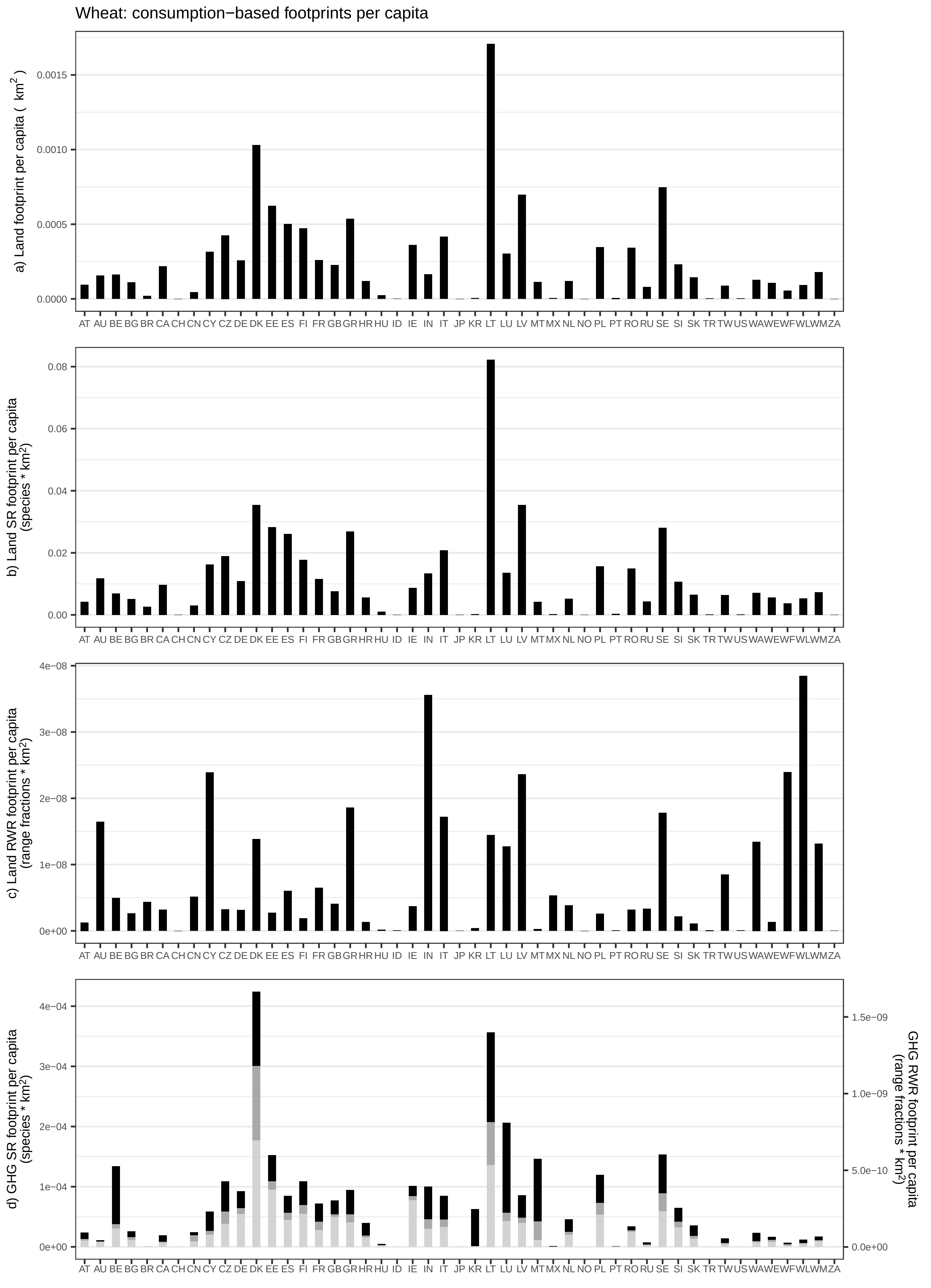

Wheat: consumption−based footprints per capita
a) Land footprint per capita ( km2 )
0.0015
0.0010
0.0005
0.0000
AT AU BE BG BR CA CH CN CY CZ DE DK EE ES FI FR GB GR HR HU ID IE IN IT JP KR LT LU LV MT MX NL NO PL PT RO RU SE SI SK TR TW US WA WE WF WL WM ZA
b) Land SR footprint per capita (species * km2)
0.08
0.06
0.04
0.02
0.00
c) Land RWR footprint per capita (range fractions * km2)
4e−08
3e−08
2e−08
1e−08
0e+00
d) GHG SR footprint per capita (species * km2)
4e−04
3e−04
2e−04
1e−04
0e+00
GHG RWR footprint per capita (range fractions * km2)
1.5e−09
1.0e−09
5.0e−10
0.0e+00

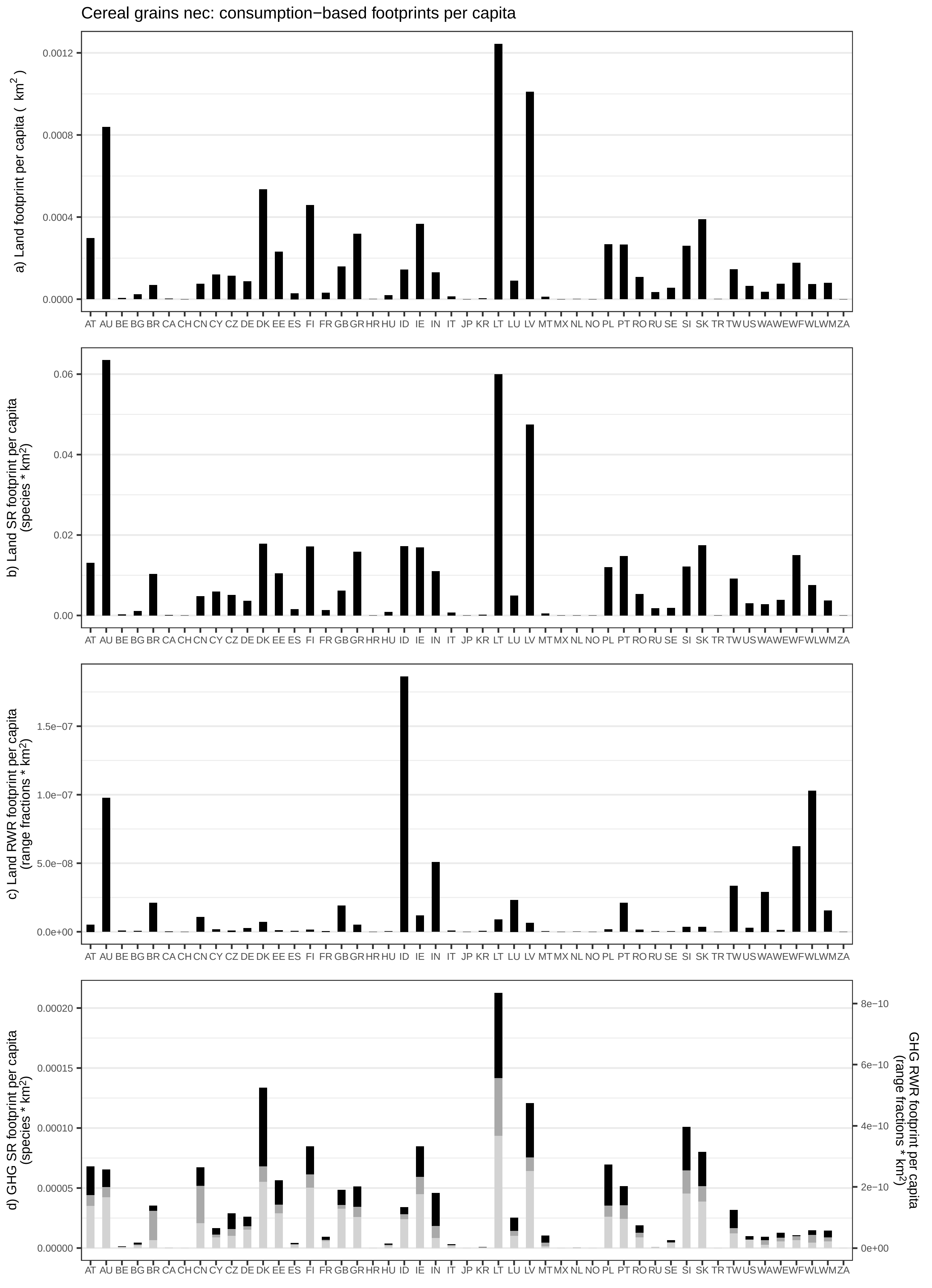
Cereal grains nec: consumption−based footprints per capita
a) Land footprint per capita ( $km^2$ )
0.0012
0.0008
0.0004
0.0000
AT AU BE BG BR CA CH CN CY CZ DE DK EE ES FI FR GB GR HR HU ID IE IN IT JP KR LT LU LV MT MX NL NO PL PT RO RU SE SI SK TR TW US WA WE WF WL WM ZA
b) Land SR footprint per capita (species * $km^2$)
0.06
0.04
0.02
0.00
AT AU BE BG BR CA CH CN CY CZ DE DK EE ES FI FR GB GR HR HU ID IE IN IT JP KR LT LU LV MT MX NL NO PL PT RO RU SE SI SK TR TW US WA WE WF WL WM ZA
c) Land RWR footprint per capita (range fractions * $km^2$)
1.5e−07
1.0e−07
5.0e−08
0.0e+00
AT AU BE BG BR CA CH CN CY CZ DE DK EE ES FI FR GB GR HR HU ID IE IN IT JP KR LT LU LV MT MX NL NO PL PT RO RU SE SI SK TR TW US WA WE WF WL WM ZA
d) GHG SR footprint per capita (species * $km^2$)
0.00020
0.00015
0.00010
0.00005
0.00000
GHG RWR footprint per capita (range fractions * $km^2$)
8e−10
6e−10
4e−10
2e−10
0e+00
AT AU BE BG BR CA CH CN CY CZ DE DK EE ES FI FR GB GR HR HU ID IE IN IT JP KR LT LU LV MT MX NL NO PL PT RO RU SE SI SK TR TW US WA WE WF WL WM ZA

Veg/Fruit/Nuts: consumption−based footprints per capita

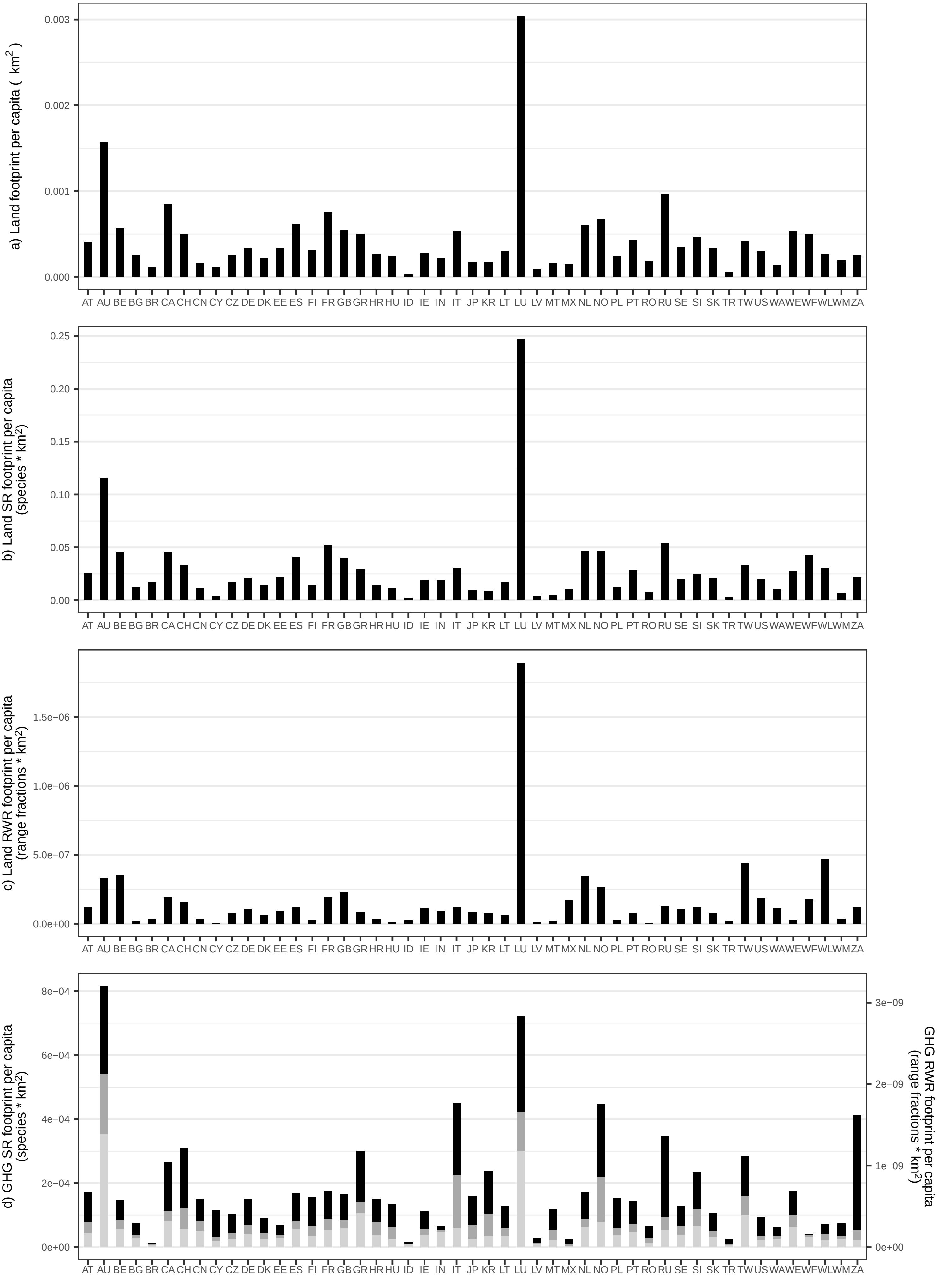

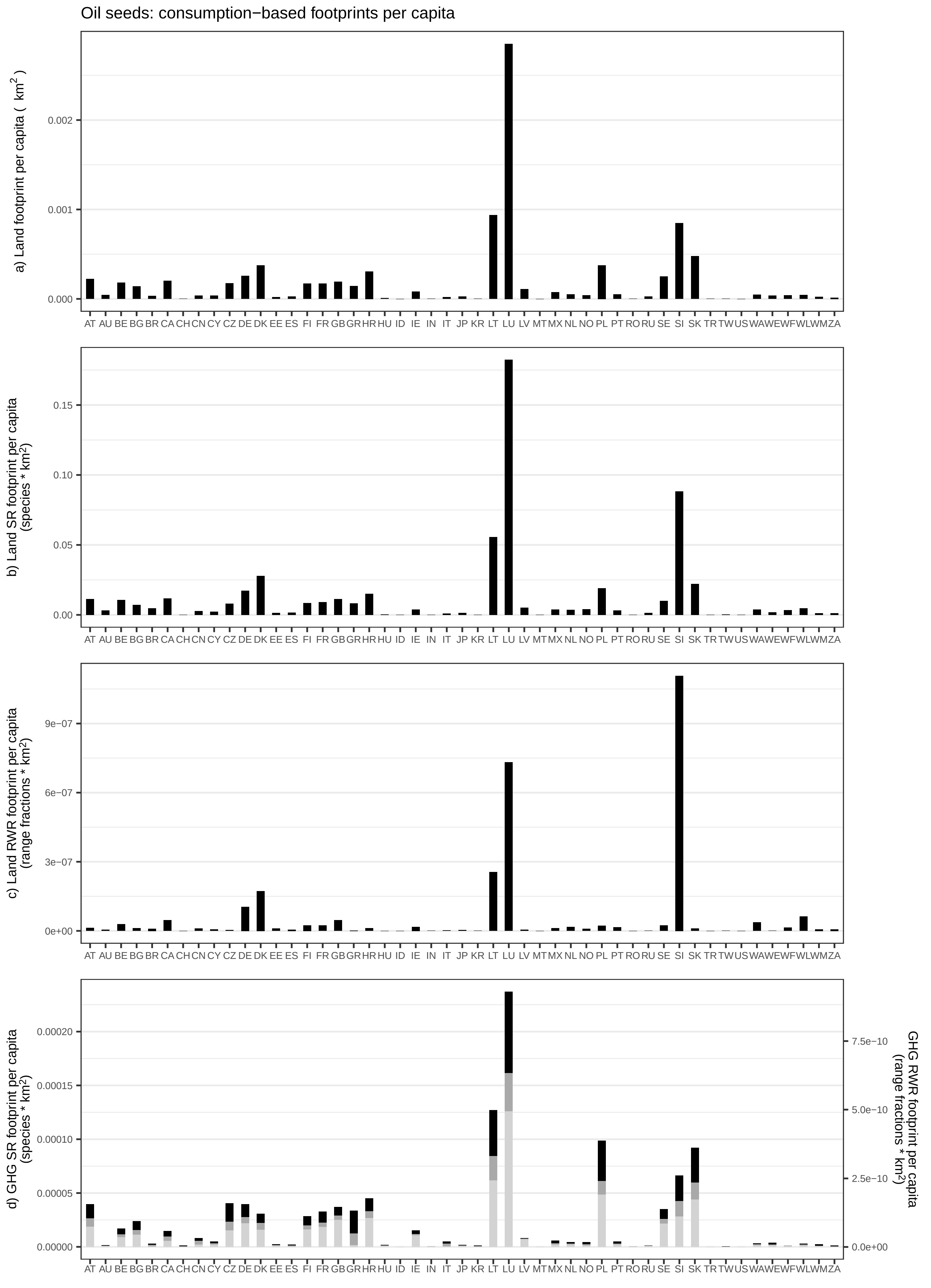

Oil seeds: consumption−based footprints per capita
a) Land footprint per capita ( km²)
0.000
0.001
0.002
b) Land SR footprint per capita (species * km²)
0.00
0.05
0.10
0.15
c) Land RWR footprint per capita (range fractions * km²)
0e+00
3e−07
6e−07
9e−07
d) GHG SR footprint per capita (species * km²)
0.00000
0.00005
0.00010
0.00015
0.00020
GHG RWR footprint per capita (range fractions * km²)
0.0e+00
2.5e−10
5.0e−10
7.5e−10
AT AU BE BG BR CA CH CN CY CZ DE DK EE ES FI FR GB GR HR HU ID IE IN IT JP KR LT LU LV MT MX NL NO PL PT RO RU SE SI SK TR TW US WA WE WF WL WM ZA

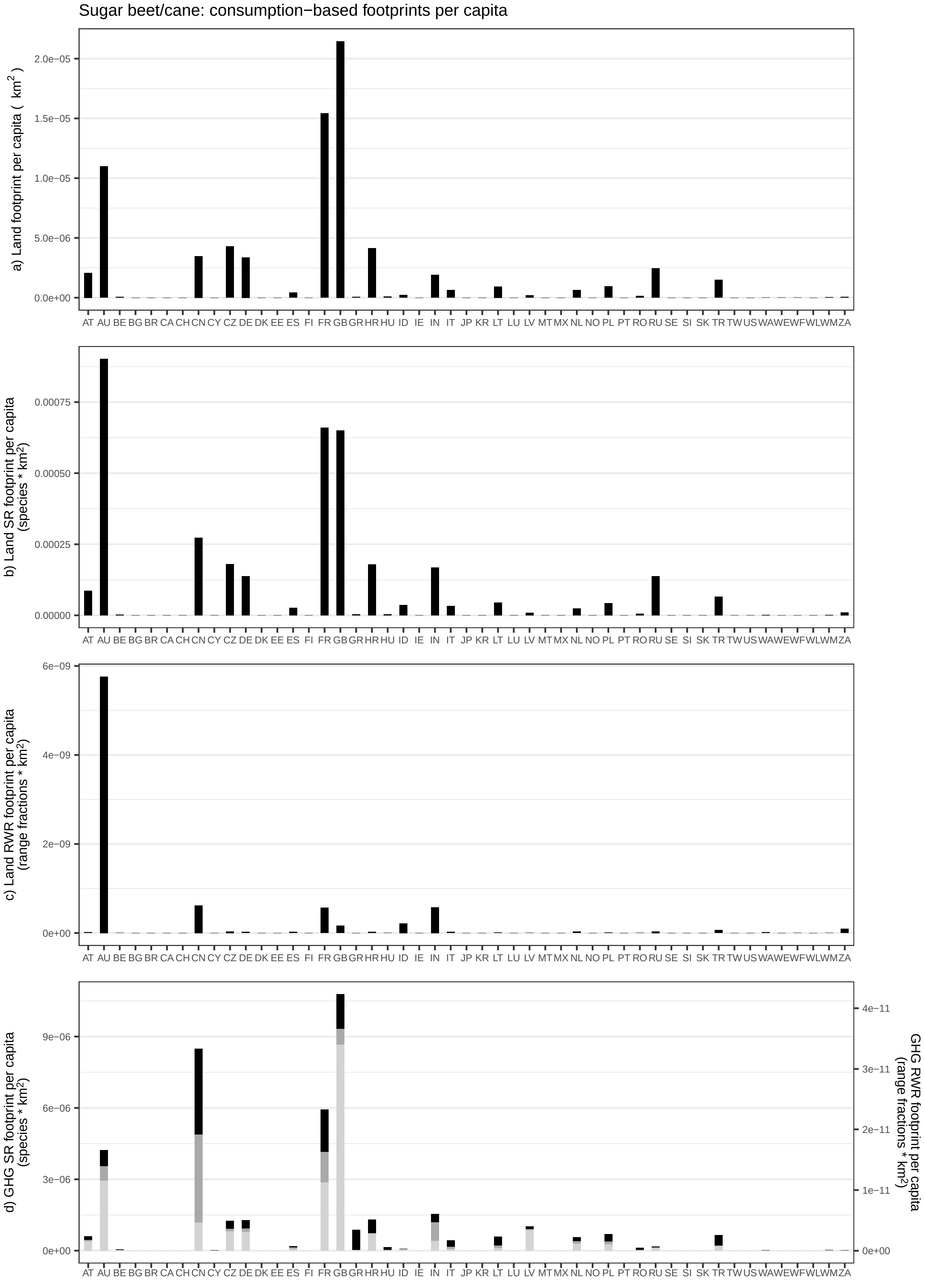
Sugar beet/cane: consumption−based footprints per capita
a) Land footprint per capita ( km2 )
b) Land SR footprint per capita (species * km2)
c) Land RWR footprint per capita (range fractions * km2)
d) GHG SR footprint per capita (species * km2)
GHG RWR footprint per capita (range fractions * km2)
AT AU BE BG BR CA CH CN CY CZ DE DK EE ES FI FR GB GR HR HU ID IE IN IT JP KR LT LU LV MT MX NL NO PL PT RO RU SE SI SK TR TW US WA WE WF WL WM ZA

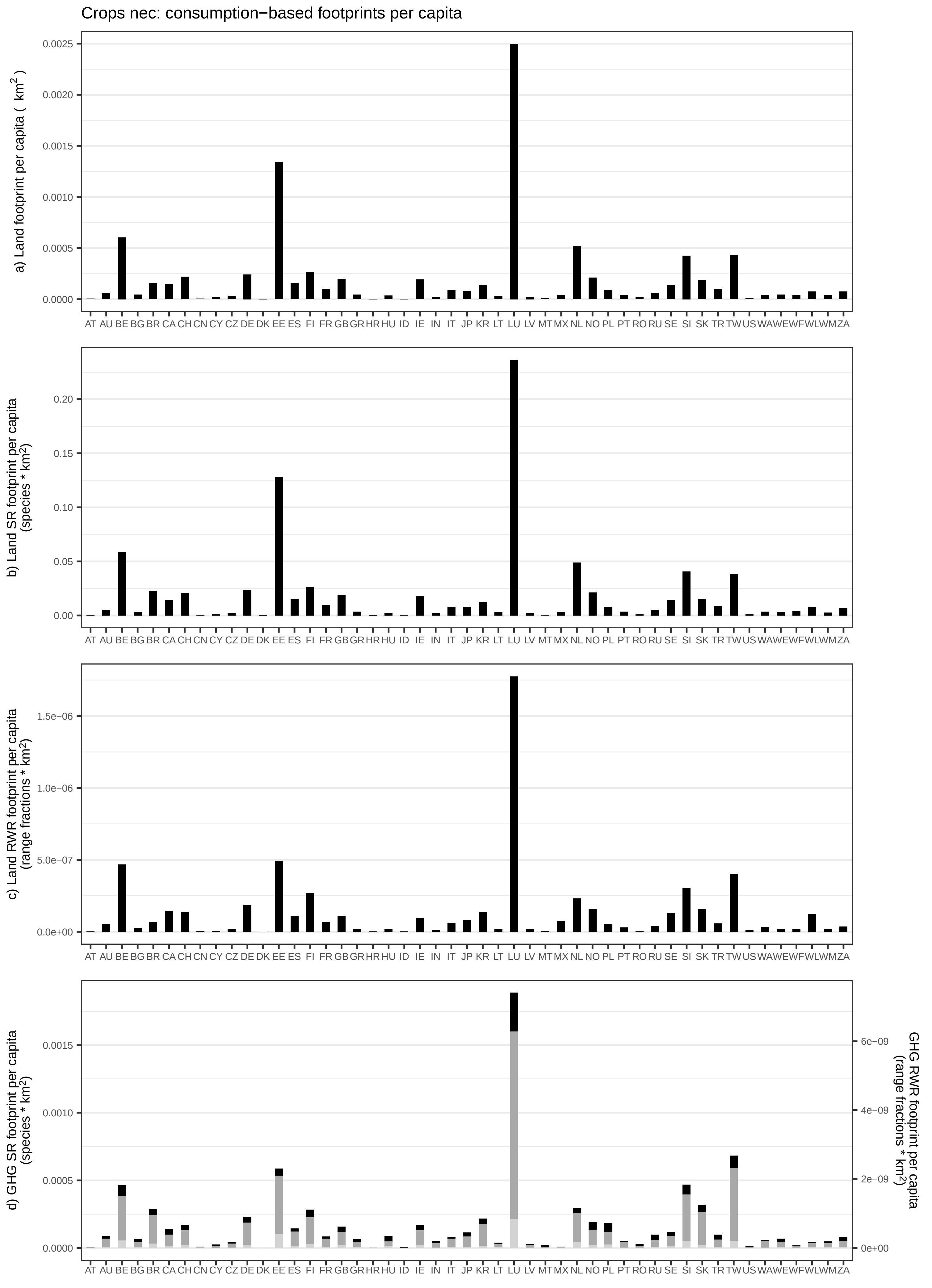
Crops nec: consumption−based footprints per capita
a) Land footprint per capita ( km²)
b) Land SR footprint per capita (species * km²)
c) Land RWR footprint per capita (range fractions * km²)
d) GHG SR footprint per capita (species * km²)
GHG RWR footprint per capita (range fractions * km²)
AT AU BE BG BR CA CH CN CY CZ DE DK EE ES FI FR GB GR HR HU ID IE IN IT JP KR LT LU LV MT MX NL NO PL PT RO RU SE SI SK TR TW US WA WE WF WL WM ZA

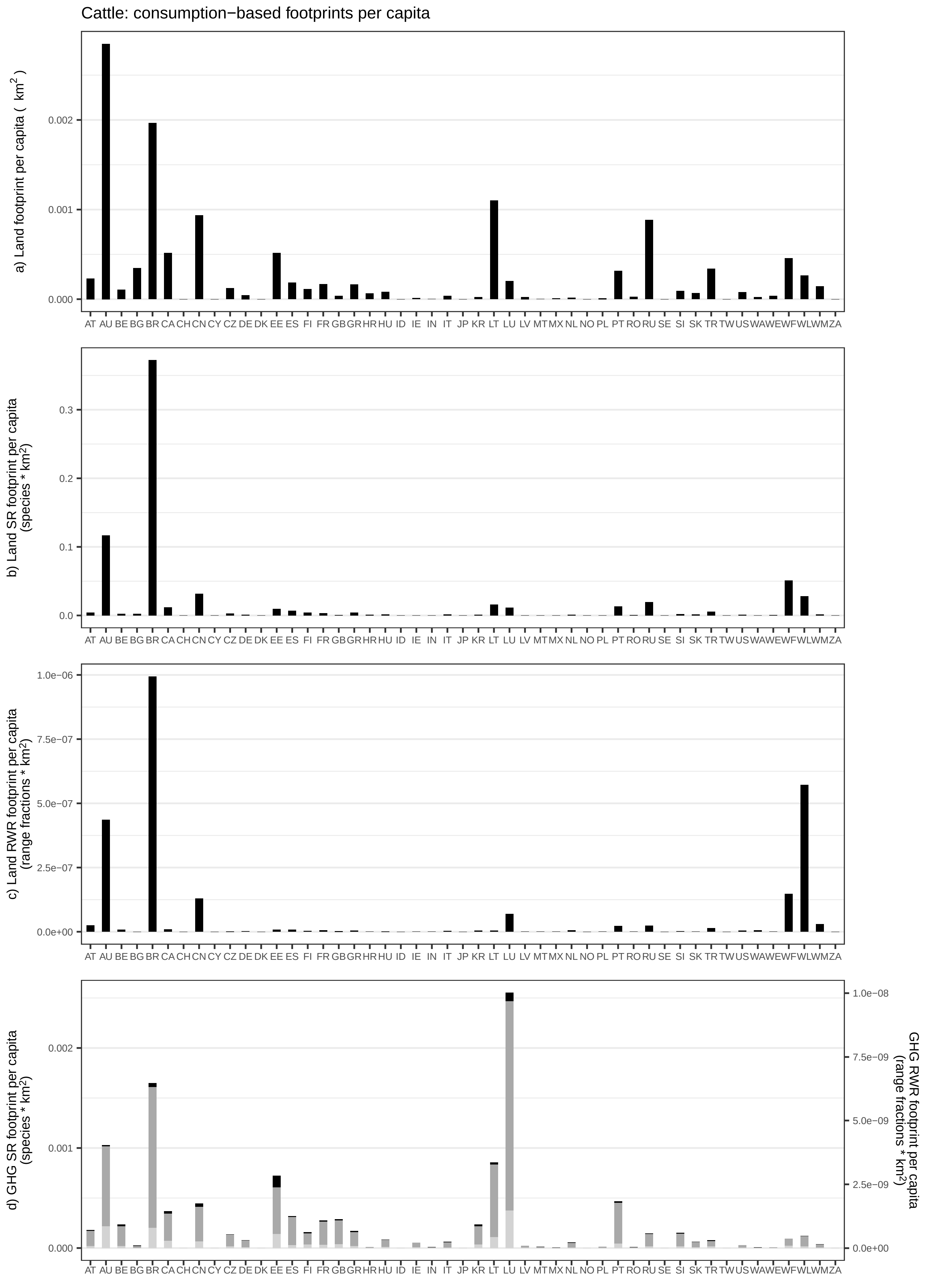

Cattle: consumption−based footprints per capita
a) Land footprint per capita ( km$^2$ )
0.000
0.001
0.002
AT AU BE BG BR CA CH CN CY CZ DE DK EE ES FI FR GB GR HR HU ID IE IN IT JP KR LT LU LV MT MX NL NO PL PT RO RU SE SI SK TR TW US WA WE WF WL WM ZA
b) Land SR footprint per capita (species * km$^2$)
0.0
0.1
0.2
0.3
c) Land RWR footprint per capita (range fractions * km$^2$)
0.0e+00
2.5e−07
5.0e−07
7.5e−07
1.0e−06
d) GHG SR footprint per capita (species * km$^2$)
0.000
0.001
0.002
GHG RWR footprint per capita (range fractions * km$^2$)
0.0e+00
2.5e−09
5.0e−09
7.5e−09
1.0e−08

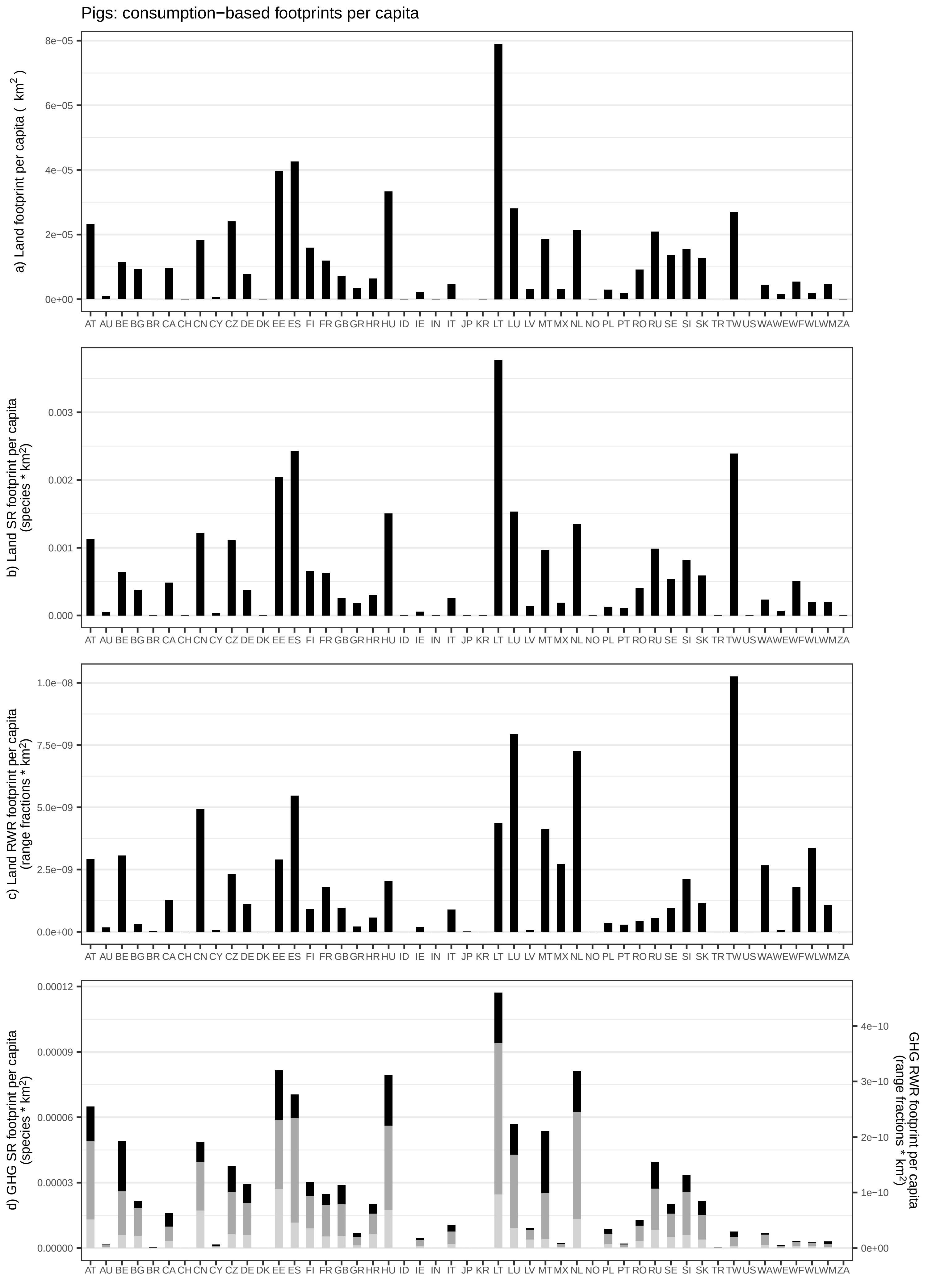

Pigs: consumption−based footprints per capita
a) Land footprint per capita ( km$^2$ )
8e−05
6e−05
4e−05
2e−05
0e+00
AT AU BE BG BR CA CH CN CY CZ DE DK EE ES FI FR GB GR HR HU ID IE IN IT JP KR LT LU LV MT MX NL NO PL PT RO RU SE SI SK TR TW US WA WE WF WL WM ZA
b) Land SR footprint per capita (species * km$^2$)
0.003
0.002
0.001
0.000
c) Land RWR footprint per capita (range fractions * km$^2$)
1.0e−08
7.5e−09
5.0e−09
2.5e−09
0.0e+00
d) GHG SR footprint per capita (species * km$^2$)
0.00012
0.00009
0.00006
0.00003
0.00000
GHG RWR footprint per capita (range fractions * km$^2$)
4e−10
3e−10
2e−10
1e−10
0e+00

Poultry: consumption−based footprints per capita

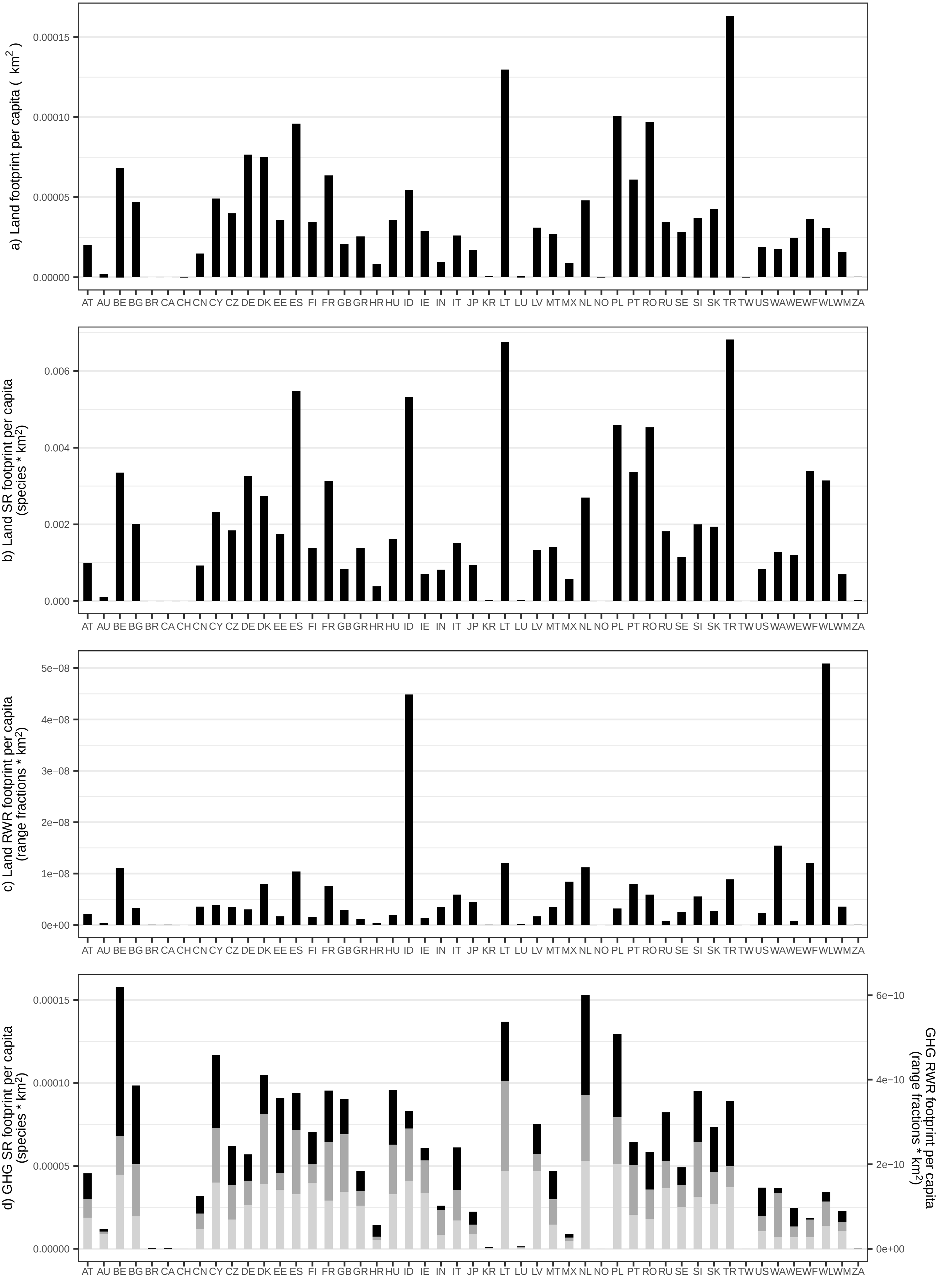

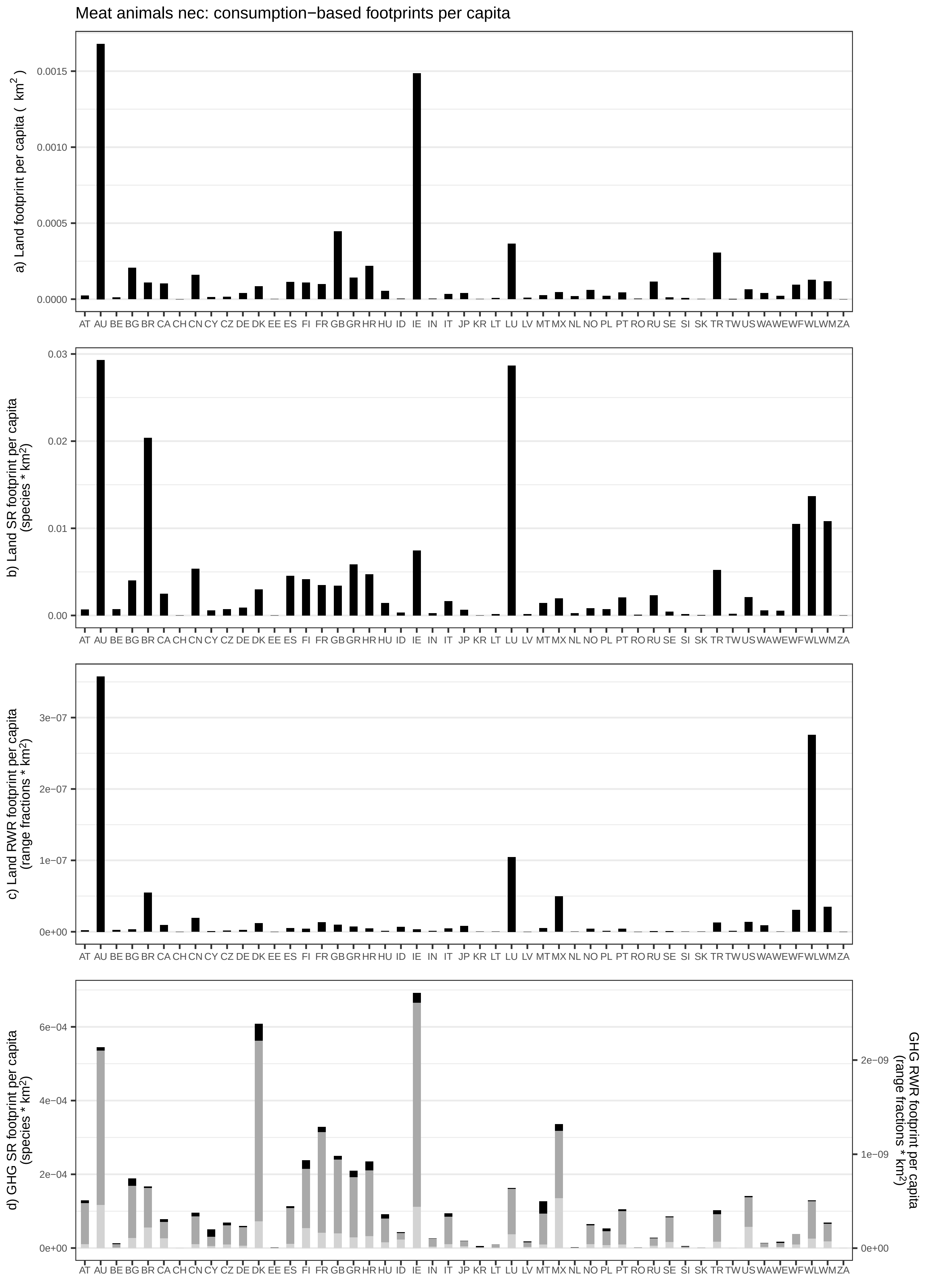

Meat animals nec: consumption−based footprints per capita
a) Land footprint per capita ( km2 )
b) Land SR footprint per capita (species * km2)
c) Land RWR footprint per capita (range fractions * km2)
d) GHG SR footprint per capita (species * km2)
GHG RWR footprint per capita (range fractions * km2)
AT AU BE BG BR CA CH CN CY CZ DE DK EE ES FI FR GB GR HR HU ID IE IN IT JP KR LT LU LV MT MX NL NO PL PT RO RU SE SI SK TR TW US WA WE WF WL WM ZA

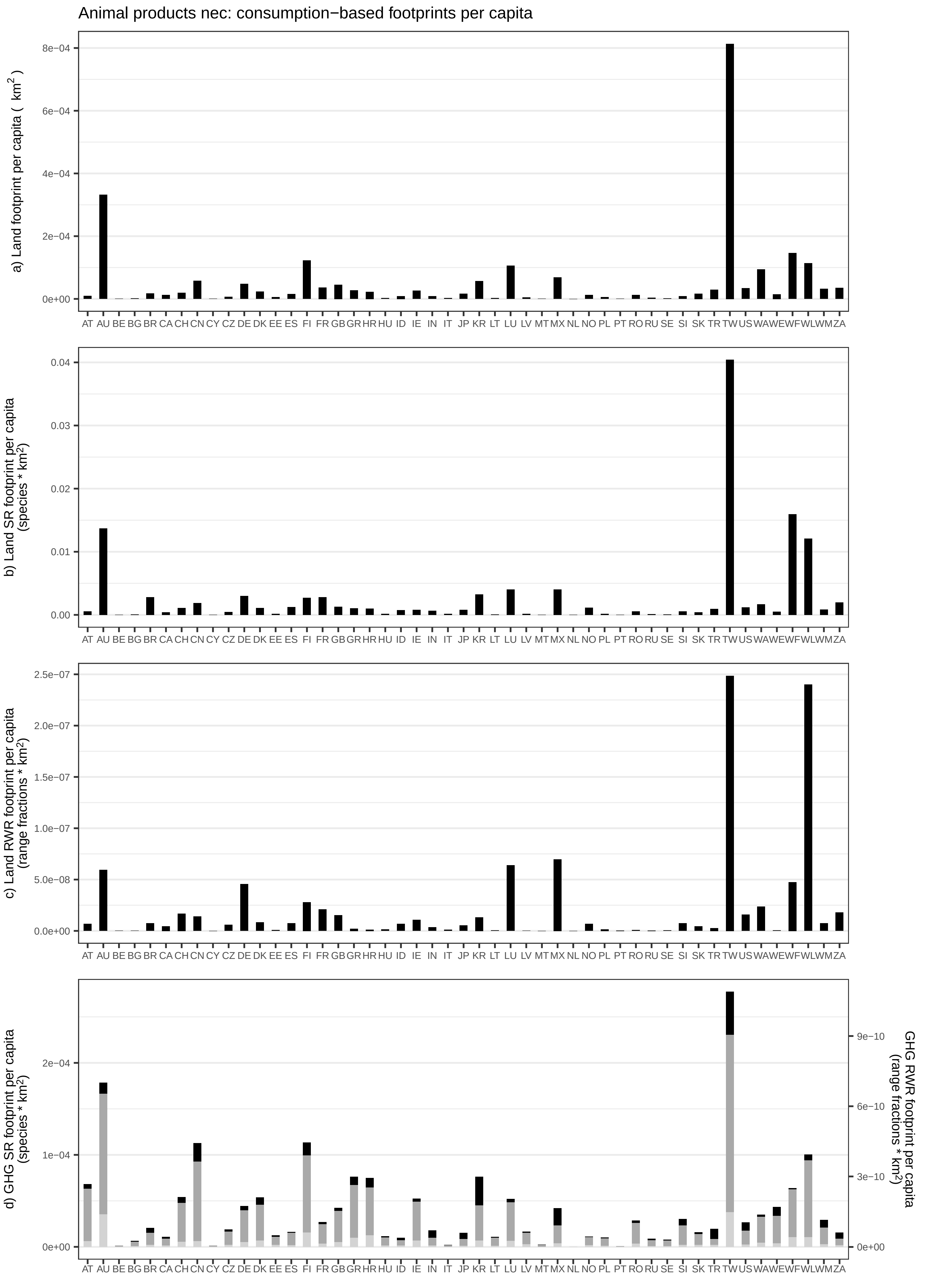
Animal products nec: consumption−based footprints per capita
a) Land footprint per capita ( km$^2$ )
8e−04
6e−04
4e−04
2e−04
0e+00
b) Land SR footprint per capita (species * km$^2$)
0.04
0.03
0.02
0.01
0.00
c) Land RWR footprint per capita (range fractions * km$^2$)
2.5e−07
2.0e−07
1.5e−07
1.0e−07
5.0e−08
0.0e+00
d) GHG SR footprint per capita (species * km$^2$)
2e−04
1e−04
0e+00
GHG RWR footprint per capita (range fractions * km$^2$)
9e−10
6e−10
3e−10
0e+00
AT AU BE BG BR CA CH CN CY CZ DE DK EE ES FI FR GB GR HR HU ID IE IN IT JP KR LT LU LV MT MX NL NO PL PT RO RU SE SI SK TR TW US WA WE WF WL WM ZA

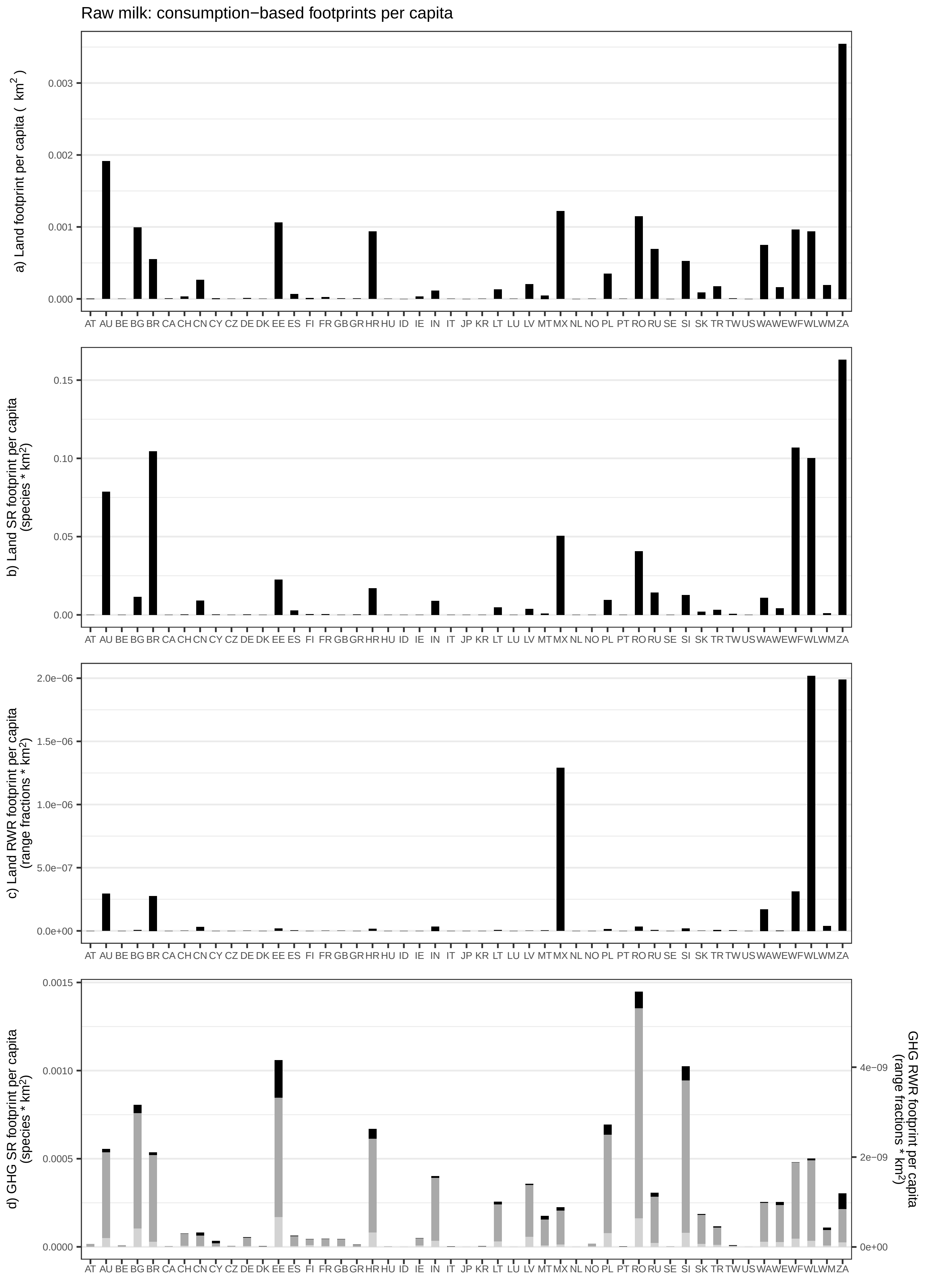
Raw milk: consumption−based footprints per capita
a) Land footprint per capita ( km2 )
b) Land SR footprint per capita (species * km2)
c) Land RWR footprint per capita (range fractions * km2)
d) GHG SR footprint per capita (species * km2)
GHG RWR footprint per capita (range fractions * km2)
AT AU BE BG BR CA CH CN CY CZ DE DK EE ES FI FR GB GR HR HU ID IE IN IT JP KR LT LU LV MT MX NL NO PL PT RO RU SE SI SK TR TW US WA WE WF WL WM ZA

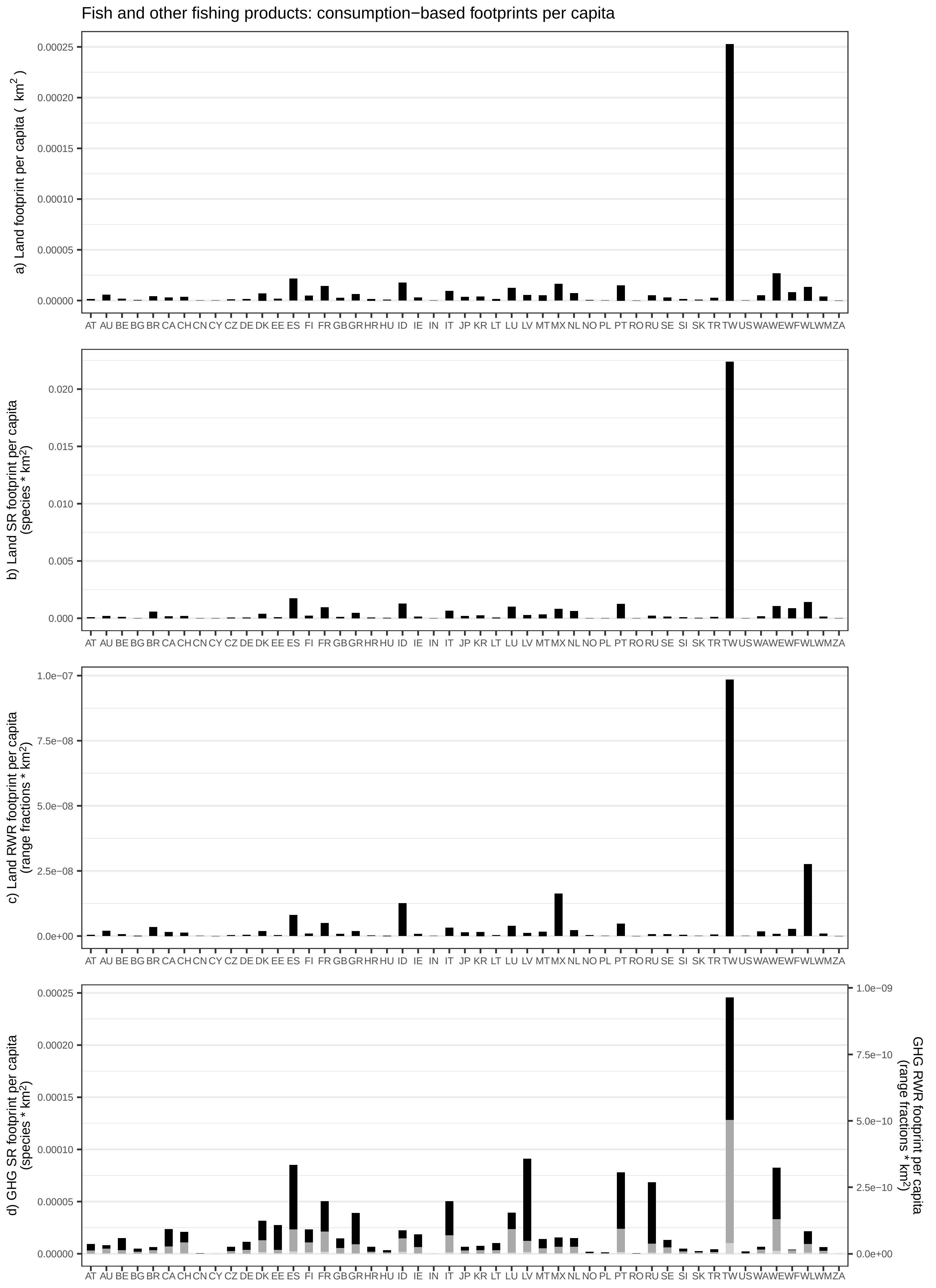

Fish and other fishing products: consumption−based footprints per capita
a) Land footprint per capita ( km²)
0.00025
0.00020
0.00015
0.00010
0.00005
0.00000
AT AU BE BG BR CA CH CN CY CZ DE DK EE ES FI FR GB GR HR HU ID IE IN IT JP KR LT LU LV MT MX NL NO PL PT RO RU SE SI SK TR TW US WA WE WF WL WM ZA
b) Land SR footprint per capita (species * km²)
0.020
0.015
0.010
0.005
0.000
c) Land RWR footprint per capita (range fractions * km²)
1.0e−07
7.5e−08
5.0e−08
2.5e−08
0.0e+00
d) GHG SR footprint per capita (species * km²)
GHG RWR footprint per capita (range fractions * km²)
1.0e−09
7.5e−10
5.0e−10
2.5e−10
0.0e+00

Chemical and fertiliser minerals: consumption−based footprints per capita

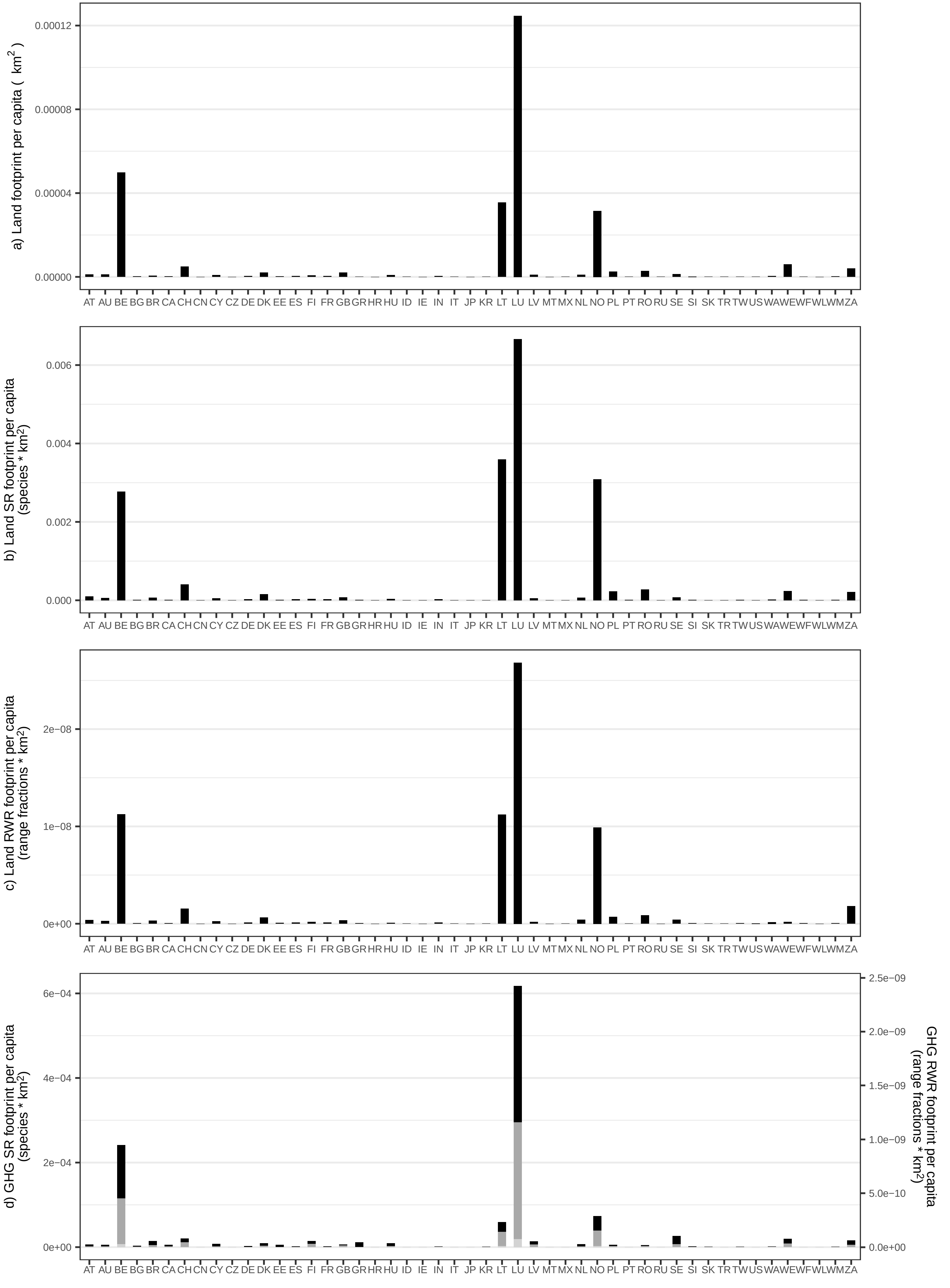

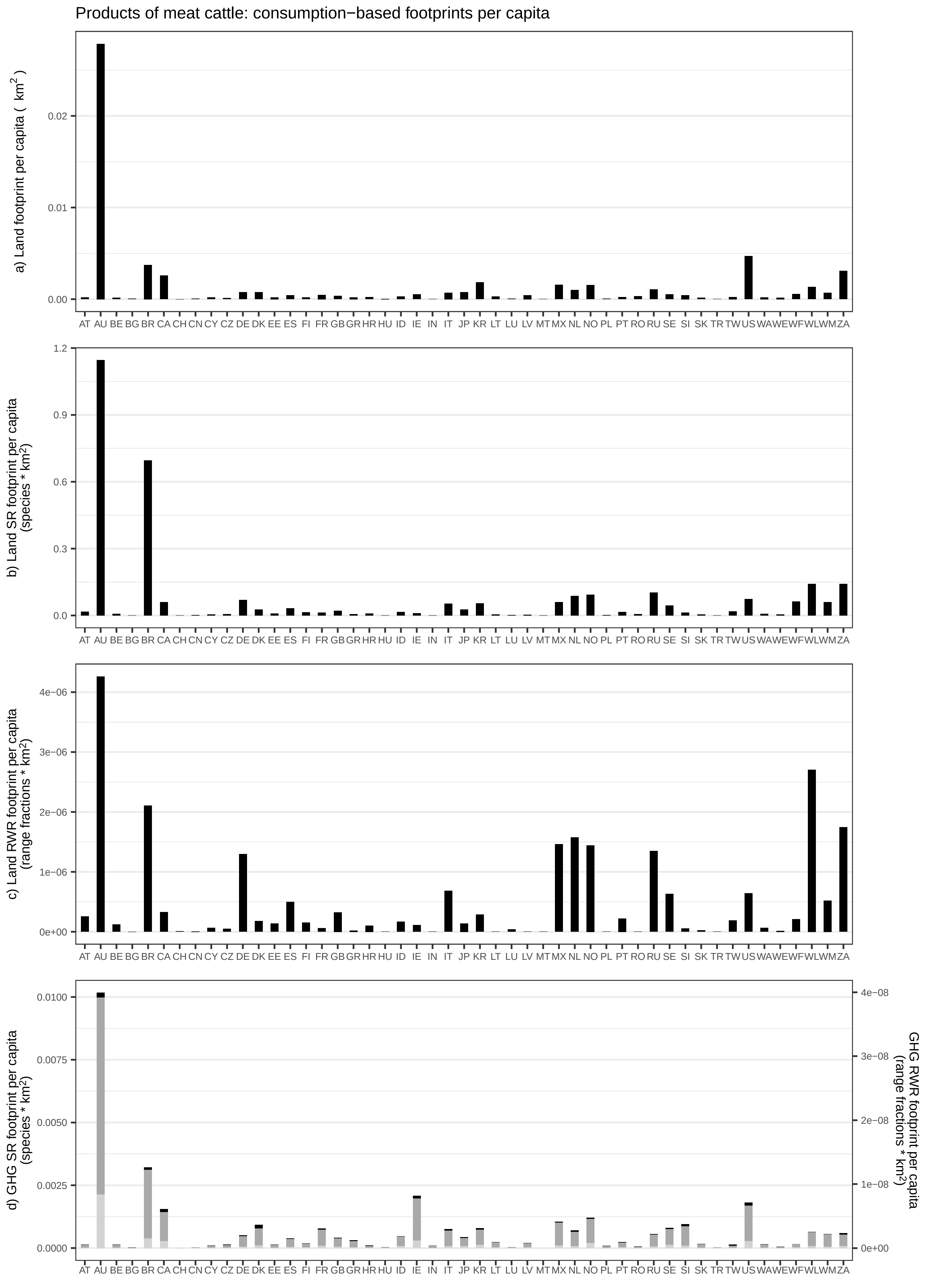

Products of meat cattle: consumption−based footprints per capita
a) Land footprint per capita ( km²)
b) Land SR footprint per capita (species * km²)
c) Land RWR footprint per capita (range fractions * km²)
d) GHG SR footprint per capita (species * km²)
GHG RWR footprint per capita (range fractions * km²)
AT AU BE BG BR CA CH CN CY CZ DE DK EE ES FI FR GB GR HR HU ID IE IN IT JP KR LT LU LV MT MX NL NO PL PT RO RU SE SI SK TR TW US WA WE WF WL WM ZA

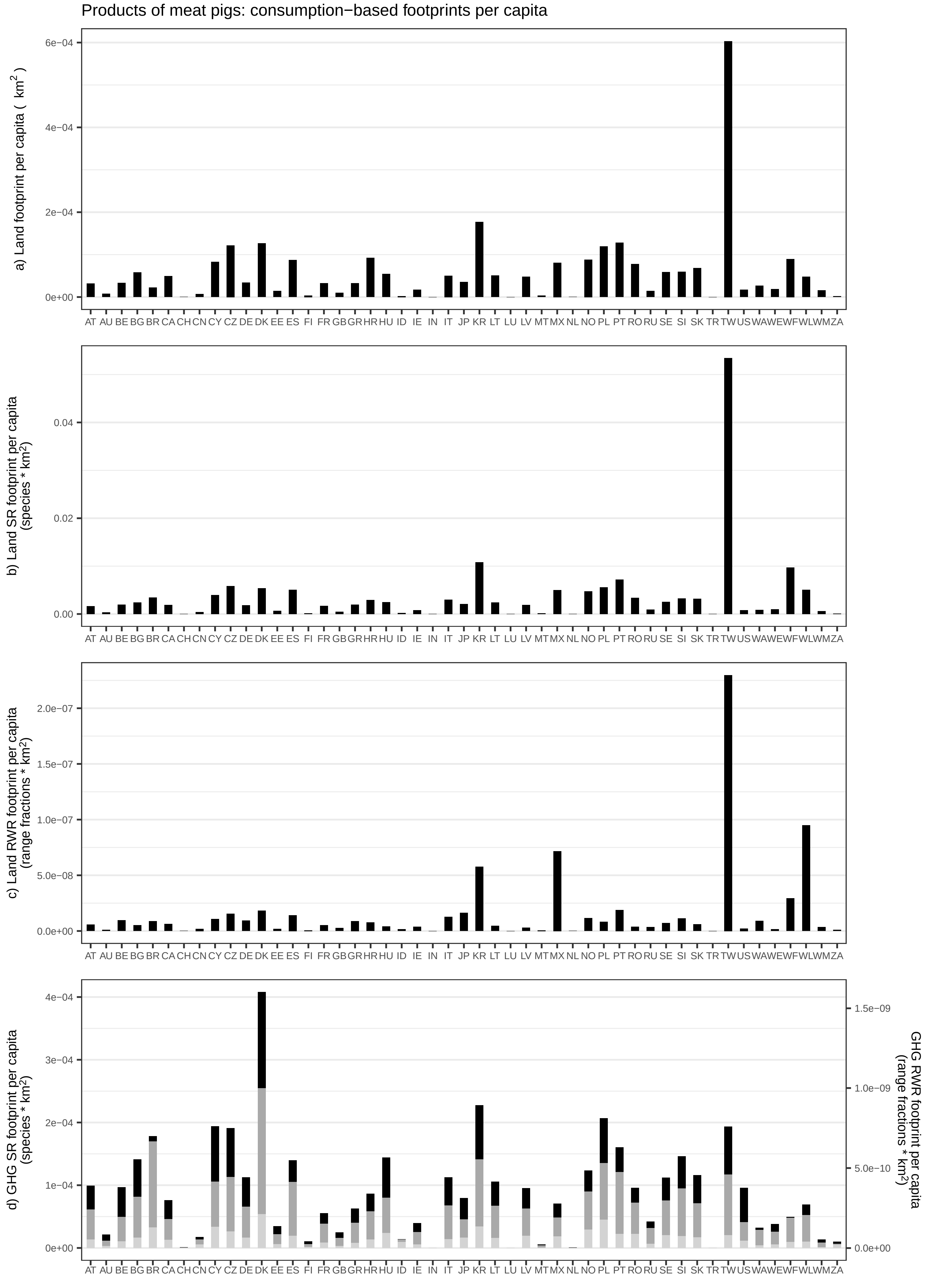

Products of meat pigs: consumption−based footprints per capita
a) Land footprint per capita ( km$^2$ )
b) Land SR footprint per capita (species * km$^2$)
c) Land RWR footprint per capita (range fractions * km$^2$)
d) GHG SR footprint per capita (species * km$^2$)
GHG RWR footprint per capita (range fractions * km$^2$)
AT AU BE BG BR CA CH CN CY CZ DE DK EE ES FI FR GB GR HR HU ID IE IN IT JP KR LT LU LV MT MX NL NO PL PT RO RU SE SI SK TR TW US WA WE WF WL WM ZA

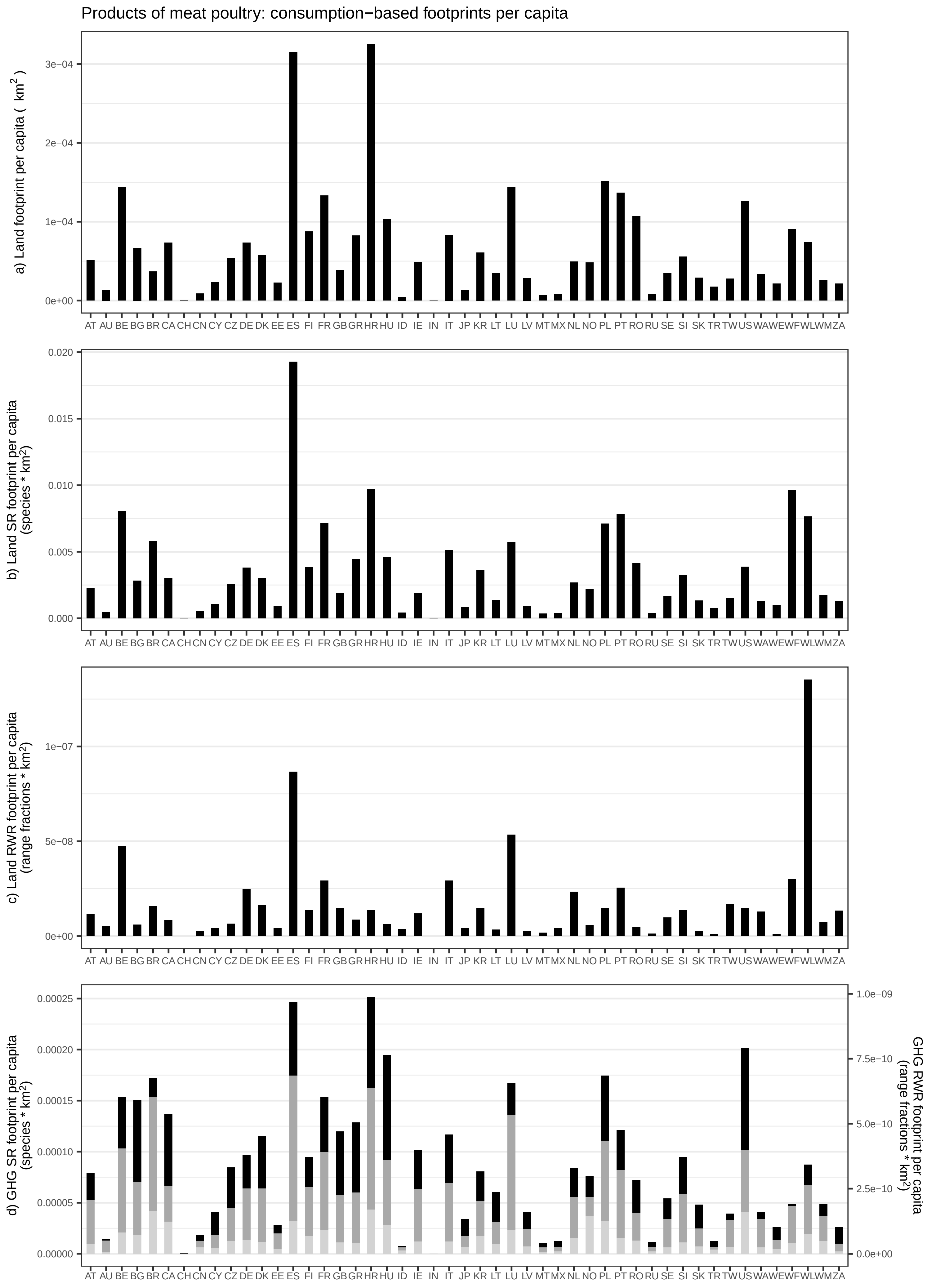
Products of meat poultry: consumption−based footprints per capita
a) Land footprint per capita ( km2 )
3e−04
2e−04
1e−04
0e+00
AT AU BE BG BR CA CH CN CY CZ DE DK EE ES FI FR GB GR HR HU ID IE IN IT JP KR LT LU LV MT MX NL NO PL PT RO RU SE SI SK TR TW US WA WE WF WL WM ZA
b) Land SR footprint per capita (species * km2)
0.020
0.015
0.010
0.005
0.000
c) Land RWR footprint per capita (range fractions * km2)
1e−07
5e−08
0e+00
d) GHG SR footprint per capita (species * km2)
0.00025
0.00020
0.00015
0.00010
0.00005
0.00000
GHG RWR footprint per capita (range fractions * km2)
1.0e−09
7.5e−10
5.0e−10
2.5e−10
0.0e+00

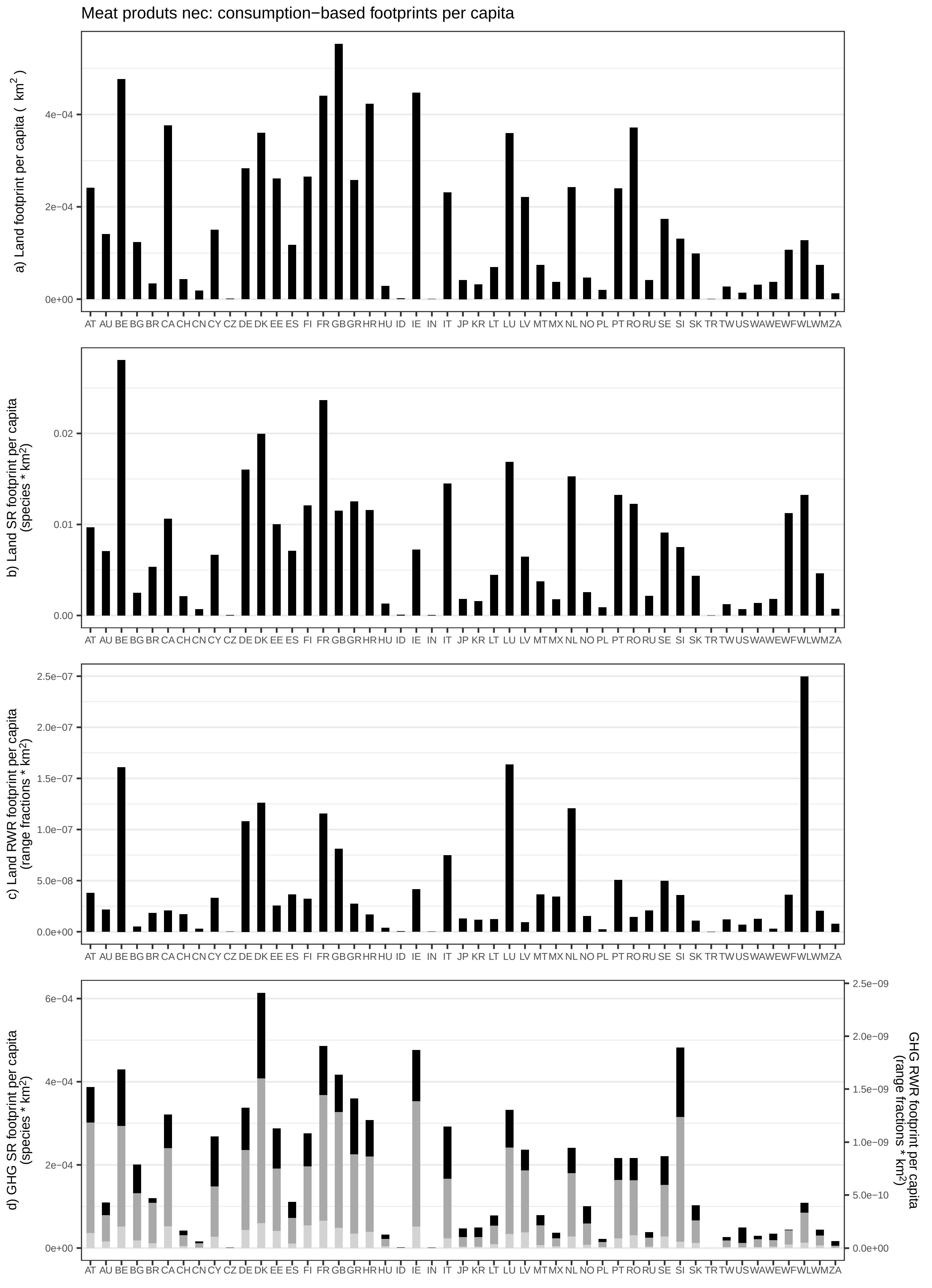

Meat produts nec: consumption−based footprints per capita
a) Land footprint per capita ( km2 )
0e+00
2e−04
4e−04
b) Land SR footprint per capita (species * km2)
0.00
0.01
0.02
c) Land RWR footprint per capita (range fractions * km2)
0.0e+00
5.0e−08
1.0e−07
1.5e−07
2.0e−07
2.5e−07
d) GHG SR footprint per capita (species * km2)
0e+00
2e−04
4e−04
6e−04
GHG RWR footprint per capita (range fractions * km2)
0.0e+00
5.0e−10
1.0e−09
1.5e−09
2.0e−09
2.5e−09
AT AU BE BG BR CA CH CN CY CZ DE DK EE ES FI FR GB GR HR HU ID IE IN IT JP KR LT LU LV MT MX NL NO PL PT RO RU SE SI SK TR TW US WA WE WF WL WM ZA

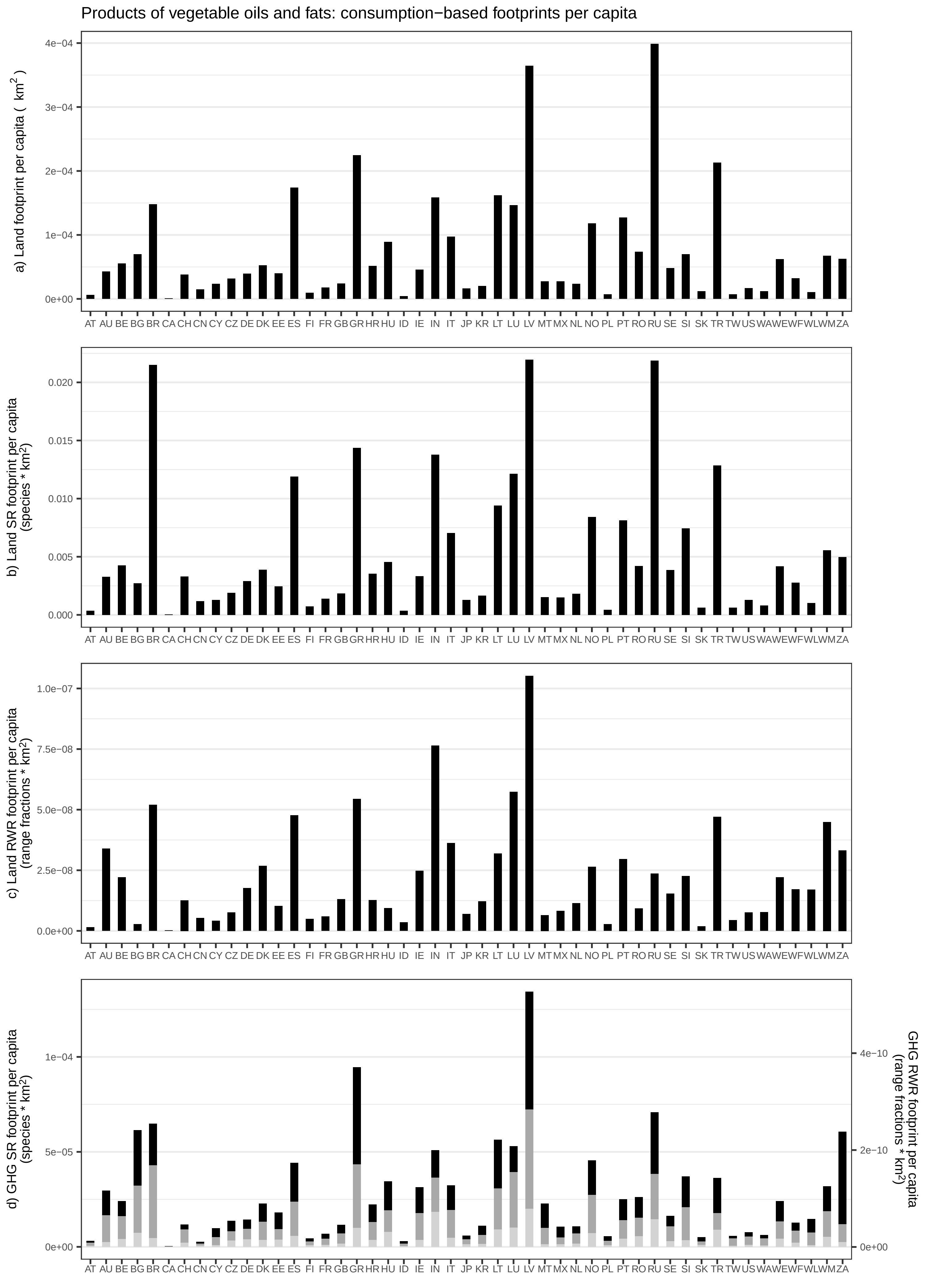
Products of vegetable oils and fats: consumption−based footprints per capita
a) Land footprint per capita ( km$^2$ )
4e−04
3e−04
2e−04
1e−04
0e+00
AT AU BE BG BR CA CH CN CY CZ DE DK EE ES FI FR GB GR HR HU ID IE IN IT JP KR LT LU LV MT MX NL NO PL PT RO RU SE SI SK TR TW US WA WE WF WL WM ZA
b) Land SR footprint per capita (species * km$^2$)
0.020
0.015
0.010
0.005
0.000
AT AU BE BG BR CA CH CN CY CZ DE DK EE ES FI FR GB GR HR HU ID IE IN IT JP KR LT LU LV MT MX NL NO PL PT RO RU SE SI SK TR TW US WA WE WF WL WM ZA
c) Land RWR footprint per capita (range fractions * km$^2$)
1.0e−07
7.5e−08
5.0e−08
2.5e−08
0.0e+00
AT AU BE BG BR CA CH CN CY CZ DE DK EE ES FI FR GB GR HR HU ID IE IN IT JP KR LT LU LV MT MX NL NO PL PT RO RU SE SI SK TR TW US WA WE WF WL WM ZA
d) GHG SR footprint per capita (species * km$^2$)
1e−04
5e−05
0e+00
GHG RWR footprint per capita (range fractions * km$^2$)
4e−10
2e−10
0e+00
AT AU BE BG BR CA CH CN CY CZ DE DK EE ES FI FR GB GR HR HU ID IE IN IT JP KR LT LU LV MT MX NL NO PL PT RO RU SE SI SK TR TW US WA WE WF WL WM ZA

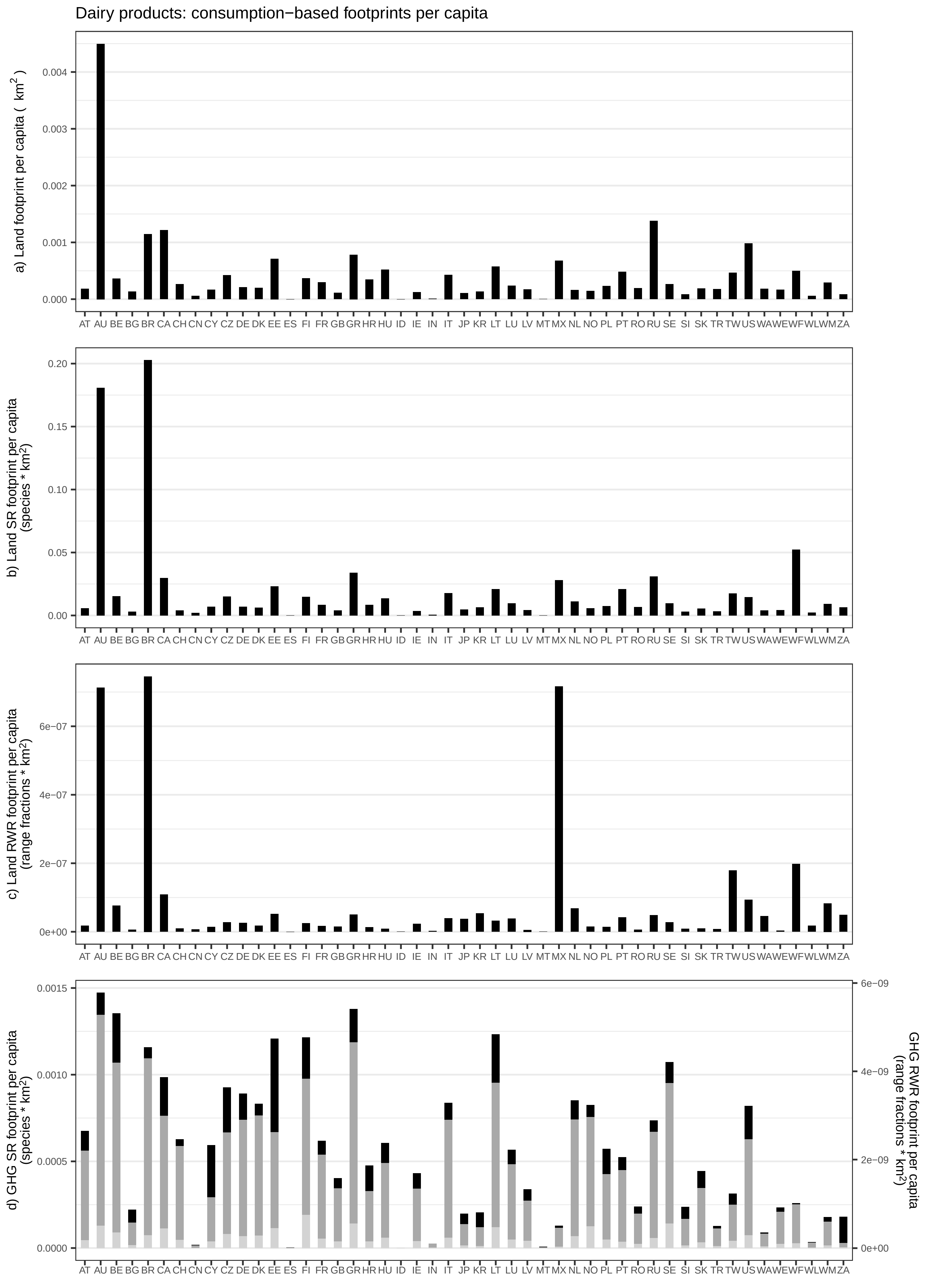

Dairy products: consumption−based footprints per capita
a) Land footprint per capita ( km²)
b) Land SR footprint per capita (species * km²)
c) Land RWR footprint per capita (range fractions * km²)
d) GHG SR footprint per capita (species * km²)
GHG RWR footprint per capita (range fractions * km²)
AT AU BE BG BR CA CH CN CY CZ DE DK EE ES FI FR GB GR HR HU ID IE IN IT JP KR LT LU LV MT MX NL NO PL PT RO RU SE SI SK TR TW US WA WE WF WL WM ZA

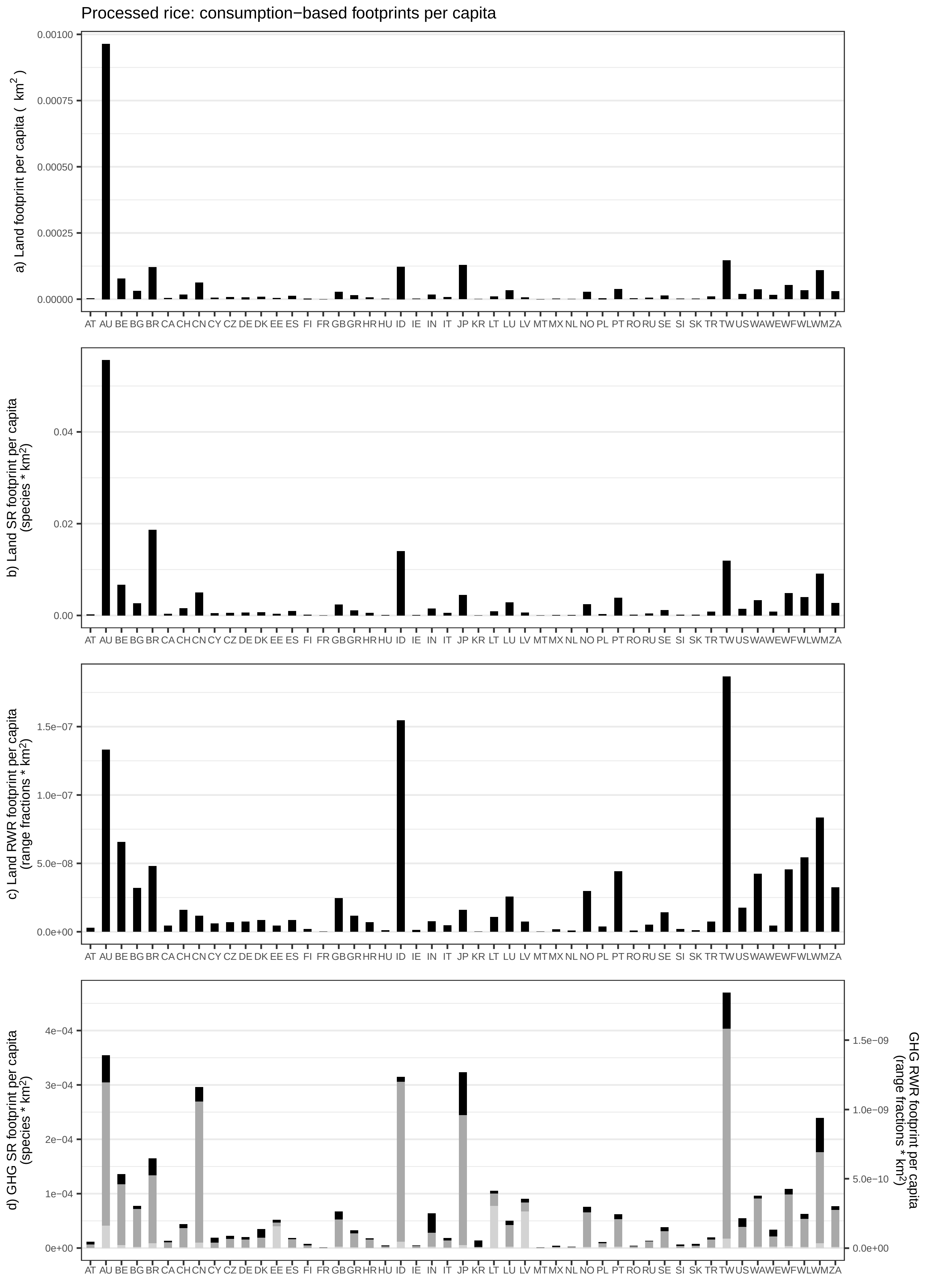

Processed rice: consumption−based footprints per capita
a) Land footprint per capita ( km2 )
0.00100
0.00075
0.00050
0.00025
0.00000
b) Land SR footprint per capita (species * km2)
0.04
0.02
0.00
c) Land RWR footprint per capita (range fractions * km2)
1.5e−07
1.0e−07
5.0e−08
0.0e+00
d) GHG SR footprint per capita (species * km2)
4e−04
3e−04
2e−04
1e−04
0e+00
GHG RWR footprint per capita (range fractions * km2)
1.5e−09
1.0e−09
5.0e−10
0.0e+00
AT AU BE BG BR CA CH CN CY CZ DE DK EE ES FI FR GB GR HR HU ID IE IN IT JP KR LT LU LV MT MX NL NO PL PT RO RU SE SI SK TR TW US WA WE WF WL WM ZA

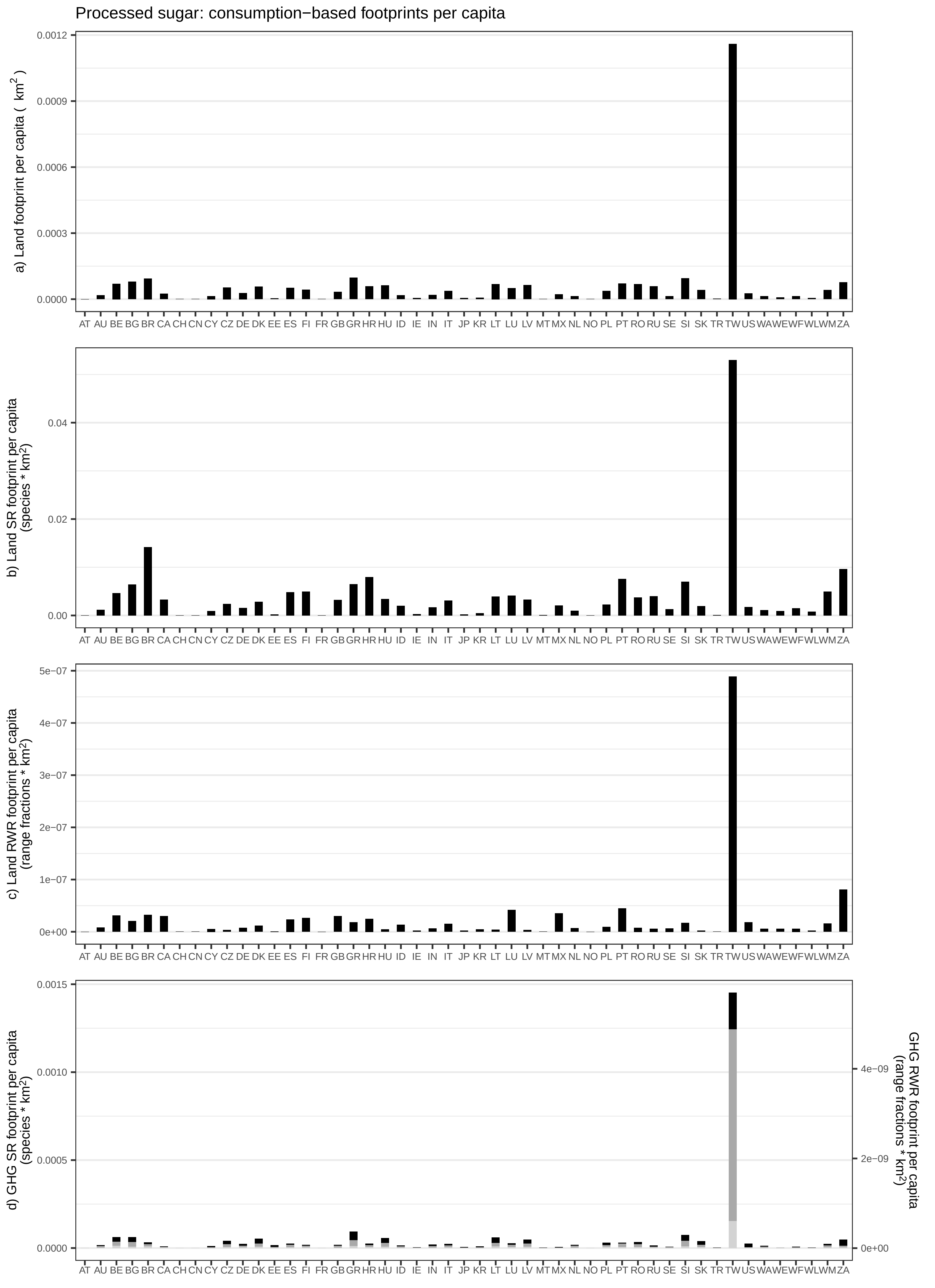

Processed sugar: consumption−based footprints per capita
a) Land footprint per capita ( km$^2$ )
b) Land SR footprint per capita (species * km$^2$)
c) Land RWR footprint per capita (range fractions * km$^2$)
d) GHG SR footprint per capita (species * km$^2$)
GHG RWR footprint per capita (range fractions * km$^2$)
AT AU BE BG BR CA CH CN CY CZ DE DK EE ES FI FR GB GR HR HU ID IE IN IT JP KR LT LU LV MT MX NL NO PL PT RO RU SE SI SK TR TW US WA WE WF WL WM ZA

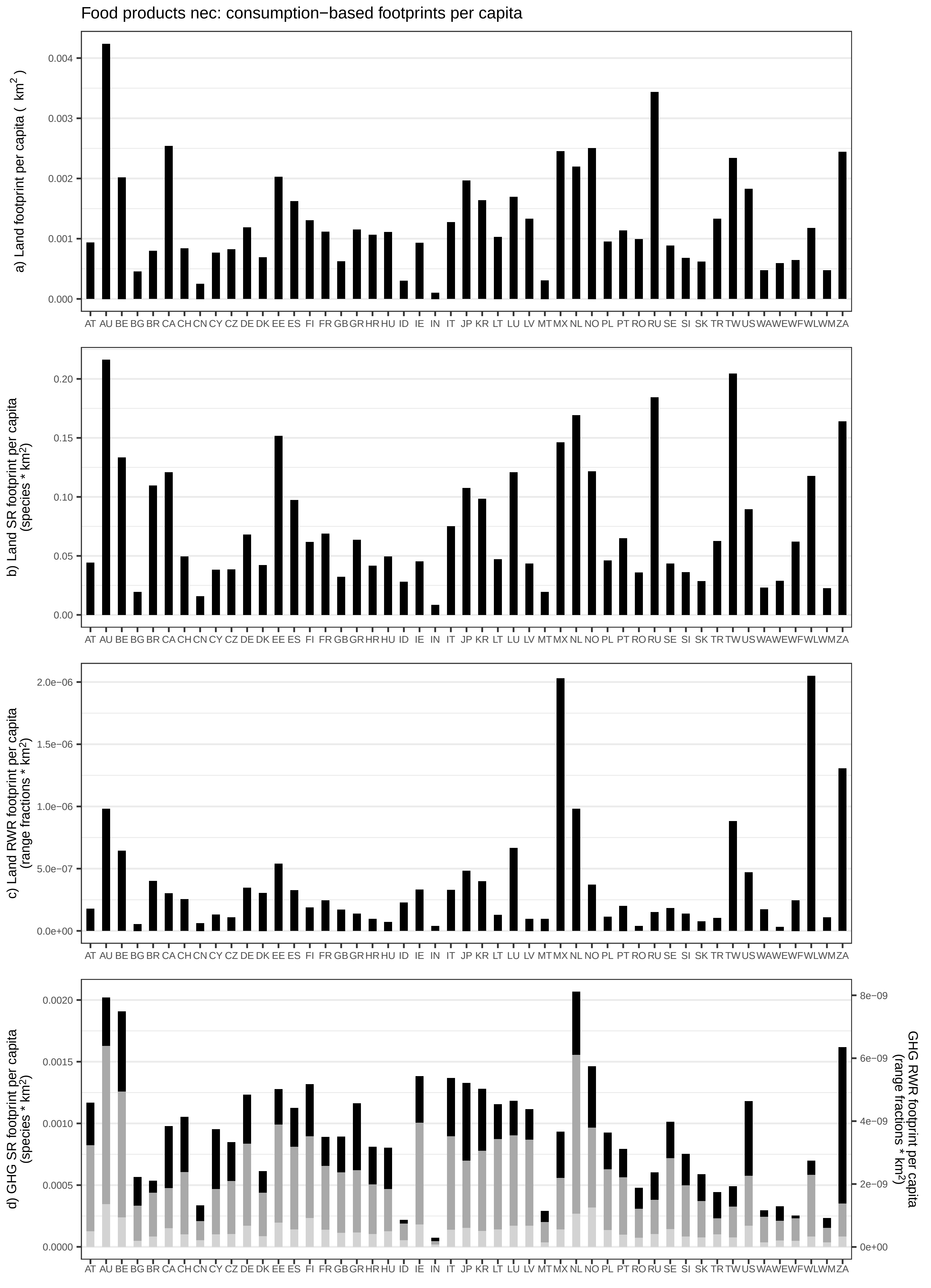
Food products nec: consumption−based footprints per capita
a) Land footprint per capita ( km$^2$ )
b) Land SR footprint per capita (species * km$^2$)
c) Land RWR footprint per capita (range fractions * km$^2$)
d) GHG SR footprint per capita (species * km$^2$)
GHG RWR footprint per capita (range fractions * km$^2$)
AT AU BE BG BR CA CH CN CY CZ DE DK EE ES FI FR GB GR HR HU ID IE IN IT JP KR LT LU LV MT MX NL NO PL PT RO RU SE SI SK TR TW US WA WE WF WL WM ZA

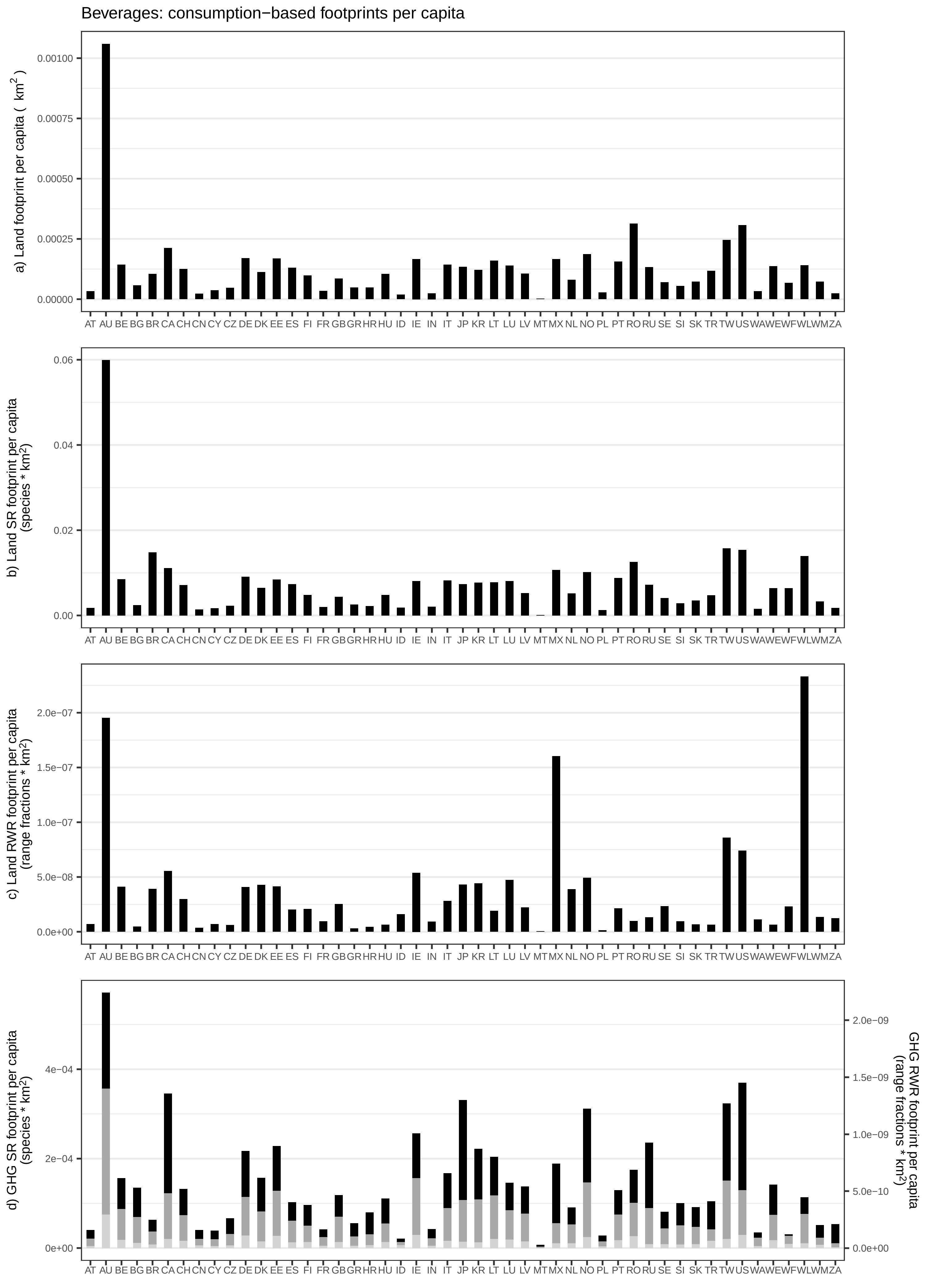
Beverages: consumption−based footprints per capita
a) Land footprint per capita ( km2 )
0.00100
0.00075
0.00050
0.00025
0.00000
b) Land SR footprint per capita (species * km2)
0.06
0.04
0.02
0.00
c) Land RWR footprint per capita (range fractions * km2)
2.0e−07
1.5e−07
1.0e−07
5.0e−08
0.0e+00
d) GHG SR footprint per capita (species * km2)
4e−04
2e−04
0e+00
GHG RWR footprint per capita (range fractions * km2)
2.0e−09
1.5e−09
1.0e−09
5.0e−10
0.0e+00
AT AU BE BG BR CA CH CN CY CZ DE DK EE ES FI FR GB GR HR HU ID IE IN IT JP KR LT LU LV MT MX NL NO PL PT RO RU SE SI SK TR TW US WA WE WF WL WM ZA

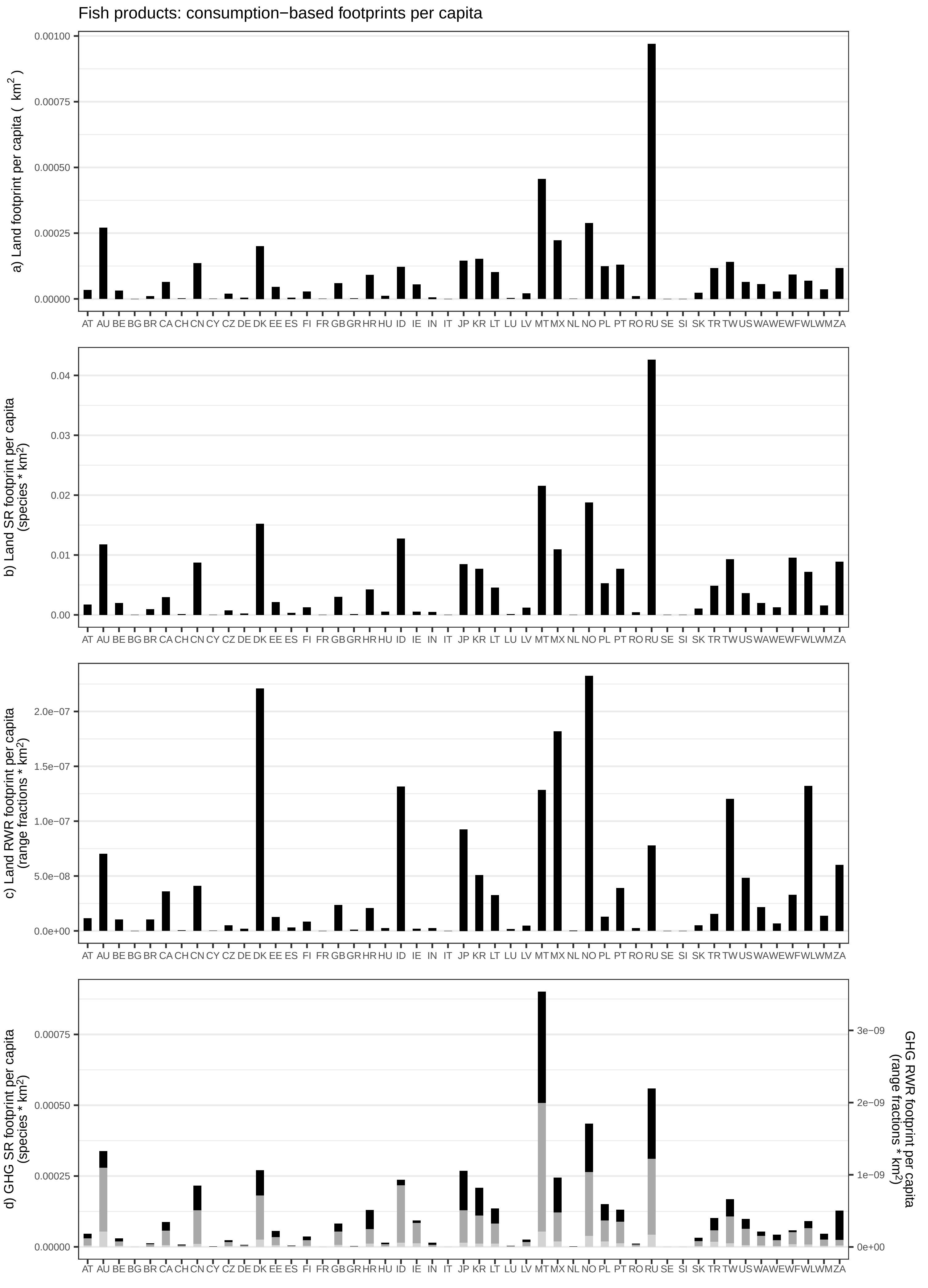

Fish products: consumption−based footprints per capita
a) Land footprint per capita ( km2 )
b) Land SR footprint per capita (species * km2)
c) Land RWR footprint per capita (range fractions * km2)
d) GHG SR footprint per capita (species * km2)
GHG RWR footprint per capita (range fractions * km2)
AT AU BE BG BR CA CH CN CY CZ DE DK EE ES FI FR GB GR HR HU ID IE IN IT JP KR LT LU LV MT MX NL NO PL PT RO RU SE SI SK TR TW US WA WE WF WL WM ZA

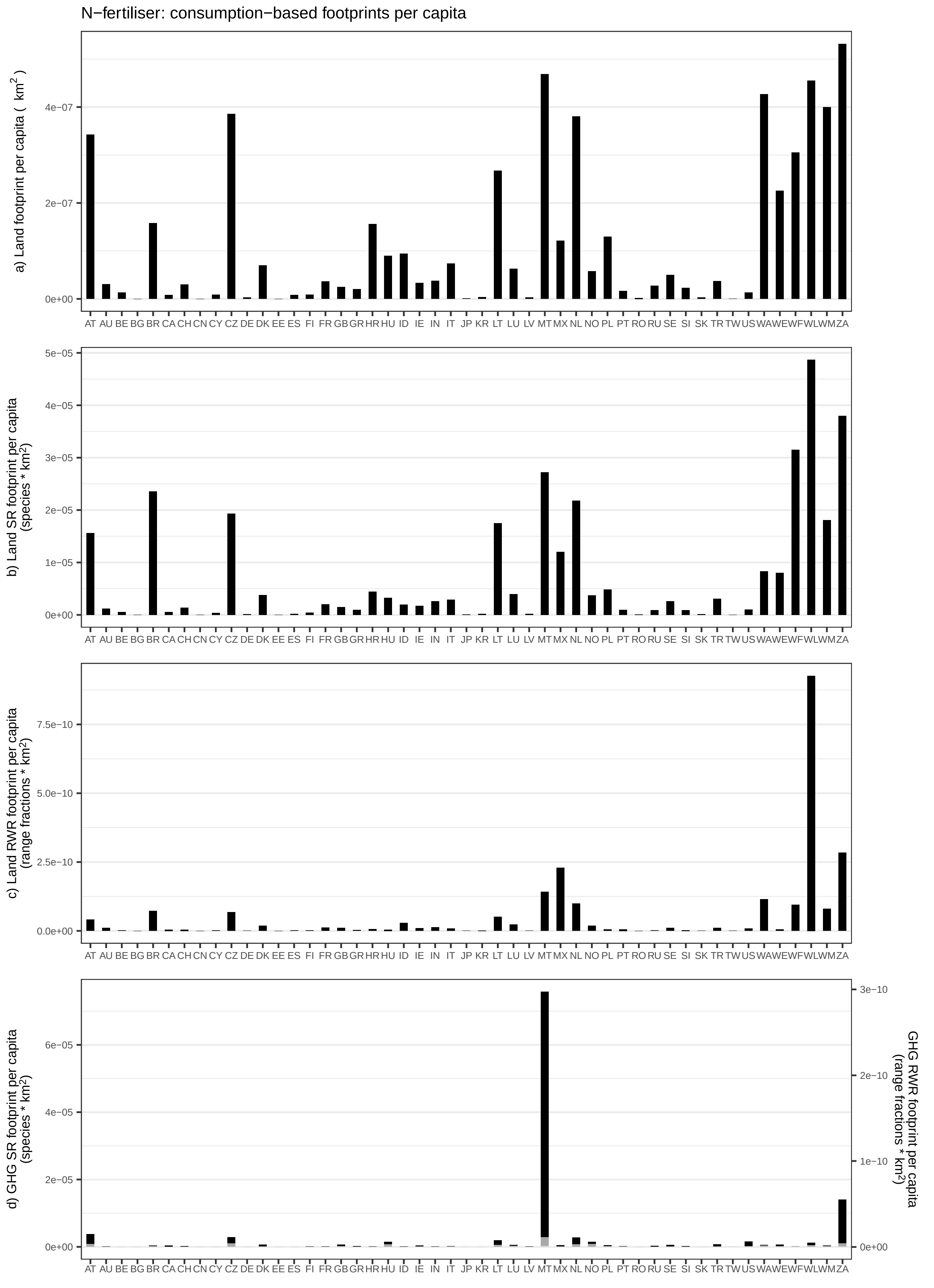

N−fertiliser: consumption−based footprints per capita
a) Land footprint per capita ( km2 )
4e−07
2e−07
0e+00
b) Land SR footprint per capita (species * km2)
5e−05
4e−05
3e−05
2e−05
1e−05
0e+00
c) Land RWR footprint per capita (range fractions * km2)
7.5e−10
5.0e−10
2.5e−10
0.0e+00
d) GHG SR footprint per capita (species * km2)
6e−05
4e−05
2e−05
0e+00
GHG RWR footprint per capita (range fractions * km2)
3e−10
2e−10
1e−10
0e+00
AT AU BE BG BR CA CH CN CY CZ DE DK EE ES FI FR GB GR HR HU ID IE IN IT JP KR LT LU LV MT MX NL NO PL PT RO RU SE SI SK TR TW US WA WE WF WL WM ZA

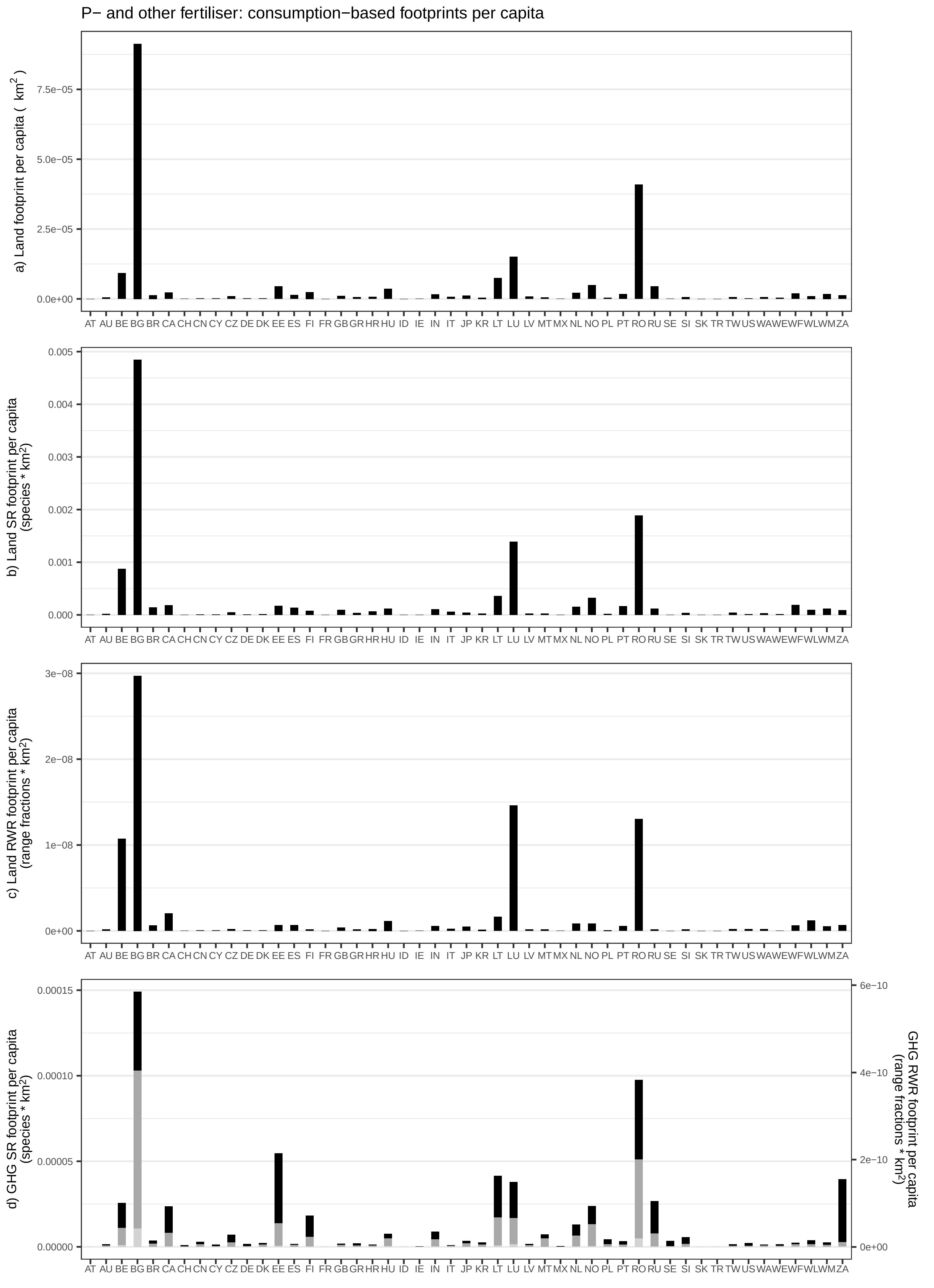

P− and other fertiliser: consumption−based footprints per capita
a) Land footprint per capita ( km2 )
7.5e−05
5.0e−05
2.5e−05
0.0e+00
b) Land SR footprint per capita (species * km2)
0.005
0.004
0.003
0.002
0.001
0.000
c) Land RWR footprint per capita (range fractions * km2)
3e−08
2e−08
1e−08
0e+00
d) GHG SR footprint per capita (species * km2)
0.00015
0.00010
0.00005
0.00000
GHG RWR footprint per capita (range fractions * km2)
6e−10
4e−10
2e−10
0e+00
AT AU BE BG BR CA CH CN CY CZ DE DK EE ES FI FR GB GR HR HU ID IE IN IT JP KR LT LU LV MT MX NL NO PL PT RO RU SE SI SK TR TW US WA WE WF WL WM ZA

Food waste: incineration: consumption−based footprints per capita

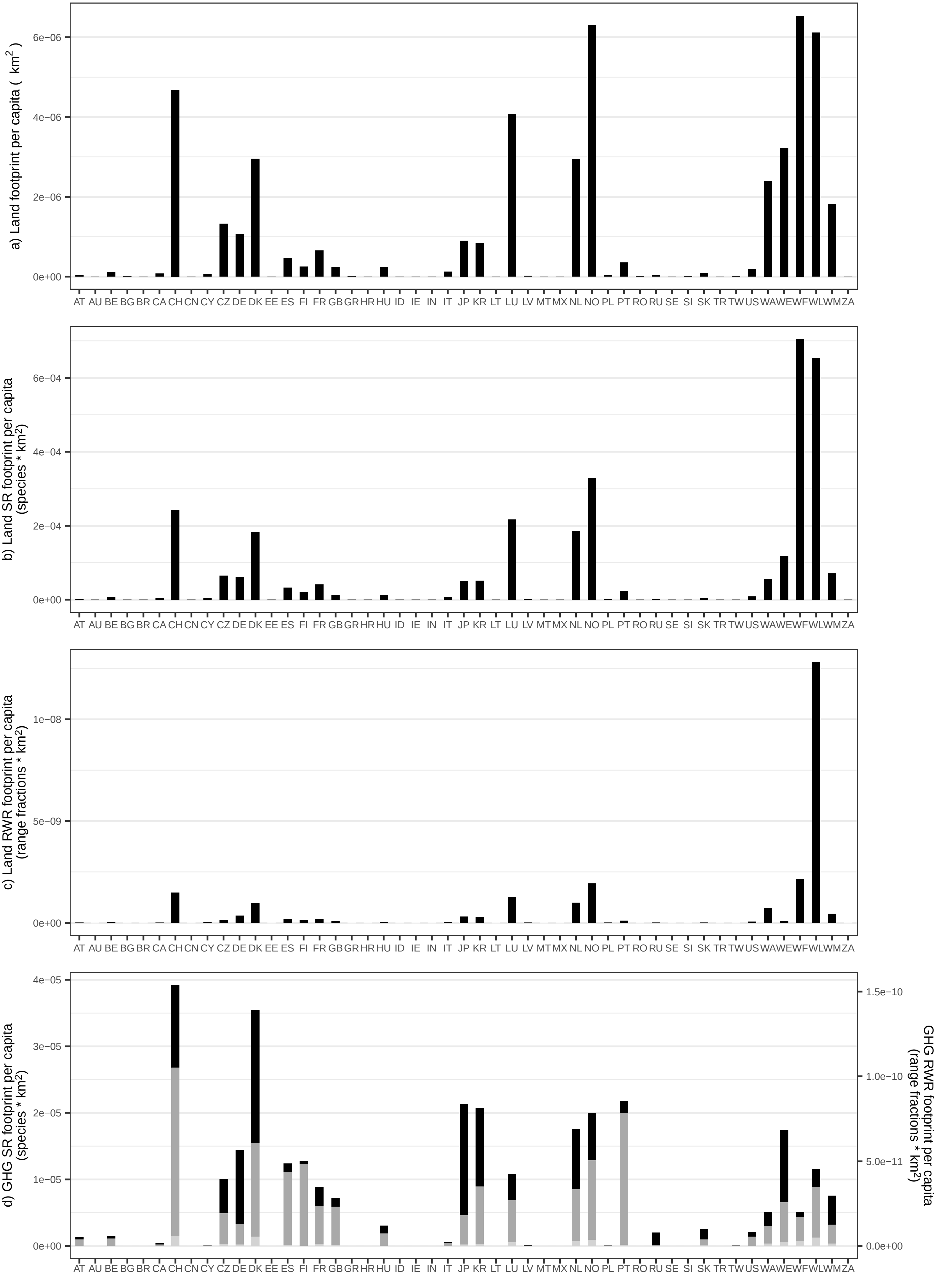

Food waste: biogasification and land application: consumption−based footprints per capita

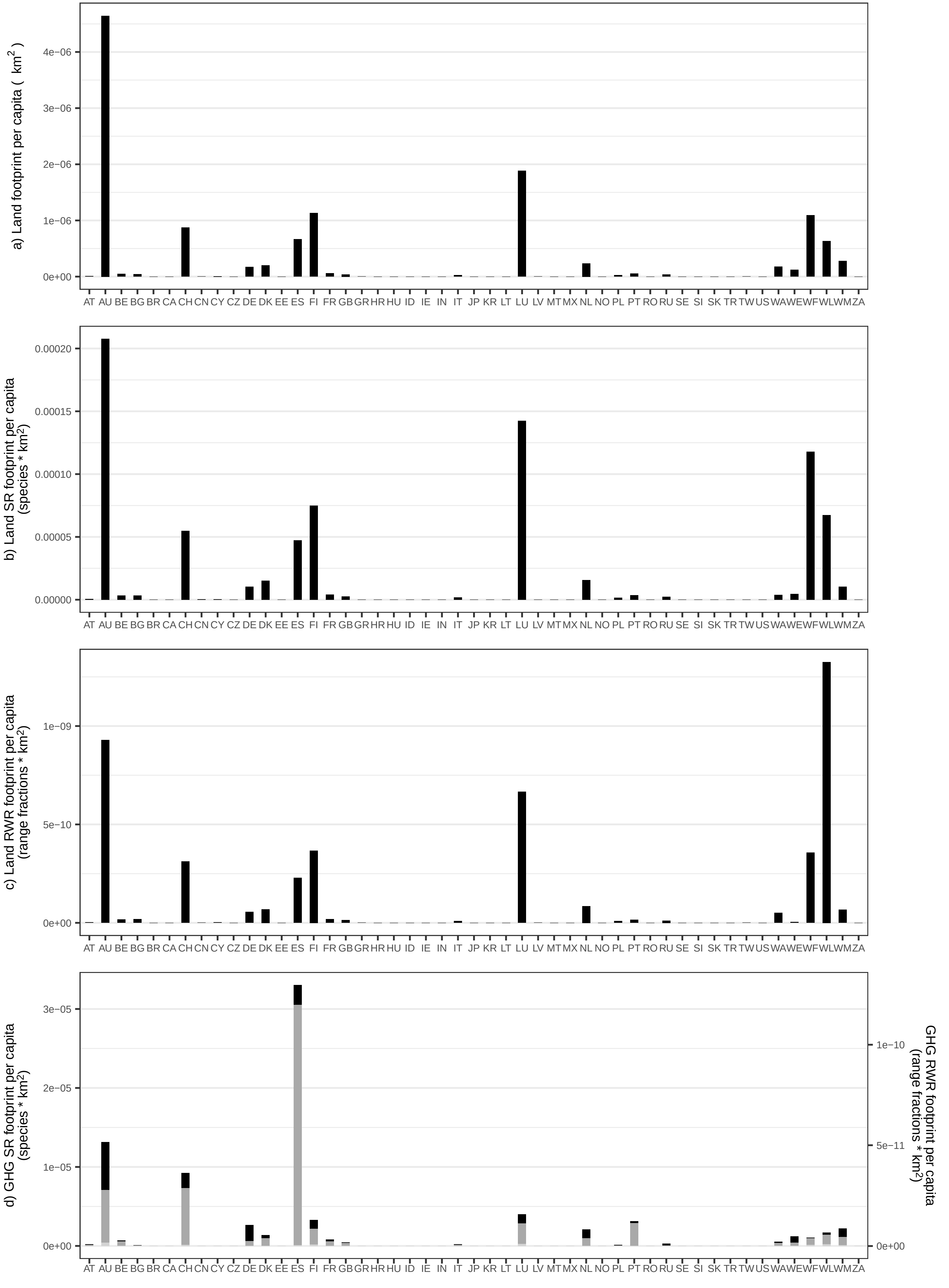

Food waste: composting and land application: consumption−based footprints per capita

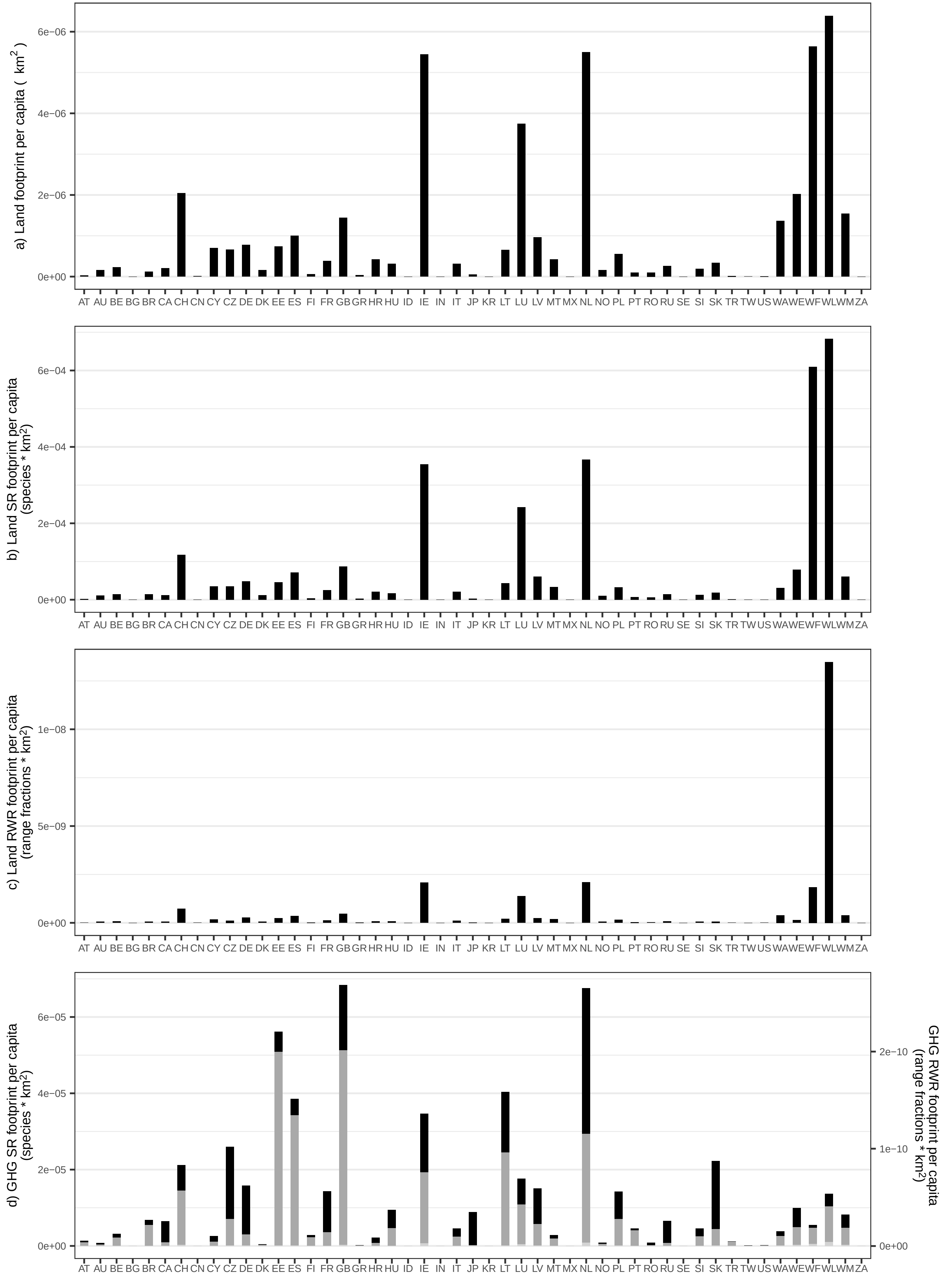

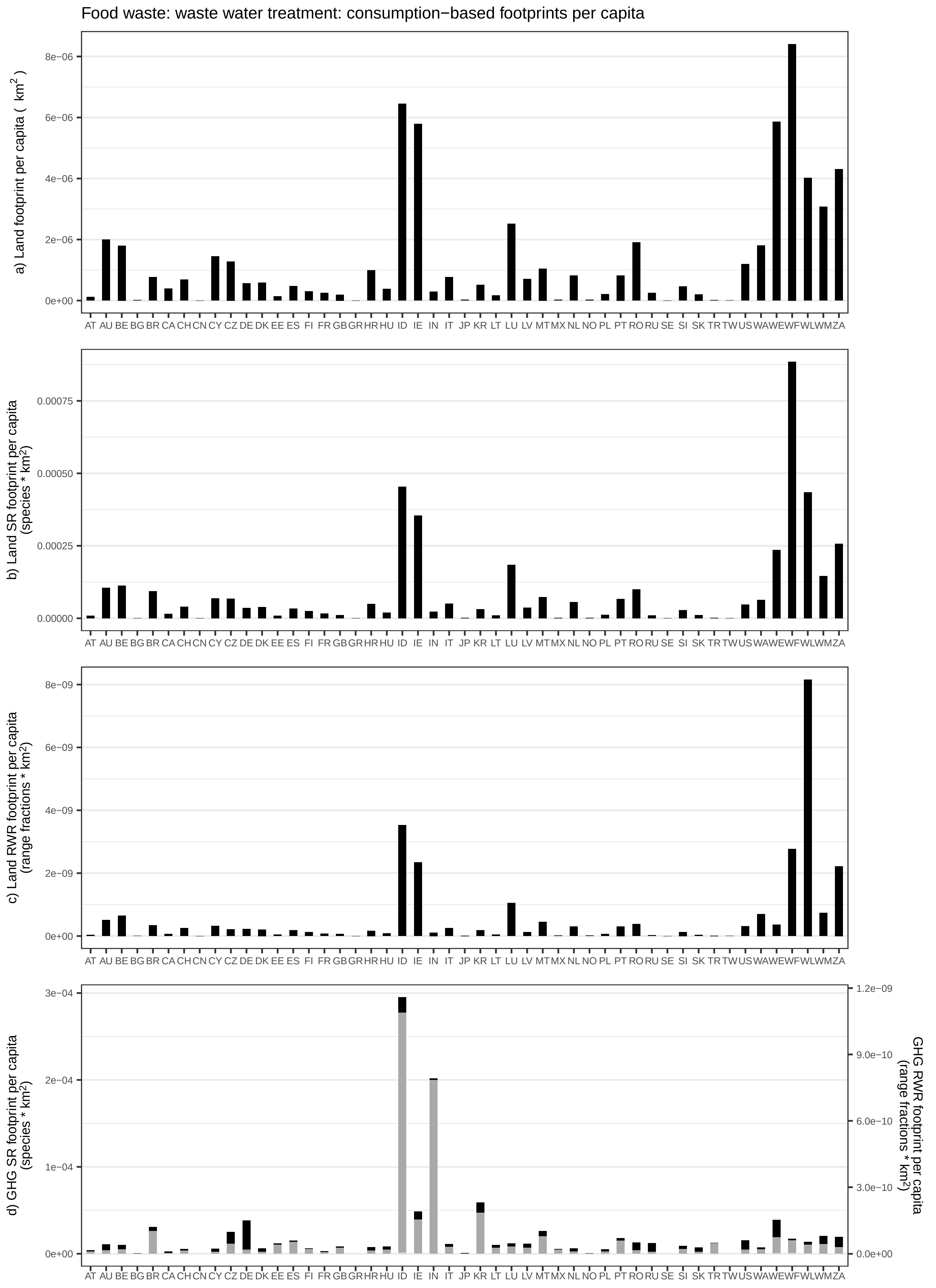
Food waste: waste water treatment: consumption−based footprints per capita
a) Land footprint per capita ( km$^2$ )
b) Land SR footprint per capita (species * km$^2$)
c) Land RWR footprint per capita (range fractions * km$^2$)
d) GHG SR footprint per capita (species * km$^2$)
GHG RWR footprint per capita (range fractions * km$^2$)
AT AU BE BG BR CA CH CN CY CZ DE DK EE ES FI FR GB GR HR HU ID IE IN IT JP KR LT LU LV MT MX NL NO PL PT RO RU SE SI SK TR TW US WA WE WF WL WM ZA

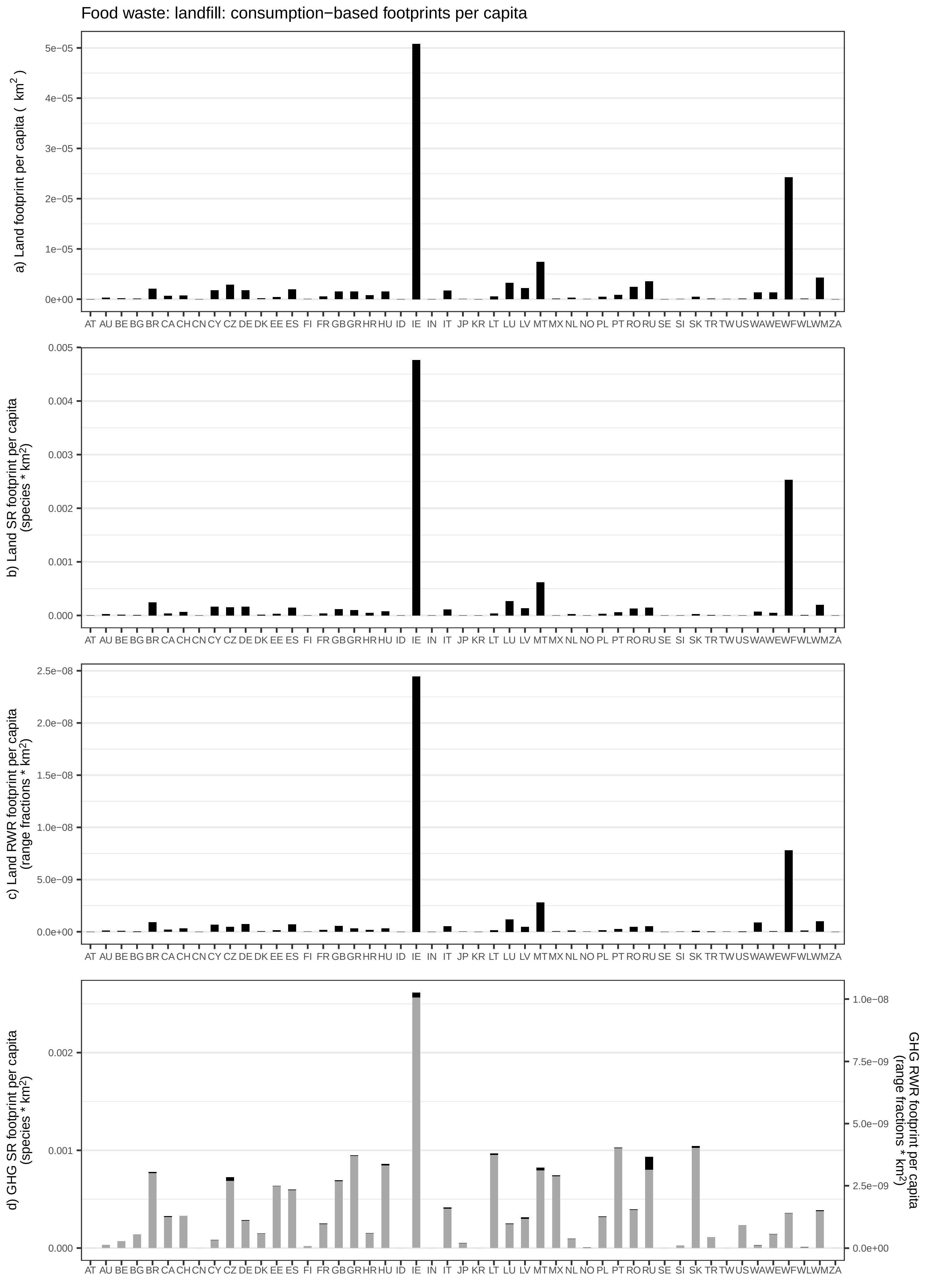

Food waste: landfill: consumption−based footprints per capita
a) Land footprint per capita ( km²)
b) Land SR footprint per capita (species * km²)
c) Land RWR footprint per capita (range fractions * km²)
d) GHG SR footprint per capita (species * km²)
GHG RWR footprint per capita (range fractions * km²)
AT AU BE BG BR CA CH CN CY CZ DE DK EE ES FI FR GB GR HR HU ID IE IN IT JP KR LT LU LV MT MX NL NO PL PT RO RU SE SI SK TR TW US WA WE WF WL WM ZA
